# High-throughput discovery of a [4+3] dearomative cycloaddition enables dual photochemical-photophysical perturbative probing of protein function

**DOI:** 10.1101/2025.10.23.684193

**Authors:** Samuel Griggs, Yibo Zeng, Stefan Lohmann, Chris Arter, Samuel Liver, Yana Demyanenko, Sivaramakrishnan Ramadurai, Georgina L. Bonney, Mariia Lopatniuk, Alasdair M. Mackenzie, Stanley Botchway, Andrew G. Leach, Graham J Hutchings, Shabaz Mohammed, Stephen P. Marsden, Ajay Jha, Adam Nelson, Benjamin G. Davis

**Affiliations:** Rosalind Franklin Institute, Harwell Campus, Didcot, OX11 0QX, UK; School of Chemistry, University of Leeds, Leeds, LS2 9JT, UK; Department of Pharmacology, University of Oxford, Oxford OX1 3QT, United Kingdom; UK Catalysis Hub, Harwell, Didcot, OX11 0FA, UK; Max Planck-Cardiff Centre on the Fundamentals of Heterogeneous Catalysis FUNCAT, Cardiff Catalysis Institute, School of Chemistry, Cardiff University, Translational Research Hub, Maindy Road, Cardiff, CF24 4HQ, UK; Central Laser Facility, Science and Technology Facilities Council, Rutherford Appleton Laboratory, Harwell, Didcot, OX11 0QX UK; Division of Pharmacy and Optometry, University of Manchester, Stopford Building, Manchester, M13 9PT UK; Department of Chemistry, University of Oxford, Oxford OX1 3TA, UK; Department of Biochemistry, University of Oxford, Oxford OX1 3QU, UK; Astbury Centre for Structural Molecular Biology, University of Leeds, Leeds, LS2 9JT, UK

## Abstract

Proteins exist in multiple conformations or states, often associated with different functional capacities. The ability to selectively identify and study these states is crucial for understanding the dynamic nature of proteins and their roles in cellular processes, yet can be currently limited by the artefacts of existing molecular labels and the lack of strategies to exploit them in diverse ways. Whilst i) methods for the photochemical *perturbation* of proteins and ii) photophysical protein probes and sensors are both known powerful tools, simultaneous methods for site– and context-specific photochemical *modification* of a protein side chain with concomitant modulation of photophysical properties would allow interrogation of protein function via diverse modes (potentially reporting simultaneously on structure and dynamics of ground– and reactive-state properties). Due to their mechanistic complexity, rational approaches for the design of suitable photochemical strategies can be difficult and non-intuitive. Here, we describe a high-throughput experimentation (HTE) strategy designed to allow the photochemical modification of a genetically-encoded, minimally disruptive (‘zero-size’), photophysically-active – yet typically unreactive and biologically-inert – unnatural amino acid (uAA) residue in proteins. The discovery of a novel dearomative, formal [4+3] photocycloaddition of imidazopyrimidine reagents which proceeds via photochemically generated C• intermediates with polyaromatic substrates enabled chemoselective application to the normally unreactive Trp-mimetic residue, 2-naphthylalanine (Npa). Npa’s site-selective installation allows dual photochemical and photophysical dearomative perturbation benignly and efficiently in proteins. Replacement of Trp in the archetypal apoptotic biosensor protein Annexin A5 (AnxV) enabled dual photochemical-photophysical determination of switched, conformationally-determined reactive states induced by Ca^2+^-binding, without any apparent functional perturbation of its *in vitro* or cellular recognition of phospholipids, indicating an ordered annexin mechanism. This ready and benign unification of photochemistry and photophysics suggests a general approach for using high-throughput light-mediated experimentation in artefact-free chemical biology to capture different functional states of proteins and to interrogate those states using combined photochemical-photophysical methods.

**Highlights:**

- Development of a general, high-throughput experimentation (HTE) approach enabled identification of an unprecedented photochemical reaction with potential for editing amino acid side chains within proteins.
- Genetic incorporation of 2-naphthylalanine, a minimally-disruptive (‘zero-size’) Trp mimic, enabled photochemical side chain editing with concomitant modulation of its distinctive photophysical properties in proteins.
- A ‘naphthalene-tagged’ variant of the apoptotic biomarker protein AnxV enabled dual photochemical-photophysical exploration of its dynamic conformational states induced by Ca^2+^-binding whilst fully maintaining its functional capacity for *in vitro* or cellular recognition of phospholipids.

## Introduction

Despite striking advances in observing and understanding static structures, the interrogation of the protein dynamics that determine function remains a key ongoing challenge.^1–5^ The probing of biology with photophysical methods, including sensing and perturbing, is now a dominant strategic approach. Fluorescence microscopies and spectroscopies, in conjunction with protein-specific labelling approaches, enable non-invasive protein monitoring, and even sensing,^6^ in living cells,^7^ typically through genetic fusion of fluorescent protein (FP) domains, and can be exploited in a photoactivatable manner.^8^ However, given the considerable size (>25 kDa) of FPs, it is clear^9^ that such adjuncts are likely to impair the usual function and distribution of target proteins and they cannot be obviously applied in a residue-specific manner. Alternative genetic approaches utilize the cell’s translational machinery to label proteins by incorporating, via stop-codon suppression, small molecule fluorophores as unnatural amino acids (uAA).^10^ However, the efficiency of incorporation can be limited (for most current variants^11–14^ to <2 mg/L of culture) and potentially hampered by degradation of precursor uAAs during biosynthesis, with a few important examples of higher incorporation efficiencies.^10,15,16^ The expansion of the range of introducible fluorescent uAAs has been explicitly identified as an ongoing challenge.^17^ Whilst there is one example of a fluorescent uAA that is photoactivatable^18^ (through C=S → C=O desulfurization), there is, to our knowledge, no fluorescent uAA that can be reactively probed, for example based on accessibility of a second ‘co-factor’, as a reporter of protein state.

We considered that a combination of light-mediated, genetic and chemical methods could prove powerful if a suitably *reactive* fluorescent uAA, and a corresponding chemoselective photochemical reaction to modify it, could be found. Applied strategically, this could provide opportunities not only for observing proteins, but also for capturing functionally-relevant conformational transitions based on reactive capacity. In this way, proteins engaging in functional interactions with other biomolecules, ligands or cofactors that induce local conformational changes might be discerned through the use of suitably fast reactions to trap specific conformational states and/or changes within a given protein of interest (**Figure 1**) – such ‘reactive trapping’ is a highlighted goal in protein dynamics.^5^ With suitable perturbation of an existing fluoro– or chromophore, the resulting chemical functionality might simultaneously also be investigated through photophysical probing of its altered discernible spectral characteristics. Here, through agnostic discovery of a *bis*-radical mediated reaction that then dearomatises a distinct fluorophore, we show that photochemical modification of proteins can drive switching between different observable photophysical states in a conformation-specific manner.

**Figure 1.**
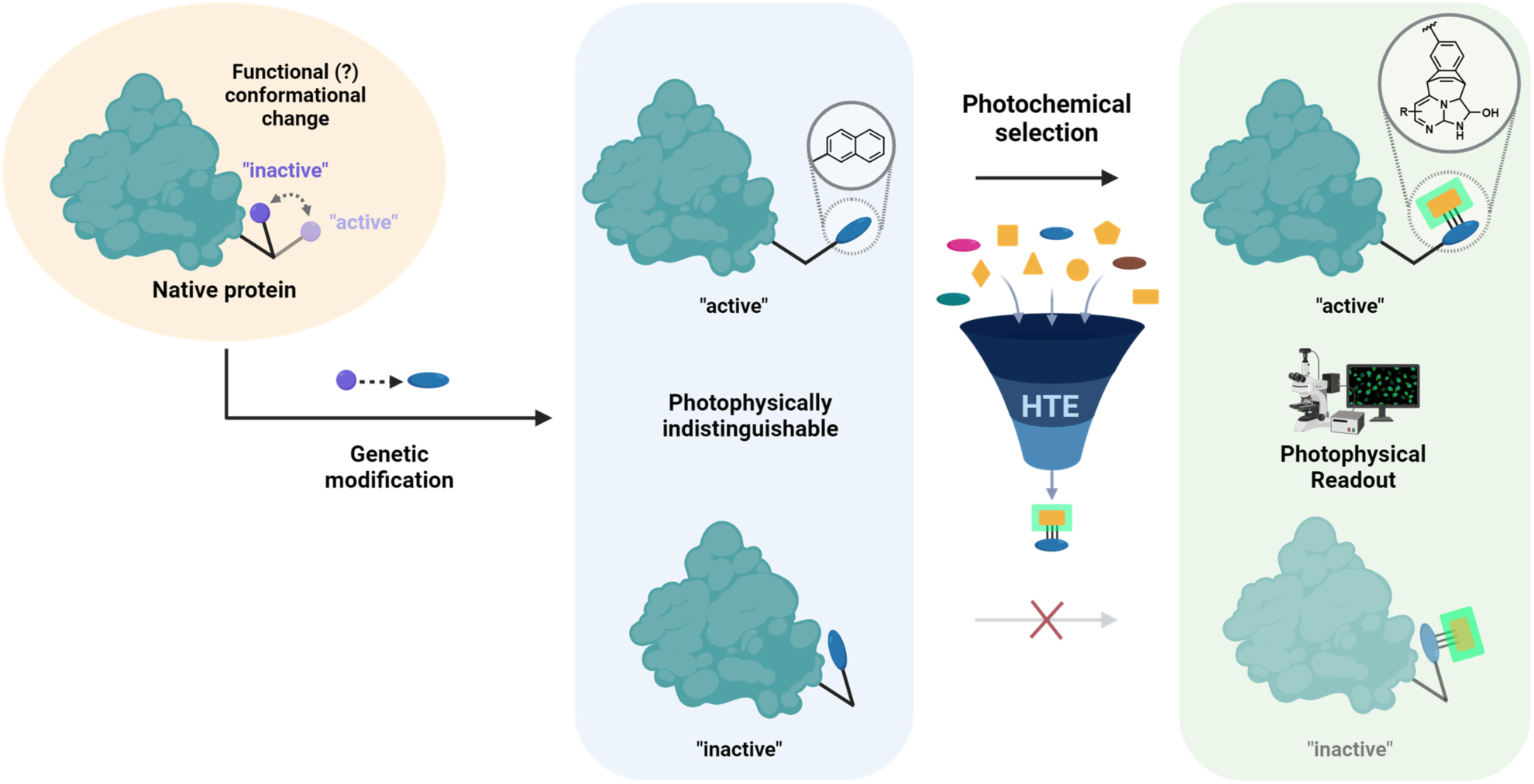
A Dual Photochemical-Photophysical Strategy for Probing of Protein Dynamics. Following site-selective genetic introduction of a suitably minimally perturbative (‘zero-size’) uAA, proteins may interact with other biomolecules, ligands or cofactors in a functional manner, leading to localized conformational changes. Specific conformational states might be reactively-trapped rapidly using selective *photochemical* probes, and the resulting chemically-modified uAA examined *photophysically* based on their distinct altered spectral characteristics. A suitable photochemical reaction capable of photophysical pertubation was discovered using a high throughput experimentation (HTE) approach.

## Results

### High-throughput Discovery of a Photochemical [4+3] Cycloaddition-Oxidation

Our strategy was reliant on the availability of a photochemical reaction that would chemoselectively modify the side chain of an introduced uAA ‘tag’^19^, resulting in perturbation of its photophysical properties. The development of photochemical methods for the synthetic modification of proteins is gaining in popularity.^20–22^ However, the *a priori* design of new photocatalytic reactions can be challenging due to mechanistic complexity, with the potential for the involvement of multiple excited states and the formation of alternative products. Consequently, high-throughput experimental approaches have been developed to drive the discovery of new small-molecule photochemical reactions.^23^ ^24^

We used here a high-throughput experimental approach to discover new photocatalytic reactions applicable to protein side-chain editing (**Figure 2**). We assembled a set of substrates (**Figure S1**) that were comparable in size to amino acid side chains (potentially bioisosteric), and that contained at least one functional group (e.g. hetarene, amine, aniline, alkene) with precedented reactivity in photochemical reactions (e.g. photoredox-catalysed dehydrogenative couplings^25,26^ and hydroaminations^27–30^). Tryptophan (Trp) contains a natural indolic fluorophore in its side chain. We recognised that reactions for modifying Trp-mimics might have particular value because Trp occurs at low frequency in proteins, is often found at key interfaces^31^ (e.g. with membranes and in ligand binding sites), and Trp-selective chemistries that modify its photophysical properties are rare.^32^ We therefore deliberately included in the substrate set not only modified indoles but also bicyclic aromatic compounds comparable in size to Trp’s indole side chain (near ‘zero-size’ perturbation).

**Figure 2.**
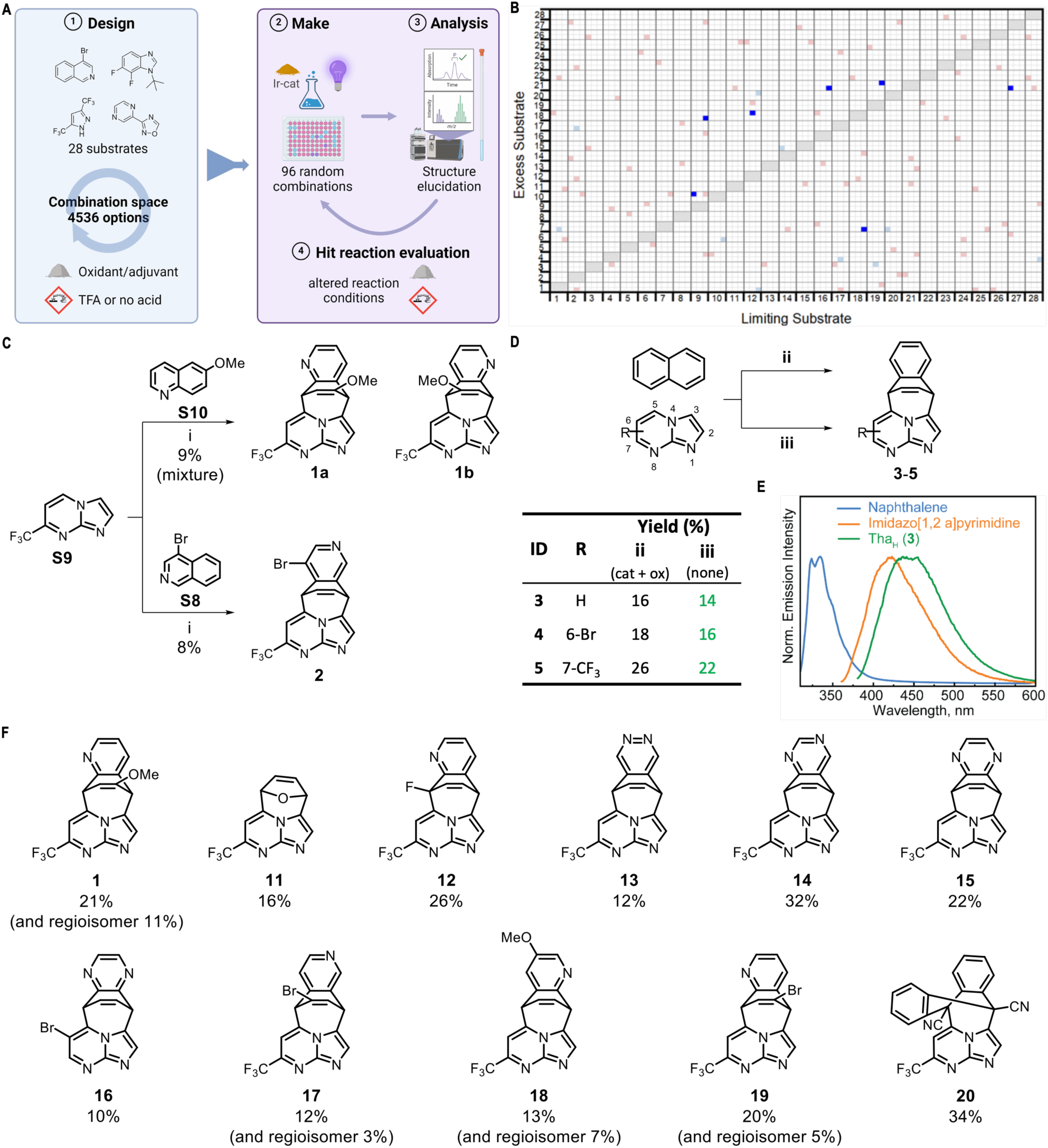
Discovery of a novel [4+3] cycloaddition-oxidation reaction. **A**) Overview of high-throughput experimental (HTE) approach. The virtual reaction space of 4536 possible combinations comprised all pairwise combinations of 28 substrates and six alternative reaction conditions (with TBPA, TRIP-thiol or no additive; with/without TFA, shown in each subarray). In each round, an array of 96 reactions was executed, and promising reaction products were purified by mass-directed HPLC and structurally elucidated. **B**) Overview of the virtual reaction space with limiting and excess substrate identities on the x– and y-axes, respectively. Reactions executed in the first round are highlighted with coloured squares (**light red**: unproductive combinations; **light blue**: productive combinations without validated/elucidated products; **dark blue**: productive reaction combinations with validated products). **C**) A novel [4+3] cycloaddition-oxidation was discovered in the first round (top scheme), which was also found to be successful with an alternative co-substrate in a second array (bottom scheme). This critically disrupted the chromophoric aromatic core (i: 2 mol% Ir[dF(CF_3_)ppy]_2_(dtbpy)PF_6_, TBPA, MeCN, 365 nm, 16 h). **D**) Further investigation of co-substrate scope revealed that the reaction was successful even with naphthalene as limiting substrate. Notably, this reaction proceeded without added oxidant or catalyst, suggesting enhanced feasibility for application in protein modification. Conditions: ii: TBPA (2 eq.), 2 mol% Ir(ppy)_3_, 4:1 MeCN−H_2_O, 365 nm, 16 h; iii: DMSO, 365 nm, 16 h. **E**) Comparison of the fluorescence emission spectra of the reactants naphthalene (**blue**), imidazo[1,2-*a*]pyrimidine (**orange**) and the reaction product **3** (**green**) revealed clear wanted photophysical pertubation. The spectra have been normalised to their respective peak maxima. **F)** Structures and yields of cycloaddition-oxidation products generated in scoping experiments (utilizing conditions ii, see above).

**Figure 3.**
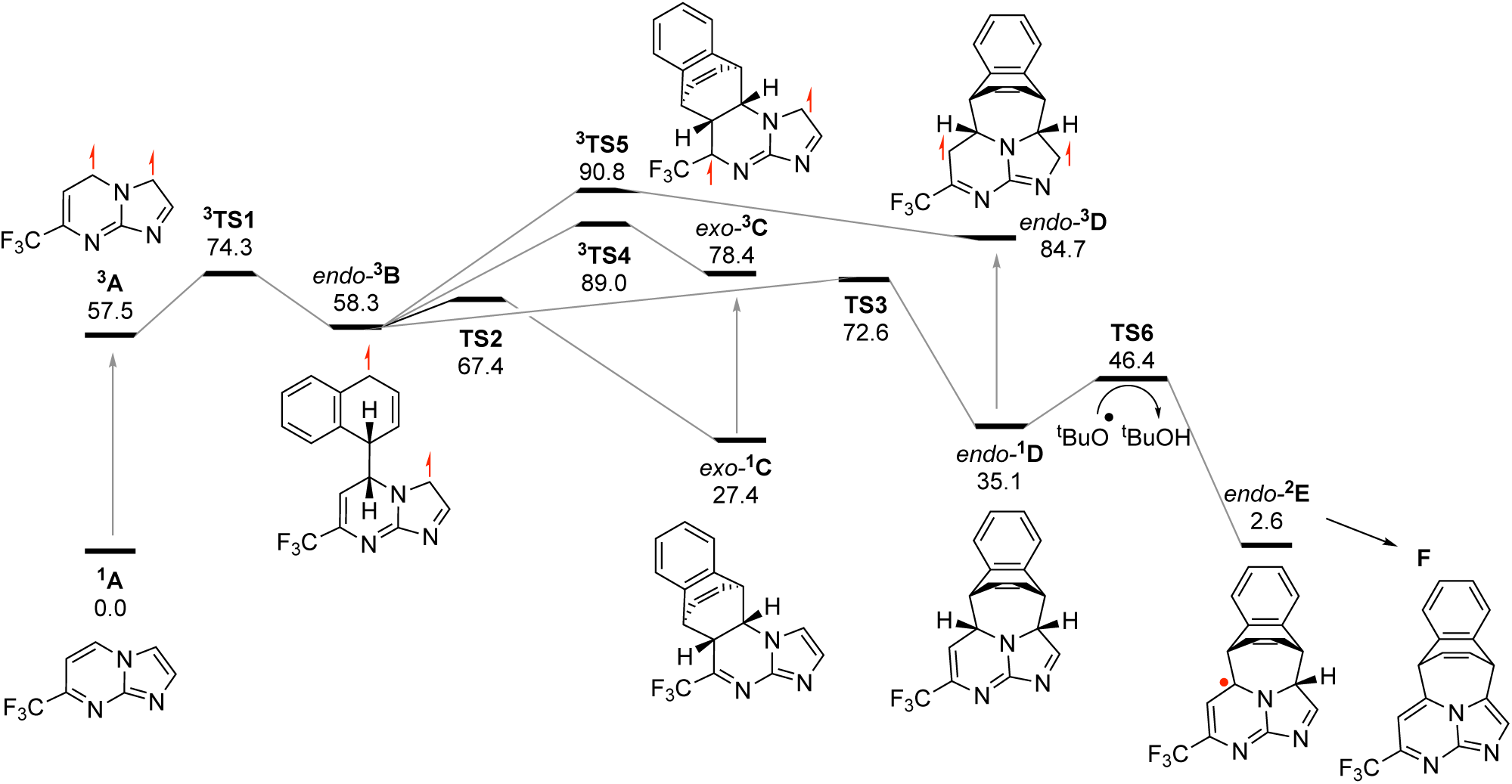
Calculated Mechanism of *bis*-Radical 4+3 photochemical Cycloaddition. Free energies (in kcal.mol^−1^ for 1M standard state at 298K) relative to singlet ground states of 7-trifluoromethyl imidazo[1,2-*a*]pyrimidine **S9** (^1^A) and naphthalene for their reaction computed at the M06-2X/def2-TZVP level including IEFPCM toluene. Superscripts correspond to spin states.

For possible ultimate application in reactive trapping, an emergent photochemical protein modification reaction would take place between the side chain of an introduced uAA and an external reagent. Each explored reaction combination therefore involved a pair of the substrates (one in threefold excess) and a photocatalyst (initially Ir[dF(CF_3_)ppy]_2_(dtbpy)PF_6_) ^33^ under one of six reaction conditions (**Figure 2A**). Initially, we randomly selected 96 reactions (**Figure 2B**) from 4536 possible permutations (28 substrates x 27 co-substrates in excess x 6 reaction conditions; see **Methods** for full details of the experimental conditions). The success of the reactions was assessed by analytical UPLC-MS, together with evaporative light scattering (ELS) detection to enable product quantification.^34^ Reactions were deemed promising if any product with mass ≥30 Da higher than either substrate was produced in >8 (±2) % yield. Such products were then isolated by preparative mass-directed HPLC and structurally elucidated using 500 MHz ^1^H NMR spectroscopy.

In total, five distinct product families were successfully identified (**Figure 2B** and **Figure S2**) from seven of the reactions. Three included homodimerization of the co-substrate (e.g. **S21**) via [2+2] cycloaddition without incorporation of the (varied) limiting substrate (→ **8**, **Figure S2**) and direct addition with thiol adjuvant (e.g. for vinyl pyrazine **S18**) rather than with the co-substrate (→ **7**, **Figure S2**). **S18** also underwent hydroamination with **S10** (→ **6**) and **S7** (→ **9**) (**Figure S2**). The most intriguing and promising product, however, stemmed from reaction between quinoline **S10** and the imidazo[1,2-*a*]pyrimidine **S9** to give two regioisomeric products (**1a** and **1b**, **Figure 2C**). Notably, these arose from a dearomative [4+3] cycloaddition−oxidation (see **Figure 2C**). Crucially, consistent with the intended design, we considered that such a dearomative reaction would likely strongly perturb the photophysical properties of this, or possibly similar, functionality incorporated into a uAA side chain. Variation of the reaction in the same microscale format (**Table S1**) allowed estimation of initial efficiency through larger scale isolation (**Figure 2F**, **S9**+**S10**, 2 mol% Ir(ppy)_3_, 2 eq. TBPA, 4:1 MeCN:H_2_O, 365 nm, 16 h, regioisomeric products **1a** and **1b** in 21% and 11% yield, respectively).

Informed by these validated reactions from the first discovery round, a second reaction array was then iteratively designed (**Supplementary Notes**, Figure SN1.2 and Table SN1.2). The validated reactions were adjusted in three ways: (a) swapping the limiting and excess substrates; (b) exchanging either substrate with a related substrate from the initial set (e.g. hetarene for another hetarene); or (c) exploring other sub-array reaction conditions. A consequently expanded range of validated reactions were thus identified in the second round (**Figure S2**), including further [4+3] cycloaddition−oxidation reaction of isoquinoline **S8** (in place of the quinoline **S10**) leading to the formation of **2** (**Figure 2C**). Crucially, this suggested significant possible latitude in the choice of substrate, and so eventual uAA that might be modified.

### Photochemical [4+3] Reaction has Broader Scope

We reasoned that in our design this discovered [4+3] reaction could correspond to an [aromatic sidechain + reagent] reaction applied to proteins in which a photophysical active 4π electron ‘diene’ moiety is altered. This cycloaddition-oxidation proved successful with a range of ‘diene’ components, including several other substituted quinolines, 9,10-dicyanoanthracene, furan, phthalazine, and quinazoline (**Figure 2F**). Encouraged by this and the observation in particular that both quinoline and an isoquinoline were viable co-substrates, we investigated the viability even of naphthalene (‘de-aza-quinoline’) as a substrate. Napthylalanine (Npa) and analogues display photophysical uniqueness due to the presence of distinctive fine structure in fluorescence emission, which is a manifestation of both Franck–Condon and vibronic effects,^35^ offering the potential opportunity to differentiate Npa even amidst the overlapping complex spectral landscape of other aromatic amino acids that might be present in proteins. Npa has little to no reactivity under normal biological conditions, or indeed in aqueous solution, using classical chemistries, making it an ideal masked, condition–responsive ‘tag’ amino acid. To our knowledge, Npa has not been used before as a chemically-reactive group in proteins.^19^ Pleasingly, under the optimal conditions, a corresponding [4+3] product was obtained when naphthalene was used in excess (26% yield) and as the limiting component (14% yield) (**Figure 2D**). Notably, the reaction was also successful without either the photocatalyst or the oxidant – this further suggested that it may ultimately be possible to identify even simpler conditions suitable for protein modification that would avoid their potentially damaging^36^ role.

Next, we examined the initial scope of the imidazo[1,2-*a*]pyrimidines that might be used as reagents to address naphthalene as a target sidechain. Besides the initially discovered 7-trifluoromethyl-substituted substrate (**S9**), 6-bromo– and unsubstituted imidazopyrimidine also reacted with naphthalene to give the expected cycloadducts, albeit in reduced yield (**Figure 2D**). With 6-bromo-imidazopyrimidine, a small amount of debrominated adduct was also observed in reactions with naphthalene and quinazoline (**Figures 2D** and **2F**).

### Mechanism

To gain an insight into the mechanism of the [4+3] cycloaddition-oxidation, we performed density functional theory (DFT) calculations on the reaction between naphthalene and the imidazopyrimidine **S9** (M06-2X/def2-TZVP level of theory^37–41^, including SMD solvation^42,43^). These suggested a putative mechanism in which, initially, triplet **^3^A** is generated, either by energy transfer from an excited photosensitiser, or by direct excitation to the first excited singlet state, followed by inter-system crossing. Consistent with these calculations, steady-state and time-resolved spectroscopic measurements revealed that imidazopyrimidine forms aggregates at concentrations used for the reaction, showing new optical transitions and weak, red-shifted emissions (**Figure S4**). These aggregates may facilitate the [4+3] cycloaddition by tuning the singlet-triplet energy gap highlighted by calculations. Additional titration experiments with naphthalene showed no ground state complexation (**Figure S5**), indicating diffusion-mediated energy transfer from excited imidazopyrimidine to naphthalene, which necessitates the role of long-lived triplet states.

Calculations further suggested that **^3^A** then reacts readily with naphthalene, essentially by C• radical addition, at either the 3– or 5-position of the imidazopyrimidine ring: the most favourable pathway proceeds via transition state **^3^TS1** to yield *endo-***^3^B**. Notably, the mechanism of a recently-reported *peri*-[3+2] cycloaddition between quinolones and alkynes is also proposed to be stepwise, and to be initiated by triplet energy transfer.^44^ Cyclisation of *endo-***^3^B** (or of one of the three products of alternative pathways, see **Supporting Information Figure SN2.3**) can yield either a [4+2] (*exo*-**^1^C** *via* **TS2**) or a [4+3] cycloadduct (*endo-***^1^D** *via* **TS3**). Whilst the formation of the [4+2] adducts is kinetically and thermodynamically favoured, the cycloadducts (*exo-***^1^C** and *endo-***^1^D**) can ring-open *via* re-excitation to triplet intermediates (e.g. *exo-***^3^C** or *endo-***^3^D**) and, thus, revert to *endo*-**^3^B** (via **TS4** and **TS5** respectively). Crucially, the calculations indicate that re-excitation occurs significantly more readily than excitation of **^1^A** and, hence, the initially favoured species would be rapidly generated and destroyed. The overall reaction outcome is therefore determined by the kinetically-controlled abstraction of a hydrogen atom from the alternative cycloadducts (with alkoxyl radical a likely proficient agent) *via* relatively low lying **TS6**. In this way, the most favourable [4+3] pathway is seemingly determined by the hydrogen abstraction from *endo-* **^1^D** to give *endo-***^2^E**, and, hence, the observed product **F** (**5**).

### Naphthalene and [4+3] Adducts Display Distinct Spectral Properties

Having established the *photochemical* feasibility of this novel [4+3] cycloaddition, we next probed the *photophysical* perturbation of naphthalene when converted to its corresponding [4+3] cycloadducts. Comparison of the steady state absorption spectra of naphthalene and its [4+3] products showed clear spectral changes induced upon cycloaddition with the conversion of naphthalene from vibronically-split bands (λ < 300 nm) to notably intense absorption bands centered at ∼338 nm (in **5**), ∼339 nm (in **4**) and ∼329 nm (in **3**) (**Figure S5, S6**). The corresponding fluorescence emission measurements also showed clear perturbation and revealed markedly red-shifted emission bands with λ_max_ ∼440 nm for **3**-**5** (**Figure 2E and S6**) compared to naphthalene (λ_max_ ∼334 nm, **Figure 2E**). Finally, napthalene exhibited a long fluorescence lifetime (>10 ns, **Figure S7**) that was also significantly perturbed upon cycloaddition to a regime ≤ 4ns. Together these observations of three significant *photophysical* perturbations (excitation, emission and lifetime), as observed by fluorescence, upon dearomative [4+3] *photochemical* addition suggested promise for their designed implementation in proteins.

### Discovered [4+3] Cycloaddition Allows Dual Site-selective Photochemical-Photophysical Modification-Perturbation in Proteins

Next, we investigated the incorporation of Npa into a model protein, as both a minimal naphthalene-bearing uAA and potentially close-mimetic of Trp, and its subsequent photochemical perturbation. One powerful mode of incorporation of uAAs into proteins relies on the use of orthogonal aminoacyl-tRNA synthetases (aaRS) and their corresponding supressing tRNAs from different origins.^45–50^ A number of unique aromatic (including some polycyclic) uAAs, including Npa,^47,51,52^ have been introduced into proteins using either paired tyrosyl amber suppressor tRNA_CUA_^Tyr^ and cognate tyrosyl-tRNA synthetases (TyrRS) or corresponding pyrrolysyl tRNA_CUA_^Pyl^-PylRS pairs where the aaRS synthetases have been made active and, ideally, highly selective for charging of the chosen uAA after directed evolution.^47^ In our hands, screening suggested limits on the utility of currently evolved tRNA_CUA_^Pyl^-PylRS pairs for Npa incorporation^52^ and we failed to see useful incorporation (see **Supplementary Information** for details). However, the *Mj* tRNA_CUA_^Tyr^-SS12TyrRS pair^47,51^ showed clear promise. We therefore enhanced its efficiency to allow extension of its applicability through the construction of suitable plasmids for use in multi-open reading frame (ORF), single vector systems (so-called pEVOL^53^). Co-transformation of a suitable histone *eh3 gene* (encoding an amber TAG codon at position 9) together with generation of a novel pEVOL-SS12-TyrRS/tRNA^Tyr^ plasmid allowed highly efficient expression (41.5 mg/L of culture) in *E. coli* of a full-length epitope (HA-FLAG-C-terminus)-tagged human histone H3 protein (eH3) bearing Npa at site 9 – eH3-Npa9 – in a manner dependent on the addition of 2 mM Npa to cell culture (**Figure 4B,C**). LC-MS, MS-MS and Western blot analyses confirmed incorporation of Npa at site 9 with essentially no background suppression by endogenous amino acids (**Figure 4A, step 1**). Importantly, the presence of Npa and its potential in photophysical methods were further confirmed by fluorescence emission spectra (**Figure 4D, blue**) that displayed the same characteristic vibronic emission features observed in small molecule model systems, setting the stage for designed perturbative probing.

**Figure 4.**
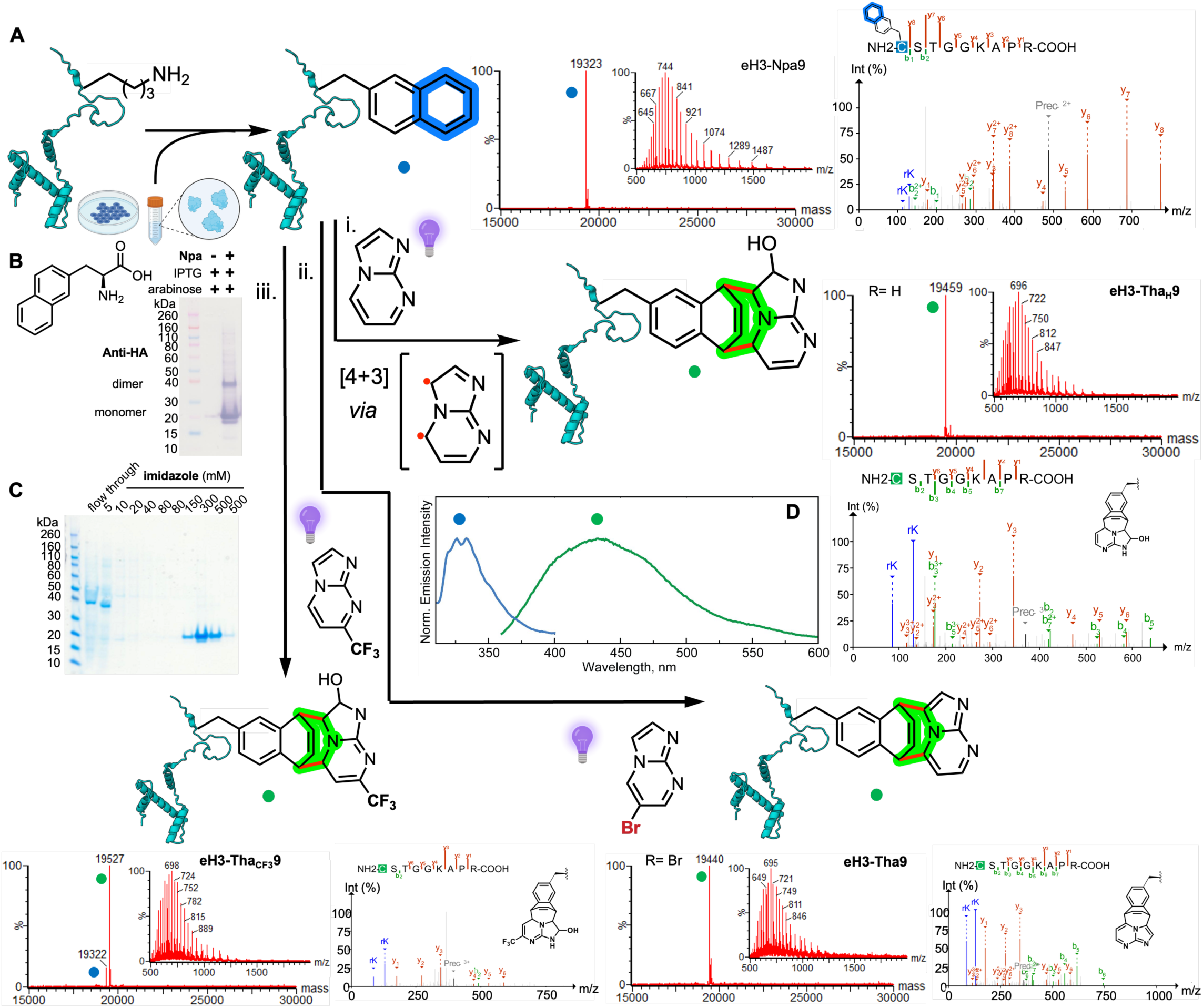
Insertion and dearomative [4+3] photochemical perturbation of Npa in proteins. **A-D**) Efficient site-selective incorporation and use of Npa into eH3 protein at position 9 of histone protein eH3 (**A step 1**) relied upon use of a bespoke pEVOL-SS12-TyrRS/tRNA^Tyr^ plasmid system encoding the SS12-TyrRS and tRNATyr pair to effectively suppress an amber codon at site in the *eH3* gene sequence in an Npa-dependent manner (see **B**, Western blot anti-HA) giving good yields of homogenous eH3-Npa9 (see **C**, Coomassie-stained SDS-PAGE). Intact protein MS (ion series and deconvoluted signal) revealed excellent incorporation efficiency (>95%). The identity of Npa and site of modification were further confirmed by LC-MSMS (using higher-energy collision dissociation (HCD)). **A step 2i)**. [4+3] photocycloaddition reaction of eH3-Npa9 with imidazole[1,2-*a*]pyrimidine yields protein tetrahydroazepine (Tha) product eH3-Tha_H_9 as a hydrated cycloaddition product, as indicated by the deconvoluted signals of intact protein mass spectra (middle panel) and LC-MSMS (right panel). Reactions were performed under argon atmosphere in denaturing buffer (500 mM ammonium acetate, 3 M guanidine hydrochloride, pH 6.2, 2% residual DMSO) without any photocatalyst or oxidant. **A step 2iii)**. Reaction with 7-CF_3_ imidazole[1,2-*a*]pyrimidine yields the corresponding hydrated eH3-Tha_CF3_9 [4+3] cycloadduct. **A step 2ii)**. Reaction with 6-bromo imidazo[1,2-*a*]pyrimidine allows elimination via dehydrobromination to yield rearomatized unsubstituted protein [4+3] cycloadduct eH3-Tha9. The intended site of all modifications was again confirmed by respective LC-MSMS (HCD). **D**) Photochemical modification leads to clear photophysical perturbation. The emission spectrum of eH3-Npa9 after excitation at 280 nm (**blue**) displays the characteristic vibronic features of naphthalene, thereby confirming the successful incorporation of Npa into the histone protein. The emission spectrum of the cycloaddition product on the protein, eH3-Tha_H_9 (**green**) obtained after excitation at 350 nm is also shown. These showed the presence of a strong, broad emission band centered at 435 nm, essentially identical to those found in small-molecule model photophysical systems (see **Figure 2**). Thus, together photophysical perturbation alters λ_abs_, λ_em_, vibronic fine structure and τ.

Following efficient generation of eH3-Npa9 bearing a site-selectively installed napthyl ‘tag’ site, we next tested it as a model protein substrate for [4+3] reaction. Successful on protein reaction of eH3-Npa9 with unsubstituted imidazopyrimidine to give eH3-Tha_H_9 bearing the corresponding tetrahydroazepinyl (Tha)-cycloadduct sidechain without photocatalyst immediately confirmed the utility of our high-throughput experimental approach involving model ‘side-chain’ substrates (**Figure 4A, step 2**). With focus on complete and rapid conversion of the Npa ‘tag’ to a core-disrupted dearomatized adduct, the scope of the reaction was initially explored under conditions favoring ‘tag’ accessibility (50 µM protein, 500 mM ammonium acetate, 3 M guanidine-HCl, pH 6.2). Corresponding cycloadducts were also observed for different imidazopyrimidine reagents (100 equiv., 5 mM) with >90% conversion (**Figure 4A, step 2i,ii,iii** and **Figure S8**). Notably, under these aqueous conditions, both un– and 7-CF_3_-substituted imidazopyrimidine reagents, gave eH3-Tha_H_ and eH3-Tha_CF3_ in their hydrated forms, consistent with trapping of the initial cycloadduct (which is an imine) rather than oxidation (**Figure 4A, step 2i,iii**). This again paralleled our observations with small molecules where non-hydrate cycloadducts were formed in 4:1 acetonitrile−water, whereas hydrates were formed in buffer (500 mM ammonium acetate, 3 M guanidine hydrochloride, pH 6.2, administering substrates via DMSO stock solutions). Characterization by fluorescence spectroscopy (with excitation at 350 nm) revealed both the disappearance of emissions signals associated with the naphthyl group and the appearance of emission that correlates with rearomatized adduct (**Figure 4**). Together this suggested *in situ* eliminative generation, via the hydrate as an intermediate. Consistent with this, we were also able to drive essentially similar concomitant elimination through the use of 6-bromo-imidazolopyrimidine (**Figure 4A, step 2ii**); this allowed direct access to a non-hydrated and oxidized protein eH3-Tha9 via debrominative elimination to the rearomatized unsubstituted adduct, **Figure 4A, step 2ii**).

The unsubstituted imidazopyrimidine exhibited fastest conversion (100% to eH3-Tha_H_ after 15 min) compared to the substituted analogues (90-95% after 30 min). Moreover, the proteins exhibited high stability even with increased light intensity levels, thereby allowing reactions to be run even at higher power levels (305 mW/well) to reduce reaction times yet further. Consistent with intended, designed selectivity, no reaction was observed for wildtype H3 (eH3-Lys9) lacking Npa nor in the absence of the imidazopyrimidine reagents. The latter also vitally confirmed the stability of both protein and Npa chromophore to light exposure (i.e. to ‘photobleaching’). Tryptic LC-MSMS of all proteins confirmed site-selective cycloaddition on eH3 protein at position 9 (**Figure 4A, step 2i,ii,iii** and **Figure S9**).

Finally, we characterized, the designed (see above) photophysical perturbation induced by these observed dearomatizations. Pleasingly, spectral changes were found to be essentially identical to those obtained on small molecule analogues (**Figure 2**), allowing useful predictive application. The sharp vibronic features, characteristic for naphthalene as a small molecule (excitation absorption and emission), were found to be retained in eH3-Npa9 (see emission spectrum **Figure 4D, blue**). Moreover, as well as perturbative disruption of this (and other) Npa photophysical features in corresponding eH3-Tha9 cycloadducts, we also observed ‘switch on’ spectral features: clear, novel emission bands were observed at λ_max_ ∼435 nm (eH3-Tha_H_9, **Figure 4D, green**) that are wholly absent in unmodified eH3-Npa9. Together these confirmed not only the designed dearomative ‘switch off’ perturbation of the unique photophysical features of the Npa tags (including fine-structure), but also the ‘switch on’ of new features due to photochemical [4+3] cycloadduct formation, as well as the preservation of the endogenous spectral properties of the protein. In particular, given that the emission spectrum of naphthalene overlaps with that of aromatic amino acids, the modulation of vibronic features suggested that they might be effectively utilized as spectral fingerprints for imaging Npa-tagged proteins, as ‘*vibronic imaging*’. Thus, employing spectral unmixing algorithms in conjunction with these sharp vibronic features might in principle allow enhancement of the ability to accurately separate and distinguish mixed signals (see also **Discussion**).

### Dual Photochemical-Photophysical Probing of Membrane Sensor Protein Annexin A5 in a Conformation– and Site-Selective Photochemical Protein Manifold

The repair and sensing of cellular membranes by corresponding proteins underpins cellular survival and death. As one example, apoptosis as a programmed mode of cell death^54^ is strictly organised. Under normal physiological conditions, the negatively charged phospholipid phosphatidylserine (PS) is located exclusively on the inner leaflet of the mammalian cell membrane (**Figure 6A**, top right). The apoptotic translocation of PS to the outer leaflet of the cell marks the cell for recognition by macrophages.^55^

The annexins are a family of switchable membrane binding proteins with conserved repeating motifs that engage phospholipids to not only facilitate membrane transport but, it has been suggested, to act as sensors of cellular and organismal stress and as modulators of cellular stress responses including perhaps even repair of membrane rupture.^56^ Structural observations have suggested that sequential functions are induced in annexins in response to various stimuli to drive assembly into higher order, two-dimensional ‘mats’, via ligand (lipid) and cofactor (e.g. Ca(II)) engagement and associated conformational changes.^57^

To distinguish early phases of apoptosis, the phospholipid binding protein Annexin A5 (AnxV) in particular has become widely used as a tool for probing the extracellular membrane in, for example, flow cytometry or confocal microscopy^58^ due to its affinity (K_d_ ∼10^-7^-10^−8^ M) for PS;^59,60^ this is fast, non-invasive, non-toxic and can be extended to use in tissues. However, neither the protein itself, nor its complex with PS, can be imaged directly; imaging normally requires labeling or recognition by an intermediary, labelled antibody.^55,58,61^ Moreover, despite its vital utility, the underlying mechanisms and order of events that drive AnxV’s membrane engagement remain unclear.^56,57^

The single Trp residue (Trp187) in AnxV is suggested to play a central role in membrane binding and in related channel activities.^57,62^ It is known that at low calcium concentrations, Trp187 is buried in a hydrophobic niche at the convex surface of AnxV. In the presence of a higher concentration of Ca^2+^ ions^55^ the conformation of the protein is changed, causing the Trp187 loop to be ‘flipped out’ to a presumed solvent-exposed form (**Figure 6A**, orange panel). In this conformation, AnxV also binds the PS headgroup and associates with cell membrane,^63^ however, the precise order of events that lead to full engagement remain unclear, with suggestions that it is in fact PS binding that induces the outward motion of Trp187 to drive oligomerization at critical three-fold axes.^57^ Together these not only raise questions on ordering (PS first or Ca(II) first) but also as to whether an open conformation of AnxV is sufficient for function (or if binding remains Ca-dependent), as well as the question of Trp187’s precise functional role.

We reasoned that putative direct analysis and [4+3]-mediated trapping of Npa187 as a potentially functional Trp187 analogue, as well as the spectral changes of Npa187 in both ‘open’ and ‘closed’ conformation, could provide a robust test of dual photochemical-photophysical probing. Specifically, this could allow us to survey these dynamic conformational changes through accessibility-dependent reactivity (reactive accessibility^64,65^) (**Figure 6A**, blue and green panels).

First, using the same efficient system that we had established for human histone H3 protein (see above), amber codon suppression was used to incorporate Npa into AnxV. Thus, amber stop codon TAG was introduced into the *annexin V* gene at position Trp187 and a corresponding plasmid harboring *annexin V/trp187tag* gene was co-transformed with the generated pEVOL-SS12-TyrRS/tRNA^Tyr^ plasmid into *E. coli* cells. As for H3, induced expression of TyrRS and tRNACUA^Tyr^ in the presence of 2 mM Npa led to efficient production (9 mg/L culture) of full-length annexin AnxV-Npa187 protein, which was purified by ion-exchange and size-exclusion chromatography and fully characterized (**Supplementary Figure S11**).

Next, we created and characterized a ‘trapped’ variant of AnxV-Npa187 in an ‘open’ conformation (**Figure 5**). Pleasingly [4+3] reaction with denaturation (500 mM NH_4_OAc, 3 M guanidine hydrochloride) proceeded with >95% conversion (**Figure 5A,D**) confirming efficient reactivity. The site of cycloaddition at Npa187 was confirmed by LC-MS-MS (**Figure 5,E,F** and **Supplementary Figure S12**) and, as for all prior systems, the photophysical properties of AnxV-Npa187 were shown to be concomitantly perturbed. A clear intense emission band centre at ∼430 nm was generated (**Figure 5B**), and the excitation spectrum of the cycloadduct exhibited a characteristic feature centred at 320 nm (**Supplementary Figure S13**). Fluorescence lifetime was also altered (**Figure 5C**).

**Figure 5.**
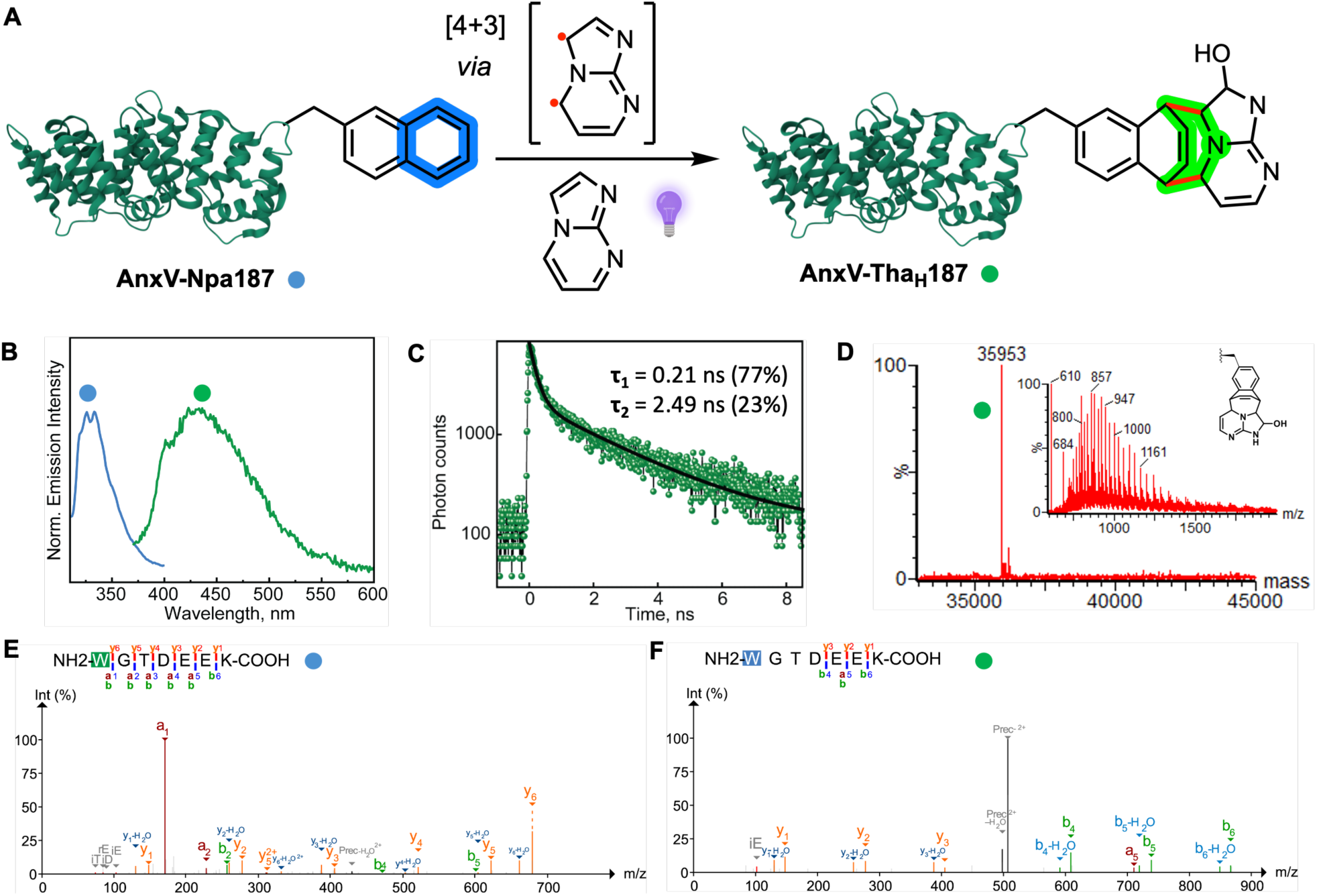
Annexin A5 (AnxV) is Photophysically Perturbed via a Site-Selective Photochemical 4+3 Cycloaddition. **A**) [4+3] cycloaddition of AnxV-Npa187 (blue dot) with unsubstituted imidazolopyrimidine. Conditions: 500 mM NH_4_OAc, 3 M guanidine hydrochloride. **B**) The emission spectrum of AnxV-Npa187 after excitation at 280 nm (**blue**) shows the characteristic vibronic features of naphthalene, thereby confirming the successful incorporation of Npa into the AnxV protein. The emission spectrum of the cycloaddition product on the protein, Anx-Tha_H_187 (**green**) obtained after excitation at 350 nm is also shown. The spectra have been normalized to their respective peak maxima. The presence of a strong, broad emission band centered at ∼430 nm confirms perturbation upon the formation of the cycloaddition product. **C**) Altered AnxV-Tha_H_187 fluorescence lifetime displays bi-exponential decay ι−_1_ = 0.21 ns (77%), ι−_2_ = 2.49 ns (23%). Fluorescence lifetime was obtained after excitation at 680 nm (equivalent to one-photon excitation at 340 nm). **D**) Reaction is efficient with greater than 95% conversion as judged by deconvoluted signal of the respective intact protein mass spectra. **E,F**) LC-MSMS confirms both site of Npa187 incorporation (**E**) and site-selective [4+3] reaction (**F**) in Anx-Npa187.

**Figure 6.**
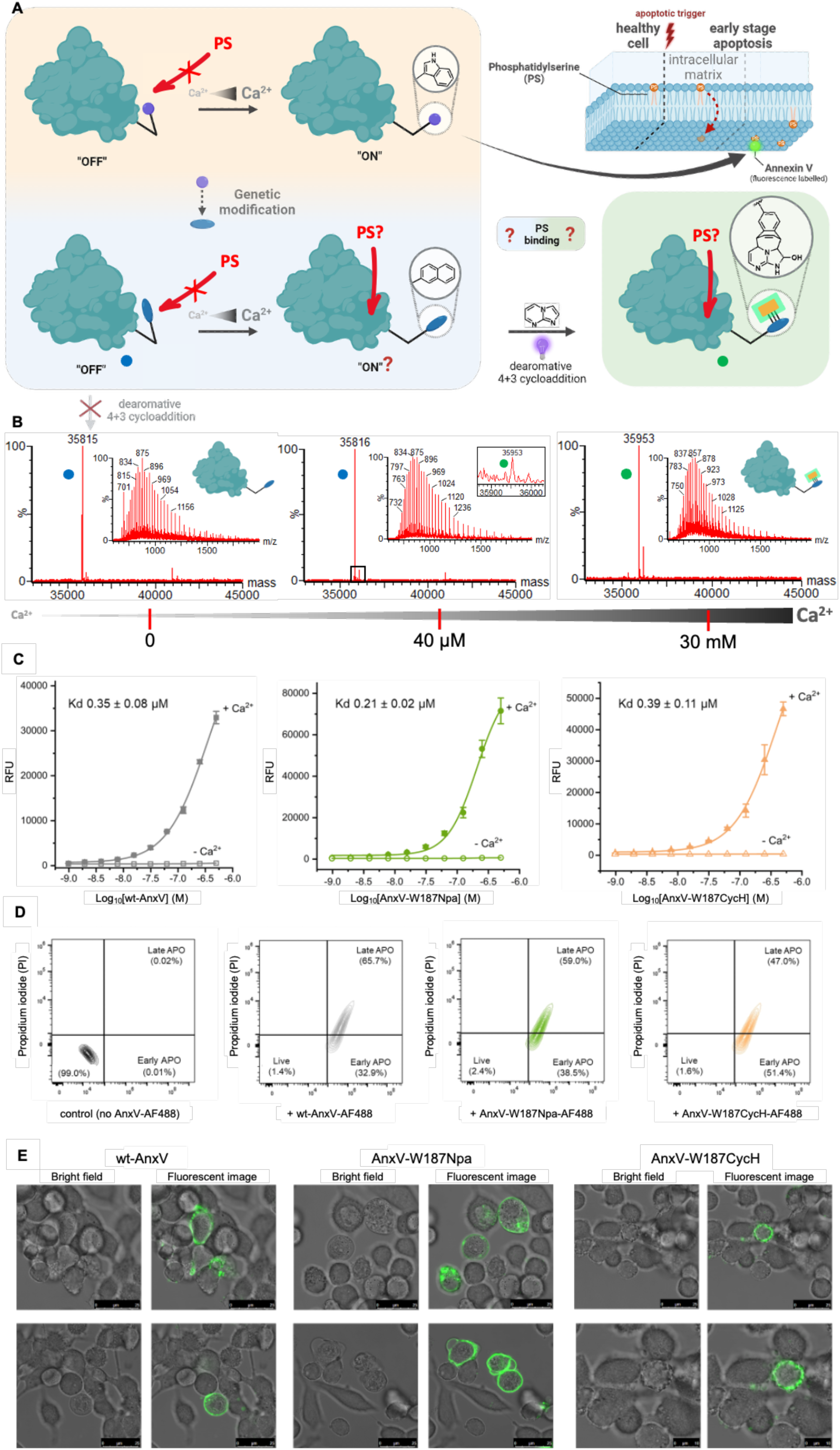
AnxV Function is Unperturbed. **A**) Illustration of the Ca^2+^-dependent conformational change of the Trp-side chain of AnxV, facilitating its usage as indicator of early apoptosis (orange panel). AnxV binds to phosphatidylserine (PS, orange phospholipid) which is translocated to the outer layer of the membrane once apoptosis is initiated by internal or external triggers. This process can be visualized with fluorescent AnxV. **B**) The [4+3] cycloaddition reactions of AnxV-Npa187 (blue dot) with unsubstituted imidazolopyrimidine proceeds by adding Ca^2+^, giving traces of product (green dot) after addition of 40 μM CaCl_2_ (middle) and progressing to up to >95% after addition of 30 mM of CaCl_2_ (right) as illustrated by the deconvoluted signal of the respective intact protein mass spectra; no reaction is observed in the absence of Ca^2+^ (left). **C**) Fluorescence assay of binding affinities of AnxV variants (wild type, AnxV-Npa187, Anx-Tha_H_187) to arrayed PS in the presence or the absence of 5 mM Ca^2+^; the relative fluorescent units are shown as mean ± standard deviation of triplicated experiments. The quantity of bound annexin V obtained from the relative fluorescence unit values was plotted against the logarithmic concentration of total annexin V input. The curve was fitted according to a sigmoidal dose-response profile using Origin 2022. **D**) Representative flow cytometry contour plots using propidium iodide (PI) counterstaining for apoptosis (AnxV +PI). HeLa cells were treated with apoptosis reagent actinomycin D (ActD) for 30 minutes, and then stained with AnxV variants and PI. The FSC-SSC (forward scatter-side scatter) plots were gated according to untreated cells and percentages into four regions. **E**) Localization of AnxV analogues on apoptotic HeLa cells. Laser-scanning confocal microscope shows bright field (grey) and AnxV variants (green) fluorescent images.

### Calcium-dependent Photochemical [4+3] Traps Annexin A5 in an ‘Open’ state that Retains PS– and Cell-binding Function

Next, to interrogate triggers and ordering of annexin mechanism we probed the stimuli-dependent reactivities and functions of AnxV-Npa187 in various ways (**Figure 6**). In the absence of calcium, ‘closed’ AnxV-Npa187 did not react (Tris buffer 20 mM, pH 7.2) with any of the three imidazopyrimidines, even over extended periods. However, with increasing levels of added Ca^2+^(**Figure 6B**) conversion to AnxV-Tha187 cycloadduct formed from imidazo[1,2-*a*]pyrimidine (100 equiv.) increased in a dose-dependent manner: initial indications of product started at 40 μM CaCl_2_ with trace but detectable conversions after 30 min; greater than 95% conversion was observed at 30 mM after just 15 min (**Figure 6B**). This unprecedented, calcium-dependent reactivity of AnxV-Npa187 suggested that conversion of Trp187 to Npa187 had proven sufficiently artefact-free, allowing close mimicry of Trp187 by a zero-size yet perturbable ‘tag’. Moreover, the unique optical signatures of Npa and its cycloadduct allow photo-selective chemical probing of Ca^2+^-bound AnxV-Npa187 through cycloadduct formation, which can be distinctly captured using excitation at 350 nm and collecting emission at ∼430 nm. Together, these results excitingly demonstrated the proof-of-principle of our designed dual photochemical selectivity coupled with photophysical perturbation (**Figure 1**), in a manner apparently governed by reactive accessibility that allows covalent, observational trapping of a corresponding, functionally important dynamic conformational equilibrium (here the calcium cofactor-dependent conformation of the site 187 loop of AnxV).

To further explore the possibility of artefacts in *function* (as opposed to designed perturbation of photophysics), we probed the function of three AnxV variants: wild type AnxV (i.e. with Trp187), AnxV-Npa187 and AnxV-Tha_H_187. Although potentially capable of direct and distinct fluorescence detection using endogenous (wild type AnxV) or introduced (AnxV-Npa187) or induced (AnxV-Tha_H_187) fluorophores, we chose to normalize comparison within a similar dynamic range through the attachment of Alexa Fluor 488 (AF488) dye to Lys residues through use of a corresponding *N*-hydroxysuccinimide ester; modification is distal to the PS binding site at positions Lys286/Lys290.^61^

First, dose-response binding of the AnxV variants directly to arrayed PS was tested. Although the three AnxV variants possess differing side-chains at critical position 187, in the presence of 5 mM Ca^2+^, they all showed similar affinities of binding to PS (*K*_D_: wt-AnxV, 0.35 µM; AnxV-Npa187, 0.21 µM; AnxV-cycloadduct, 0.39 µM). In the absence of Ca^2+^, no binding was observed with any of the variants (**Figure 6C**).

Second, combined AnxV-plus-Propidium iodide (PI) counterstaining was used to monitor apoptotic cells and to quantitatively discriminate apoptotic and necrotic cell profiles via flow cytometry analysis (**Figure 6D**). HeLa cells untreated with apoptosis-inducing reagent actinomycin D and without AnxV staining, as controls, showed 99% viability. HeLa cells incubated with AnxV variants and actinomycin D were detected in both early– and, particularly, late-stages of apoptosis. The populations of cells labelled by AnxV-Npa187 and AnxV-Tha_H_187 variants were counted within similar ranges as that for wild type AnxV; no apparent difference for cell labelling was observed among all three AnxV variants (**Figure 6D**).

Finally, having shown essentially identical cell binding levels, confocal microscopy analysis was performed to determine the cellular localization of each of the AnxV variants. All three annexin V analogues showed selective binding only to the extracellular membrane of apoptotic HeLa cells, whereas control measurements showed no binding to healthy cells (**Figure 6E**).

Together these experiments showed that despite dual photochemical-photophysical perturbation at critical site 187, no apparent alteration of AnxV’s *function* was observed. The site-selective, calcium-dependent reactive trapping shown here of AnxV-Npa187, consistent with an accessible open state that is formed even in the absence of PS, now shows that unlike prior suggestions^57^, AnxV is not dependent on the presence of PS for induced conformational change. Instead, calcium establishes a conformational equilibrium in which an open conformer may be trapped. Furthermore, the continued dependence of PS binding on calcium that we observe for this trapped state suggests both that: a) calcium mediates PS binding; and b) this is possible with even a bulky disrupted [4+3] Tha adduct at site 187. Together, these observations suggest an order of events as follows in Annexin A5 function at membranes: i), calcium binding establishes a dynamic equilibrium that exposes site 187; ii), this and the presence of calcium allows and mediates PS binding, respectively; and iii) the role of site 187 is not critical to these functions and may instead contribute to later establishment of two-dimensional arrays (e.g. in a transition from p6-symmetry array to p3-symmetry array^57^) in which site 187 at protein interfaces might be critical for ‘tightened’ tiling at membranes.

## Discussion

These studies open the door to a system of dual photochemical and photophysical interrogation based upon small, artefact-free, zero-size tags placed into proteins. The approach may enable examination of dynamic biology using photophysical perturbation driven by photochemistry without concomitant artefact generation through any associated functional perturbation.

The methodology we describe introduces a powerful two-mode optical readout system that combines the molecular-level specificity of vibronic fingerprint detection with the dynamic functional reporting of a switchable fluorescence signal. At the core of this approach is a genetically-encoded reactive Npa tag, which now provides a unique opportunity to exploit both the photophysical properties of the parent fluorophore and its chemically-induced transformations by ‘reactive trapping’ using the novel photochemical [4+3] reaction discovered in this work. A defining advantage of naphthalene as a probe is its emission profile. While conventional small molecule fluorophores and intrinsic aromatic residues such as tryptophan, tyrosine, and phenylalanine exhibit broad, featureless bands, Npa fluorescence contains structured vibronic features. These sharp ‘spectral fingerprints’ are particularly advantageous in complex biological environments, where cellular autofluorescence can obscure small molecular probes. By employing spectral unmixing algorithms, the vibronic contributions of the naphthalene tag could be computationally separated from overlapping signals, thereby enabling high-confidence localization and tracking of labelled proteins.^66,67^ This fingerprint-like behaviour is conceptually analogous to Raman spectroscopy, which relies on unique vibrational signatures for molecular identification, but here it can be implemented within a fluorescence framework, making it compatible with bioimaging platforms with spectral-detection capabilities.

In addition to spectral structure, Npa possesses favourable photophysical properties relative to natural amino-acid tryptophan, which has also been used for fluorescence application. Naphthalene exhibits a fluorescence quantum yield of ∼0.23 (in cyclohexane) and a single-exponential lifetime >10 ns (**Figure S7**).^66^ By contrast, tryptophan, the intrinsic amino acid most commonly used for fluorescence application, has a lower quantum yield (∼0.14) and a complex, multiexponential decay typically <5 ns due to heterogeneous local environments and rapid nonradiative relaxation pathways. These differences are highly relevant for imaging. The longer fluorescence lifetime of Npa makes it more amenable to time-resolved detection techniques such as fluorescence lifetime imaging microscopy (FLIM)^68^, where it can be separated from shorter-lived autofluorescence by lifetime gating. Furthermore, the higher quantum yield ensures brighter signals per molecule, increasing sensitivity in low-abundance or high-background contexts. Thus, vibronic fingerprinting could be complemented by lifetime contrast, providing an additional orthogonal parameter for signal discrimination in cells and tissues.

The second dimension of this system is the chemically induced switch from Npa to its [4+3] cycloaddition product Tha. Dearomatization results in the loss of vibronic structure (‘switch-off’) and the emergence of a novel, red-shifted emission band ∼435 nm (‘switch-on’). This not only provides a binary readout of chemical reactivity but also shifts the emission into the visible region. The red-shifted absorption spectrum of the cycloadduct (**Figure S6**) further allows selective excitation of the product without exciting the parent tag, enabling background-free detection of protein states or interactions that trigger cycloaddition (such as protein-protein interaction).

These optical modes could be exploited in a variety of biological contexts. In mechanistic protein studies, the vibronic fingerprint could be used for precise localization of tagged proteins, while the switch-on emission reports conformational changes or intermolecular interactions that enable or induce cycloaddition. Importantly, the separation of the two signals offers unique opportunities for multiplexed and ratiometric detection.^69^ Multiplexing is possible because the vibronic signal (in the near-UV with structured features) and the product emission (in the visible) occupy distinct spectral regimes. Both signals can be collected simultaneously, providing two independent channels from a single labelling event. Moreover, ratiometric analysis between the disappearing naphthalene features and the emerging cycloadduct band can provide a quantitative measure of reaction progression, conformational changes, or protein interactions. Thus, application of combined spectral unmixing (to isolate the naphthalene vibronic peaks) with selective excitation (to isolate the cycloadduct emission) would allow a high degree of optical orthogonality conceptually complementary to multiplexed use of orthogonal spectral channels in autofluorescent environments.^70,71^

The discovery of a suitable, perturbative photochemical protein reaction was critically enabled by high throughput experimentation (HTE). Whilst now prevalent in modes of synthetic chemical methodology,^72,73^ HTE is essentially unexplored in the context of expanding the available chemistry for protein modification. In this way, the use of chemistry with relatively few constraints led to the discovery of a new class of cycloaddition reactions with an intriguing and apparently novel mechanism. This further highlights the benefit of a semi-agnostic approach to push the bounds of discovery not only in chemical biology but also mechanistic chemistry.

Our results here therefore suggest that designed protein editing can be accomplished through a focused lens of *functional* discovery, here in the realm of photochemically-driven modification of photophysics. Through a semi-agnostic approach which had few constraints of design – simply, desired photophysical perturbation enabled by photochemistry – we were able to discover not only a novel reactive ‘tag’, displaying unprecedented Npa reactivity, that may be genetically incorporated into proteins, and then photochemically and selectively modified. Remarkably, the novel photochemical reaction enables selective modification of an uAA that is typically unreactive whilst concomitantly disrupting its fluorophore. The chemical modification apparently creates no functional alteration upon the ensuing biomolecular system (here PS binding by AnxV), thereby allowing elucidation of mechanistic ordering.

Whilst we have not applied this approach here, we note that our combined use of light-based methods for chemistry, spectroscopic interrogation and microscopy could, in the future, even enable simultaneous concomitant discovery and application in certain scenarios. This would be dependent on appropriate to-be-discovered combined photo-devices but would allow perhaps even discovery of functionally relevant photochemistry as perturbations *in situ* (e.g. in or on cells) with a direct read-out of effect also on-the-fly. This type of incontext high throughput methodology is the subject of ongoing experiments in our laboratories.

Finally, the endogenous roles of annexins are diverse both intra– and extra-cellularly.^56^ They have been suggested as modulators of inflammation and even enhancers of the immunogenicity of apoptotic cells through their reduction of macrophage recognition.^74^ Our results here, including mechanistic insight, suggest that holistic synthetic engineering, application and monitoring of Anx variants as synthetic theranostics to both monitor and treat may now be possible in a manner that exploits critical residues, such as Trp187 or mimics, yet without loss of function.

## Methods

### High-Throughput Experimentation

Reactions were performed in micro-scale vials in a 96-well plate format on 100 µL scale. Each reaction comprised of a limiting substrate (final concentration 100 mmol), a distinct cosubstrate (300 mmol) and the photocatalyst Ir[dF(CF_3_)ppy]_2_(dtbpy)PF_6_ (2 mM). Importantly, each of the 28 substrates was included at least once as the limiting as well as the excessive substrate, respectively. Further possible additives were TRIP-thiol (50 mM) or TBPA (200 mM) and TFA (200 mM). To perform the reactions the array of reactants was irradiated with two 40 W Kessil A160WE Tuna blue lamps for 16 hours.

The success of each reaction was evaluated by analytical UPLC-MS, with an additional evaporative light-scattering detector (ELSD) allowing for estimating the product yield.^34^ No assumption was made on the molecular mass of product; all reactions, that had potentially led to at least 0.75 µmol of a mass signal more than 30 higher than any of the starting material substrates (as such corresponding to >8% yield) were deemed productive. The respective reactions yielding products are highlighted in blue in **Figure 2B**. Each of these products were purified by mass directed HPLC followed by elucidating the structures *via* 500 MHz NMR. In total, seven reactions yielded eight validated reaction products (dark blue squares, **Figure 2B**).

### *4+3* cycloaddition-oxidation reaction on histone

In the initial reaction set a 5 mM solution of 7-trifluoromethyl imidazolo[1,2 *a*]pyrimidine in DMSO (100 equivalents, 1% final DMSO concentration) was added to a 50 µM solution of eH3-Npa9 protein in a microscale vial (100 µL). In parallel, a negative control was prepared using wt-eH3 protein. The vials were placed into a DeSyre reaction block in a Zinsser Lumidox lightbox and irradiated for 1 h at 365 nm (65 mW/well) with agitation. Reaction progress was evaluated by intact-protein LC-MS analysis.

Subsequently, where applied, reduction of oxygen in the reaction was achieved by gentle flushing of solutions with nitrogen and degassing in a glovebox (12-16 h) prior to the reaction. Further negative controls were prepared containing eH3-Npa9 but with no addition of small molecule. In this optimized workflow, the mixtures were prepared as before followed by capping the microscale vials and placing them into the reaction block, which was then taken out of the glovebox. The block was put into a Zinsser Lumidox light box and irradiated for 15 min at 365 nm (220 mW/well) with agitation. After this initial time an aliquot was taken and analyzed, while the block was irradiated for another 15 min.

### *4+3* cycloaddition-oxidation reaction on AnxV

Initial reactions were carried out with 15 µM of protein in Tris buffer and otherwise handled as described for previous experiments in microscale vials under reduction of oxygen (in the glove box). In the samples with Ca(II), the salt solution was added prior to addition of 100 equivalents of the respective imidazopyrimidine. Negative controls were prepared accordingly using the wildtype protein instead for its modified analogue. All samples were capped, removed from the glove box, put into the DeSyre reaction block and irradiated for 30 min at 365 nm (305 mW/well) in the Zinsser Lumidox light box. Subsequently, aliquots were taken and analyzed by HRMS. If they did not show any conversion the reactions were further irradiated for 90 min followed by final analysis. Reactions performed under denaturing conditions used acetate buffer (300 mM ammonium acetate, 3 M guanidine HCl).

### Incorporation of Npa tag into proteins via amber codon suppression

Construct generation: The initial pEVOL expression vector was generously donated by Prof.

P.G. Schultz (Scripps Institute). A dual *glnS^ʹ^* (constitutive) / *araBAD* (arabinose-inducible) promoter flanks two *tyrRS* genes. A single tRNA^Tyr^ gene is located between the *proK* promoter and terminator. Mutations in the *ss12-tyrRS* (Y12L/D158P/I159A/L162Q/A167V) gene were generated in a stepwise manner via site-directed mutagenesis (QuickChange II sitedirected mutagenesis kit, Agilent). Two identical *ss12-tyrRS* genes were inserted into the pEVOL vector between BglII and SalI or NdeI and PstI sites. Human histone *eH3.1 (C96A/C110A)* gene with a C-terminal FLAG, HA and His-tag was subcloned into pET3d vector using NcoI and BamHI sites. The eH3.1-K9 → TAG and annexin V-W187 → TAG mutants were generated by site-directed mutagenesis (QuickChange II site-directed mutagenesis kit, Agilent). A list of primers for generating all constructs is provided in Supplementary Information.

L-3-(2-Naphthyl)alanine (Npa) incorporation: The plasmid harbouring human eH3-TAG gene or annexin V-TAG gene was co-transformed with pEVOL-SS12-TyrRS/tRNA^Tyr^ into *E. coli* BL21(DE3) cells. A 30 ml of overnight culture from a single colony was amplified into a 1 L fresh LB media supplemented with 100 µg/ml Ampicillin (Amp) and 25 µg/ml Chloramphenicol (Cm). Cells were grown at 37 °C until OD_600_ reached 0.4∼0.6. Then 1.0 g L-arabinose (0.2%) and 2 mM Npa were added to induce the expression of tRNA synthetase and aminoacylated tRNA and the incubator temperature was reduced to 30 ^°^C. An hour later, cells were induced with 1 mM IPTG and incubated at 30 ^°^C for 16 h. The cell pellets were harvested by centrifugation and stored at –80°C for later extractions and purification.

In order to express wild type annexin V as a control, the plasmid harboring wild type *annexin V* gene was transformed into *E. coli* BL21(DE3) competent cells. A 10 ml sample of overnight culture was added into 1 L fresh LB media in the presence of 100 µg/ml Amp. When the OD_600_ reached 0.4∼0.6, 1 mM IPTG was added, and incubation continued for a further 18 h at 30 ^°^C.

### Protein purification

For histone eH3-Npa9 protein, cell pellets were re-suspended and lysed in 20 ml of wash buffer (50 mM Tris, 100 mM NaCl, pH 7.5) with a cOmplete^TM^ protease inhibitor and 1 mg DNase. The mixture was sonicated using a microtip and centrifuged at 20,000 rpm for 20 mins at 4 ^°^C. The supernatant was discarded, and the pellets were resuspended and washed once with 20 ml of Tris/1% Triton X-100 and another 20 ml of Tris buffer. The pellets were resuspended in 1 ml DMSO at room temperature for 10 min. A 10 ml unfolding buffer (7 M guanidinium chloride, 20 mM Tris, pH 7.5) was added and the mixture was shaken vigorously at room temperature for 1 hr. Following centrifugation at 20,000 rpm for 20 mins, the supernatant was incubated with Ni-NTA resin, which was pre-equilibrated with Tris/urea buffer. The impurities and endogenous proteins were washed with Tris/urea buffer (50 mM Tris, 500 mM NaCl, 7 M urea, pH 7.5) with increased amounts of imidazole (20-60 mM). The final protein was eluted with 150 to 500 mM imidazole in Tris/urea buffer and identified on SDS-PAGE. Purified fractions were dialyzed against water (with 2 mM β-mercaptoethanol) and lyophilized.

For annexin V proteins, cell pellets were suspended in a 30 ml CaCl_2_ buffer (50 mM Tris, 10 mM CaCl_2_, pH 7.2) with a tablet of cOmplete^TM^ protease inhibitor cocktail and 1 mg DNaseI. The mixture was sonicated using a microtip and centrifuged at 20,000 rpm for 15 mins at 4 ^°^C. The supernatant was discarded. The cell pellets were re-suspended in 40 ml EDTA buffer (50 mM Tris, 20 mM EDTA, pH 7.2) and the cell debris was removed by centrifugation (20, 000 rpm, 15 mins, 4 ^°^C). The supernatant containing annexin V was dialyzed against 4 L Tris buffer (20 mM Tris, pH 7.8) 3 times at 4 ^°^C. The dialyzed protein solution was loaded onto a pre-equilibrated 5 ml HiTrap Q HP anion exchange chromatography column for FPLC (ÄKTA GE Healthcare) using solvent A (20 mM Tris, pH 7.8) and solvent B (20 mM Tris, 500 mM NaCl, pH 7.8). The purified fraction was identified by SDS-PAGE and concentrated with Vivaspin centrifugal concentrator (10 kDa cutoff, Cytiva^TM^) to 1 ml. Further purification was performed on a pre-equilibrated Superdex 75 16/600 size-exclusion column with a Tris buffer (20 mM Tris, pH 7.8). The successful incorporation of Npa in annexin V protein was confirmed by LC-MS spectra.

### Fluorescence spectroscopy measurements

All the measurements were performed on Cary Eclipse Fluorescence Spectrophotometer (Agilent Technologies) using a fluorimeter cuvette with path length of 10 mm. For different measurements, excitation and emission slits are varied. During experiments for the comparison of emission intensities, excitation and emission slits are kept constant.

### Time-resolved emission measurements

A custom two-photon microscope was built around a Nikon Ti2 inverted microscope, incorporating a modified Nikon EC2-Si confocal scanning system to support near-infrared laser wavelength transmission. The laser source was a mode-locked tunable laser (Chameleon Discovery NX, Coherent Lasers) with a wavelength range of 660–1320 nm, set to 680 nm with a 100-fs pulse width at 80 MHz. The samples were illuminated on the microscope stage using a focused, diffraction-limited laser spot through a 60x water-immersion objective (Nikon VC; numerical aperture 1.27). Fluorescence emission was collected in non-scanned mode, bypassing the confocal scanning system, and filtered through a BG39 (Comar) and a 650 nm short-pass filter to block the near-infrared laser light. Line, frame, and pixel clock signals were synchronized with an external detector, a fast hybrid photomultiplier tube (HPM100-40, Becker and Hickl, GmbH). The scanning system was connected to a time-correlated single-photon counting (TCSPC) PC module (SPC830, Becker and Hickl) to record TCSPC decays at each pixel or generate single decay curves for solution phase studies.

### UV-Vis absorption measurements

Steady state absorption measurements were performed on an Agilent Cary 60 UV-Vis instrument.

### Labelling of AnxVs with Alexa Fluor 488 NHS ester dye

Annexin V proteins were desalted into 1X PBS buffer via PD SpinTrap G25 column (Cytiva^TM^ life sciences) for reaction with NHS ester. AF488-NHS ester (1.6 µl of 2.5 mM stock in DMSO) was incubated with 40 µM Annexin V solutions at room temperature for 1 h to give a final volume of 100 µl. The reaction was monitored by LC-MS spectra. The percentage of conjugation was determined by LC-MS. Unreacted dye ester in each sample was removed by PD SpinTrap^TM^ G-25 desalting.

### Fluorescence plate assay for determination of annexin V binding affinities to phosphatidylserine (PS)

PS (POPS16:0-18:1, 1-palmitoyl-2-oleoyl-sn-glycero-3-phopho-L-serine, Avanti Polar Lipids) was immobilized on a 96-well F-bottom microplate (µCLEAR®, black, Greiner BIO-ONE) after solubilization in methanol at a concentration of 1.0 mg/ml. Each well was filled with 200 µl of PS solution. After methanol evaporation at room temperature, a film of PS was formed at the bottom of each well. 50 µl of AF488-conjugated annexin V with serial dilutions (ranging from 500 nM to 1 nM) was added to the PS-coated wells and incubated for 1 h at room temperature in the dark. Triplicate experiments were performed in Tris buffer (20 mM Tris, pH 7.8) with or without 5 mM CaCl_2_. The plates were washed three times with 50 µl Tris buffer. Bound annexin V was detected by fluorescence (ex, 488-14 nm; em, 535-30 nm; CLARIOstar^Plus^ plate reader, BMG Labtech) and CLARIOstar-Data Analysis software used for data collection. Quantity of bound AF488-annexin V on PS layer was obtained from the relative fluorescent units (RFU) and plotted against the logarithmic concentration of total AF488-annexin V proteins. Curves were fitted according to a sigmoidal dose-response profile using Origin 2022 to determine protein concentration at half-saturation corresponding to dissociation constant (K_d_).

### Analysis of apoptosis in cells stained with AF488-conjugated annexin V proteins

Confocal microscopy: Confocal microscopic analysis was performed to determine the cellular localization of each of the AF488-conjugated annexin V proteins. HeLa cells (50×10^3^ cells) were seeded into 8-well chamber (300 µl/well, µ-sldie 8 well ibiTreat, Ibidi) and incu-bated at 37 ^°^C for 24 h in an incubator humidified with 5% CO_2_. Cells were washed once with 1χ PBS, and then treated with 0.3 µg (1µg/ml/10^6^ cell) actinomycin D at 37 ^°^C for 30 min in order to induce cell death. Cells were washed twice with annexin binding buffer (10 mM HEPES, 140 mM NaCl, 5 mM CaCl_2_, pH 7.4), and AF488-conjugated annexin V protein was added (0.35 µM/well). After incubation at room temperature in the dark for 15 mins, unbound annexin was washed away with binding buffer. Cells were analyzed using a Leica SP8 confocal microscope (63X/1.40 oil, Argon laser 488).

Flow cytometry: HeLa cells (6 x 10^6^ in a T75 flask) were incubated with actinomycin D (1 µg/ml in 5 ml 1χ PBS) at 37 ^°^C for 30 mins. After 2 washes with 10 ml 1χ PBS, cells were detached from T75 flask with 5 ml 1χ PBS (containing 10 mM EDTA) by centrifugation at 300 g for 5 min and resuspended in a 2 ml annexin binding buffer (10 mM HEPES, 140 mM NaCl, 5 mM CaCl_2_, pH 7.4). Cells were aliquoted into 1.5 ml Eppendorf tubes (200 µl/tube). 4 µM of corresponding AF-conjugated annexin Vs were added to each tube and incubated at room temperature for 15 min in the dark. Unbound annexin V was removed by centrifugation. The treated cells were resuspended in FACS buffer (annexin binding buffer with 1% BSA) containing 5 µl of propidium iodide staining solution (eBioscience^TM^ propidium iodide, ThermoFisher Scientific) and analyzed by flow cytometry (CytoFLEX LX, BECKMAN COULTER). The data was collected with software CytExpert 2.4 and processed by FlowJo v10.9.0. As controls, HeLa cells were also left untreated with apoptosis reagent actinomycin D and without annexin V staining. For cells incubated with annexin V-AF488 variants, the FSCSSC (forward scatter-side scatter) plots were gated according to untreated cells and percentages into four regions. Annexin V-positive cells were denoted as early apoptotic, PI-positive cells were specified as necrotic, while positive cells for both staining indicated late apoptosis in the population.

## Acknowledgements

The authors would like to acknowledge the assistance given by Research IT and the use of the Computational Shared Facility at The University of Manchester. The Chemistry theme at the Rosalind Franklin Institute has been supported by grants from UKRI-EPSRC (EP/V011359/1, EP/V011367/1, EP/T012021/1, EP/X527245/1). We also thank UKRI-EPSRC for grant funding via EP/W002914/1 and EP/N025652/1.

## Author Contributions

Y.Z. reproduced the expression system for the MmNpaRS/tRNA_Pyl_ pair system (A302T/N346G/C348T/V401I/W417Y); reproduced the pEVOL-SS12-TyrRS/tRNA_Tyr_ pair (Y12L/D158P/I159A/L162Q/A167V) and used it to incorporate Npa into eH3-K9TAG model; developed the expression of AnxV-wt, AnxV-Npa187; characterized AnxV-Tha_H_187; and reproduced a PylRS/tRNA_Pyl_ pair system for positive control incorporation of *p*-iPhe. Y.Z. and S.R. performed confocal imaging and optimized the phospholipid binding experiments.

S.G. designed, executed, analysed and purified rounds 1 and 2 of the high throughput screen; identified the [4+3] cycloadduct through analysis, purification, scale-up and crystallisation of product(s) to obtain crystal structure(s).

S.G. and G.B. explored the scope of the [4+3] cycloaddition.

C.A. conducted scoping of small molecule reaction to suit protein conditions; scoping of initial on-protein cycloadduct reactions; and preparation of initial samples for biophysical and MS/MS characterization.

S.L. reproduced the cycloaddition products for biological and photophysical experiments; reassessed and reproduced on-protein reactions and completed characterization; reassessed and reproduced small molecule reactions and completed characterization; gathered, organized and presented revisions of manuscript contents..

A.J. conducted and analyzed all photophysical measurements for small and biomolecule systems; and analyzed all steady-state absorption and emission measurements. AJ performed spectroscopic DFT calculations

A.J., A.M. and S.B. measured time-resolved emission at the Octopus Facility. S.M., A.N. supervised studies into the synthetic scope of the [4+3] cycloaddition. A.L., S.M. and A.N. formulated the mechanistic hypothesis for the cycloadditions. AL performed mechanistic DFT calculations.

All authors contributed content to the Methods and Supplementary Information. S.G., S.L., Y.Z., A.J., A.L., A.N., B.G.D contributed to and generated content for original manuscript drafts. S.L., Y.Z., A.J., A.N. and B.G.D wrote the manuscript. All authors read, revised and approved the manuscript.

## Competing Interests statement

The authors declare no competing interests.

## Notes

### Competing Interest Statement

The authors have declared no competing interest.

