## Supplementary Information for "High-throughput discovery of a [4+3] dearomative cycloaddition enables dual photochemical-photophysical perturbative probing of protein function"

**SUPPLEMENTARY FIGURES** 4

**Figure S1. Substrate set creating the virtual reaction space explored with High-Throughput Chemical synthesis to identify novel photochemical reactions.** 4

**Figure S2. Further validated products from productive reactions of the first and second array round.** 5

**Figure S3. All products from [4+3 cycloaddition] scoping experiments** 6

**Figure S4. Aggregation in Imidazo[1,2-a]pyrimidine (IP).** 7

**Figure S5. Influence of naphthalene mixtures on absorption and fluorescence lifetime of imidazo[1,2-*a*]pyrimidine.** 8

**Figure S6. Spectral properties of cycloaddition products.** 9

**Figure S7. Fluorescence lifetime of naphthalene and cycloaddition product.** 10

**Figure S8. Scoping of Reaction Conditions for On-protein Cycloaddition.** 11

**Figure S9. LC-MS/MS experiments on eH3-Npa9 (referred to in Figure 4).** 12

**Figure S10: Emission spectra of cycloaddition products obtained after reaction with modified Histone (eH3-Npa9).** 13

**Figure S11: Purification of wt- and Npa-modified AnxV.** 14

**Figure S12. LC-MS/MS experiments on ANXV-Npa187 before and after the cycloaddition reaction.** 15

**Figure S13: Excitation spectra of cycloaddition product AnxV-Tha$_H$187.** 16

**Figure S14. Negative control of the cycloaddition reaction on wt-AnxV under denaturing conditions.** 17

**SUPPLEMENTARY TABLES** 18

**Table S1: Reaction condition screening to optimise the yield of the discovered [4+3] cycloaddition-oxidation.** 18

**SUPPLEMENTARY NOTES** 19

**Identification of photochemical reactions enabled by High-Throughput Chemical Synthesis** 19

**Further details of the computational chemistry for the putative mechanism of the 4+3 cycloaddition-oxidation reaction** 31

UV/Visible absorption spectra 31

Intermediate adducts and products 35

Reaction potential energy surface 38

Concerted cycloadditions and singlet diradicals 46

**Failed incorporation of Npa into protein models via the orthogonal MmNpaRS/tRNAPyl pair** 48

**SUPPLEMENTARY METHODS** 52

**General Experimental** 52

**Chemical Methods** 53
High-throughput synthesis arrays to discover the cycloaddition reaction 53
ELSD calibration to estimate reaction productivity from analytical liquid chromatography 54
General procedures for the [4+3] cycloaddition-oxidation with small molecules 54
Synthesized cycloaddition-oxidation products 55

**NMR Spectra** 71
General procedures for [4+3] cycloaddition on proteins 114

**Biological Methods** 117
Construction of Npa-tag *via* amber codon suppression system 117
L-3-(2-Naphthyl)alanine (Npa) incorporation into proteins 118
Protein purification 119
Labelling of annexin Vs with Alexa Fluor 488 NHS ester dye 120
Fluorescence plate assay for the determination of AnxV binding affinities to phosphatidylserine (PS) 120
Analysis of apoptosis cells stained with AF488-conjugated annexin V proteins 121

**Spectroscopy and spectrometry** 122
Fluorescence measurements 122
Tandem mass spectrometry 122

**SUPPLEMENTARY REFERENCES** 124

### Supplementary Figures

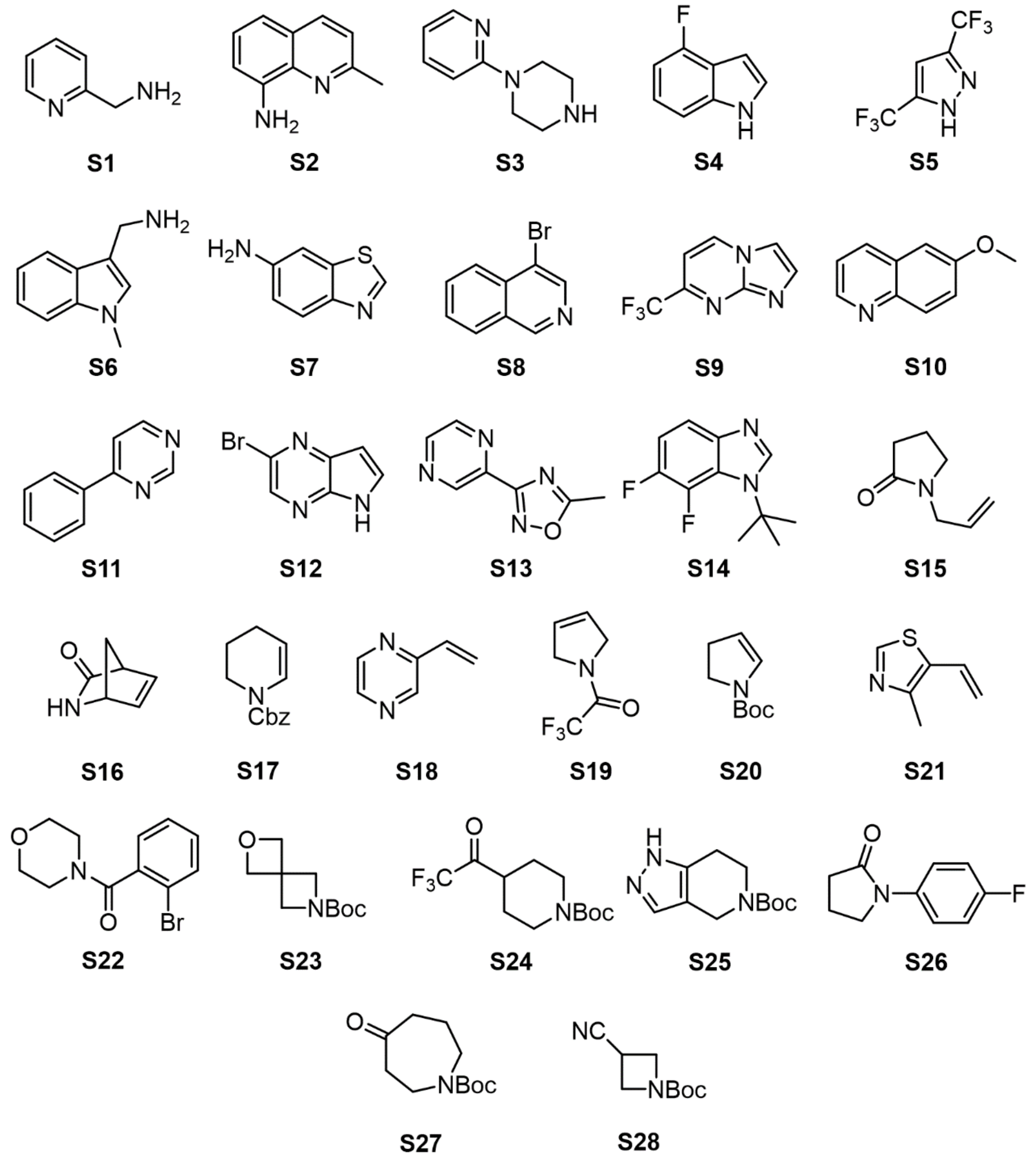

**Figure S1. Substrate set creating the virtual reaction space explored with High-Throughput Chemical synthesis to identify novel photochemical reactions.**
Substrates were picked based on the presence of at least one functional group known to react in photochemical reactions and the potential to be site-selectively incorporated into a protein as part of an unnatural amino acid.

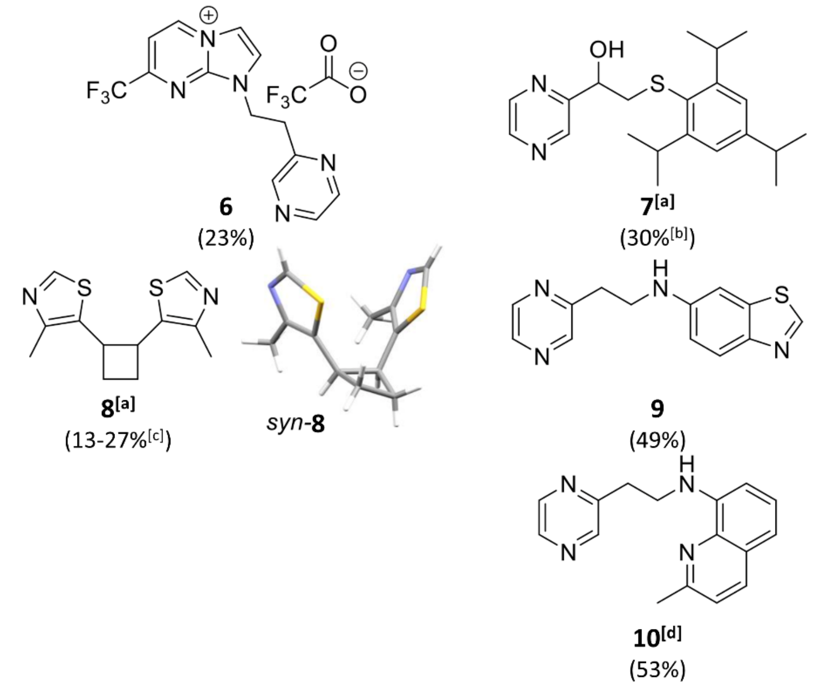

**Figure S2. Further validated products from productive reactions of the first and second array round.**
[a]Homo-dimerization of substrate **S21**; 500 MHz $^{1}H$ NMR spectroscopic analysis revealed a 65:35 mixture of diastereoisomers (*anti*-**8** and *syn*-**8**); the relative configuration of *syn*-**8** was determined by X-ray crystallography.; [b]Yield determined in respect to TRIP-thiol as limiting component; [c]Product observed in three distinct reactions involving **S21**; [d]Discovered in second round of exploration.

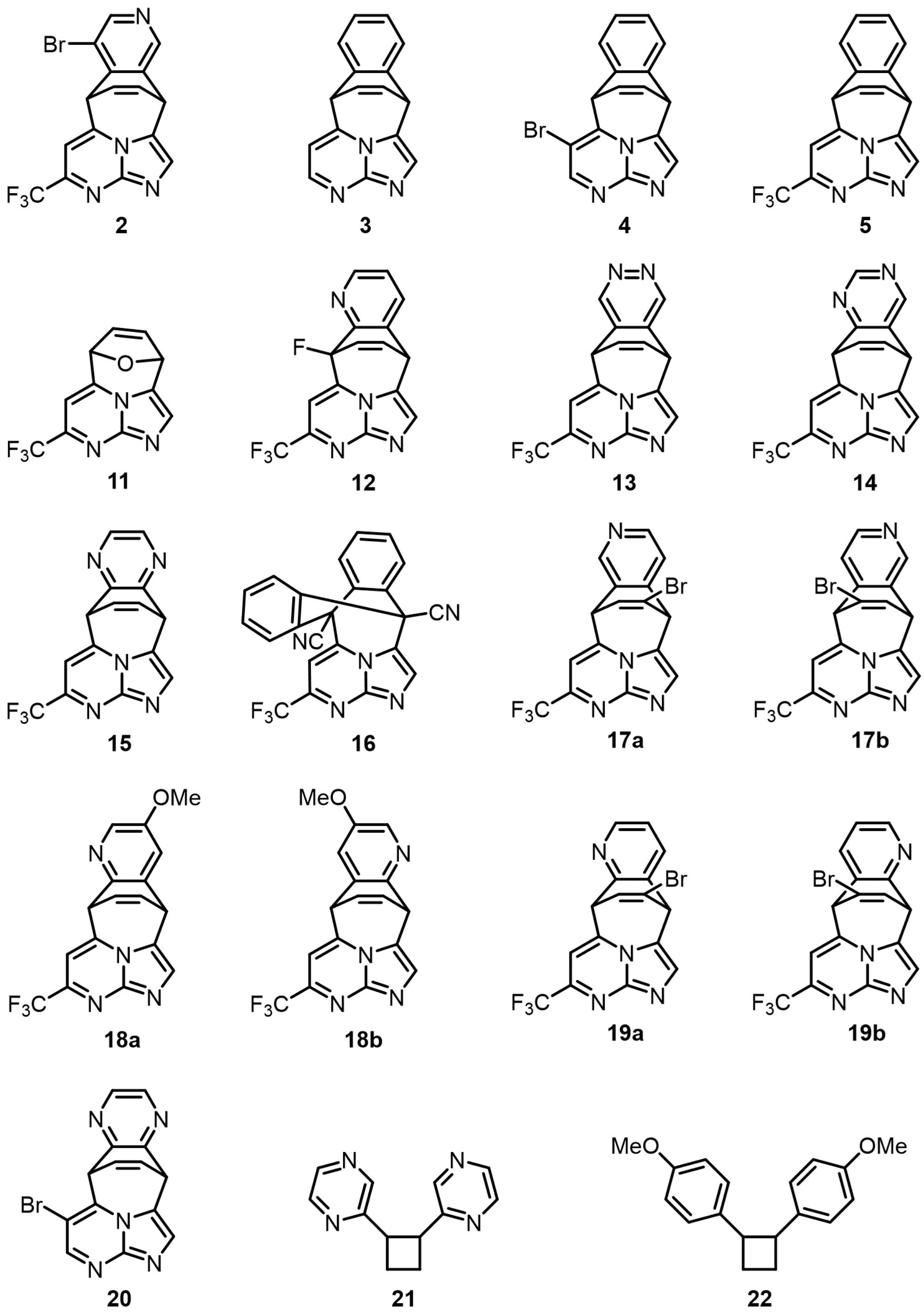

**Figure S3.** All products from [4+3 cycloaddition] scoping experiments

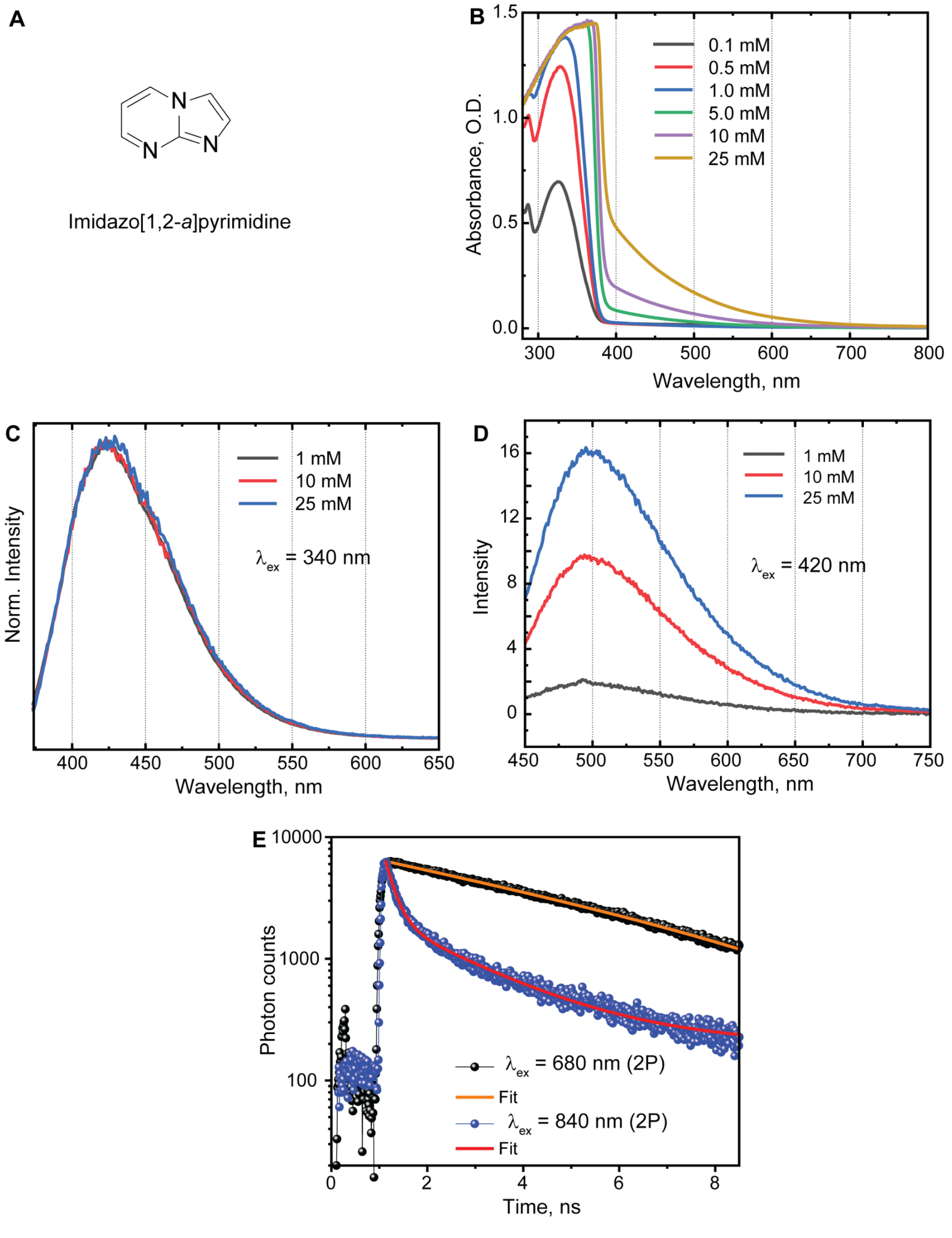

**Figure S4. Aggregation of imidazo[1,2-a]pyrimidine.**
(**A**) Structure of imidazo[1,2-a]pyrimidine (**IP**); (**B**) Steady-state absorption spectra of **IP** in DMSO at different concentrations, showing the appearance of a new lower-energy optical transition relative to the absorption band centered at 330 nm, becoming more prominent at higher concentrations; (**C**) Concentration-dependent emission spectra of **IP** in DMSO after excitation at 340 nm, normalized to compare spectral shapes. No new emission features are observed due to the strong emission band centered at 430 nm; (**D**) Concentration-dependent emission spectra of **IP** in DMSO after excitation at 420 nm (aggregate absorption band), revealing a weak red-shifted emission band centered at 500 nm, which increases in intensity with increasing **IP** concentration; (**E**) Fluorescence lifetime comparison of 25 mM **IP** in DMSO after 2-photon excitation at 680 nm and 840 nm (equivalent to 1-photon at 340 nm and 420 nm, respectively). Fluorescence decays obtained after $\lambda_{ex}$ = 680 nm (2-photon) and 840 nm (2-photon) were fitted with mono-exponential [5.59 ± 0.06 ns] and bi-exponential [0.21 ± 0.01 ns (99.6%) and 1.95 ± 0.03 ns (0.4%)] decay functions, respectively. A faster decay following excitation at 840 nm indicates a non-radiative decay pathway, possibly due to intersystem crossing to triplet states.

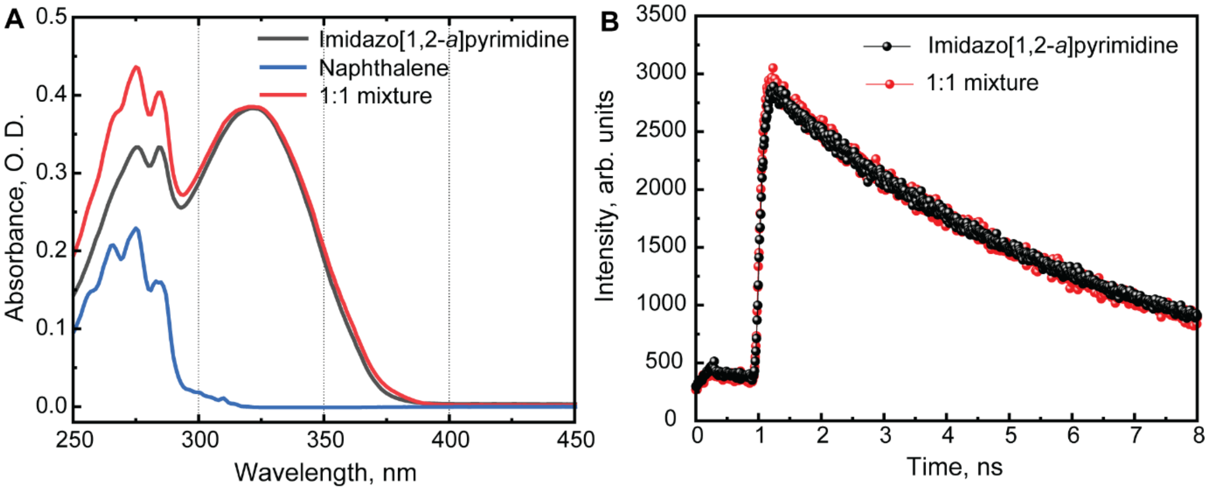

**Figure S5. Influence of naphthalene mixtures on absorption and fluorescence lifetime of imidazo[1,2-*a*]pyrimidine.** (**A**) Comparison of the absorption spectra of the reactants naphthalene (blue) and imidazo[1,2-*a*]pyrimidine (**IP**, black) in DMSO. The absorption spectrum of an equimolar mixture of both reactants (red) reveals no new optical features, indicating no interaction between the two species in the ground state. (**B**) Comparison of the fluorescence lifetimes of 25 mM **IP** and an equimolar mixture of the reactants (25 mM) in DMSO after excitation at 340 nm using two-photon excitation at 680 nm. The lack of change in fluorescence lifetimes suggests the absence of complex formation between the reactants, implying that any observed chemistry upon photoexcitation likely proceeds through a diffusion-based mechanism, potentially involving triplet states

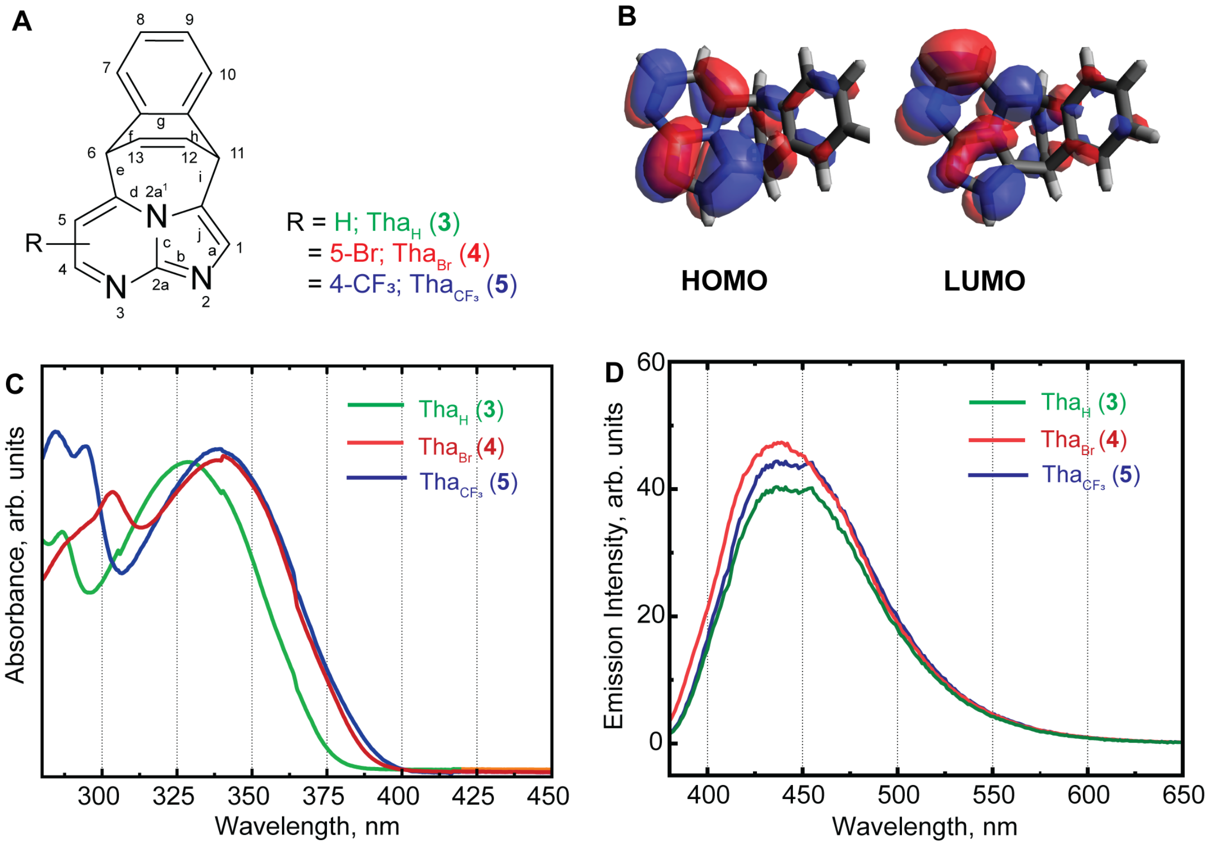

**Figure S6. Spectral properties of cycloaddition products.**
(**A**) Molecular structures of the cycloaddition products; (**B**) Frontier molecular orbitals of $Tha_H$ molecule. Lowest electronic energy transition (HOMO → LUMO) is localised at IP moiety. Molecular orbitals shown here are obtained by performing DFT calculations using B3LYP/6311G(d,p); (**C**) Steady state absorption spectra of the cycloaddition products in DMSO; (**D**) steady state emission spectra of the cycloaddition products in DMSO obtained after excitation 350 nm.

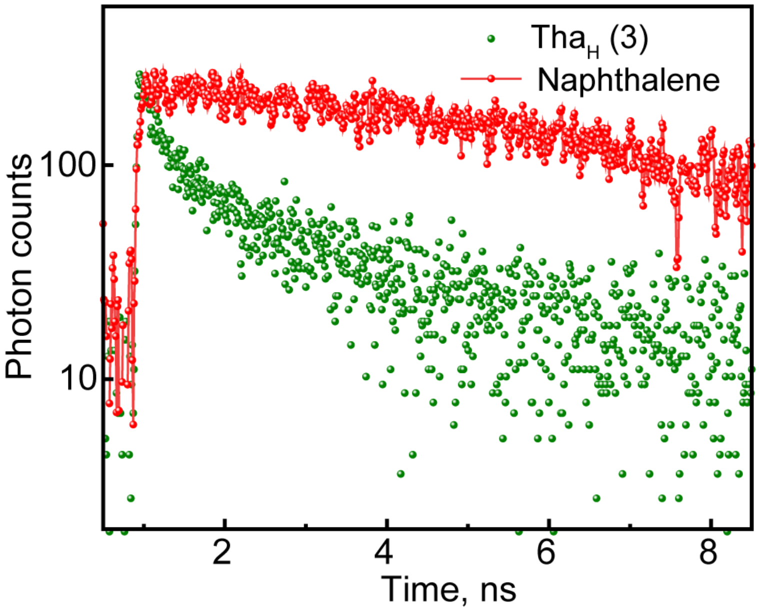

**Figure S7. Fluorescence lifetime of naphthalene and cycloaddition product.**
Comparison of the fluorescence lifetimes of naphthalene (red) and cycloaddition product, Tha$_H$ (3) (green) in DMSO obtained after excitation at 280 nm using two-photon excitation at 560 nm.

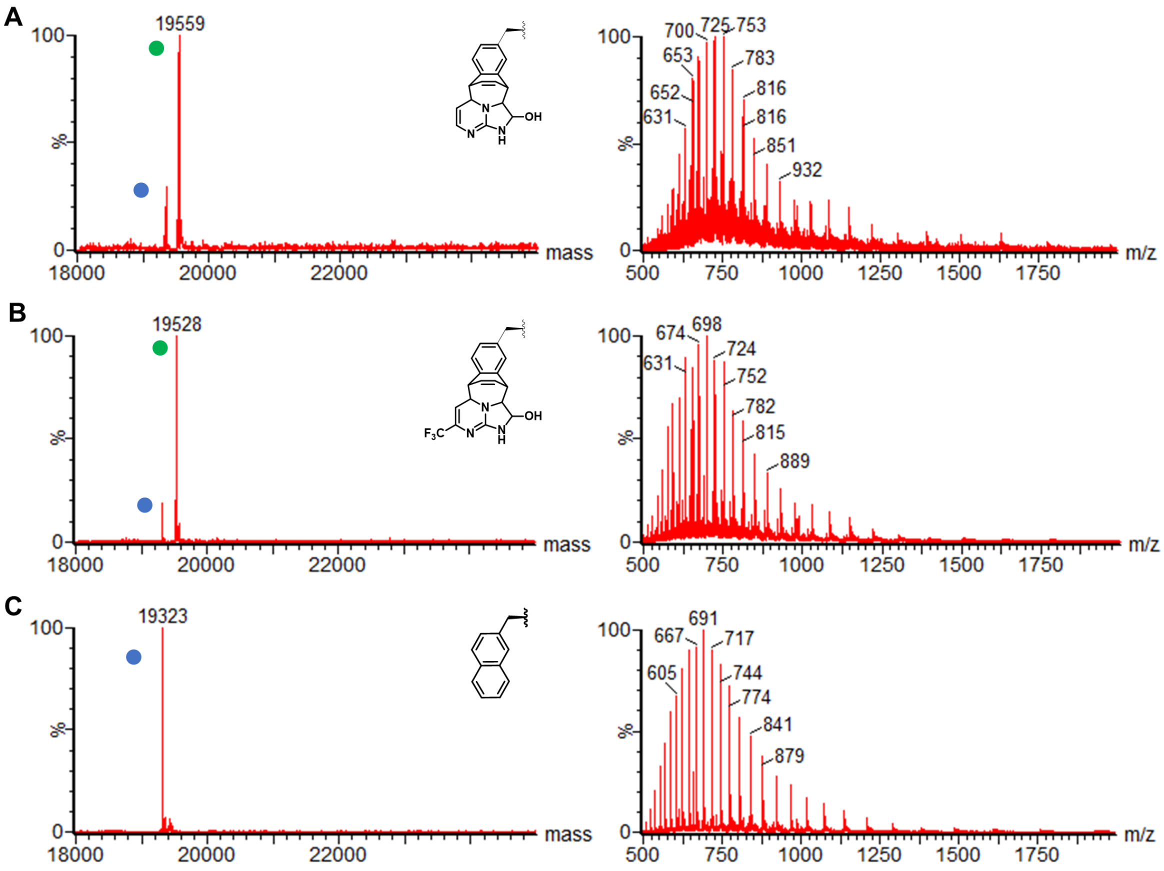

**Figure S8. Scoping of Reaction Conditions for On-protein Cycloaddition.**
Reduction of oxygen levels decreased observed concomitant oxidation of methionines within both starting eH3-Npa9 (blue dot) and products (green dot) to essentially negligible levels (see Methods).

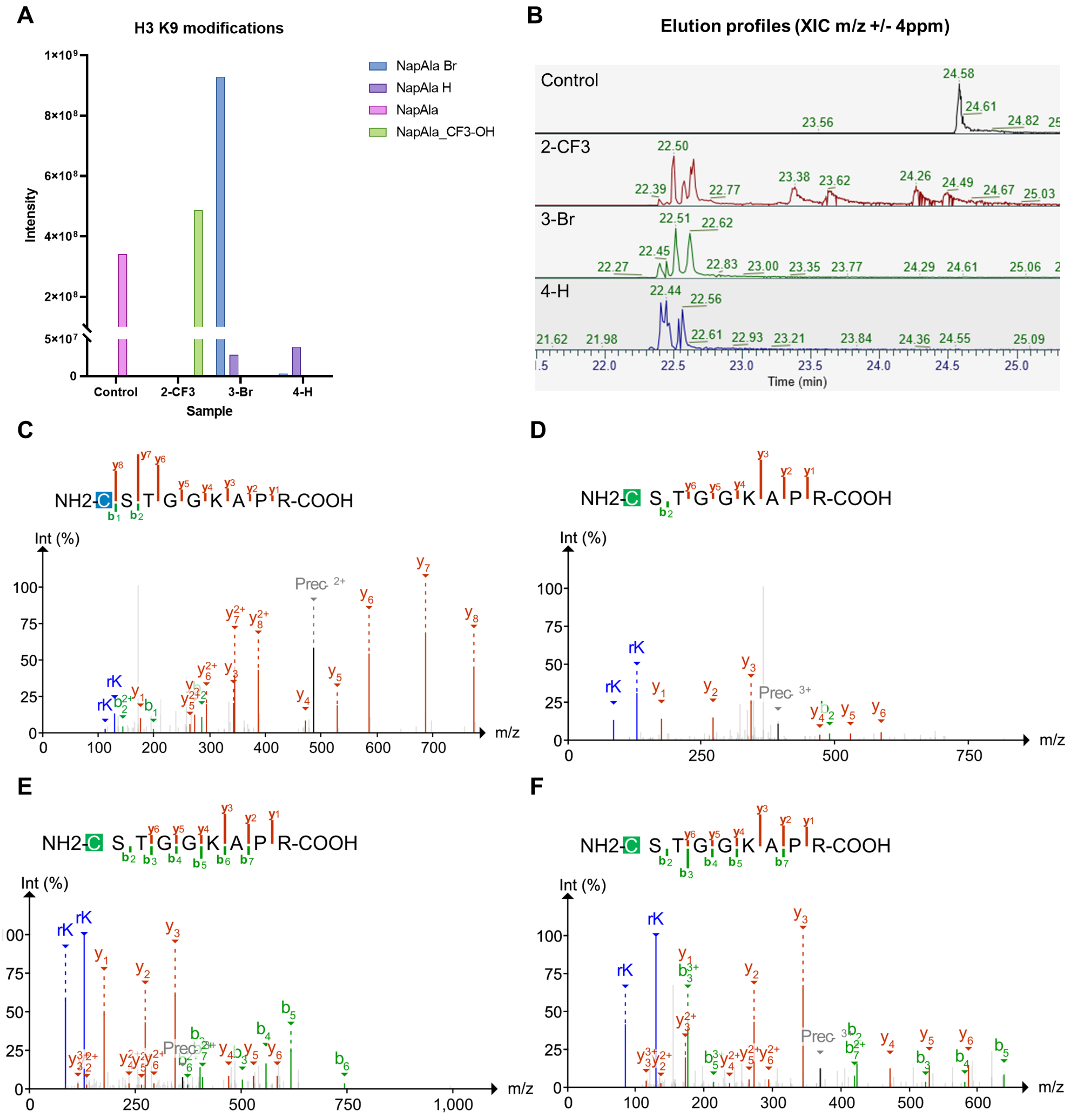

**Figure S9. LC-MS/MS experiments on eH3-Npa9 (referred to in Figure 4).**
**(A)** LC-MS/MS data confirms the quantitative incorporation of naphthyl alanine (Npa) into eH3 (control) and the presence of the cycloaddition product after reaction with each small molecule under denaturing conditions (500 mM ammonium acetate, 3 M guanidine-HCl, pH 6.2), respectively as illustrated by the extracted ion chromatogram signal intensity analysis. The product with 6-bromo imidazo[1,2-*a*]pyrimidine exhibits a minor fraction of unsubstituted product as well (purple column), suggesting that HBr has been eliminated during the course of the reaction. **(B)** Extracted ion chromatograms of all species highlighting the formation of a mixture of product isomers, causing a cohort of closely eluted peaks in the traces of protein after reaction. **(C-F)** MS2 spectra of the eH3-Npa9 before **(C)** and after reaction with 7-trifluoromethyl- **(D)**, 6-bromo- **(E)** and unsubstituted imidazo[1,2-*a*]pyrimidine **(F)** under denaturing conditions.

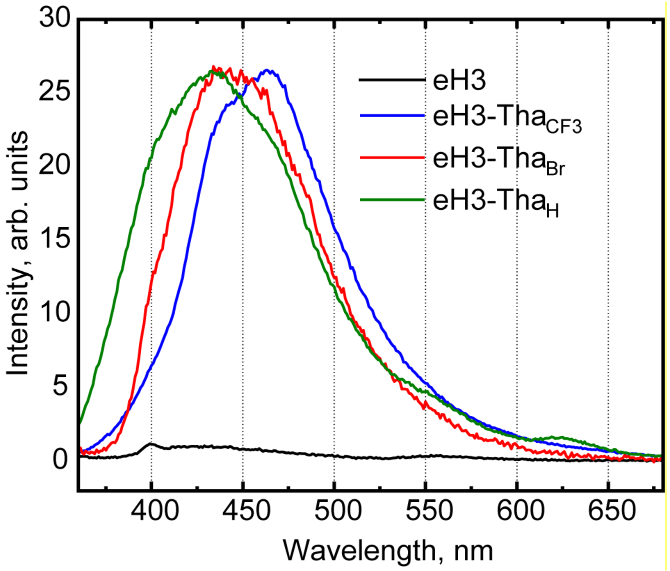

**Figure S10: Emission spectra of cycloaddition products obtained after reaction with modified Histone (eH3-Npa9).** Emission spectra of cycloaddition products with different derivatives: -$CF_3$ (blue), -Br (red) and -H (green) obtained after excitation at 350 nm. The emission spectrum of eH3-Npa9 is shown in black. The spectra have been normalised to their respective peak maxima (except spectrum for the control eH3 sample).

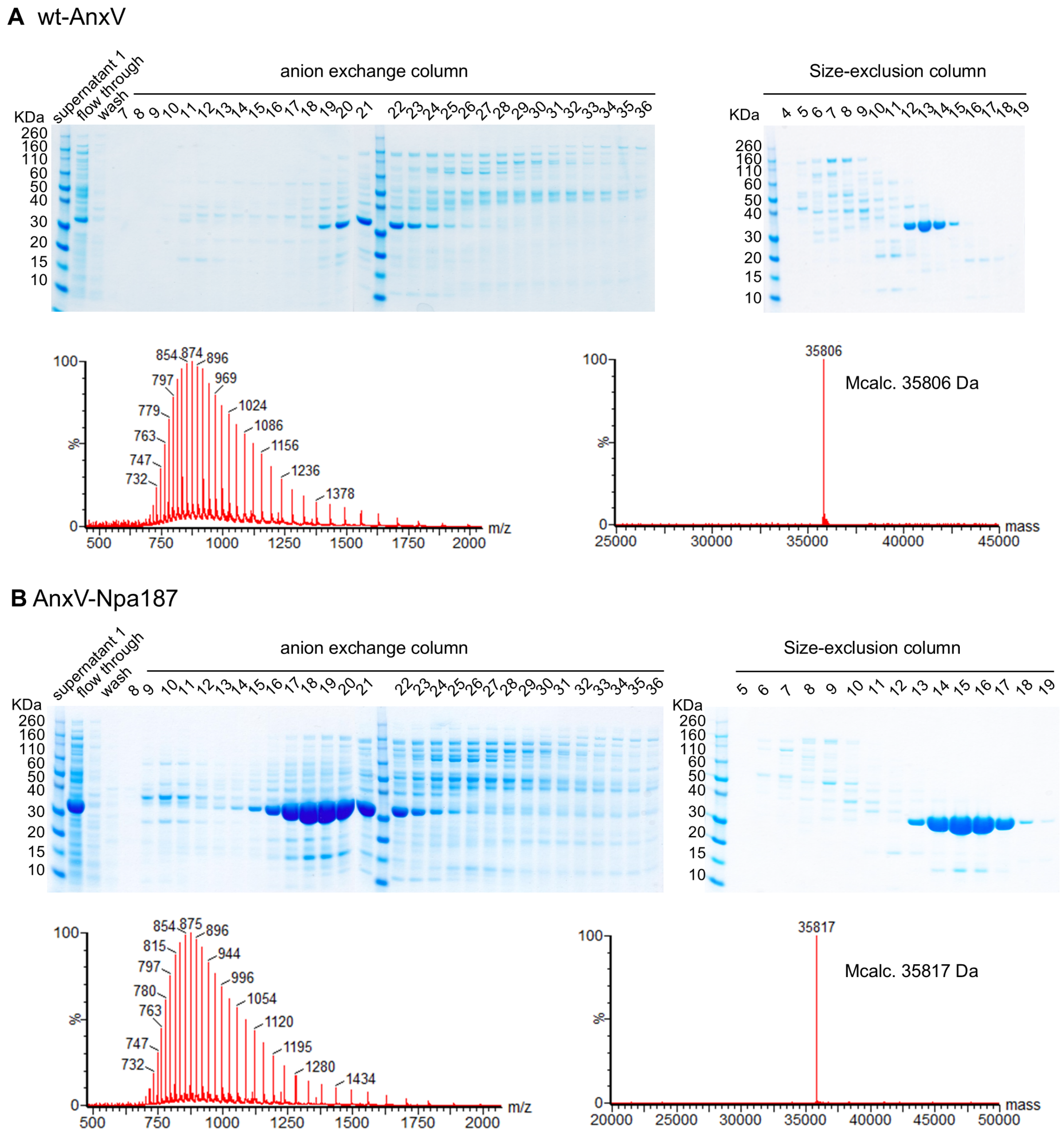

**Figure S11: Purification of wt- and Npa-modified AnxV.**
The purification of wt-Annexin V (a) and Annexin V-Npa187 (b) by anion exchange- and size-exclusion columns. The molecular size of each annexin protein was confirmed by LC-MS spectra (wt-Annexin V, $M_{calc.}$ 35806 Da, $M_{obs.}$ 35806 Da; Annexin V-Npa187, $M_{calc.}$ 35817 Da, $M_{obs.}$ 35817 Da)

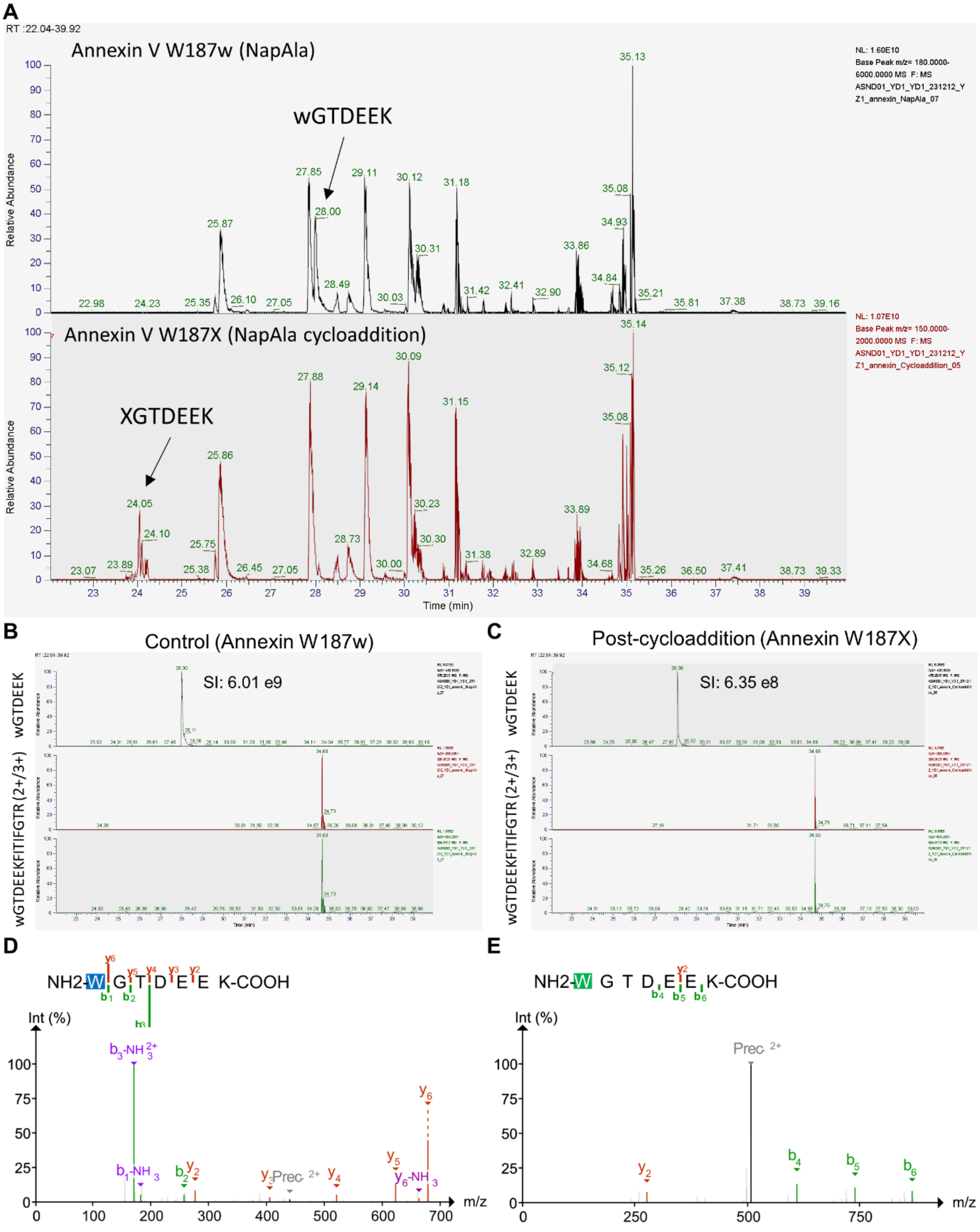

**Figure S12. LC-MS/MS experiments on ANXV-Npa187 before and after the cycloaddition reaction.**
**(A)** Base peak chromatograms of both samples exhibiting a peak ($t_R$= 28.00 min) in the protein sample before the reaction (top), which disappeared after the reaction (bottom) while a convolute of unique peaks had been formed ($t_R$= 24 min). The latter implies the formation of distinct product isomers. **(B-C)** TICs of both species indicating ca. 90% of Npa conversion by the cycloaddition reaction, apparent through the reduction of the signal intensity (SI) of the respective Npa peptide. **(D-E)** Higher-energy collision dissociation (HCD) spectra of both species.

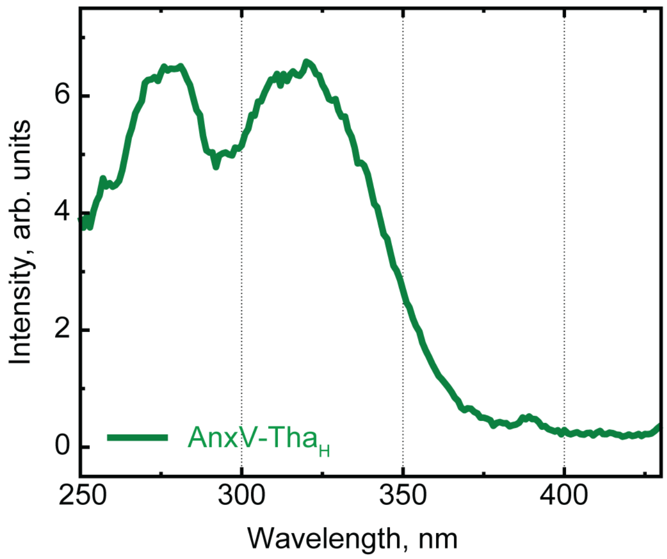

**Figure S13: Excitation spectra of cycloaddition product AnxV-Tha$_H$187.**
Excitation spectra of cycloaddition product obtained after scanning the excitation wavelength and collecting the emission photons at 450 nm.

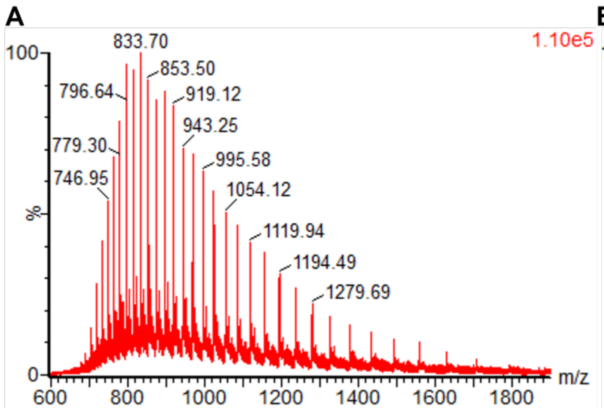

**Figure S14. Negative control of the cycloaddition reaction on wt-AnxV under denaturing conditions.**
Wt-AnxV irradiated with imidazo[1,2-*a*]pyrimidine under denaturing conditions (3 M guanidine-HCl, 500 mM ammonium acetate, pH 6.2). In contrast to Npa-bearing AnxV, only small amounts of non-protein specific (yellow and purple dots) adducts were observed under exhaustive conditions.

#### Supplementary Tables

**Table S1: Reaction condition screening to optimise the yield of the discovered [4+3] cycloaddition-oxidation.** Varying catalyst, oxidant and/or the solvent system of the reaction between the substrates **S9** and **S10**. The yield is represented by the total AUC of the ELS detector trace of the UPLC analysis calibrated to quantify the product species. Each reaction was performed in an identical setup, unless stated otherwise. According to the highest yield, the conditions of entry 14 (bold) were selected for scaled-up synthesis of discovered products.

| Entry | Catalyst | Oxidant (equiv.) | Solvent | Total AUC | Selectivity |
|---|---|---|---|---|---|
| **1** | Ir[dF($CF_3$)ppy]$_2$(dtbpy)$PF_6$ | TBPA (2) | MeCN | 960 | 52:48 |
| **2** | Ir[dF($CF_3$)ppy]$_2$(dtbpy)$PF_6$ | TBPA (5) | MeCN | 345 | 48:52 |
| **3** | Ir[dF($CF_3$)ppy]$_2$(dtbpy)$PF_6$ | TBPA (2) | MeCN/$H_2O$ (4:1) | 897 | 46:54 |
| **4** | Ir[dF($CF_3$)ppy]$_2$(dtbpy)$PF_6$ | TBPA (2) | Acetone | 588 | 54:46 |
| **5** | Ir[dF($CF_3$)ppy]$_2$(dtbpy)$PF_6$ | TBPA (2) | DMSO | 164 | 50:50 |
| **6** | Ir[dF($CF_3$)ppy]$_2$(dtbpy)$PF_6$ | TBPA (2) | DCE | 323 | 46:54 |
| **7** | Ir[dF($CF_3$)ppy]$_2$(dtbpy)$PF_6$ | $NH_4S_2O_8$ | MeCN | 519 | 52:48 |
| **8** | Ir[dF($CF_3$)ppy]$_2$(dtbpy)$PF_6$ | $K_2S_2O_8$ | MeCN | 642 | 52:48 |
| **9**[a] | Ir[dF($CF_3$)ppy]$_2$(dtbpy)$PF_6$ | $LiNO_3$ | MeCN | 548 | 46:54 |
| **10** | Ir[dF(Me)ppy]$_2$(dtbpy)$PF_6$ | TBPA (2) | MeCN | 1170 | 50:50 |
| **11** | Ir(ppy)$_3$ | TBPA (2) | MeCN | 1021 | 50:50 |
| **12** | 4CzIPN | TBPA (2) | MeCN | 914 | 52:48 |
| **13** | Ir[dF(Me)ppy]$_2$(dtbpy)$PF_6$ | TBPA (2) | MeCN/$H_2O$ (4:1) | 970 | 48:52 |
| **14** | **Ir(ppy)$_3$** | **TBPA (2)** | **MeCN/$H_2O$ (4:1)** | **1318** | **46:54** |
| **15**[b] | Ir[dF($CF_3$)ppy]$_2$(dtbpy)$PF_6$ | TBPA (2) | MeCN | 0 | - |
| **16** | Ir[dF($CF_3$)ppy]$_2$(dtbpy)$PF_6$ | - | MeCN | 538 | 52:48 |
| **17** | - | TBPA (2) | MeCN | 830 | 58:42 |

[a] Copper(II)trifluoromethanesulfonate (0.5eq) was added. [b] Reaction was performed in the dark.

### Supplementary Notes

#### Identification of photochemical reactions enabled by High-Throughput Chemical Synthesis

To enable discovery of unique photochemical reactions a reasonably sized virtual reaction space was created, which was subsequently explored by designing two automated parallel synthesis arrays. The virtual reaction space was formed by randomly combining a set of substrates along with defining several possible reaction conditions.

For this, we assembled a set of 28 substrates (Figure S1) that were comparable in size to amino acid side chains (potentially bioisosteric), and that contained at least one functional group (e.g. hetarene, amine, aniline, alkene) with precedented reactivity in photochemical reactions. Each reaction within the combination space involved a pair of those substrates (one in threefold excess) and a photocatalyst (initially Ir[dF($CF_3$)ppy]$_2$(dtbpy)$PF_6$) under one of six reaction conditions. Firstly, either or none of TBPA as oxidant or TRIP-thiol as adjuvant could be added to the reaction mix (→ three variants). Secondly, each of these three variants could be run with or without addition of TFA, yielding six variants per substrate combination.

Initially, we randomly selected 96 reactions (see below) from the 4536 possible permutations (28 substrates x 27 co-substrates in excess x 6 reaction conditions). The success of the reactions was assessed by analytical UPLC-MS, together with evaporative light scattering (ELS) detection to enable product quantification. Reactions were deemed promising if any product with mass ≥30 Da higher than either substrate was produced in >8 (±2)% yield. Such products were then isolated by preparative mass-directed HPLC and structurally elucidated using 500 MHz $^1$H NMR spectroscopy. The executed reactions of both rounds of experiments are illustrated below.

##### Reactions explored in the first round of 96 combinations

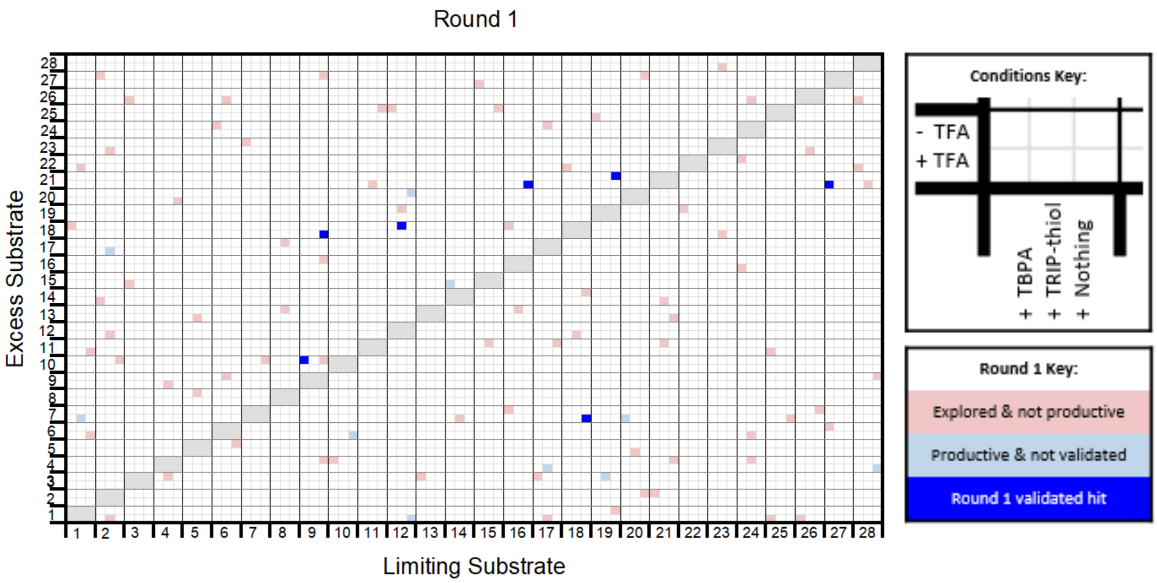

**Figure SN1.1 First round of exploration of the combination space (96 reactions).** The upper chart illustrates the combination space of the 28 substrates, which could either act as limiting or as excessive component in a combination with another non-similar substrate. The black squares depict the six possible reaction condition of each possible substrate combination (see key, top right). Executed combinations are illustrated in coloured squares. Non-productive reactions are shown in red, whereas blue squares exhibit reactions with observed unique products. The dark blue squares, finally, mark substrate combinations with isolated, validated and elucidated unique compounds.

**Table SN1.1** Reactions explored in Round 1

| Limiting Reagent | | Excess Reagent (3 eq) | | Additive? | TFA? | Observed product | | NMR? | Validated? | Compound |
|---|---|---|---|---|---|---|---|---|---|---|
| Code | *M* (g/mol) | Code | *M* (g/mol) | (2 eq TBPA/0.5 eq TRIP-SH) | (2 eq) | *M* (g/mol) | *n* (μmol) | | | number |
| **S15** | 125.2 | **S25** | 223.3 | - | | 239 | 4.86 | | | |
| **S19** | 165.1 | **S21** | 125.2 | - | | 250 | 3.98 | ✓ | ✓ | **8** |
| **S9** | 187.1 | **S27** | 213.3 | - | | | | | | |
| **S2** | 158.2 | **S27** | 213.3 | TBPA | | 407 | 0.95 | | | |
| **S9** | 187.1 | **S10** | 159.2 | TBPA | | 344 | 0.90 | ✓ | ✓ | **1a, 1b** |
| **S21** | 125.2 | **S11** | 156.2 | TRIP-thiol | | 377 | 1.10 | | | |
| **S1** | 108.1 | **S11** | 156.2 | - | TFA | 476 | 0.40 | | | |
| **S28** | 182.2 | **S4** | 135.1 | - | TFA | 417 | 0.67 | ✓ | x | |
| **S28** | 182.2 | **S26** | 179.2 | TBPA | TFA | | | | | |
| **S26** | 179.2 | **S1** | 108.1 | TBPA | TFA | | | | | |
| **S1** | 108.1 | **S7** | 150.2 | TRIP-thiol | TFA | 347 | 3.38 | ✓ | x | |
| **S2** | 158.2 | **S1** | 108.1 | TRIP-thiol | TFA | 263 | 0.81 | | | |
| **S2** | 158.2 | **S10** | 159.2 | - | | | | | | |
| **S20** | 169.2 | **S27** | 213.3 | - | | 376 | 0.52 | | | |
| **S6** | 160.2 | **S5** | 204.1 | - | | | | | | |
| **S21** | 125.2 | **S2** | 158.2 | TBPA | | | | | | |
| **S7** | 150.2 | **S23** | 199.3 | TBPA | | | | | | |

| | | | | | | | | | | |
|---|---|---|---|---|---|---|---|---|---|---|
| **S12** | 198 | **S18** | 106.1 | TRIP-thiol | | 358 | 1.48 | ✓ | ✓ | **7** |
| **S10** | 159.2 | **S6** | 160.2 | - | TFA | 301 | 6.51 | ✓ | x | |
| **S9** | 187.1 | **S18** | 106.1 | - | TFA | 293 | 2.26 | ✓ | ✓ | **6** |
| **S14** | 210.2 | **S15** | 125.2 | TBPA | TFA | 333 | 1.89 | ✓ | x | |
| **S25** | 223.3 | **S1** | 108.1 | TBPA | TFA | | | | | |
| **S21** | 125.2 | **S14** | 210.2 | TRIP-thiol | TFA | 377 | 0.46 | | | |
| **S18** | 106.1 | **S12** | 198 | TRIP-thiol | TFA | 358 | 0.94 | | | |
| **S9** | 187.1 | **S10** | 159.2 | - | | 344 | 0.64 | | | |
| **S9** | 187.1 | **S16** | 109.1 | - | | | | | | |
| **S18** | 106.1 | **S14** | 210.2 | - | | | | | | |
| **S17** | 217.3 | **S3** | 163.2 | TBPA | | | | | | |
| **S5** | 204.1 | **S8** | 208.1 | TRIP-thiol | | | | | | |
| **S12** | 198 | **S19** | 165.1 | TRIP-thiol | | | | | | |
| **S12** | 198 | **S1** | 108.1 | - | TFA | 308 | 5.36 | ✓ | x | |
| **S16** | 109.1 | **S21** | 125.2 | - | TFA | 250 | 3.08 | | | |
| **S19** | 165.1 | **S25** | 223.3 | TBPA | TFA | 261 | 7.45 | | | |
| **S2** | 158.2 | **S17** | 217.3 | TRIP-thiol | TFA | 453 | 3.78 | ✓ | x | |
| **S24** | 281.3 | **S6** | 160.2 | TRIP-thiol | TFA | 301 | 7.54 | | | |
| **S4** | 135.1 | **S9** | 187.1 | TRIP-thiol | TFA | 417 | 0.54 | | | |
| **S28** | 182.2 | **S9** | 187.1 | - | | 370 | 0.51 | | | |

| | | | | | | | | | | |
|---|---|---|---|---|---|---|---|---|---|---|
| **S19** | 165.1 | **S1** | 108.1 | - | | | | | | |
| **S13** | 162.2 | **S3** | 163.2 | TBPA | | | | | | |
| **S24** | 281.3 | **S22** | 270.1 | TBPA | | | | | | |
| **S4** | 135.1 | **S3** | 163.2 | TRIP-thiol | | | | | | |
| **S17** | 217.3 | **S24** | 281.3 | TRIP-thiol | | 473 | 0.43 | | | |
| **S18** | 106.1 | **S7** | 150.2 | - | TFA | 256 | 4.89 | ✓ | ✓ | **9** |
| **S20** | 169.2 | **S7** | 150.2 | TBPA | TFA | 488 | 1.46 | ✓ | x | |
| **S24** | 281.3 | **S16** | 109.1 | TBPA | TFA | | | | | |
| **S23** | 199.3 | **S28** | 182.2 | TRIP-thiol | TFA | | | | | |
| **S23** | 199.3 | **S18** | 106.1 | TRIP-thiol | TFA | 318 | 1.27 | | | |
| **S11** | 156.2 | **S21** | 125.2 | TRIP-thiol | TFA | 361 | 0.97 | | | |
| **S12** | 198 | **S20** | 169.2 | - | | 376 | 1.91 | ✓ | x | |
| **S21** | 125.2 | **S4** | 135.1 | - | | 250 | 1.26 | | | |
| **S1** | 108.1 | **S18** | 106.1 | TBPA | | 212 | 1.68 | | | |
| **S16** | 109.1 | **S18** | 106.1 | TBPA | | 212 | 1.52 | | | |
| **S8** | 208.1 | **S13** | 162.2 | TRIP-thiol | | | | | | |
| **S13** | 162.2 | **S13** | 162.2 | TRIP-thiol | | | | | | |
| **S21** | 125.2 | **S13** | 162.2 | - | TFA | | | | | |
| **S28** | 182.2 | **S22** | 270.1 | TBPA | TFA | | | | | |
| **S3** | 163.2 | **S26** | 179.2 | TBPA | TFA | | | | | |

| | | | | | | | | | |
|---|---|---|---|---|---|---|---|---|---|
| **S1** | 108.1 | **S22** | 270.1 | TRIP-thiol | TFA | | | | |
| **S26** | 179.2 | **S23** | 199.3 | TRIP-thiol | TFA | | | | |
| **S24** | 281.3 | **S26** | 179.2 | TRIP-thiol | TFA | | | | |
| **S26** | 179.2 | **S7** | 150.2 | - | | | | | |
| **S17** | 217.3 | **S11** | 156.2 | - | | | | | |
| **S27** | 213.3 | **S6** | 160.2 | TBPA | | | | | |
| **S6** | 160.2 | **S24** | 281.3 | TBPA | | | | | |
| **S15** | 125.2 | **S11** | 156.2 | TRIP-thiol | | 358 | 0.44 | | |
| **S6** | 160.2 | **S9** | 187.1 | TRIP-thiol | | | | | |
| **S4** | 135.1 | **S20** | 169.2 | - | TFA | 326 | 1.33 | | |
| **S25** | 223.3 | **S11** | 156.2 | TBPA | TFA | | | | |
| **S15** | 125.2 | **S27** | 213.3 | TBPA | TFA | | | | |
| **S14** | 210.2 | **S7** | 150.2 | TRIP-thiol | TFA | | | | |
| **S2** | 158.2 | **S12** | 198 | TRIP-thiol | TFA | | | | |
| **S17** | 217.3 | **S4** | 135.1 | TRIP-thiol | TFA | 270 | 2.60 | ✓ | x |
| **S20** | 169.2 | **S2** | 158.2 | - | | | | | |
| **S9** | 187.1 | **S4** | 135.1 | - | | | | | |
| **S22** | 270.1 | **S19** | 165.1 | TBPA | | | | | |
| **S12** | 198 | **S25** | 223.3 | TBPA | | | | | |
| **S16** | 109.1 | **S13** | 162.2 | TRIP-thiol | | 345 | 0.38 | | |

| | | | | | | | | | |
|---|---|---|---|---|---|---|---|---|---|
| **S8** | 208.1 | **S17** | 217.3 | TRIP-thiol | | 400 | 0.37 | | |
| **S25** | 223.3 | **S7** | 150.2 | - | TFA | | | | |
| **S2** | 158.2 | **S14** | 210.2 | TBPA | TFA | | | | |
| **S27** | 213.3 | **S21** | 125.2 | TBPA | TFA | 250 | 1.98 | | |
| **S6** | 160.2 | **S26** | 179.2 | TRIP-thiol | TFA | | | | |
| **S2** | 158.2 | **S23** | 199.3 | TRIP-thiol | TFA | | | | |
| **S17** | 217.3 | **S1** | 108.1 | TRIP-thiol | TFA | 235 | 2.48 | | |
| **S11** | 156.2 | **S25** | 223.3 | - | | 239 | 3.76 | | |
| **S7** | 150.2 | **S10** | 159.2 | - | | | | | |
| **S10** | 159.2 | **S4** | 135.1 | TBPA | | | | | |
| **S16** | 109.1 | **S7** | 150.2 | TBPA | | | | | |
| **S24** | 281.3 | **S4** | 135.1 | TRIP-thiol | | 534 | 0.66 | | |
| **S19** | 165.1 | **S3** | 163.2 | TRIP-thiol | | 301 | 4.82 | ✓ | x |
| **S1** | 108.1 | **S6** | 160.2 | - | TFA | | | | |
| **S3** | 163.2 | **S15** | 125.2 | TBPA | TFA | | | | |
| **S18** | 106.1 | **S22** | 270.1 | TBPA | TFA | 267 | 0.91 | | |
| **S5** | 204.1 | **S13** | 162.2 | TRIP-thiol | TFA | | | | |
| **S20** | 169.2 | **S5** | 204.1 | TRIP-thiol | TFA | | | | |
| **S28** | 182.2 | **S21** | 125.2 | TRIP-thiol | TFA | 377 | 0.68 | | |

#### Reactions explored in the second round of 96 combinations

Informed by the validated reactions from the first discovery round, a second reaction array was designed. The validated reactions were adjusted in three ways:

a) swapping of limiting and excess substrates;
b) exchanging of either substrate with a related substrate from the initial set (e.g. hetarene for another hetarene);
c) exploring the five other reaction conditions.

An expanded range of validated reactions were identified in the second round (**Figure S2**), including a [4+3] cycloaddition-oxidation reaction involving the isoquinoline **S8** (in place of the quinoline **S10**) leading to the formation of **2** (**Figure 2C**). The detailed information on the selected combinations and reaction conditions is illustrated in **Figure SM 2** and **Table SM 2** (see below).

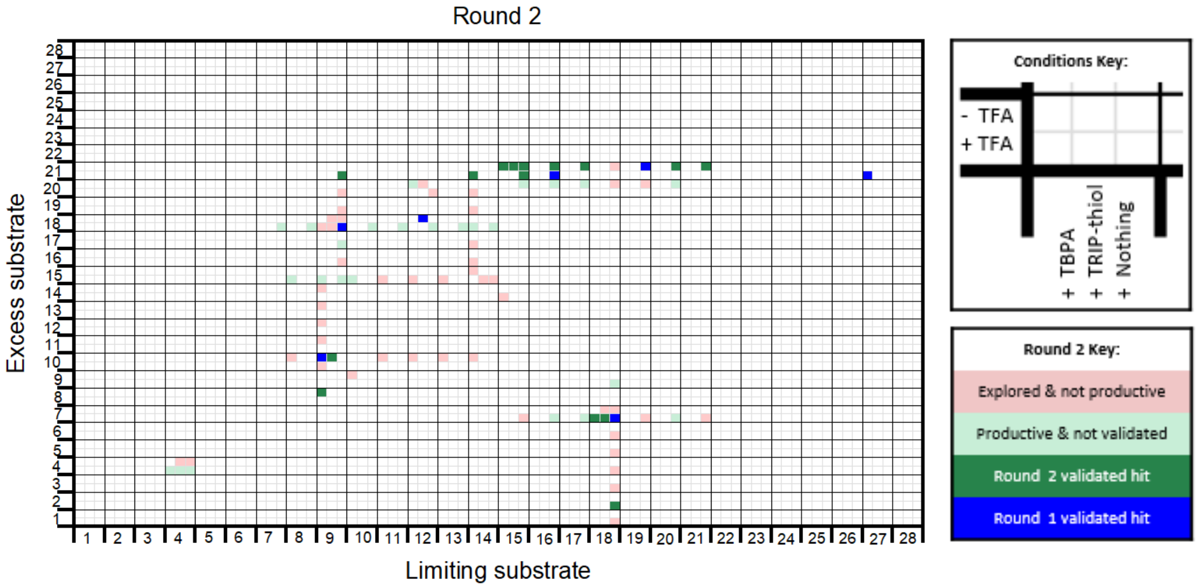

**Figure SN1.2 Second round of exploration of the combination space (96 reactions).** Informed by the results of the first round, validated substrate combinations were further investigated by either swapping of limiting and excess substrates, exchanging either one with a structurally related substrate from the initial set, or exploring of the five other reaction condition variants. For reference, the validated hit combinations from the first round are shown in blue squares. Explored combinations from the second round are highlighted in non-blue squares, with non-productive reactions depicted in red and productive but non-validated reactions in light green. Successfully validated productive reactions are illustrated as dark green squares.

**Table SN1.2** Reactions explored in Round 2.

| Limiting Reagent | | Excess Reagent (3 eq) | | Additive? | TFA? | Observed product(s) | | NMR? | Validated? | Compound |
|---|---|---|---|---|---|---|---|---|---|---|
| Code | *M* (g/mol) | Code | *M* (g/mol) | (2 eq TBPA/0.5 eq Thiol) | (2 eq) | *M* (g/mol) | *n* (µmol) | | | Number |
| **S15** | 125 | **S21** | 125 | - | | 250 | 8.35 | | | |
| **S9** | 187 | **S10** | 159 | - | | 344 | 1.00 | | | |
| **S15** | 125 | **S20** | 169 | - | | 338<br>354 | 2.22<br>1.48 | | | |
| **S9** | 187 | **S13** | 162 | TBPA | | | | | | |
| **S12** | 198 | **S20** | 169 | TBPA | | 338<br>354 | 0.83<br>1.72 | | | |
| **S18** | 106 | **S7** | 150 | - | TFA | 256 | 6.14 | | | |
| **S16** | 109 | **S7** | 150 | - | TFA | 259 | 6.81 | | | |
| **S18** | 106 | **S9** | 187 | - | TFA | 293 | 1.43 | | | |
| **S10** | 159 | **S18** | 106 | - | TFA | 265 | 1.88 | | | |
| **S18** | 106 | **S7** | 150 | TBPA | TFA | 256 | 6.32 | | | |
| **S14** | 210 | **S20** | 169 | TBPA | TFA | | | | | |
| **S9** | 187 | **S18** | 106 | TBPA | TFA | | | | | |
| | | **S21**[a] | 125 | - | | 250 | 7.13 | | | |
| **S9** | 187 | **S18** | 106 | - | | | | | | |
| | | **S4**[a] | 135 | - | | | | | | |
| **S9** | 187 | **S14** | 210 | TBPA | | | | | | |
| **S15** | 125 | **S21** | 125 | Thiol | | 250 | 7.78 | | | |

| | | | | | | | | | | |
|---|---|---|---|---|---|---|---|---|---|---|
| **S18** | 106 | **S1** | 108 | - | TFA | | | | | |
| **S17** | 217 | **S7** | 150 | - | TFA | 584 | 2.82 | | | |
| **S9** | 187 | **S15** | 125 | - | TFA | 310 | 0.49 | | | |
| **S11** | 156 | **S18** | 106 | - | TFA | 262 | 5.62 | | | |
| **S9** | 187 | **S10** | 159 | TBPA | TFA | 344 | 0.47 | | | |
| **S14** | 210 | **S21** | 125 | TBPA | TFA | 250 | 1.49 | | | |
| | | **S4**[a] | 135 | TBPA | TFA | 270 | 6.41 | | | |
| **S16** | 109 | **S21** | 125 | - | | 250 | 7.45 | | | |
| **S12** | 198 | **S20** | 169 | - | | 338<br>354 | 2.56<br>2.85 | | | |
| **S15** | 125 | **S21** | 125 | TBPA | | 250 | 6.61 | | | |
| **S8** | 208 | **S10** | 159 | TBPA | | | | | | |
| **S18** | 106 | **S7** | 150 | Thiol | | | | | | |
| **S18** | 106 | **S2** | 158 | - | TFA | 264 | 5.34 | ✓ | ✓ | **10** |
| **S19** | 165 | **S7** | 150 | - | TFA | | | | | |
| **S9** | 187 | **S16** | 109 | - | TFA | | | | | |
| **S12** | 198 | **S18** | 106 | - | TFA | 318 | 1.89 | | | |
| **S14** | 210 | **S15** | 125 | TBPA | TFA | 333 | 0.83 | | | |
| **S8** | 208 | **S15** | 125 | TBPA | TFA | 330 | 1.34 | | | |
| **S18** | 106 | **S7** | 150 | Thiol | TFA | 256 | 2.37 | | | |
| **S17** | 217 | **S21** | 125 | - | | 250 | 7.16 | | | |
| | | **S20**[a] | 169 | - | | 338 | 1.48 | | | |

| | | | | | | | |
|---|---|---|---|---|---|---|---|
| **S9** | 187 | **S10** | 159 | TBPA | | 344 | 0.83 |
| **S11** | 156 | **S10** | 159 | TBPA | | | |
| **S9** | 187 | **S10** | 159 | Thiol | | 344 | 1.01 |
| **S18** | 106 | **S3** | 163 | - | TFA | | |
| **S20** | 169 | **S7** | 150 | - | TFA | 488 | 1.53 |
| | | | | | | 486 | 0.98 |
| **S9** | 187 | **S17** | 217 | - | TFA | 404 | 1.27 |
| **S13** | 162 | **S18** | 106 | - | TFA | 318 | 1.95 |
| **S15** | 125 | **S14** | 210 | TBPA | TFA | 333 | 0.54 |
| **S9** | 187 | **S15** | 125 | TBPA | TFA | 310 | 1.40 |
| **S14** | 210 | **S15** | 125 | Thiol | TFA | 333 | 0.32 |
| **S18** | 106 | **S21** | 125 | - | | 231 | 3.93 |
| | | | | | | 250 | 6.11 |
| **S16** | 109 | **S20** | 169 | - | | 338 | 1.60 |
| | | | | | | 354 | 1.44 |
| **S10** | 159 | **S9** | 187 | TBPA | | 344 | 0.58 |
| **S12** | 198 | **S10** | 159 | TBPA | | | |
| **S9** | 187 | **S18** | 106 | Thiol | | 293 | 0.49 |
| **S18** | 106 | **S4** | 135 | - | TFA | 347 | 0.72 |
| | | | | | | 482 | 0.45 |
| | | | | | | 241 | 0.53 |
| | | | | | | 270 | 0.82 |

| | | | | | | | | | | |
|---|---|---|---|---|---|---|---|---|---|---|
| **S21** | 125 | **S7** | 150 | - | TFA | | | | | |
| **S9** | 187 | **S19** | 165 | - | TFA | | | | | |
| **S14** | 210 | **S18** | 106 | - | TFA | 316 | 1.18 | | | |
| **S14** | 210 | **S16** | 109 | TBPA | TFA | | | | | |
| **S10** | 159 | **S15** | 125 | TBPA | TFA | 282 | 1.12 | | | |
| **S9** | 187 | **S18** | 106 | Thiol | TFA | | | | | |
| **S19** | 165 | **S21** | 125 | - | | 250 | 6.56 | | | |
| **S17** | 217 | **S20** | 169 | - | | 338<br>354 | 1.21<br>0.95 | | | |
| **S9** | 187 | **S8** | 208 | TBPA | | 392 | 0.75 | ✓ | ✓ | **2** |
| **S13** | 162 | **S10** | 159 | TBPA | | | | | | |
| **S12** | 198 | **S20** | 169 | Thiol | | | | | | |
| **S18** | 106 | **S5** | 204 | - | TFA | 318 | 0.63 | | | |
| **S7** | 150 | **S18** | 106 | - | TFA | 362<br>256 | 3.23<br>2.96 | | | |
| **S9** | 187 | **S20** | 169 | - | TFA | | | | | |
| **S12** | 198 | **S20** | 169 | - | TFA | | | | | |
| **S14** | 210 | **S17** | 217 | TBPA | TFA | 441 | 0.77 | | | |
| **S11** | 156 | **S15** | 125 | TBPA | TFA | | | | | |
| | | **S4**[a] | 135 | Thiol | TFA | 270 | 7.34 | | | |
| **S20** | 169 | **S21** | 125 | - | | 250 | 8.95 | | | |
| **S18** | 106 | **S20** | 169 | - | | 338 | 0.84 | | | |

| | | | | | | | |
|---|---|---|---|---|---|---|---|
| **S9** | 187 | **S11** | 156 | TBPA | | | |
| **S14** | 210 | **S10** | 159 | TBPA | | 367 | 0.23 |
| | | **S4**[a] | 135 | Thiol | | | |
| **S18** | 106 | **S6** | 160 | - | TFA | 330 | 0.85 |
| **S14** | 210 | **S15** | 125 | - | TFA | | |
| **S9** | 187 | **S21** | 125 | - | TFA | 250 | 5.24 |
| **S28** | 182 | **S4** | 135 | - | TFA | 270 | 2.83 |
| **S14** | 210 | **S18** | 106 | TBPA | TFA | 316 | 1.16 |
| **S12** | 198 | **S15** | 125 | TBPA | TFA | | |
| **S18** | 106 | **S7** | 150 | - | | | |
| **S19** | 165 | **S20** | 169 | - | | | |
| **S9** | 187 | **S12** | 198 | TBPA | | | |
| **S14** | 210 | **S15** | 125 | TBPA | | 333 | 0.43 |
| **S15** | 125 | **S21** | 125 | - | TFA | 250 | 5.04 |
| **S15** | 125 | **S7** | 150 | - | TFA | 334 | 0.81 |
| **S9** | 187 | **S18** | 106 | - | TFA | 293 | 2.72 |
| **S8** | 208 | **S18** | 106 | - | TFA | 313 | 5.66 |
| | | **S4**[a] | 135 | - | TFA | 270 | 7.88 |
| **S14** | 210 | **S19** | 165 | TBPA | TFA | | |
| **S13** | 162 | **S15** | 125 | TBPA | TFA | | |

[a] mono-substrate reaction

#### Further details of the computational chemistry for the putative mechanism of the 4+3 cycloaddition-oxidation reaction

The computational studies described were all performed in Gaussian 16.(M. J. Frisch, G. W. Trucks, H. B. Schlegel, G. E. Scuseria, M. A. Robb, J. R. Cheeseman, G. Scalmani, V. Barone, G. A. Petersson, H. Nakatsuji, X. Li, M. Caricato, A. V. Marenich, J. Bloino, B. G. Janesko, R. Gomperts, B. Mennucci, H. P. Hratchian, J. V. Ortiz, A. F. Izmaylov, J. L. Sonnenberg, D. Williams-Young, F. Ding, F. Lipparini, F. Egidi, J. Goings, B. Peng, A. Petrone, T. Henderson, D. Ranasinghe, V. G. Zakrzewski, J. Gao, N. Rega, G. Zheng, W. Liang, M. Hada, M. Ehara, K. Toyota, R. Fukuda, J. Hasegawa, M. Ishida, T. Nakajima, Y. Honda, O. Kitao, H. Nakai, T. Vreven, K. Throssell, J. A. Montgomery Jr., J. E. Peralta, F. Ogliaro, M. J. Bearpark, J. J. Heyd, E. N. Brothers, K. N. Kudin, V. N. Staroverov, T. A. Keith, R. Kobayashi, J. Normand, K. Raghavachari, A. P. Rendell, J. C. Burant, S. S. Iyengar, J. Tomasi, M. Cossi, J. M. Millam, M. Klene, C. Adamo, R. Cammi, J. W. Ochterski, R. L. Martin, K. Morokuma, O. Farkas, J. B. Foresman and D. J. Fox, Gaussian 16, Revision C.01, Gaussian, Inc., Wallingford CT, **2016**) The calculations employed the same method previously employed by Glorius to study a photo-cycloaddition.(R. Kleinmans, T. Pinkert, S. Dutta, T. O. Paulisch, H. Keum, C. G. Daniliuc and F. Glorius, *Nature* **2022**, *605*, 477-482) This entailed the use of the M06-2X functional and the def2-TZVP basis set.(Y. Zhao and D. G. Truhlar, *Theor. Chem. Account.* **2008**, *120*, 215-241; Y. Zhao and D. G. Truhlar, *Acc. Chem. Res.* **2008**, *41*, 157-167; F. Weigend and R. Ahlrichs, *Phys. Chem. Chem. Phys.* **2005**, *7*, 3297-3305). Solvation was incorporated via the IEFPCM protocol with default settings for toluene. (J. Tomasi, B. Mennucci and E. Cancès, *J. Mol. Struct.: Theochem* **1999**, *464*, 211-226; J. Tomasi, B. Mennucci and R. Cammi, *Chem. Rev.* **2005**, *105*, 2999-3094). UV/Visible spectra were computed via the TD keyword and included allowed transitions to excited singlet states.(C. Adamo and D. Jacquemin, *Chem. Soc. Rev.* **2013**, *42*, 845-856; A. D. Laurent, C. Adamo and D. Jacquemin, *Phys. Chem. Chem. Phys.* **2014**, *16*, 14334-14356). Open shell singlet diradicals were computed via the guess=mix keyword that invokes the mixing of HOMO and LUMO to remove spatial symmetry for electrons of opposing spin; optimizations for open shell singlet species were initiated from optimized geometries of triplet species. Free energies were obtained using Goodvibes v. 2.0.2 and employing concentrations of 1 M at 298 K.(G. Luchini, J. V. Alegre-Requena, I. Funes-Ardoiz and R. S. Paton. *F1000Research* **2020**, *9(Chem Inf Sci)*, 291 DOI: 10.12688/f1000research.22758.1; S. Grimme, *Chem. Eur. J.* **2012**, *18*, 9955-9964).

**UV/Visible absorption spectra**

The first property to be studied computationally was the excitation via photon absorption that would be necessary to initiate the reaction in the absence of catalyst. It is assumed that such a process would occur via the allowed transition to an excited singlet state followed by inter-system crossing to the triplet state. As will be described later, the singlet excited states were found likely to be unproductive reactive intermediates.

The computed spectra along with the values of the longest wavelength absorbance maximum and corresponding oscillator strength are provided in Table SN2.1. Alongside these, the relative energies of the geometry optimized triplet state (NB not vertical excitation) are provided.

It became clear that the absorption behaviour of all long-lived species that might be formed during the reaction must be considered; the values for a range of the reactants (**S9, SI1-5**), intermediate adducts (**C**, **D** and **SI6**) and final products (**F** and **SI7-8**) that are relevant are therefore included in Table SN2.1.

**Table SN2.1** Computed UV/visible absorbance spectra for a range of relevant species and energies of lowest triplet states.

| **Species** | **Spectrum** | **Longest wavelength** (nm) [oscillator strength] | **Relative energy of optimized triplet** (kcal/mol) |
|---|---|---|---|
| 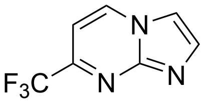 **S9** | 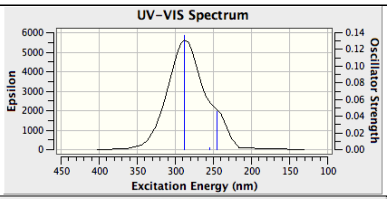  | 287.78 [0.14] | 57.5 |
| 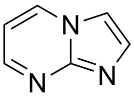 **SI1** | 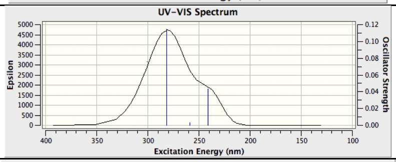  | 281.89 [0.11] | 59.0 |
| 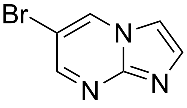 **SI2** | 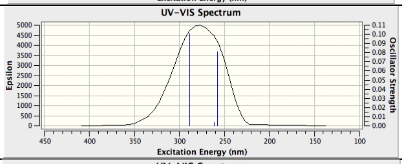  | 288.87 [0.10] | 57.0 |
| 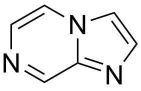 **SI3** | 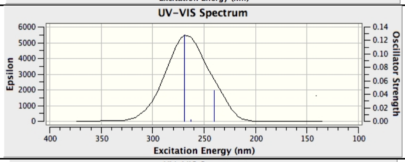  | 269.23 [0.13] | 60.8 |
| 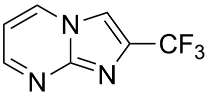 **SI4** | 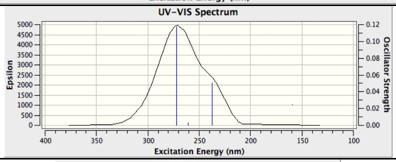  | 272.26 [0.12] | 61.2 |
| 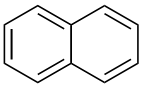 **SI5** | 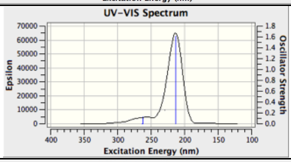  | 263.47 [0.00] | 64.4 |
| 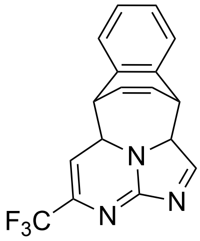 *endo*-**D** | 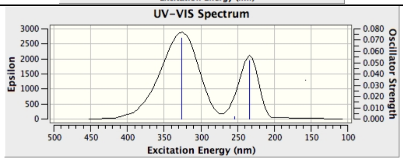  | 325.93 [0.07] | 49.6 |
| 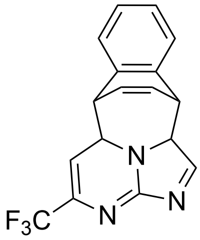 *exo*-**D** | 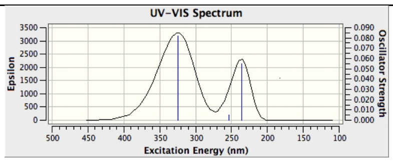  | 324.88 [0.08] | 49.9 |

| | | | |
|---|---|---|---|
| 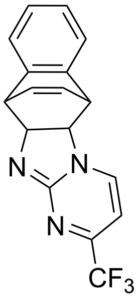 *endo*-**SI6** | 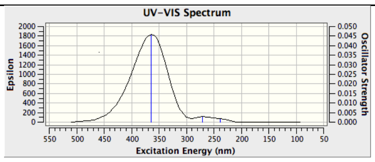  | 363.66 [0.05] | 44.0 |
| 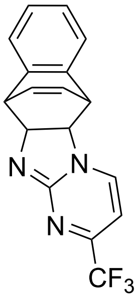 *exo*-**SI6** | 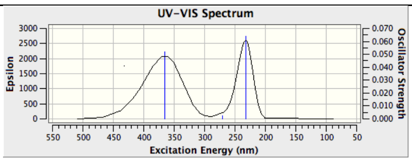  | 325.93 [0.07] | 43.8 |
| 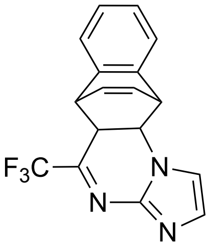 *endo*-**C** | 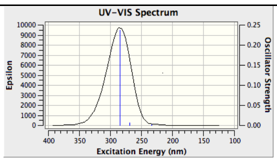  | 284.83 [0.24] | 50.9 |
| 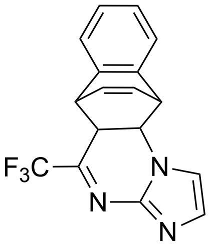 *exo*-**C** | 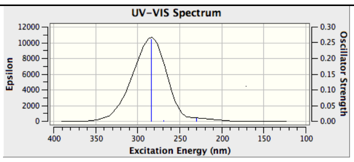  | 284.19 [0.26] | 51.0 |
| 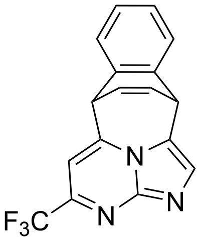 **F** | 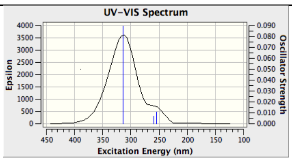  | 313.4 [0.09] | |
| 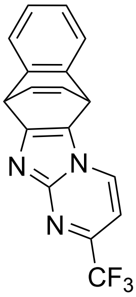 **SI7** | 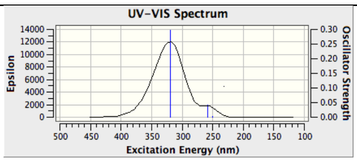  | 319.9 [0.30] | |

| | | |
|---|---|---|
| 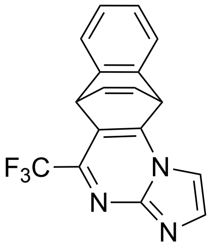 **SI8** | 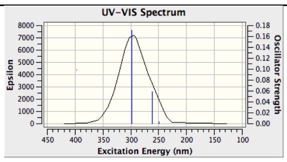  | 298.7 [0.17] |

It is clear from the computed spectra of the aromatic bicyclic compounds that there is a distinction between those that are experimentally found to be viable substrates in the reaction and those that are not. Somewhere in the range of 272 to 282 nm lies the point at which the excitation becomes infeasible. A similar correlation can be found in terms of the calculated triplet state energies ,which would impact the photocatalysed energy transfer process; somewhere between 59 and 61 kcal/mol lies the point at which the triplet state becomes inaccessible.

The computed spectra of the cycloaddition adducts reveal that these species generally absorb at a lower wavelength than the unreacted aromatics. Their excited triplet states are also significantly lower in energy than those of the unreacted aromatics. Both of these findings seem surprising given the usual expectations that more extended aromatic systems are likely to be more photochemically active and to have lower energy excited states. To explore this effect, the HOMO and LUMO energies of the ground state as well as of the two SOMOs in the triplet states were extracted from the calculations for some of the key species. These are given in atomic units in Table SN2.2.

***Table SN2.2*** *Energies of key orbitals in the species indicated (in Hartrees).*

| **Species** | **HOMO energy** (ground state) | **LUMO energy** (ground state) | **SOMO energy 1** (triplet) | **SOMO energy 2** (triplet) |
|---|---|---|---|---|
| 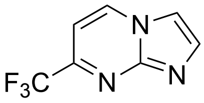 **S9** | -0.293 | -0.041 | -0.319 | -0.198 |
| 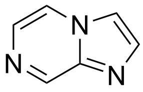 **SI3** | -0.284 | -0.020 | -0.304 | -0.186 |
| 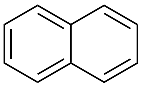 **SI5** | -0.269 | -0.013 | -0.281 | -0.158 |
| 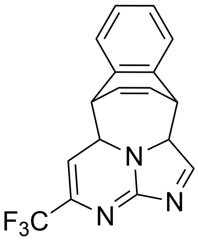 *endo*-**D** | -0.267 | -0.031 | -0.286 | -0.183 |

| | | | | |
|---|---|---|---|---|
| 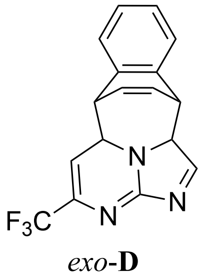 *exo*-**D** | -0.268 | -0.032 | -0.286 | -0.183 |

The orbital energies reveal that the adducts have the surprising property that, whereas their HOMOs are higher in energy (as might be expected given the loss of aromaticity), their LUMOs are of comparable energy. This leads to a narrowed HOMO-LUMO gap and explains the longer wavelength absorption and lower singlet-triplet energy difference. In order to understand this effect, the HOMO and LUMO for the ground state and two SOMOs for the triplet were visualized for **S9** and **D** (Table SN2.3). This reveals that a defining feature of both the LUMO and the higher energy SOMO that corresponds to the LUMO after excitation is the presence of an interaction through the sigma framework that permits the LUMO to benefit from a more extended conjugation than might otherwise be expected.

***Table SN2.3*** *Visualisations of key molecular orbitals for the species indicated.*

| **Species** | **HOMO** (ground state) | **LUMO** (ground state) | **SOMO 1** (triplet) | **SOMO 2** (triplet) |
|---|---|---|---|---|
| 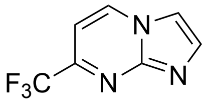 **S9** | 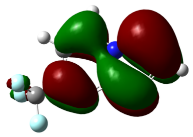 | 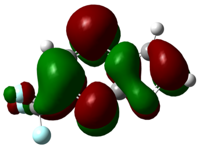 | 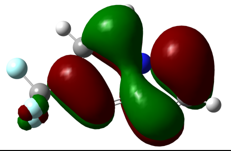 | 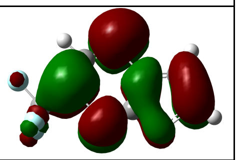 |
| 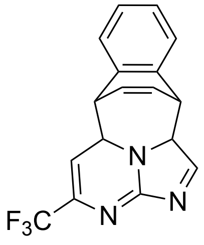 *endo*-**D** | 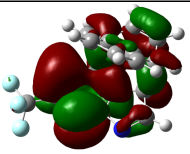 | 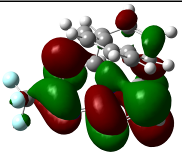 | 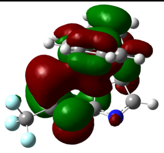 | 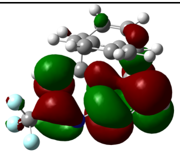 |
| 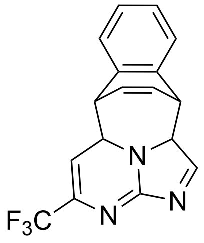 *exo*-**D** | 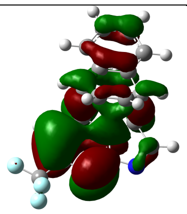 | 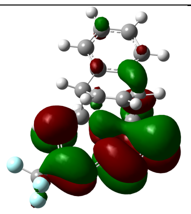 | 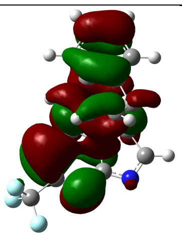 | 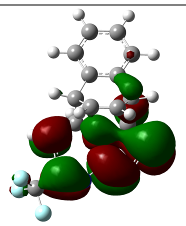 |

**Intermediate adducts and products**

One of the key observations to explain is the absence of alternative products, particularly those for alternative cycloadditions, such as [4+2] or [2+2]. The first step to exploring this was to compute the energies of alternative species to identify those that need to be considered and to quantify any influence that overall reaction thermodynamics might exert on the reaction outcome. In Table SN2.4, the energies of each species relative to separate reactants is given (N.B. oxidation products not included). This reveals that the thermodynamically preferred species are those arising from [4+2] cycloadditions (**C** and **SI6**), with those from [4+3] (**D**) and [2+2] (**SI9-12**) being disfavoured.

***Table SN2.4*** *Free energy of reaction to form the indicated species (for each stereoisomer) from ground state reactants.*

| Type | Adduct | $\Delta G_{reaction}$(endo) / kcal/mol | $\Delta G_{reaction}$(exo) / kcal/mol |
|---|---|---|---|
| [4+3] (3,5-imidazo-pyrimidine/1,4-naphthalene) | 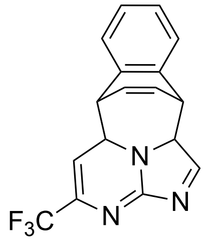  **D** | +35.1 | +35.4 |
| [4+2] (5,6-imidazo-pyrimidine/1,4-naphthalene) | 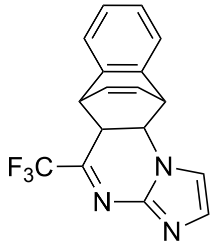  **C** | +27.7 | +27.4 |
| [4+2] (2,3-imidazo-pyrimidine/1,4-naphthalene) | 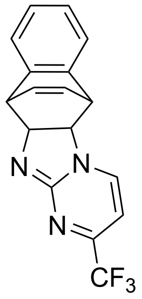  **SI6** | +29.9 | +29.6 |
| [2+2] (5,6-imidazo-pyrimidine/1,2-naphthalene) | 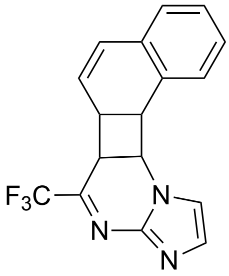  **SI9** | +35.4 | +34.3 |
| [2+2] (5,6-imidazo-pyrimidine/1,2-naphthalene) | 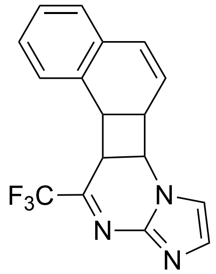  | +37.5 | +35.6 |

| | | | |
|---|---|---|---|
| | SI10 | | |
| [2+2] (2,3-imidazo-pyrimidine/1,2-naphthalene) | 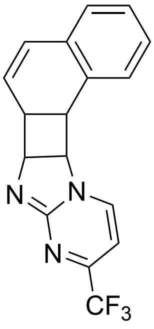  **SI11** | +41.3 | +40.7 |
| [2+2] (2,3-imidazo-pyrimidine/1,4-naphthalene) | 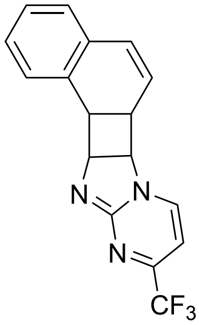  **SI12** | +42.7 | +40.2 |

Among the oxidation products, by contrast (Figure SN2.1), there is a preference for the [4+3] reaction product **F** compared to either of the [4+2] possibilities (**SI7-8**) and a very marked preference compared to the lowest energy [2+2] product **SI13**.

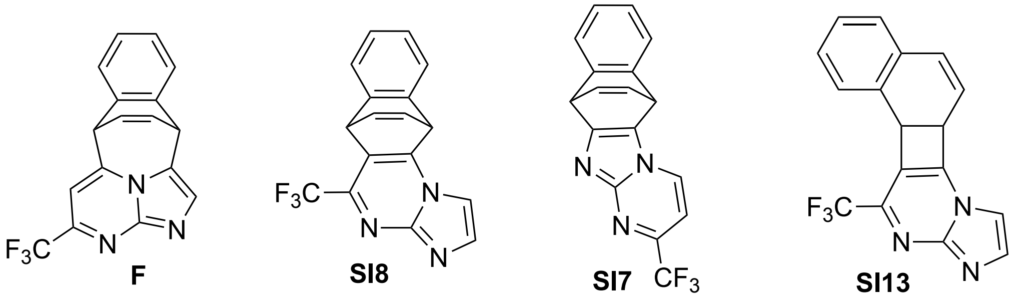

0.0 +10.8 +17.5 +25.5

**Figure SN2.1** Relative free energies (in kcal/mol) of oxidised products relative to that arising from a [4+3] cycloaddition.

The origins of these differences can be found in the geometries of the various species, shown in Figure SN2.2. The C-C bond lengths of the two bonds formed between the two reactants are shown and indicate that an increase in steric clash with the $CF_3$ group in **SI8** upon oxidation is one contributor disfavouring the oxidation of **C**. Meanwhile, the bond angle changes between the reactant and oxidized products reveal significant angle strain in the product **SI7** or **SI8** arising from [4+2]-cycloadditions whereas in the [4+3]-derived product **F** there is essentially no change from the angles to the CH bonds in the reactant. After restoration of aromaticity in both rings after oxidation, there is a much reduced strain in the [4+3]-derived product compared to that in the [4+2]-derived products. This is likely a dominant factor in the change in selectivity upon going from cycloadducts to oxidized products. Another key factor is that the [4+2] cycloadducts involve disruption of aromaticity of only one ring whereas the [4+3] disrupts both rings. Upon oxidation, the [4+3] can benefit from partial rearomatization (see below).

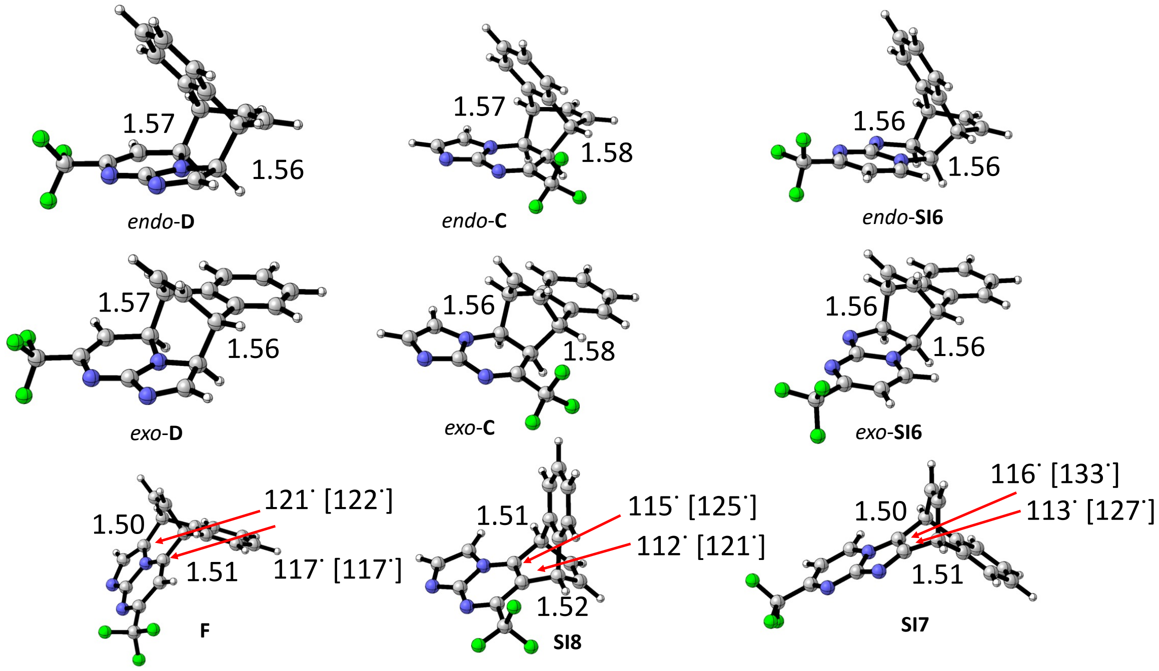

**Figure SN2.2** Geometries of key species: Left – [4+3] addition adducts **D** and oxidized product **F**, Centre – [4+2] addition on the 6-ring adducts **C** and oxidized product **SI8**, Right – [4+2] addition on the 5-ring adduct **SI6** and oxidized product **SI7**. Key distances are indicated in Å. Angles shown are the internal angle at the atom indicated by the arrow and is between the attached exo-cyclic carbon and the next atom around in the reactive aza-aromatic towards the other exo-cyclic bond. In brackets are the corresponding angle to the C-H bond in the reactant aza-aromatic.

#### Reaction potential energy surface

Once the reactant aromatic has been excited to the triplet state, it can then undergo a sequence of reactions, as outlined in the main text. The first step is an addition to the other coupling partner to form an open-chain diradical intermediate **B** or **SI14**. This can yield a number of regioisomers each of which can exist as two relative stereochemical arrangements. The free energy changes corresponding to the barrier to formation and the reaction free energy change upon addition to naphthalene of the triplet diradical $^{3}$**A** are given in Table SN2.5 for the four key diradicals. In each case, higher energy conformations were also explored and their free energies also given.

***Table SN2.5*** *Free energy barriers and reaction energies for the addition of triplet diradical to naphthalene. Energies are given in kcal/mol relative to the triplet excited state.*

| Type | Species | Lowest $\Delta G^{\ddagger}$ (kcal/mol) | Alternative conformations $\Delta G^{\ddagger}$ (kcal/mol) | Lowest $\Delta G_{reaction}$ (kcal/mol) | Alternative conformations $\Delta G_{reaction}$ (kcal/mol) |
|---|---|---|---|---|---|
| 1,5-addition (endo) | 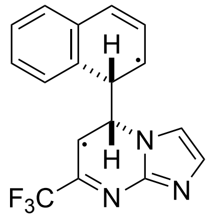 *endo*-$^{3}$**B** | 16.8 | 18.9, 20.5 | +0.8 | +2.3, +2.9 |
| 1,3-addition (endo) | 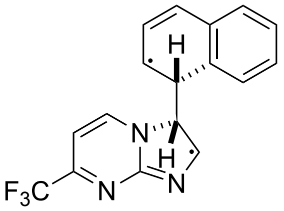 *endo*-**SI14** | 19.8 | 20.5, 21.3 | +4.9 | +6.0, +6.4 |
| 1,5-addition (exo) | 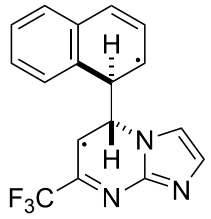 *exo*-$^{3}$**B** | 17.3 | 20.5, 21.1 | +0.6 | +1.5, +4.1 |
| 1,3-addition (exo) | 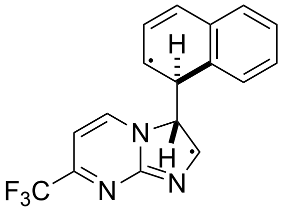 *exo*-**SI14** | 19.5 | 20.8, 22.3 | +4.8 | +5.4, +7.2 |

This reveals that the kinetically and thermodynamically dominant products would arise from addition at the 6-membered ring of the reactant triplet diradical to give $^{3}$**B**; only a small stereochemical preference might be expected. In addition, only trace amounts of the alternative regio-isomers **SI14** might be expected. Critically, all reactions are readily reversible with equilibria expected to be towards separated reactants and thus these initial addition products are unlikely to be observable.

The possibility of the formation of diradical adducts on the singlet surface (i.e. without the requirement for inter-system crossing) was also considered but optimization of the low energy triplet diradicals as singlets usually led to their fragmentation to reactants. This suggests that open-chain singlet diradicals are unlikely to be important contributors to the reaction as they will have either no or very short lifetimes. Those singlet diradicals that could be obtained had energies that were identical to those of the triplet diradical (difference < 0.1 kcal/mol).

The fate of the initial addition products was then considered (see Figure SN2.3). In blue are shown steps leading to [4+3]-type adducts; in orange those leading to [4+2] adducts on the 6-membered ring; and in green those leading to the [4+2] adducts on the 5-membered ring. Each species is labelled as either T for triplet, S for closed-shell singlet or S/T for points at which singlet and triplet are degenerate. These

latter points are not geometry optimized but are obtained by scanning the forming C-C bond in 0.1 Å steps. In addition, their electronic energies are computed relative to that for the lowest energy [4+3] adduct and this difference is added to the free energy of that adduct in order to obtain the energy given. As a consequence, the S/T-labelled points come with more uncertainty in their free energies than other points although this is unlikely to affect the interpretation of these reaction free energy schemes.
Beginning at the left of each scheme, the lowest barrier process for each diradical is to undergo closure through a surface-crossing point leading to a [4+2] adduct. As mentioned previously, this species is readily re-excited to the triplet state which can re-open through a low barrier to regenerate $^{3}$**B**. Alternatively, the next lowest free energy path involves the initial addition product cyclizing to give the [4+3] adduct $^{1}$**D** which likewise can be re-excited to yield $^{3}$**D** that has a low barrier to re-opening to also regenerate $^{3}$**B**. Alternatively, $^{3}$**D** can open in the alternative direction to yield the higher energy diradical **SI14**. This diradical could also be obtained from initial addition and readily cyclises to give $^{1}$**SI16**. These two schemes suggest that the dominant products both kinetically and thermodynamically should be the various [4+2] adducts and not the observed [4+3] adduct. However, it is also clear that while the exposure to light continues, the cycloadducts **C**, **D** and **SI6** will be selectively re-activated and recycled to either separated reactants or to diradical addition products. This likely explains why none of these intermediate adducts are observed in the reaction mixture.

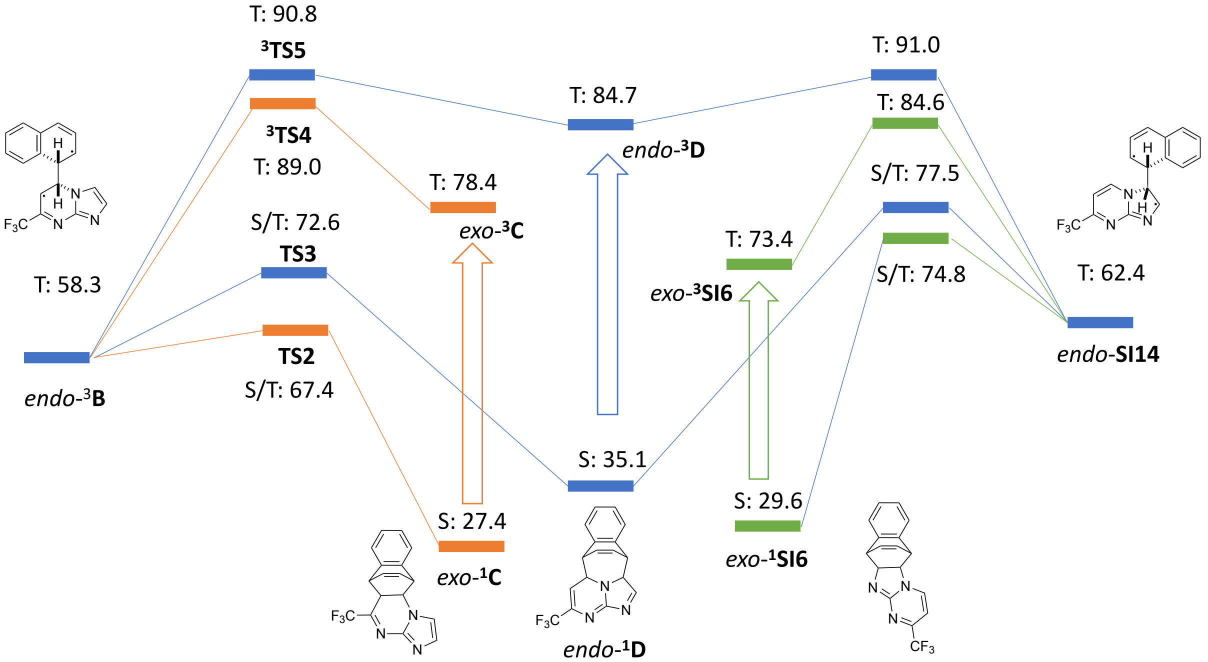

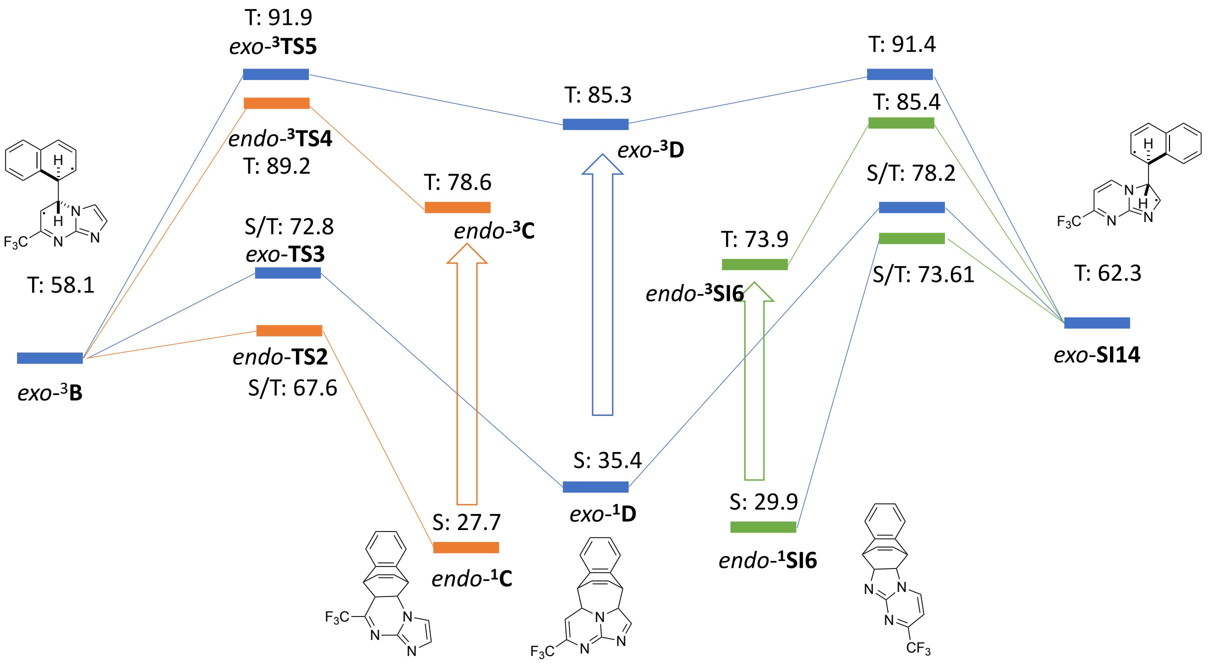

**Figure SN2.3** Free energies of species relative to separated, ground state reactants aza-aromatic and naphthalene. T indicates free energy of a triplet species, S a singlet and S/T a point at which singlet and triplet are degenerate. Arrows represent likely reactivation. Blue – [4+3]; Orange – [4+2] on the 6-ring; Green – [4+2] on the 5-ring. Top scheme: species leading to the *endo*-[4+3] adduct *endo*-**D**; Bottom scheme: species leading to the exo-[4+3] adduct *exo*-**D**.

The reaction outcome in these circumstances would depend on the relative rate of any reactions that permit these adducts to be trapped. Hence, the next process to be studied was that of hydrogen abstraction by any radicals present in the system; when tBuOOAc is present, it seems reasonable that tBuO radicals might perform this function. The barrier height for the radical abstraction of each candidate hydrogen by tBuO radical was computed (Table SN2.6).

***Table SN2.6*** *Barrier height and reaction energy for abstraction of a hydrogen radical by tBuO radical for the hydrogens indicated by the letters A and B.*

| | $\Delta G^{\ddagger}$(HA) in kcal/mol | $\Delta G^{\ddagger}$(HB) in kcal/mol | $\Delta G_{rxn}$(HA) in kcal/mol | $\Delta G_{rxn}$(HB) in kcal/mol |
|---|---|---|---|---|
| 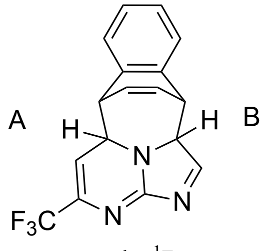  endo-$^1$**D** | 11.3 | 13.5 | -32.6 | -37.1 |

| | | | | |
|---|---|---|---|---|
| 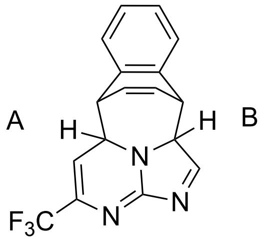 exo-$^{1}$**D** | 11.7 | 12.9 | -33.1 | -37.2 |
| 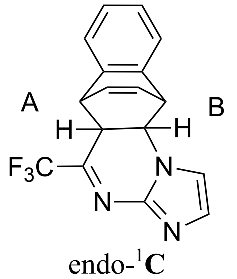 endo-$^{1}$**C** | 15.2 | 14.4 | -24.9 | -14.9 |
| 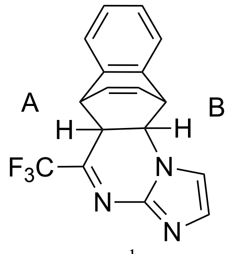 exo-$^{1}$**C** | 16.0 | n.c. | -24.6 | -14.7 |
| 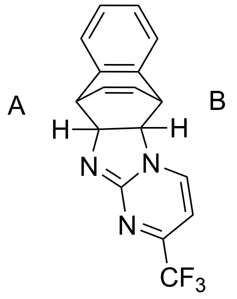 endo-$^{1}$**SI6** | 13.0 | n.c. | -22.9 | -14.1 |
| 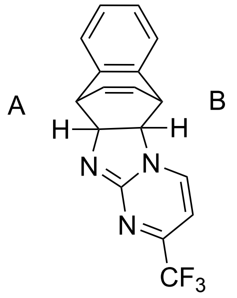 exo-$^{1}$**SI6** | 12.9 | 13.6 | -22.8 | -13.7 |

n.c. = not computed.

Several things are clear from these computed radical abstractions. The first is that the influence of stereochemistry is modest and so even where the two transition states have not yet been obtained, the barriers can be imputed. The second and most important is that there is a clear preference for abstraction from $^{1}$**D** and the radicals that arise from this process $^{2}$**SI15** and $^{2}$**SI16** are the most stable ones. This likely explains why the [4+3] product **F** is observed and not the **SI7** or **SI8**; the precursors to **F** are formed more slowly but irreversibly oxidised more quickly.

It is interesting to note that for the [4+3] cycloadducts, the radicals formed by abstracting hydrogen, $^{2}$**SI15** and $^{2}$**SI16**, feature cyclic delocalization of electrons that is not possible in the other regioisomeric cycloadducts. Combined with the decrease in geometric strain on going from cyclo-adduct towards oxidized product, this likely explains the preference for the formation of **F**. This can be visualized by examining the bond lengths in the ring at various stages of the reaction. For the **D**, the bonds in the ring that contains the radical (Figure SN2.4) are significantly changed towards more closely resembling those in the aromatic (oxidized) product. By contrast, for the two [4+2] cycloadducts (**C** in Figure SN2.5 and **SI6** in Figure SN2.6), the bonding patterns resemble those expected for a radical stabilized only by delocalization. This often comes at the cost of a small disruption of the extant aromaticity in the ring that was not involved in the cycloaddition (Figure SN2.4-6).

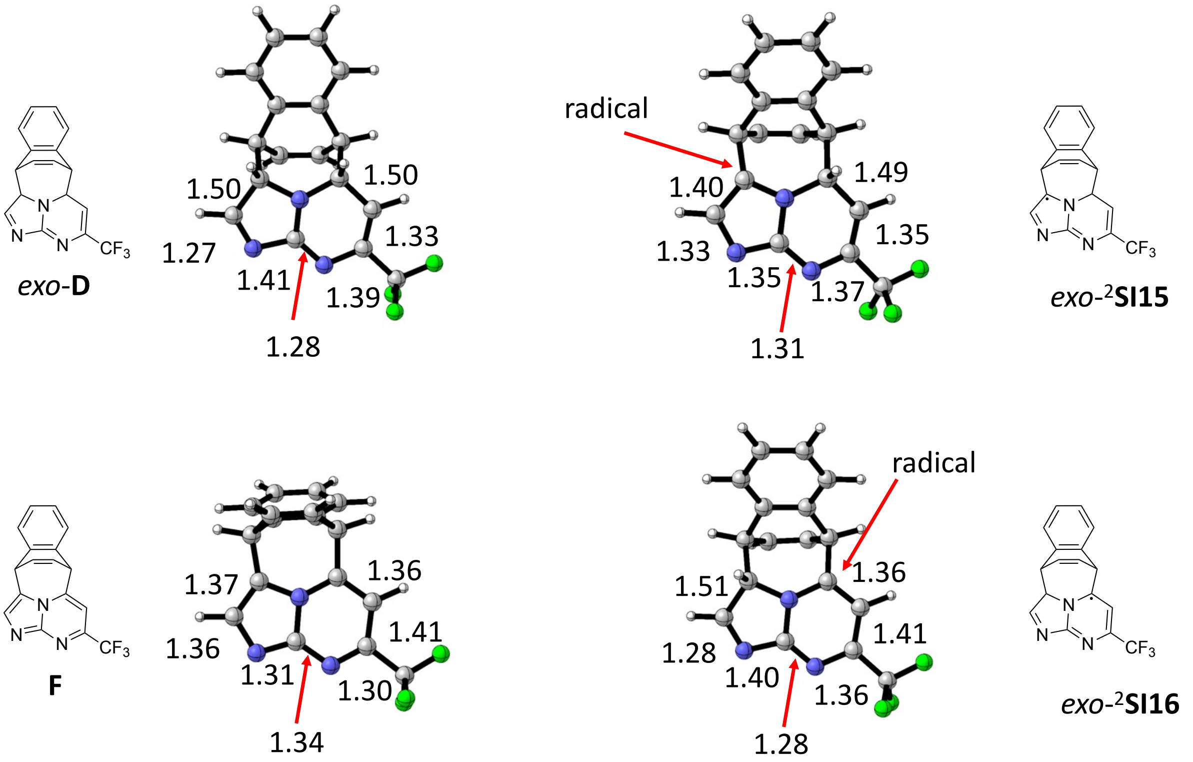

**Figure SN2.4.** Key bond lengths (in Å) for the species shown that are relevant to the [4+3] reaction. Only the *exo* isomer is shown because this geometry is clearer to visualize. The same general features are present in the *endo* equivalent (where relevant).

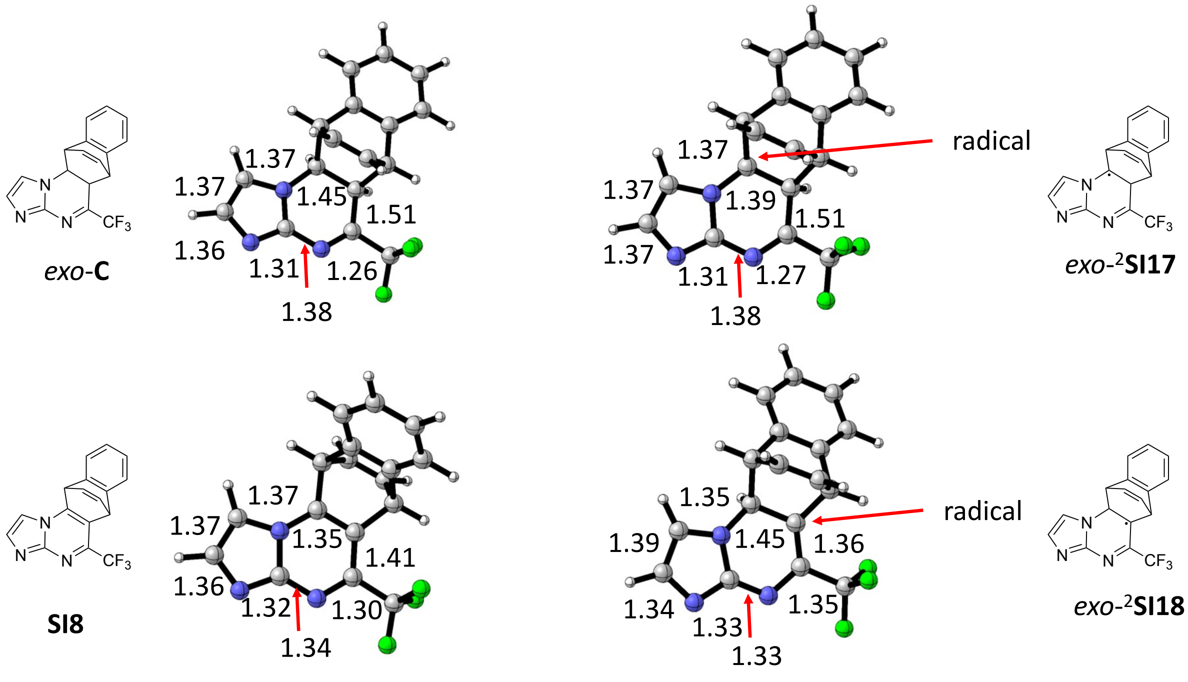

**Figure SN2.5** Key bond lengths (in Å) for the species shown that are relevant to the [4+2] reaction in the 6-ring. Only the *exo* isomer is shown because this geometry is clearer to visualize. The same general features are present in the *endo* equivalent (where relevant).

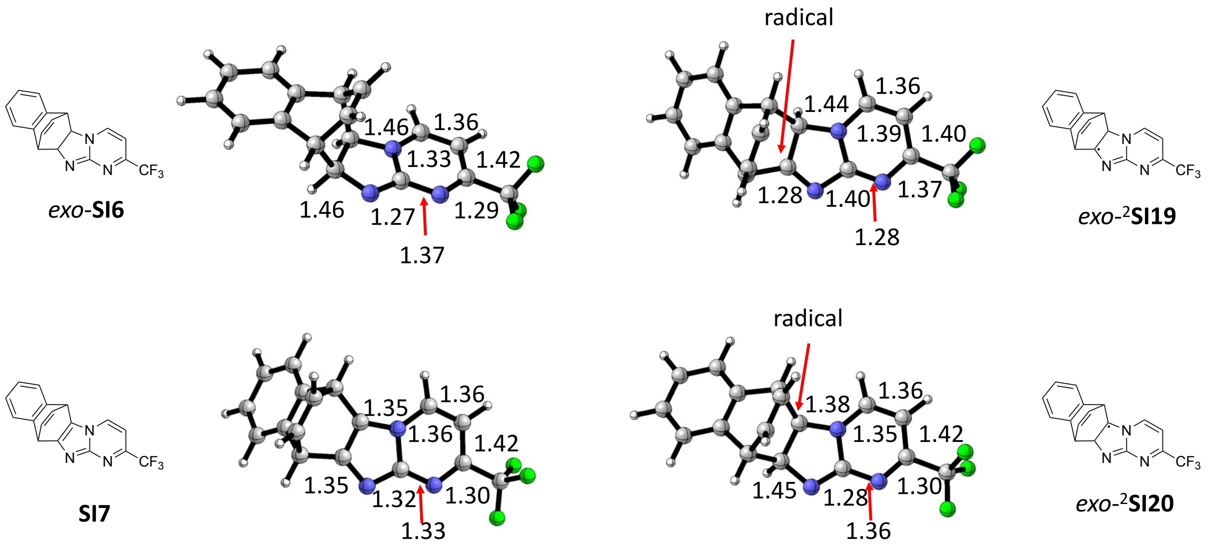

**Figure SN2.6** Key bond lengths (in Å) for the species shown that are relevant to the [4+2] reaction in the 5-ring. Only the *exo* isomer is shown because this geometry is clearer to visualize. The same general features are present in the *endo* equivalent (where relevant).

Finally, the radicals obtained by hydrogen abstraction from the cycloadducts must be converted to the observed products. There are a number of possible mechanisms for this process. One is that the radical products are trapped by AcO or other radicals. Another is that the radical products undergo electron transfer to species such as AcO radicals to obtain a readily deprotonated cation and acetate anion or a second tBuO radical might abstract the second hydrogen. All of these have been computed to be feasible but the details of this step will depend on the composition of the solution at the time that each radical is generated. The computed free energy change for a number of candidate reactions for the two kinetically most accessible radicals are given in Figure SN2.7.

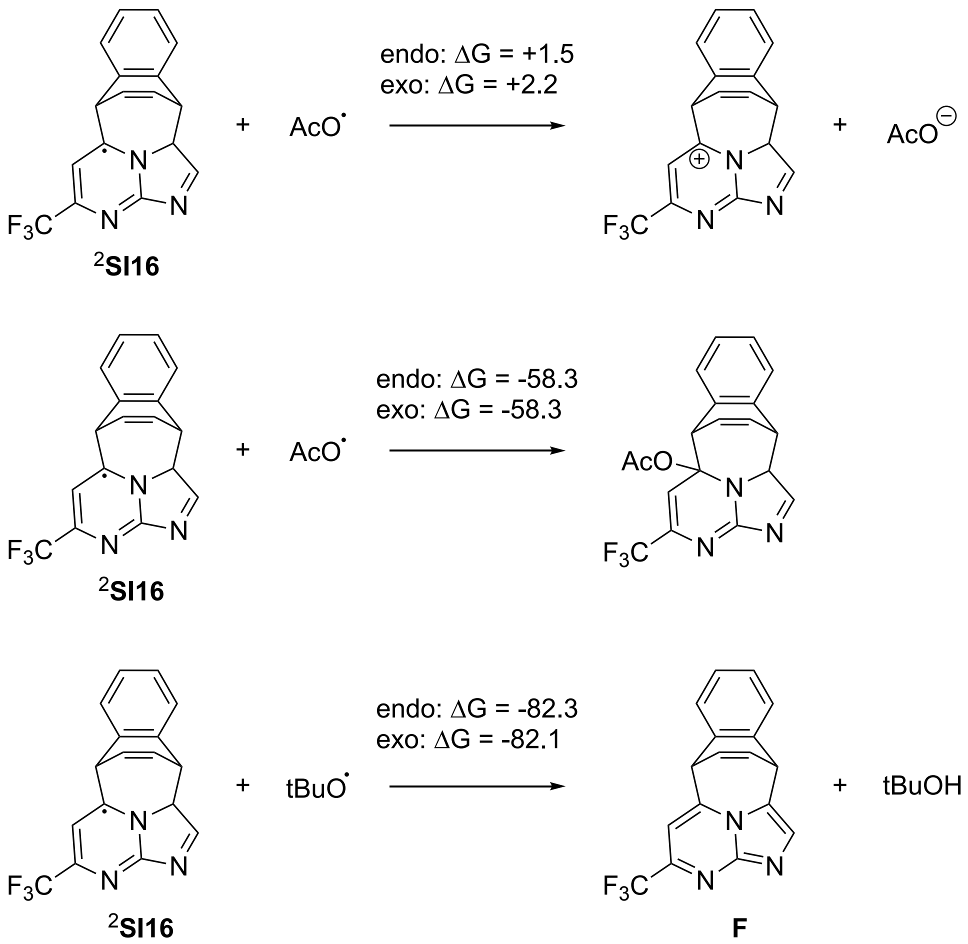

**Figure SN2.7** Processes that could explain the formation of the observed oxidation products from the initially formed radicals.

#### Concerted cycloadditions and singlet diradicals

A concerted cycloaddition transition state could be obtained for each of the [4+2] cycloaddition reactions and for one isomer of the [4+3] cycloaddition. These are high barrier processes (Table S12). The interplay between concerted and stepwise (via diradicals) processes in cycloadditions has long been a subject of interest.(K. Black, P. Liu, C. Doubleday and K. N. Houk, *Proc. Natl. Acad. Sci. USA* **2012**, *109*, 12860-12865). It is curious that when attempting to obtain transition states for singlet diradical formation and cyclization for the [4+2] processes, no cyclization transition states could be identified. By contrast, such barriers could be obtained for the [4+3] processes. These are significantly higher in energy than those for the [4+2]. These results, combined with the weakly bound nature of singlet diradical addition products, suggest that any reactions on the singlet excited state surface are also most likely to yield [4+2] cycloadducts.

***Table SN2.7*** *Free energy barriers for concerted cycloadditions or for closure of singlet diradicals.*

| Species | $\Delta G^{\ddagger}$(endo) in kcal/mol | $\Delta G^{\ddagger}$(exo) in kcal/mol |
|---|---|---|
| 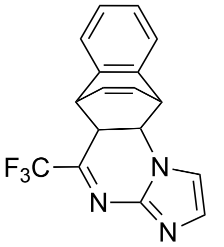<br>**C**<br>Concerted TS | 51.3 | 51.0 |
| 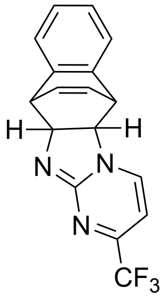<br>**SI6**<br>Concerted TS | 55.8 | 55.9 |
| 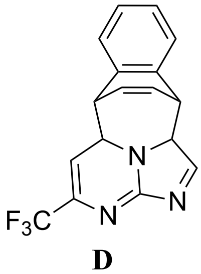<br>**D**<br>Concerted TS | 68.3 | |
| 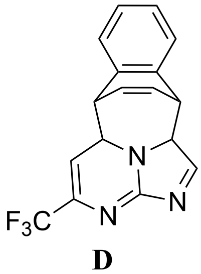<br>**D** | Closing at 5-ring: 62.5<br>($\langle S^2 \rangle = 0.095$)<br>Closing at 6-ring: 64.7<br>($\langle S^2 \rangle = 0.089$) | Closing at 5-ring: 68.8<br>($\langle S^2 \rangle = 0.076$)<br>Closing at 6-ring: 67.2<br>($\langle S^2 \rangle = 0.090$) |

| |
|---|
| Singlet diradical ring closure TS |

#### Failed incorporation of Npa into protein models via the orthogonal MmNpaRS/tRNAPyl pair

Two systems have been previously suggested for Npa insertion. First, following analysis of the crystal structure of the homologous TyrRS from *Bacillus stearothermophilus*, Schultz and co-workers saturatively mutated and selected a five-point mutant SS12-TyrRS (Y32L-D158P-I159A-L162Q-A167V) variant of the *M. jannaschii* TyrRS. This *Mj* $tRNA_{CUA}^{Tyr}$-SS12TyrRS pair successfully facilitated introduction of Npa into mouse dihydrofolate reductase (DHFR; via an amber stop codon at site 163) through expression in *E. coli*.[1,2] Then, in a more recent study, a system with potential application to both *E. coli* and mammalian cell expression, has been proposed based on a five point mutant of *M. mazei* PylRS in (A302T-N346G-C348T-V401I-W417Y).[3] With this *Mm* $tRNA_{CUA}^{Pyl}$-'Nap'RS pair, introduction of Npa into sperm-whale myoglobin (via an amber stop codon at site 4) was enabled, again, through expression in *E. coli*. Cellular fluorescence in HeLa also suggested low (5.6% of reference Boc-Lys with PylRS) but significant expression of GFP where Tyr182 is replaced with Npa.

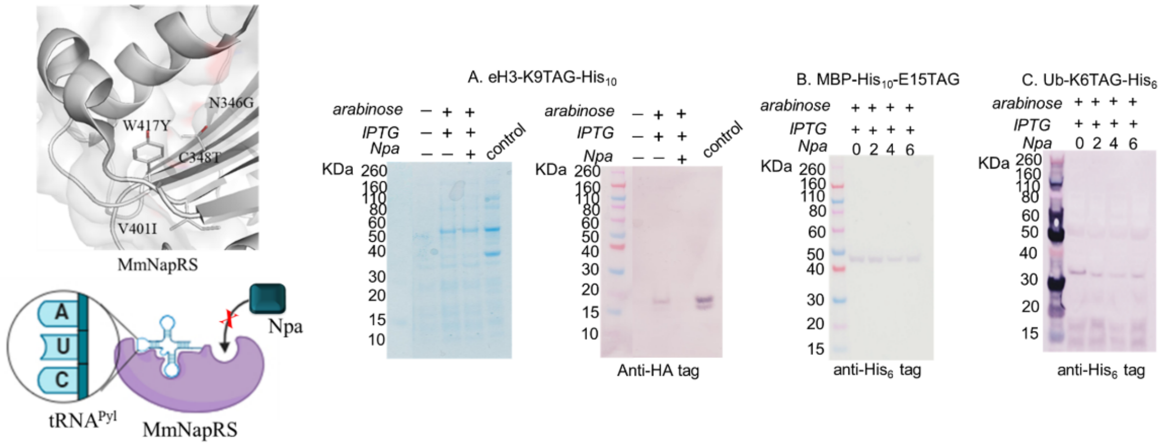

**Figure SN3.1:** Failed incorporation of Npa into different protein models via orthogonal MmNapRS/$tRNA^{Pyl}$ pair.[3] SDS-PAGE and western blotting experiments for the comparation of the expression of (A) protein H3-K9TAG, (B) MBP-E15TAG or (C) Ub-K6TAG in the presence of increasing amounts of Npa. The successfully incorporated iodophenylalanine onto K9 site of the same eH3 protein with PylRS(N346A/C348A)/$tRNA^{Pyl}$ ref[4] as used as positive control.

To evaluate both strategies in an unbiased manner, we constructed new plasmids based on the increased efficiency pEVOL vector systems[5] and used this to evaluate incorporation via an amber stop codon at site 9 in epitope (HA-FLAG-C-terminus)-tagged human H3 protein (eH3). Crucially, we were unable to see incorporation of Npa into epitope-tagged human histone H3 (eH3) with the *Mm* $tRNA_{CUA}^{Pyl}$-'Npa'RS pair (**Figure SN3.1**). The *Mm* $tRNA_{CUA}^{Pyl}$-'Nap'RS pair also failed in our hands for attempted incorporation into other amber codon-bearing targets (maltose-binding protein and ubiquitin); notably Western blot analysis suggested reduction of a background expression upon addition of Npa to culture in these systems.

Further codon optimization of the PylRS gene, which was intended to enhance its expression in *E. coli*., did not lead to a functional enzyme.

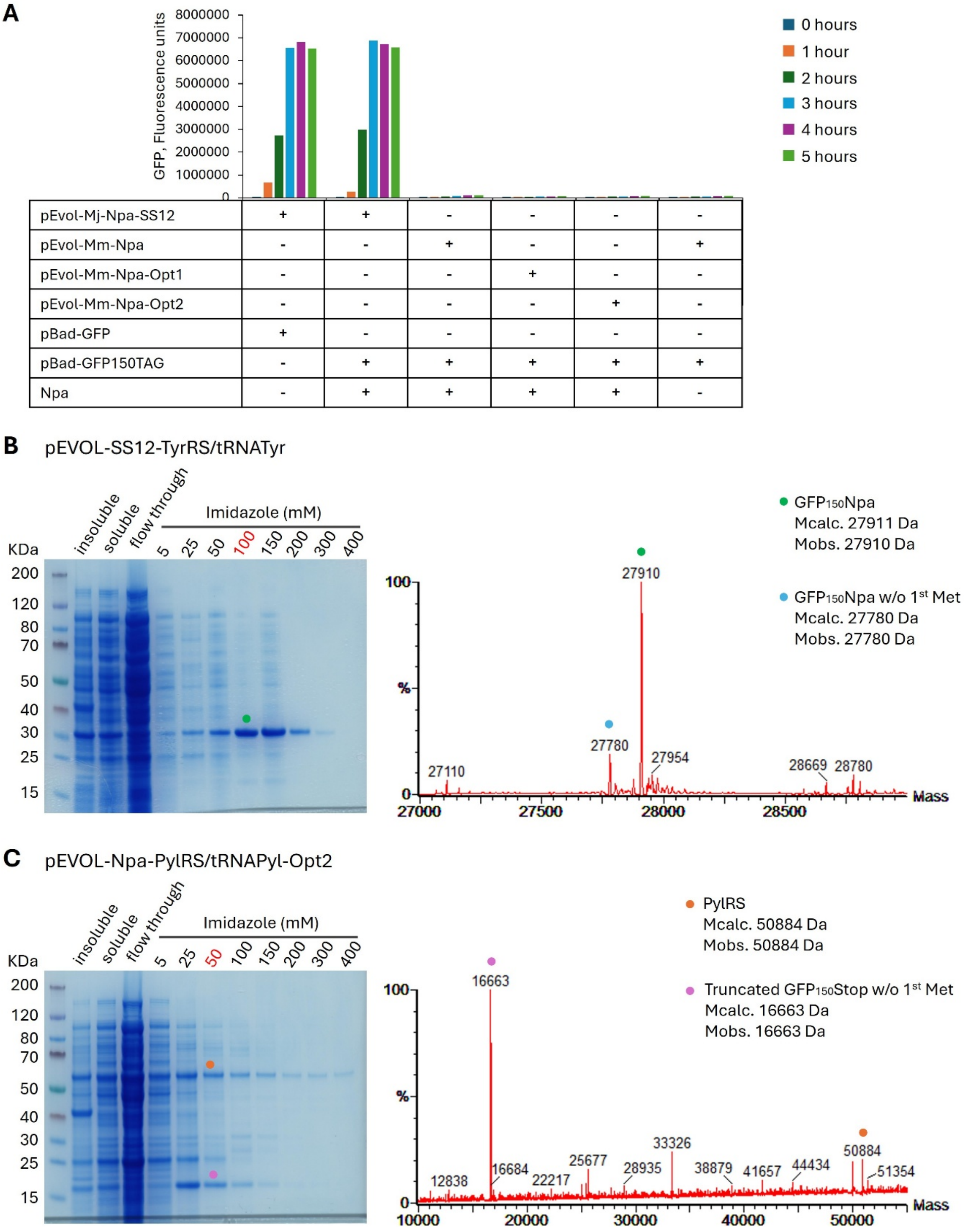

**Figure SN3.2. Analysis of functionality of Npa-specific aminoacyl tRNA-synthetases used in this study through incorporation of Npa into GFP reporter protein**. **A**) Amber codon suppression efficiency was tested using *Mm*-Npa-PylRS constructs generated in this study and compared to *Mj*-Npa-TyrRS-SS12. Three versions of codon optimized *Mm*-Npa-PylRS (including one previously published by the L. Wang group [ref. 44], see SI for sequences) were evaluated. GFP fluorescence was measured hourly over a 5-hour period. All *Mm*-Npa-PylRS variants failed to incorporate Npa into GFP at detectable levels, while *Mj*-Npa-TyrRS-SS12 supported efficient Npa incorporation, comparable to the positive control expressing wild-type (WT) GFP. **B**) Efficient site-specific incorporation of Npa at position 150 of GFP protein (green dot) was achieved using pEVOL-SS12-TyrRS/tRNA$^{Tyr}$ plasmid, which encodes the SS12-TyrRS and tRNA$^{Tyr}$ pair. Coomassie-stained SDS-PAGE (left) of the fractions collected after Ni-NTA purification shows protein band corresponding to the expected GFP mass. LC-MS analysis (right) of fraction eluted with 100mM imidazole confirms Npa incorporation, with a deconvoluted protein mass matching GFP with Npa. A minor mass corresponding to GFP lacking the initial methionine (blue dot) was also observed. **C**) Failed incorporation of Npa into GFP protein using the pEVOL-Npa-PylRS/$tRNA^{Pyl}$ plasmid encoding the *Mm*-Npa-PylRS-Opt2. Coomassie-stained SDS-PAGE (left) of the fractions collected after Ni-NTA purification shows protein bands that do not correspond to the expected GFP mass and display weak binding to the Ni-NTA resin. LC-MS analysis (right) of the fraction eluted with 50mM imidazole revealed two prominent masses: 16,663 Da and 50,884 Da. The 16,663 Da peak corresponds to a truncated $GFP_{150TAG}$ product, indicating translation termination at the amber stop codon and thus confirming lack of suppression. The 50,884 Da mass matches the *Mm*-Npa-PylRS-Opt2 protein, which was expressed at high levels. Despite this, no full-length GFP was detected, suggesting that Mm-Npa-PylRS-Opt2 is either non-functional or lacks specificity to Npa under these conditions, and thus cannot facilitate its incorporation into the protein of interest.

Nucleotide sequences used in this study. Codons representing amino acids in the evolved amino acid binding pocket are highlighted in yelow:

Mm-Npa-PylRS [44] nucleotide sequence:
ATGGATAAAAAGCCTCTGAACACTCTGATTTCTGCGACCGGTCTGTGGATGTCCCGCACCGGCACC
ATCCACAAAATCAAACACCATGAAGTTAGCCGTTCCAAAATCTACATTGAAATGGCTTGCGGCGAT
CACCTGGTTGTCAACAACTCCCGTTCTTCTCGTACCGCTCGCGCACTGCGCCACCACAAATATCGC
AAAACCTGCAAACGTTGCCGTGTTAGCGATGAGGACCTGAACAAATTCCTGACCAAAGCTAACGA
GGATCAGACCTCCGTAAAAGTGAAGGTAGTAAGCGCTCCGACCCGTACTAAAAAGGCTATGCCAA
AAAGCGTGGCCCGTGCCCCGAAACCTCTGGAAAACACCGAGGCGGCTCAGGCTCAACCATCCGGT
TCTAAATTTTCTCCGGCGATCCCAGTGTCCACCCAAGAATCTGTTTCCGTACCAGCAAGCGTGTCTA
CCAGCATTAGCAGCATTTCTACCGGTGCTACCGCTTCTGCGCTGGTAAAAGGTAACACTAACCCGA
TTACTAGCATGTCTGCACCGGTACAGGCAAGCGCCCCAGCTCTGACTAAATCCCAGACGGACCGTC
TGGAGGTGCTGCTGAACCCAAAGGATGAAATCTCTCTGAACAGCGGCAAGCCTTTCCGTGAGCTG
GAAAGCGAGCTGCTGTCTCGTCGTAAAAAGGATCTGCAACAGATCTACGCTGAGGAACGCGAGAA
CTATCTGGGTAAGCTGGAGCGCGAAATTACTCGCTTCTTCGTGGATCGCGGTTTCCTGGAGATCAA
ATCTCCGATTCTGATTCCGCTGGAATACATTGAACGTATGGGCATCGATAATGATACCGAACTGTC
TAAACAGATCTTCCGTGTGGATAAAAACTTCTGTCTGCGTCCGATGCTGACGCCGAACCTGTACAA
CTATCTGCGTAAACTGGACCGTGCCCTGCCGGACCCGATCAAAATTTTCGAGATCGGTCCTTGCTA
CCGTAAAGAGTCCGACGGTAAAGAGCACCTGGAAGAATTCACCATGCTGGGCTTCACCCAGATGG
GTAGCGGTTGCACGCGTGAAAACCTGGAATCCATTATCACCGACTTCCTGAATCACCTGGGTATCG
ATTTCAAAATTGTTGGTGACAGCTGTATGGTGTATGGCGATACGCTGGATGTTATGCACGGCGATC
TGGAGCTGTCTTCCGCAATCGTGGGCCCAATCCCGCTGGATCGTGAGTGGGGTATCGACAAACCTT
ACATCGGTGCGGGTTTTGGTCTGGAGCGTCTGCTGAAAGTAAAACACGACTTCAAGAACATCAAA
CGTGCTGCACGTTCCGAGTCCTATTACAATGGGATTTCAACCAATCTATAA
Mm-Npa-PylRS[44] nucleotide sequence codon optimized for *E. coli* expression using Basebuddy codon optimization tool (https://basebuddy.lbl.gov/), Mm-NpaRS-Opt1:
ATGGATAAAAAACCACTCAATACATTAATAAGTGCGACCGGTTTATGGATGTCACGCACCGGCAC
CATCCACAAAATCAAACACCATGAAGTTAGCCGCAGTAAAATCTATATTGAAATGGCCTGCGGCG
ATCACTTAGTTGTCAACAACTCACGCTCGTCTCGCACAGCGCGCGCATTGCGCCATCACAAATATC
GCAAAACCTGTAAACGATGCCGTGTTAGCGATGAGGACCTGAACAAATTTCTGACAAAAGCGAAT
GAAGATCAGACCTCCGTAAAAGTGAAGGTAGTAAGCGCTCCGACACGTACTAAAAAGGCTATGCC
CAAAAGTGTGGCCCGTGCCCCGAAACCCCTGGAAAACACAGAAGCGGCGCAGGCCCAACCATCCG
GCTCTAAATTTTCTCCGGCGATCCCCGTGTCCACGCAAGAAAGTGTTTCAGTCCCAGCAAGCGTGT
CAACCAGTATTAGCAGCATTTCGACCGGGGCTACCGCTTCTGCGCTGGTAAAAGGCAATACTAACC
CGATTACTAGCATGTCGGCACCGGTCCAGGCCAGCGCCCCAGCACTAACTAAATCCCAGACGGAC
CGTCTGGAGGTGCTTCTGAACCCAAAGGATGAAATTTCGCTGAACAGCGGCAAGCCTTTCCGGGA
ACTGGAAAGCGAGTTACTGTCTCGTCGGAAAAAGGATTTGCAACAGATCTACGCGGAAGAACGCG
AAAACTATCTGGGAAAGCTGGAGCGCGAAATTACGCGCTTTTTCGTGGATCGCGGTTTTCTGGAGA
TCAAATCTCCGATTCTTATTCCGCTGGAATACATTGAAAGAATGGGCATCGATAATGATACCGAAT
TGTCGAAACAGATCTTCCGTGTCGATAAAAATTTCTGTCTCCGACCGATGCTCACGCCGAATCTTT
ATAACTATCTGCGTAAACTGGACCGTGCCCTGCCGGACCCGATCAAAATTTTCGAGATTGGTCCTT
GCTACCGTAAAGAGTCAGACGGTAAAGAACATTTGGAAGAATTTACCATGTTGGGCTTCACCCAG
ATGGGTAGCGGGTGCACGAGGGAAAACCTTGAATCCATTATCACCGACTTTCTCAATCATCTGGGG

ATTGATTTTAAAATTGTTGGTGACAGCTGTATGGTGTATGGCGATACGCTGGATGTTATGCATGGC
GATTTAGAGTTAAGTAGTGCAATCGTGGGCCCAATACCGCTCGATCGCGAATGGGGAATCGACAA
ACCTTACATCGGTGCGGGTTTTGGACTTGAGCGTTTACTGAAAGTCAAACATGACTTTAAGAATAT
AAAACGTGCTGCACGGTCCGAGTCGTATTACAATGGGATTTCAACTAATCTATAA

Mm-Npa-PylRS[44] nucleotide sequence codon optimized for *E. coli* expression using IDT codon optimization tool (https://www.idtdna.com/pages/tools/codon-optimization-tool), Mm-NpaRS-Opt2:

ATGGATAAAAAACCACTAAACACTCTGATCTCTGCTACTGGTCTGTGGATGAGTCGTACCGGAACC
ATTCATAAAATCAAACACCACGAGGTTAGCCGTTCGAAAATCTATATTGAGATGGCGTGTGGCGAT
CATCTGGTTGTGAACAATAGCCGCTCTTCTCGTACAGCACGTGCACTGCGTCACCACAAATATCGT
AAAACCTGTAAACGTTGCCGTGTGTCCGATGAGGATCTGAACAAATTCCTGACAAAAGCCAATGA
GGACCAAACAAGCGTGAAAGTGAAAGTCGTTAGCGCTCCTACCCGTACTAAAAAAGCAATGCCGA
AATCCGTTGCTCGTGCCCCTAAACCACTGGAAAACACTGAAGCAGCACAGGCACAGCCGTCTGGA
AGCAAATTCTCTCCGGCCATTCCTGTTTCTACCCAGGAGTCCGTTTCTGTTCCAGCAAGTGTGAGCA
CCAGCATTAGCAGTATTAGCACCGGTGCCACCGCTAGCGCCCTGGTTAAAGGCAATACCAATCCG
ATTACAAGCATGTCTGCCCCGGTTCAAGCATCAGCTCCAGCACTGACAAAATCCCAAACCGATCGT
CTGGAGGTTCTGCTGAATCCGAAAGACGAAATCAGCCTGAATTCCGGCAAACCGTTTCGTGAACTG
GAGAGCGAACTGCTGTCACGTCGTAAAAAAGACCTGCAACAAATCTATGCCGAAGAACGTGAGAA
CTATCTGGGGAAACTGGAACGTGAAATCACCCGCTTTTTCGTGGATCGTGGCTTTCTGGAGATCAA
ATCCCCGATTCTGATTCCTCTGGAGTATATCGAGCGTATGGGCATCGACAATGATACCGAACTGAG
CAAACAAATTTTCCGTGTGGATAAAAACTTCTGTCTGCGCCCTATGCTGACACCAAATCTGTATAA
CTATCTGCGCAAACTGGACCGTGCCCTGCCTGATCCTATCAAAATCTTCGAGATCGGCCCGTGTTA
TCGTAAAGAGTCCGACGGTAAAGAACATCTGGAGGAGTTTACCATGCTGGGCTTTACCCAAATGG
GTTCAGGTTGTACTCGTGAGAACCTGGAAAGCATCATCACCGATTTTCTGAACCACCTGGGCATTG
ACTTCAAAATTGTGGGCGACAGCTGTATGGTGTATGGCGACACCCTGGATGTCATGCACGGCGACC
TGGAACTGTCTAGTGCCATTGTTGGACCAATTCCGCTGGACCGTGAGTGGGGTATCGACAAACCGT
ATATCGGAGCAGGATTCGGTCTGGAACGCCTGCTGAAAGTGAAACACGACTTCAAAAACATCAAA
CGTGCCGCCCGTTCTGAATCGTATTATAACGGGATTTCTACCAACCTGTAATAA

### Supplementary Methods

#### General Experimental

Commercially available starting materials were obtained from SigmaAldrich, Fluorochem, Acros, Apollo Scientific, Alfa Aesar and Strem. Anhydrous acetonitrile was purchased from Acros. All solvents used in purification were of chromatography or analytical grade. Thin layer chromatography (TLC) was performed using aluminium backed silica (Merck silica gel 60 F254) plates obtained from Merck. Ultraviolet lamp ($\lambda_{max}$ = 254 nm) and $KMnO_4$ were used for visualization. Analytical LC-MS was performed using a system comprising a Waters Acquity H-CLASS UPLC with a PDA detector, ELS detector and SQD2 with electrospray ionisation. The system ran with a positive and negative switching mode using a Waters Acquity UPLC BEH C18 (50 mm × 2.1 mm × 1.7 µm) column and gradient elution with a binary solvent system: MeCN plus 0.1% formic acid and $H_2O$ plus 0.1% formic acid.
Mass directed auto-purification (MDAP) of the hits of the high-throughput array was performed using a Waters Autopurifcation system comprising a PDA detector, ELS detector and SQD2 with electrospray ionisation. The system ran in positive mode using a Waters XBridge Prep C18 (100 mm × 19 mm × 5 µm) or Waters XBridge Prep C18 (50 mm × 19 mm × 5 µm) column and gradient elution with a binary solvent system: MeCN plus 0.1% formic acid and $H_2O$ plus 0.1% formic acid.
Mass directed auto-purification (MDAP) of the scale-ups of the [4+3] cycloaddition-oxidation reaction was performed using an Agilent 1290 Infinity II Preparative HPLC system with mass spectrometer (LC/MSD XT) and fraction collector. The system ran in positive mode with an Agilent Technologies PLRP-S, 300 Å, 8 µm particle size, 150 × 25 mm column at ambient temperature with a binary solvent system: MeCN and $H_2O$ with 0.1% formic acid.
Flash column chromatography was performed using silica gel 60 (35-70 µm particles) supplied by Merck. A Bruker Daltonics micrOTOF spectrometer with electrospray (ES) ionisation source was used for high-resolution mass spectrometry (HRMS). Proton ($^1H$) and carbon ($^{13}C$) NMR data was collected on a Bruker 300, 400 or 500 MHz spectrometer using $CDCl_3$, $CD_3CN$, or DMSO-d6 as solvent. Data was collected at 298 K unless otherwise stated. Chemical shifts (δ) are given in parts per million (ppm) and they are referenced to the residual solvent peak. Coupling constants (J) are reported in Hertz (Hz) and splitting patterns are reported in an abbreviated manner: app. (apparent), s (singlet), d (doublet), t (triplet), q (quartet), pent (pentet), m (multiplet), br. (broad). Assignments and identifications were made using COSY, DEPT, HSQC, HMBC and NOESY experiments.

#### Chemical Methods

##### High-throughput synthesis arrays to discover the cycloaddition reaction

Stock solutions in MeCN were prepared for each substrate (**S1-S28**) (0.67 M), Ir[dF($CF_3$)ppy]$_2$(dtbpy)$PF_6$ (0.02 M), TBPA (1 M), TRIP-thiol (0.25 M) and TFA (2 M). Initially, 96 reactions were identified for execution through random selection of a limiting reactant (from 28), an excess reactant (from 28) and reaction condition (from six). The reactions were performed in micro-scale vials in 96-well plates on 100 µL scale. Hence, to each reaction vial in a 96-well plate was added a limiting reagent (15 µL, 0.01 mmol), an excess reagent (45 µL) and Ir[dF($CF_3$)ppy]$_2$(dtbpy)$PF_6$ (10 µL), followed by the randomly selected combinations of TBPA (20 µL), TRIP-thiol (20 µL) and TFA (10 µL). Finally, where necessary, MeCN was added to the reaction vials to bring the final volume of each reaction to 100 µL. Each reaction vial was capped, and the plate was, subsequently, irradiated with 2x 40 W Kessil A160WE Tuna Blue lamps from a distance of 17 cm for 16 h. Herein, a fan was used to avoid overheating. At the end of the run each corner of the plate was determined to be at a uniform 25 °C.

The reaction mix was then concentrated in the vials and prepared for analysis. The success of each reaction was evaluated by analytical UPLC/MS, with an additional evaporative light-scattering detector (ELSD) allowing for estimating the approximate product yield (see below). No assumption was made on the molecular weight of a product. All reactions, which had potentially led to at least 0.75 µmol of a mass signal >30 higher than any of the reaction substrates (as such corresponding to >8% yield). Each products of these hit reactions were purified by mass-directed HPLC followed by elucidating the structures *via* 500 MHz $^1$H NMR.

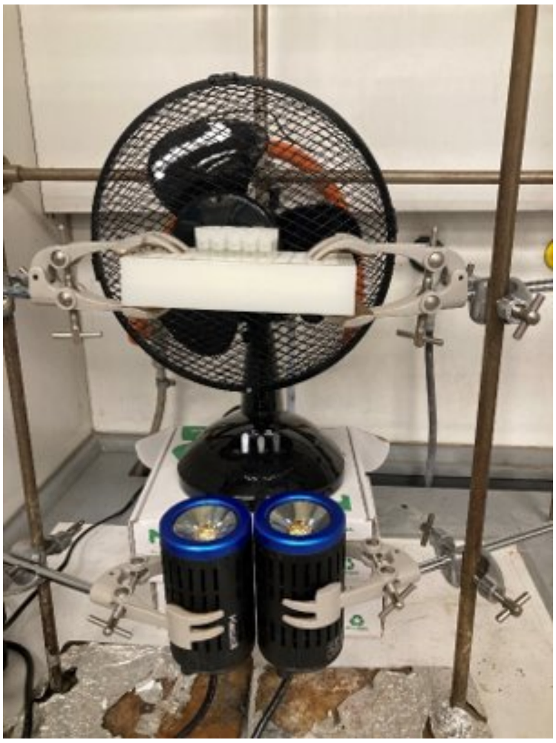

**Figure SC1.** Example set up for the high-throughput array

**ELSD calibration to estimate reaction productivity from analytical liquid chromatography**

The UPLC-ELSD calibration analytes were purchased from SigmaAldrich, Fluorochem and Fisher Scientific. The calibration mixes comprised of equal mass concentrations of 7-hydroxyethyl theophylline, hydrocortisone, dibenzyl 2,3-dihydroxysuccinate, dibenzyl succinate and dibenzyl phthalate. The calibration mixes were prepared at 7 concentration levels: 12.5, 10, 7.5, 5, 2.5, 1.0 and 0.5 mg/mL. The calibration mixes were all prepared by weighing on a calibrated 1 decimal place balance and diluted to volume in 10 ml volumetric flasks with dimethylsulfoxide:water (80:20). Where necessary, flasks were immersed in an ultrasonic bath to aid solubility.
1 µL of each of the seven calibration concentration mixes were injected in triplicate onto the UPLC-MS system and the ELSD peak areas and retention times were manually extracted from the generated results files. 3-dimensional calibration equations were produced in origin with retention time ($t_R$, in minutes) as the X-axis, log ELSD peak area (AUC) as the Y-axis and log mass concentration as the Z-axis. The 3D calibration curve, produced, was described by the equation[6]:

$$\log(c) = A + B \times t_R + C \times \log_{10}(AUC) + D \times t_R^2 + E \times t_R \times \log_{10}(AUC) + F \times (\log_{10}(AUC))^2$$

$R^2$ was calculated as a goodness-of-fit measure ($R^2 = 0.998$)

A series of test compounds (*M* range: 250-400 g/mol) was prepared at concentrations (0.5-10 mg/mL). 1 µL of each test compound was injected onto the analytical system and the ELSD AUC and $t_R$ extracted as before. Values were substituted into the described equation to give calculated concentrations. All test compounds returned calculated concentrations within 10% of the actual concentration.

**General procedures for the [4+3] cycloaddition-oxidation with small molecules**

***General procedure A for scaled-up [4+3] cycloaddition-oxidation reaction:*** To a solution of $Ir(ppy)_3$ (1.3 mg, 0.002 mmol) in MeCN (0.8 mL) and $H_2O$ (0.2 mL) in a 7 mL scintillation vial was added substituted-imidazo[1,2-*a*]pyrimidine (0.1 mmol), component to be dearomatised (0.3 mmol) and tert-butyl peracetate (50 wt% in mineral spirits, 64 µL, 0.2 mmol). The mixture was stirred for 16 h with a blue LED lamp ~10 cm away, then concentrated in vacuo to give the crude product.

***General procedure B for catalyst- and oxidant-free [4+3] cycloaddition-oxidation reaction:*** To a solutions of imidazo[1,2-*a*]pyrimidine analog (0.03 mmol, 100 mM, 1 eq) in 0.3 mL of DMSO in 1.2 mL reaction glass vials was added naphthalene (0.09 mmol, 300 mM, 3 eq). The vials were placed in a 96-position DeSyre reaction block and irradiated at 365 nm (115 mW/well) for 16 h with agitating (400 rpm) and cooling the reaction block holder (chiller temperature at 4 °C). Subsequently, the reactions were cooled to ambient temperature, diluted five times (to 20 mM of maximum anticipated product concentration) in DMSO, and analysed by analytical liquid chromatography (UPLC). Reaction products were isolated and purified by means of preparative mass directed HPLC and compared to existing standards of the expected products utilizing NMR spectroscopy. Those experiments were also performed with identical amounts of reagents in an oxygen-free environment within the Belle epoque glove box, thereby not yielding any product.

#### Synthesized cycloaddition-oxidation products

*18-Methoxy-7-(trifluoromethyl)-4,6,15,19-tetraazapentacyclo[8.6.2.12,5.011,16.09,19]nonadeca-2,4,6,8,11,13,15,17-octaene* ***(1a)*** *and 17-Methoxy-7-(trifluoromethyl)-4,6,12,19-tetraazapentacyclo[8.6.2.12,5.011,16.09,19]nonadeca- 2,4,6,8,11,13,15,17-octaene* ***(1b)***

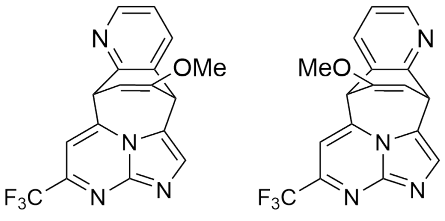

Following the general procedure **A** with 7-(trifluoromethyl)imidazo[1,2-*a*]pyrimidine (18.7 mg, 0.1 mmol) and quinazoline (47.7 mg, 0.3 mmol). The reaction gave a 45:55 ratio of regioisomers. The crude product was purified by column chromatography (80:20 EtOAc/hexane, then EtOAc) to give cycloadducts **1a** and **1b**.

**1a** (7.8 mg, 21% yield) as a yellow oil; $\delta_H$ (500 MHz, $CDCl_3$) 8.42 (1H, dd, J = 5.0, 1.6 Hz, H14), 7.90 (1H, s, H3), 7.70 (1H, dd, J = 7.6, 1.6 Hz, H12), 7.22 (1H, dd, J = 7.6, 5.0 Hz, H13), 7.16 (1H, s, H8), 5.46 (1H, dd, J = 7.3, 2.2 Hz, H17), 5.04 (1H, d, J = 7.3 Hz, H1), 4.66 (1H, d, J = 2.2 Hz, H10), 3.58 (3H, s, OMe) ppm; $\delta_C$ (126 MHz, $CDCl_3$) 158.7 (C16), 155.4 (C18), 148.9 (C14), 147.2 (C5), 147.1 (q, J = 36.9 Hz, C7), 142.5 (C9), 133.7 (C12), 133.6 (C3), 128.6 (C11), 122.8 (C13), 120.8 (q, J = 275.4 Hz, $CF_3$), 119.5 (C2), 101.3 (q, J = 2.1 Hz, C8), 96.2 (C17), 56.3 (OMe), 49.4 (C10), 40.2 (C1) ppm; $\delta_F$ (376 MHz, $CDCl_3$) −68.0 ppm; HRMS Found M+Na 367.0775 *m/z*; $C_{17}H_{11}F_3N_4NaO$ requires M+Na 367.0777 *m/z*.

**1b** (3.7 mg, 11% yield) as a yellow oil; $\delta_H$ (500 MHz, $CD_3CN$) 8.35 (1H, dd, J = 5.0, 1.6 Hz, H13), 7.85 (1H, s, H3), 7.78 (1H, dd, J = 7.6, 1.6 Hz, H15), 7.41 (1H, s, H8), 7.25 (1H, dd, J = 7.6, 5.0 Hz, H14), 5.30 (1H, dd, J = 7.4, 2.4 Hz, H18), 5.00 (1H, d, J = 7.4 Hz, H10), 4.99 (1H, d, J = 2.4 Hz, H1), 3.56 (3H, s, OMe) ppm; $\delta_C$ (126 MHz, $CD_3CN$) 160.8 (C17), 156.8 (C11), 148.8 (C13), 148.2 (C5), 147.4 (q, J = 36.0 Hz, C7), 146.8 (C9), 134.1 (C16), 133.9 (C3), 133.3 (C15), 124.2 (C14), 122.1 (q, J = 274.2 Hz, $CF_3$), 118.6 (C2), 101.8 (q, J = 2.2 Hz, C8), 91.6 (C18), 56.9 (OMe), 47.3 (C10), 43.1 (C1) ppm; $\delta_F$ (376 MHz, $CDCl_3$) −68.2 ppm; HRMS found M+H, 345.0957 *m/z*. $C_{17}H_{12}F_3N_4O$ requires 345.0958 *m/z*.

The configuration of the 18-methoxy-regioisomer **1a** was confirmed using $^1$H-NOESY, and through the collection of X-ray crystallographic data. CCDC deposition number 2157884.

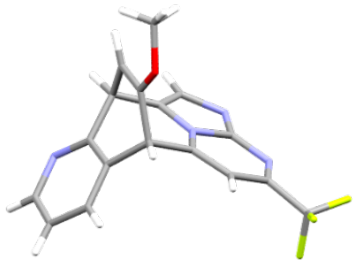

**Figure SC2:** X-ray crystallographic data of 1a to confirm the configuration of the 18-methoxy-regioisomer

*12-Bromo-7-(trifluoromethyl)-4,6,14,19-tetraazapentacyclo[8.6.2.12,5.011,16.09,19]nonadeca-2,4,6,8,11,13,15,17-octaene (**2**)*

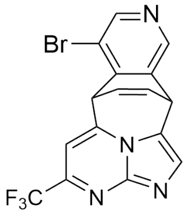

**2**

Following the general procedure **A** with 7-(trifluoromethyl)imidazo[1,2-*a*]pyrimidine (18.7 mg, 0.1 mmol) and 3-bromoisoquinoline (61.8 mg, 0.3 mmol). The reaction gave a 97:3 ratio of regioisomers. The crude product was purified by column chromatography (80:20 EtOAc/hexane to give cycloadduct **2** (7.8 mg, 20% yield) as a yellow oil. Only data for the major regioisomer is presented. $\delta_H$ (500 MHz, $CDCl_3$) 8.67 (1H, s, H13), 8.53 (1H, s, H15), 7.89 (1H, s, H3), 7.24 (1H, s, H8), 6.71 (1H, ddd, J = 7.9, 6.6, 1.0 Hz, H17), 6.37 (1H, ddd, J = 7.9, 6.6, 1.1 Hz, H18), 5.28 (1H, br. d, J = 6.6 Hz, H10), 5.03 (1H, dd, J = 6.6, 1.1 Hz, H1) ppm; $\delta_C$ (126 MHz, $CDCl_3$) 151.2 (C13), 147.9 (q, J = 37.1 Hz, C7), 147.3 (C5), 143.9 (C15), 141.8 (C9), 140.7 (C11), 135.4 (C16), 133.8 (C3), 132.0 (C17), 124.2 (C18), 120.7 (q, J = 275.3 Hz, $CF_3$), 119.7 (C12), 116.8 (C2), 102.1 (q, J = 2.0 Hz, C8), 43.8 (C10), 36.5 (C1) ppm; $\delta_F$ (376 MHz, $CDCl_3$) -68.1 ppm; HRMS Found M+H 392.9968 and 394.9948 *m/z*. $C_{16}H_9BrF_3N_4$ requires M+H 392.9957 and 394.9937 *m/z*.

The configuration of the major regioisomer was confirmed using $^1$H-NOESY. A through-space interaction was observed between H1 and H3, and H1 and H15.

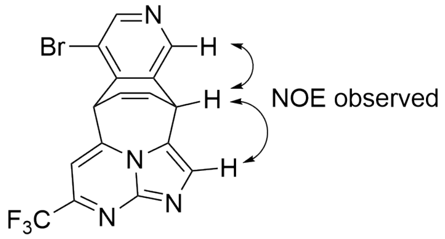

**Figure SC3:** Structure of **2** highlighting the through-space interactions between H1 and H3, as well as H3 and H15 observed to confirm the conformation.

*4,6,19-Triazapentacyclo[8.6.2.12,5.011,16.09,19]nonadeca-2,4,6,8,11,13,15,17-octaene (**3**)*

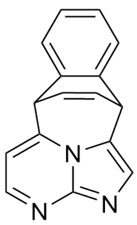

**3**

Following the general procedure **A** with imidazo[1,2-*a*]pyrimidine (IP, 11.9 mg, 0.1 mmol) and naphthalene (Nap, 38.5 mg, 0.3 mmol). The crude product was purified by column chromatography (95:5 DCM/MeOH) to give cycloadduct **3** (4.0 mg, 16% yield) as a yellow oil.

A second reaction following the general procedure **B** culminated in **3**, exhibiting similar spectroscopic results (IP: 3.6 mg, 0.03 mmol; Nap:11.54 mg, 0.09 mmol; **3**:1.0 mg 14% yield).

$\delta_H$ (500 MHz, $CD_3CN$) 8.42 (1H, d, J = 4.2 Hz, H7), 7.58 (1H, s, H3), 7.42 (1H, dd, J = 7.0, 1.5 Hz, H12), 7.37 (1H, dd, J = 7.0, 1.5 Hz, H15), 7.28 (1H, ddd, J = 7.5, 7.0, 1.5 Hz, H14), 7.24 (1H, ddd, J = 7.5, 7.0, 1.5 Hz, H13), 6.94 (1H, d, J = 4.2 Hz, H8), 6.63 (1H, ddd, J = 8.2, 6.6, 1.0 Hz, H17), 6.41 (1H, ddd, J = 8.2, 6.6, 0.8 Hz, H18), 4.99 (1H, dd, J = 6.6, 0.8 Hz, H1), 4.71 (1H, br. d, J = 6.6 Hz, H10); $\delta_C$ (126 MHz, $CD_3CN$) 151.5 (C7), 140.5 (C11), 136.8 (C16), 132.5 (C17), 128.9 (C14), 128.1 (C13), 126.8 (C12 and C18), 125.6 (C15), 105.7 (C8), 45.7 (C10), 39.9 (C1) ppm. 12 of 16 expected signals observed, C2 underneath solvent peak; HRMS found M+H 246.1025 *m/z*. $C_{16}H_{12}N_3$ requires 246.1026 *m/z*.

*8-Bromo-4,6,19-triazapentacyclo[8.6.2.12,5.011,16.09,19]nonadeca-2,4,6,8,11,13,15,17-octaene (**4**)*

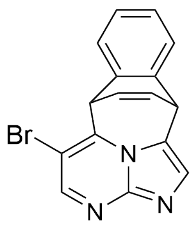

**4**

Following the general procedure **A** with 6-bromoimidazo[1,2-*a*]pyrimidine (IPBr, 19.8 mg, 0.1 mmol) and naphthalene (Nap, 38.5 mg, 0.3 mmol). The crude product was purified by column chromatography (100% EtOAc) to give cycloadduct **4** (5.9 mg, 18% yield) as a yellow oil.

A second reaction following the general procedure **B** culminated in **4**, exhibiting similar spectroscopic results (IPBr: 5.9 mg, 0.03 mmol; Nap:11.54 mg, 0.09 mmol; **4**:1.6 mg 16% yield).

$\delta_H$ (500 MHz, $CDCl_3$) 8.62 (1H, s, H7), 7.74 (1H, s, H3), 7.41 (1H, dd, J = 7.3, 1.5 Hz, H12), 7.36 (1H, dd, J = 7.3, 1.6 Hz, H15), 7.35-7.26 (2H, m, H13 and H14), 6.72 (1H, ddd, J = 8.0, 6.6, 1.0 Hz, H17), 6.40 (1H, ddd, J = 8.0, 6.6, 1.0 Hz, H18), 5.41 (1H, d, J = 6.6 Hz, H10), 4.92 (1H, d, J = 6.6 Hz, H1) ppm; $\delta_C$ (126 MHz, $CDCl_3$) 154.7 (C7), 145.3 (C9), 141.7 (C5), 138.2 (C16), 134.1 (C11), 133.1 (C17), 129.3 (C14), 128.3 (C13), 126.5 (C12), 125.7, 125.5 (C3, C15 and C18), 118.9 (C2), 104.9 (C8), 44.1 (C10), 39.1 (C1) ppm; HRMS found M+H 324.0126 and 326.0108 *m/z*. $C_{16}H_{11}N_3Br$ requires 324.0131 and 326.0110 *m/z*.

*7-(Trifluoromethyl)-4,6,19-triazapentacyclo[8.6.2.12,5.011,16.09,19]nonadeca-2,4,6,8,11,13, 15,17-octaene (**5**)*

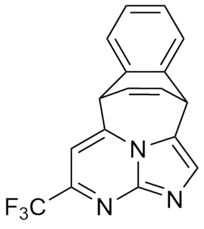

**5**

Following the general procedure **A** with 7-(trifluoromethyl)imidazo[1,2-*a*]pyrimidine (IPF, 18.7 mg, 0.1 mmol) and naphthalene (Nap, 38.5 mg, 0.3 mmol). The crude product was purified by column chromatography (100% EtOAc) to give cycloadduct **5** (8.2 mg, 26% yield) as a yellow oil.

A second reaction following the general procedure **B** culminated in **5**, exhibiting similar spectroscopic results (IPF: 5.6 mg, 0.03 mmol; Nap:11.54 mg, 0.09 mmol; **5**:2.1 mg, 22% yield).

$\delta_H$ (500 MHz, $CDCl_3$) 8.01 (1H, s, H3), 7.40 (1H, dd, J = 7.4, 1.5 Hz, H12), 7.39 (1H, dd, J = 7.4, 1.5 Hz, H15), 7.34 (1H, s, H8), 7.33 (1H, app. td, J = 7.4, 1.5 Hz, H14), 7.29 (1H, app. td, J = 7.4, 1.5 Hz, H13), 6.75-6.70 (1H, m, H17), 6.48 (1H, ddd, J = 8.0, 6.5, 1.0 Hz, H18), 5.05 (1H, br. d, J = 6.5 Hz, H1), 4.88 (1H, br. d, J = 6.5 Hz, H10) ppm; $\delta_C$ (126 MHz, $CDCl_3$) 150.0 (q, J = 37.8 Hz, C7), 146.0 (C9), 145.2 (C5), 138.1 (C16), 134.4 (C11), 132.4 (C17), 129.2 (C3), 128.4, 128.2 (C13 and C14), 126.3, 126.2, 125.5 (C12, C15 and C18), 120.3 (q, J = 275.9 Hz, $CF_3$), 119.9 (C2), 102.7 (q, J = 1.0 Hz, C8), 45.9 (C10), 39.1 (C1) ppm; $\delta_F$ (376 MHz, $CDCl_3$) −68.3 ppm; HRMS found M+H 314.0896 *m/z*. $C_{17}H_{11}N_3F_3$ requires 314.0900 *m/z.*

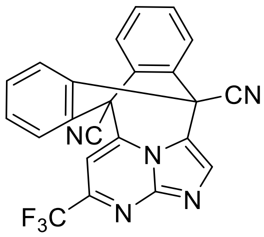

7-(Trifluoromethyl)-4,6,23-triazahexacyclo[8.6.6.$1^{2,5}$.$0^{11,16}$.$0^{17,22}$.$0^{9,23}$]tricosa-2,4,6,8, 11(16),12,14,17(22),18,20-decaene-1,10-dicarbonitrile **11**

Following the general procedure with 7-(trifluoromethyl)imidazo[1,2-a]pyrimidine (18.7 mg, 0.1 mmol) and 9,10-dicyanoanthracene (68.4 mg, 0.3 mmol). The crude product was purified by column chromatography (10:90 EtOAc/hexane) to give *cycloadduct* 11 (14.0 mg, 34% yield) as a colourless solid. $\delta_H$ (500 MHz, $CDCl_3$) 8.21 (1H, s, 3-H), 8.02 (2H, dd, *J* = 7.5, 1.5 Hz, 12-H and 21-H), 7.98 (2H, dd, *J* = 7.5, 1.5 Hz, 15-H and 18-H), 7.72 (1H, s, 8-H), 7.64 (2H, app. td, *J* = 7.5, 1.5 Hz, 14-H and 19-H), 7.60 (2H, app. td, *J* = 7.5, 1.5 Hz, 13-H and 20-H) ppm; $\delta_C$ (126 MHz, $CDCl_3$) 148.6 (q, *J* = 37.7 Hz, C-7), 148.1 (C-5), 140.9 (C-9), 133.2 (C-3), 133.0 (C-11 and C-22), 131.8 (C-14 and C-19), 130.8 (C-13 and C-20), 128.5 (C-16 and C-17), 126.0 (C-12 and C-21), 124.5 (C-15 and C-18), 120.3 (q, *J* = 275.9 Hz, $CF_3$), 117.5 (C-2), 114.7 (CN), 114.6 (CN), 99.8 (q, *J* = 2.1 Hz, C-8), 52.2 (C-10), 47.3 (C-1) ppm; $\delta_F$ (376 MHz, $CDCl_3$) −68.2 ppm; HRMS found M+H 414.0971. $C_{23}H_{11}N_3F_3$ requires 414.0961.

The structure was confirmed by the collection of X-ray crystallographic data. CCDC deposition number 2157886.

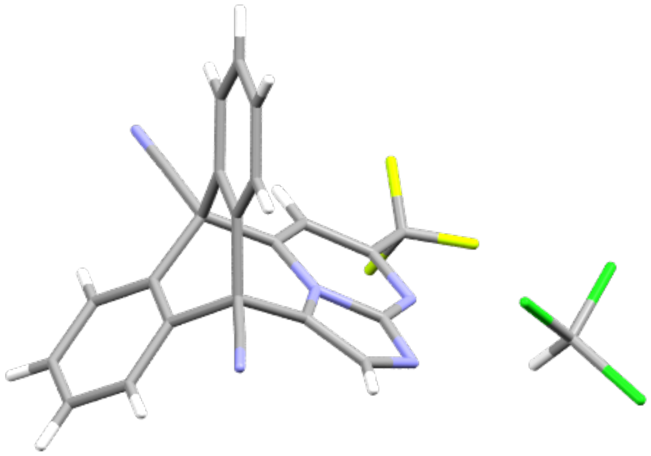

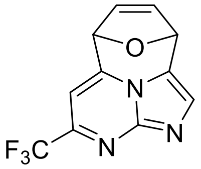

6-(Trifluoromethyl)-14-oxa-3,5,13-triazatetracyclo[6.4.1.$1^{9,12}$.$0^{4,13}$]tetradeca-1,3,5,7,10-pentaene **12**

Following the general procedure with 7-(trifluoromethyl)imidazo[1,2-a]pyrimidine (18.7 mg, 0.1 mmol) and furan (20.4 mg, 0.3 mmol). The crude product was purified by column chromatography (100% EtOAc) to give *cycloadduct* 12 (4.0, 16% yield) as a yellow oil. $\delta_H$ (500 MHz, $CDCl_3$) 7.98 (1H, s, 2-H), 7.28 (1H, s, 7-H), 6.30 (1H, br. d, *J* = 5.5 Hz, 10-H), 6.13 (1H, br. d, *J* = 1.7 Hz, 9-H), 6.06 (1H, dd, *J* = 5.5, 1.7 Hz, 11-H), 5.82 (1H, br. d, *J* = 1.3 Hz, 12-H) ppm; $\delta_C$ (126 MHz, $CDCl_3$) 148.7 (q, *J* = 36.5 Hz, C-6), 145.0 (C-4), 141.8 (C-8), 131.4 (C-2), 129.7 (C-10), 122.4 (C-11), 120.7 (q, *J* = 275.6 Hz, $CF_3$), 114.7 (C-1), 99.7 (q, *J* = 1.6 Hz, C-7), 77.1, 77.0 (C-9 and C-12) ppm; $\delta_F$ (376 MHz, $CDCl_3$) −67.8 ppm; HRMS found M+H 254.0530. $C_{11}H_7N_3F_3O$ requires 254.0536.

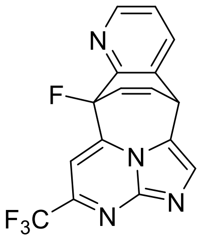

10-Fluoro-7-(trifluoromethyl)-4,6,15,19-tetraazapentacyclo[8.6.2.$1^{2,5}$.$0^{11,16}$.$0^{9,19}$]nonadeca-2,4,6,8,11,13,15,17-octaene **13**

Following the general procedure with 7-(trifluoromethyl)imidazo[1,2-a]pyrimidine (18.7 mg, 0.1 mmol) and 8-fluoroquinoline (44.1 mg, 0.3 mmol). The reaction gave a 90:10 ratio of regioisomers. The crude product was purified by column chromatography (75:25 up to 80:20 EtOAc/hexane) to give *cycloadduct* 13 (8.6 mg, 26% yield) as a yellow oil. Only data for the major regioisomer is presented. $\delta_H$ (500 MHz, $CD_3CN$) 8.51 (1H, dd, *J* = 4.8, 1.6 Hz, 13-H), 7.90 (1H, s, 3-H), 7.85 (1H, app. dt, *J* = 7.7, 1.6 Hz, 15-H), 7.53 (1H, d, *J* = 1.4 Hz, 8-H), 7.37 (1H, dd, *J* = 7.7, 4.8 Hz, 14-H), 6.67 (1H, ddd, *J* = 8.9, 6.7, 3.5 Hz, 17-H), 6.45 (1H, ddd, *J* = 13.6, 8.9, 1.3 Hz, 18-H), 5.26 (1H, dd, *J* = 6.7, 1.3 Hz, 1-H) ppm; $\delta_C$ (126 MHz, $CD_3CN$) 152.8 (d, *J* = 16.3 Hz, C-11), 149.3 (C-13), 147.8 (C-5), 147.4 (qd, *J* = 36.4, 1.3 Hz, C-7), 143.6 (d, *J* = 35.6 Hz, C-9), 134.9 (C-3), 133.6 (d, *J* = 3.0 Hz, C-15), 131.3 (d, *J* = 4.0 Hz, C-16), 129.9 (d, *J* = 11.6 Hz, C-17), 128.8 (d, *J* = 28.2 Hz, C-18), 125.6 (C-14), 122.0 (q, *J* = 274.2 Hz, $CF_3$), 98.4 (dq, *J* = 7.5, 2.2 Hz, C-8), 90.9 (d, *J* = 198.1 Hz, C-10), 38.0 (d, *J* = 0.9 Hz, C-1) ppm. 15 of 16 expected signals observed, C-2 found underneath the solvent peak; $\delta_F$ (376 MHz, $CDCl_3$) −182.4, −68.2 ppm; HRMS Found M+Na 355.0578. $C_{16}H_8F_4N_4Na$ requires M+Na 355.0577.

The configuration of the major regioisomer was confirmed using $^1$H-NOESY, and through the collection of X-ray crystallographic data. CCDC deposition number 2157887.

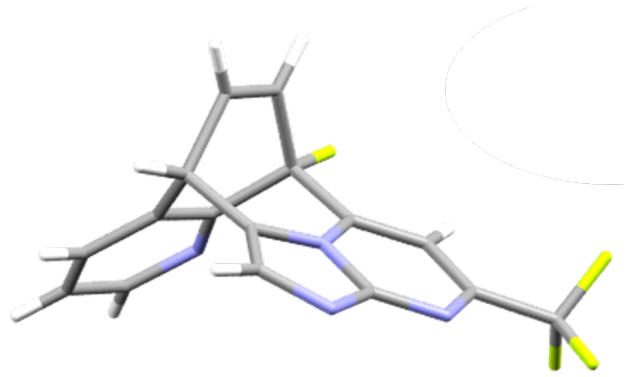

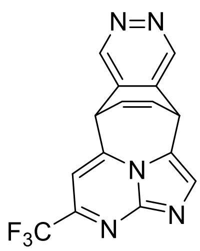

7-(Trifluoromethyl)-4,6,13,14,19-pentaazapentacyclo[8.6.2.1$^{2,5}$.0$^{11,16}$.0$^{9,19}$]nonadeca-2,4,6,8,11,13,15,17-octaene **14**

Following the general procedure with 7-(trifluoromethyl)imidazo[1,2-a]pyrimidine (18.7 mg, 0.1 mmol) and quinazoline (39.0 mg, 0.3 mmol). The crude product was purified by MDAP, with a gradient of 85:15→65:35 $H_2O$ (0.1% formic acid)/MeCN over 12 mins to give *cycloadduct* 14 (3.9 mg, 12% yield) as a yellow oil. $\delta_H$ (500 MHz, $CD_3CN$) 9.27-9.26 (2H, m, 12-H and 15-H), 7.88 (1H, s, 3-H), 7.42 (1H, s, 8-H), 6.65 (1H, ddd, $J$ = 8.2, 6.5, 1.0 Hz, 18-H), 6.45 (1H, ddd, $J$ = 8.2, 6.4, 1.0 Hz, 17-H), 5.19 (1H, dd, $J$ = 6.5, 1.0 Hz, 10-H), 5.09 (1H, dd, $J$ = 6.4, 1.0 Hz, 1-H) ppm; $\delta_C$ (126 MHz, $CD_3CN$) 149.4, 148.5 (C-12 and C-15), 143.1 (C-9), 138.5 (C-16), 134.93, 134.91 (C-3 and C-11), 131.6 (C-18), 125.8 (C-17), 116.7 (C-2), 103.0 (q, $J$ = 2.0 Hz, C-8), 42.0 (C-10), 36.2 (C-1) ppm. 12 of 15 expected signals observed; $\delta_F$ (376 MHz, $CDCl_3$) −68.2 ppm; HRMS found M+H, 316.0804. $C_{15}H_9F_3N_5$ requires 316.0805.

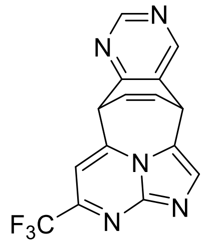

7-(Trifluoromethyl)-4,6,12,14,19-pentaazapentacyclo[8.6.2.$1^{2,5}$.$0^{11,16}$.$0^{9,19}$]nonadeca-2,4,6,8,11,13,15,17-octaene **15**

Following the general procedure with 7-(trifluoromethyl)imidazo[1,2-a]pyrimidine (18.7 mg, 0.1 mmol) and quinazoline (39.0 mg, 0.3 mmol). The reaction gave a 95:5 ratio of regioisomers. The crude product was purified by column chromatography (80:20 EtOAc/hexane, then 95:5 $CH_2Cl_2$/MeOH) to give *cycloadduct* 15 (10.1 mg, 32% yield) as a yellow oil. Only data for the major regioisomer is presented. $\delta_H$ (500 MHz, $CDCl_3$) 9.10 (1H, s, 13-H), 8.74 (1H, s, 15-H), 7.93 (1H, s, 3-H), 7.27 (1H, s, 8-H), 6.67 (1H, ddd, $J$ = 8.2, 6.5, 1.0 Hz, 17-H), 6.41 (1H, ddd, $J$ = 8.2, 6.5, 1.0 Hz, 18-H), 5.02 (1H, dd, $J$ = 6.5, 1.0 Hz, 1-H), 4.97 (1H, dd, $J$ = 6.5, 1.0 Hz, 10-H) ppm; $\delta_C$ (126 MHz, $CDCl_3$) 162.3 (C-11), 158.1 (C-13), 151.8 (C-15), 148.1 (q, $J$ = 37.1 Hz, C-7), 147.4 (C-5), 140.9 (C-9), 133.9 (C-3), 131.2 (C-16), 131.1 (C-17), 124.2 (C-18), 120.7 (q, $J$ = 275.5 Hz, $CF_3$), 116.3 (C-2), 102.5 (q, $J$ = 2.1 Hz, C-8), 48.0 (C-10), 35.3 (C-1) ppm; $\delta_F$ (376 MHz, $CDCl_3$) −68.2 ppm; HRMS Found M+H 316.0811. $C_{16}H_8F_4N_4Na$ requires M+H 316.0805.

The configuration of the major regioisomer was confirmed using $^1$H-NOESY. A through-space interaction was observed between 1-H and 3-H, and 1-H and 15-H.

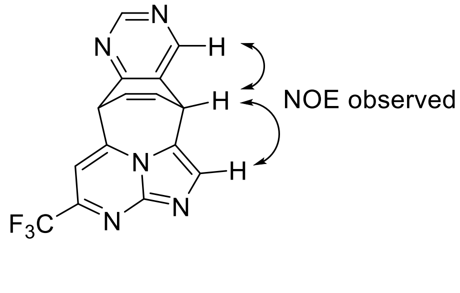

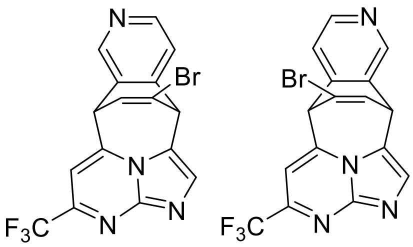

17-Bromo-7-(trifluoromethyl)-4,6,13,19-tetraazapentacyclo[8.6.2.$1^{2,5}$.$0^{11,16}$.$0^{9,19}$]nonadeca-2,4,6,8,11,13,15,17-octaene **18a** and 18-bromo-7-(trifluoromethyl)-4,6,14,19-tetraazapentacyclo[8.6.2.$1^{2,5}$.$0^{11,16}$.$0^{9,19}$]nonadeca-2,4,6,8,11,13,15,17-octaene **18b**

Following the general procedure with 7-(trifluoromethyl)imidazo[1,2-a]pyrimidine (18.7 mg, 0.1 mmol) and 6-bromoisoquinoline (62.4 mg, 0.3 mmol). The reaction gave a 25:75 ratio of regioisomers. The crude product was purified by column chromatography (100% EtOAc) to give *cycloadducts* 18a and 18b. 18a required further purification by MDAP, with a gradient of 76:24→56:44 $H_2O$ (0.1% formic acid)/MeCN over 12 mins. 18a (1.2 mg, 3% yield) as a yellow oil; $\delta_H$ (500 MHz, $CD_3CN$) 8.63 (1H, s, 12-H), 8.60 (1H, d, $J$ = 4.9 Hz, 14-H), 7.95 (1H, s, 3-H), 7.43 (1H, d, $J$ = 4.9 Hz, 15-H), 7.42 (1H, s, 8-H), 6.67 (1H, dd, $J$ = 7.1, 1.9 Hz, 18-H), 5.30 (1H, d, $J$ = 1.9 Hz, 1-H), 5.14 (1H, d, $J$ = 7.1 Hz, 10-H) ppm; $\delta_C$ (126 MHz, $CD_3CN$) 150.9 (C-14), 147.4 (C-12), 147.2 (C-5), 144.4 (C-16), 135.0 (C-3), 130.4 (C-11), 127.8 (C-18), 120.7 (C-17), 120.4 (C-15), 116.3 (C-2), 102.8 (q, $J$ = 2.3 Hz, C-8), 48.6 (C-1), 37.9 (C-10) ppm. 13 of 16 expected signals observed; $\delta_F$ (376 MHz, $CD_3CN$) −69.0 ppm; HRMS found M+H 392.9951 and 394.9931. $C_{16}H_9N_4BrF_3$ requires 392.9957 and 394.9937. **18b** (4.9 mg, 12% yield) as a yellow oil; $\delta_H$ (500 MHz, $CD_3CN$) 8.61 (1H, s, 15-H), 8.51 (1H, d, $J$ = 4.9 Hz, 13-H), 7.88 (1H, s, 3-H), 7.48-7.45 (2H, m, 8-H and 12-H), 6.92 (1H, dd, $J$ = 7.1, 1.8 Hz, 17-H), 5.24 (1H, d, $J$ = 1.8 Hz, 10-H), 5.23 (1H, d, $J$ = 7.1 Hz, 1-H) ppm; $\delta_C$ (126 MHz, $CDCl_3$) 149.9 (C-13), 147.9 (q, $J$ = 37.1 Hz, C-7), 147.4 (C-5), 145.4 (C-15), 142.2 (C-11), 140.2 (C-9), 134.0 (C-3), 132.5 (C-16), 132.0 (C-17), 120.7 (C-12), 120.6 (q, $J$ = 275.6 Hz, $CF_3$), 117.3 (C-2), 114.2 (C-18), 102.2 (q, $J$ = 2.1 Hz, C-8), 54.8 (C-10), 38.0 (C-1) ppm; $\delta_F$ (376 MHz, $CDCl_3$) −68.0 ppm; HRMS found M+H 392.9957 and 394.9939. $C_{16}H_9N_4BrF_3$ requires 392.9957 and 394.9937.

The configuration of the 17-bromo-regioisomer 18a was confirmed using $^1$H-NOESY. A through-space interaction was observed between 8-H and 10-H, and 10-H and 12-H.

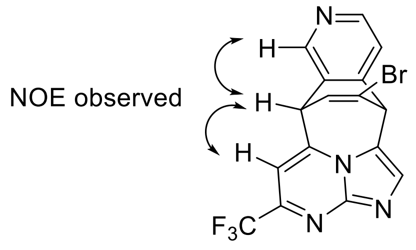

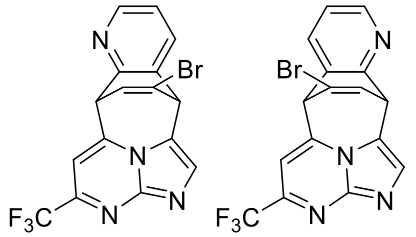

17-Bromo-7-(trifluoromethyl)-4,6,12,19-tetraazapentacyclo[8.6.2.$1^{2,5}$.$0^{11,16}$.$0^{9,19}$]nonadeca-2,4,6,8,11,13,15,17-octaene **19a** and 18-bromo-7-(trifluoromethyl)-4,6,15,19-tetraaza pentacyclo[8.6.2.$1^{2,5}$.$0^{11,16}$.$0^{9,19}$]nonadeca-2,4,6,8,11,13,15,17-octaene **19b**

Following the general procedure with 7-(trifluoromethyl)imidazo[1,2-a]pyrimidine (18.7 mg, 0.1 mmol) and 6-bromoquinoline (62.4 mg, 0.3 mmol). The reaction gave a 77:23 ratio of regioisomers. The crude product was purified by column chromatography (80:20 EtOAc/hexane) to give *cycloadducts* 19a and 19b. 19a (8.0 mg, 20% yield) as a yellow oil; $\delta_H$ (400 MHz, $CDCl_3$) 8.46 (1H, dd, *J* = 5.0, 1.4 Hz, 13-H), 7.98 (1H, s, 3-H), 7.69 (1H, dd, *J* = 7.7, 1.4 Hz, 15-H), 7.29 (1H, dd, *J* = 7.7, 5.0 Hz, 14-H), 7.28 (1H, s, 8-H), 6.59 (1H, dd, *J* = 7.1, 1.9 Hz, 18-H), 5.04-5.02 (1H, m, 1-H), 5.02 (1H, d, *J* = 7.1 Hz, 10-H) ppm; $\delta_C$ (126 MHz, $CDCl_3$) 152.6 (C-11), 149.0 (C-13), 147.7 (q, *J* = 37.9 Hz, C-7), 147.7 (C-5), 142.1 (C-9), 134.4 (C-3), 132.9 (C-16), 132.6 (C-15), 125.5 (C-18), 123.9 (C-14), 121.4 (C-17), 120.7 (q, *J* = 269.8 Hz, $CF_3$), 115.6 (C-2), 102.3 (br. s, C-8), 50.2 (C-10), 48.5 (C-1) ppm; $\delta_F$ (376 MHz, $CDCl_3$) −68.2 ppm; HRMS found M+H 392.9958 and 394.9940. $C_{16}H_9N_4BrF_3$ requires 392.9957 and 394.9937. **19b** (1.9 mg, 5% yield) as a yellow oil; $\delta_H$ (500 MHz, $CDCl_3$) 8.48 (1H, dd, *J* = 5.0, 1.6 Hz, 14-H), 7.97 (1H, s, 3-H), 7.72 (1H, dd, *J* = 7.7, 1.6 Hz, 12-H), 7.27 (1H, dd, *J* = 7.7, 5.0 Hz, 13-H), 7.25 (1H, s, 8-H), 6.84 (1H, dd, *J* = 7.1, 1.8 Hz, 17-H), 5.15 (1H, d, *J* = 7.1 Hz, 1-H), 4.88 (1H, d, *J* = 1.8 Hz, 10-H) ppm; $\delta_C$ (126 MHz, $CDCl_3$) 155.7 (C-16), 149.7 (C-14), 147.5 (C-5), 147.5 (q, *J* = 37.1 Hz, C-7), 140.8 (C-9), 134.8 (C-3), 133.7 (C-12), 132.1 (C-17), 128.7 (C-11), 123.1 (C-13), 120.7 (q, *J* = 275.6 Hz, $CF_3$), 116.3 (C-2), 114.8 (C-18), 101.9 (q, *J* = 2.1 Hz, C-8), 54.6 (C-10), 43.8 (C-1) ppm; $\delta_F$ (376 MHz, $CDCl_3$) −67.9 ppm; HRMS found M+H 392.9955 and 394.9935. $C_{16}H_9N_3BrF_3$ requires 392.9957 and 394.9937.

The configuration of the 17-bromo-regioisomer 19a was confirmed using $^1$H-NOESY. A through-space interaction was observed between 1-H and 3-H, and 1-H and 15-H.

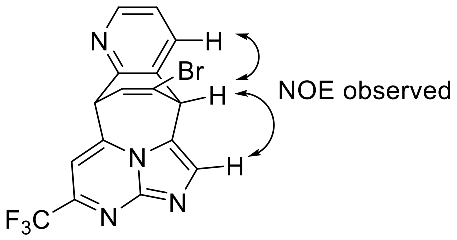

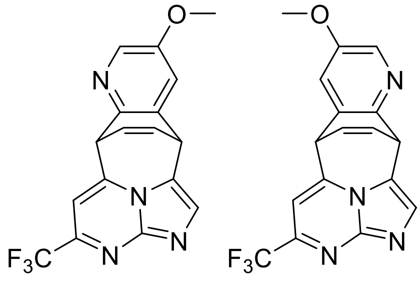

14-Methoxy-7-(trifluoromethyl)-4,6,12,19-tetraazapentacyclo[8.6.2.$1^{2,5}$.$0^{11,16}$.$0^{9,19}$]nonadeca-2,4,6,8,11,13,15,17-octaene **20a** and 13-methoxy-7-(trifluoromethyl)-4,6,15,19-tetraazapentacyclo[8.6.2.$1^{2,5}$.$0^{11,16}$.$0^{9,19}$]nonadeca-2,4,6,8,11,13,15,17-octaene **20b**

Following the general procedure with 7-(trifluoromethyl)imidazo[1,2-a]pyrimidine (18.7 mg, 0.1 mmol) and 3-methoxyquinoline (47.7 mg, 0.3 mmol). The reaction gave a 56:44 ratio of regioisomers. The crude product was purified by column chromatography (80:20 EtOAc/hexane) to give *cycloadducts* 20a and 20b. 20a required further purification by MDAP, with a gradient of 71:29→51:49 $H_2O$ (0.1% formic acid)/MeCN over 12 mins. 20a (2.4 mg, 7% yield) as a yellow oil; $\delta_H$ (501 MHz, $CD_3CN$) 8.08 (1H, d, *J* = 2.8 Hz, 13-H), 7.84 (1H, br. s, 3-H), 7.39 (1H, s, 8-H), 7.35 (1H, d, *J* = 2.8 Hz, 15-H), 6.62 (1H, ddd, *J* = 8.2, 6.5, 1.0 Hz, 17-H), 6.44 (1H, ddd, *J* = 8.2, 6.5, 1.2 Hz, 18-H), 5.05 (1H, dd, *J* = 6.5, 1.2 Hz, 1-H), 5.04 (1H, br. d, *J* = 6.5 Hz, 10-H), 3.81 (3H, s, OMe); $\delta_C$ (126 MHz, $CD_3CN$) 156.9 (C-14), 147.5 (q, *J* = 34.5 Hz, C-7), 147.2 (C-11), 147.1 (C-5), 145.1 (C-9), 136.1 (C-13), 135.4 (C-16), 133.9 (C-3), 132.0 (C-17), 126.8 (C-18), 122.1 (q, *J* = 24.2 Hz, $CF_3$), 102.2 (q, *J* = 1.9 Hz, C-8), 56.5 (OMe), 48.3 (C-10), 38.5 (C-1), 15 of 17 expected signals observed, C-2 and C-15 underneath solvent peak; $\delta_F$ (376 MHz, $CDCl_3$) −68.2 ppm; HRMS found M+H 345.0959. $C_{17}H_{12}N_4F_3O$ requires 345.0958. **20b** (4.5 mg, 13% yield) as a yellow oil; $\delta_H$ (500 MHz, $CDCl_3$) 8.12 (1H, d, *J* = 2.7 Hz, 14-H), 7.91 (1H, s, 3-H), 7.22 (1H, d, *J* = 2.7 Hz, 12-H), 7.16 (1H, s, 8-H), 6.69 (1H, ddd, *J* = 8.2, 6.7, 1.0 Hz, 17-H), 6.34 (1H, ddd, *J* = 8.2, 6.5, 1.0 Hz, 18-H), 5.10 (1H, dd, *J* = 6.7, 1.0 Hz, 1-H), 4.70 (1H, dd, *J* = 6.5, 1.0 Hz, 10-H), 3.84 (3H, s, OMe) ppm; $\delta_C$ (126 MHz, $CDCl_3$) 155.6 (C-13), 149.5 (C-16), 147.1 (q, *J* = 36.9 Hz, C-7), 147.3 (C-5), 1431 (C-9), 135.7 (C-14), 134.0 (C-3), 132.3 (C-17), 129.9 (C-11), 124.3 (C-18), 120.8 (q, *J* = 275.4 Hz, $CF_3$), 119.3 (C-12), 117.3 (C-2), 101.0 (q, *J* = 2.1 Hz, C-8), 56.1 (OMe), 45.0 (C-10), 41.5 (C-1) ppm; $\delta_F$ (376 MHz, $CDCl_3$) −68.1 ppm; HRMS found M+H 345.0961. $C_{17}H_{12}N_4F_3O$ requires 345.0958.

The configuration of the 13-methoxy-regioisomer 20b was confirmed using $^{1}$H-NOESY. A through-space interaction was observed between 8-H and 10-H, and 10-H and 11-H.

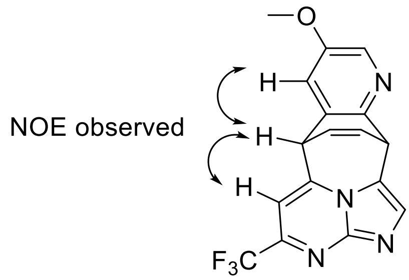

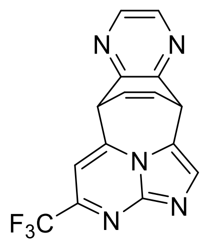

7-(Trifluoromethyl)-4,6,12,15,19-pentaazapentacyclo[8.6.2.1$^{2,5}$.0$^{11,16}$.0$^{9,19}$]nonadeca-2,4,6,8,11,13,15,17-octaene **21**

Following the general procedure with 7-(trifluoromethyl)imidazo[1,2-a]pyrimidine (18.7 mg, 0.1 mmol) and quinoxaline (39.0 mg, 0.3 mmol). The crude product was purified by MDAP with a gradient of 79:21→59:41 $H_2O$ (0.1% formic acid)/MeCN over 12 mins to give *cycloadduct* 21 (7.1 mg, 22% yield) as a yellow oil. $\delta_H$ (500 MHz, $CD_3CN$) 8.41-8.39 (2H, m, 13-H and 14-H), 7.92 (1H, s, 3-H), 7.46 (1H, s, 8-H), 6.68 (1H, ddd, $J$ = 8.2, 6.6, 1.1 Hz, 17-H), 6.44 (1H, ddd, $J$ = 8.2, 6.5, 1.2 Hz, 18-H), 5.19 (1H, dd, $J$ = 6.6, 1.2 Hz, 1-H), 5.13 (1H, dd, $J$ = 6.5, 1.1 Hz, 10-H) ppm; $\delta_C$ (126 MHz, $CD_3CN$) 153.5 (C-16), 150.6 (C-11), 148.2 (C-5), 147.4 (q, $J$ = 36.0 Hz, C-7), 144.4, 144.0 (C-13 and C-14), 143.6 (C-9), 134.8 (C-3), 132.0 (C-17), 126.1 (C-18), 122.1 (q, $J$ = 274.1 Hz, $CF_3$), 117.3 (C-2), 102.9 (q, $J$ = 2.2 Hz, C-8), 47.9 (C-10), 41.5 (C-1) ppm; $\delta_F$ (376 MHz, $CD_3CN$) −68.9 ppm; HRMS found M+H, 316.0804. $C_{15}H_9F_3N_5$ requires 316.0805.

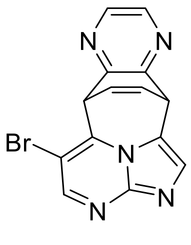

8-Bromo-4,6,12,15,19-pentaazapentacyclo[8.6.2.$1^{2,5}$.$0^{11,16}$.$0^{9,19}$]nonadeca-2,4,6,8,11,13,15,17-octaene **22**

Following the general procedure with 6-bromoimidazo[1,2-a]pyrimidine (19.8 mg, 0.1 mmol) and quinazoline (39.0 mg, 0.3 mmol). The crude product was purified by mass directed auto-purification (100% EtOAc) to give *cycloadduct* 22 (3.3 mg, 10% yield) as a yellow oil. $\delta_H$ (500 MHz, $CD_3CN$) 8.54 (1H, s, 7-H), 8.41-8.40 (2H, m, 13-H and 14-H), 7.70 (1H, s, 3-H), 6.69 (1H, ddd, $J$ = 8.1, 6.6, 1.1 Hz, 17-H), 6.41 (1H, ddd, $J$ = 8.1, 6.6, 1.3 Hz, 18-H), 5.58 (1H, dd, $J$ = 6.6, 1.1 Hz, 10-H), 5.13 (1H, dd, $J$ = 6.6, 1.3 Hz, 1-H) ppm; $\delta_C$ (126 MHz, $CD_3CN$) 153.8 (C-16), 152.2 (C-7), 150.7 (C-11), 148.8 (C-9), 144.5, 144.0 (C-13 and C-14), 139.2 (C-5), 132.8 (C-3), 132.6 (C-17), 125.2 (C-18), 115.1 (C-2), 104.1 (C-8), 46.8 (C-10), 41.5 (C-1) ppm; HRMS found M+H 326.0034 and 328.0015. $C_{15}H_9N_5Br$ requires 326.0036 and 328.0015.

Synthesis and characterisation of array hits 2-5, 7-9

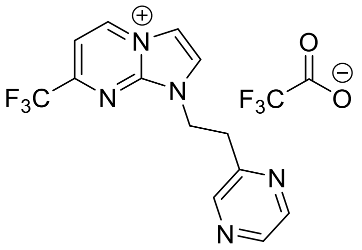

1-[2-(Pyrazin-2-yl)ethyl]-7-(trifluoromethyl)-1H-4$\lambda^5$-imidazo[1,2-a]pyrimidin-4-ylium trifluoroacetate **2**

To a solution of Ir[dF($CF_3$)ppy]$_2$(dtbpy)$PF_6$ (2.2 mg, 0.002 mmol) in MeCN (1 mL) in a 7 mL scintillation vial was added 7-(trifluoromethyl)imidazo[1,2-a]pyrimidine (18.7 mg, 0.1 mmol), 2-vinylpyrazine (31.8 mg, 0.3 mmol) and TFA (15.3 µL, 0.2 mmol). The mixture was stirred for 16 h in darkness, then concentrated *in vacuo*. The residue was vigorously stirred in a biphase of $CH_2Cl_2$ (2 mL) and $H_2O$ (2 mL) and the layers were separated. The aqueous layer was concentrated *in vacuo* to give *trifluoroacetate salt* 2 (40.5 mg, 100% yield) as a brown oil. $\delta_H$ (500 MHz, $D_2O$) 9.44 (1H, dd, $J$ = 7.0, 0.5 Hz, 5-H), 8.53 (1H, d, $J$ = 1.5 Hz, pyrazine 3-H), 8.49 (1H, d, $J$ = 2.8 Hz, pyrazine 6-H), 8.47 (1H, dd, $J$ = 2.8, 1.5 Hz, pyrazine 5-H), 8.34 (2H, s, 2-H and 3-H), 8.01 (1H, d, $J$ = 7.0 Hz, 6-H), 5.06 (2H, t, $J$ = 6.6 Hz, ethyl 1-$H_2$), 3.55 (2H, t, $J$ = 6.6 Hz, ethyl 2-$H_2$) ppm; $\delta_C$ (126 MHz, $D_2O$) 162.9 (q, $J$ = 35.5 Hz, TFA C=O), 153.5 (q, $J$ = 38.6 Hz, C-7), 152.9 (pyrazine C-2), 144.5 (pyrazine C-3), 144.2 (pyrazine C-6), 142.9 (pyrazine C-5), 141.4 (C-5), 127.8 (C-2), 119.3 (q, $J$ = 275.5 Hz, $CF_3$), 116.3 (q, $J$ = 291.8 Hz, TFA $CF_3$), 114.6 (C-3), 110.4 (q, $J$ = 2.2 Hz, C-6), 46.3 (ethyl C-1), 34.2 (ethyl C-2) ppm;

$\delta_F$ (376 MHz, $D_2O$) −73.2, −66.3 ppm; HRMS Found M-$CF_3CO_2^-$ 294.0961. $C_{13}H_{11}F_3N_5$ requires M-$CF_3CO_2^-$ 294.0961.

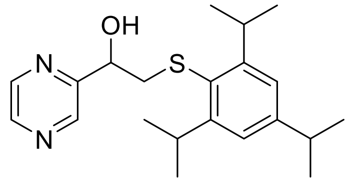

1-(Pyrazin-2-yl)-2-{[2,4,6-tris(propan-2-yl)phenyl]sulfanyl}ethan-1-ol 3

To a solution of Ir[dF($CF_3$)ppy]$_2$(dtbpy)$PF_6$ (2.2 mg, 0.002 mmol) in MeCN (1 mL) in a 7 mL scintillation vial was added 2,4,6-triisopropylbenzenethiol (11.8 mg, 0.05 mmol) and 2-vinylpyrazine (31.8 mg, 0.3 mmol). The mixture was stirred for 16 h with a blue LED lamp ~10 cm away, then concentrated *in vacuo*. The crude residue was purified by column chromatography (30:70 EtOAc/hexane) to give *thioether* 3 (10.4 mg, 58% yield) as a yellow oil. $\delta_H$ (500 MHz, $CDCl_3$) 8.75 (1H, d, $J$ = 1.5 Hz, pyrazine 3-H), 8.51 (1H, d, $J$ = 2.6 Hz, pyrazine 6-H), 8.49 (1H, dd, $J$ = 2.6, 1.5 Hz, pyrazine 5-H), 7.00 (2H, s, phenyl 3-H and 5-H), 4.89 (1H, ddd, $J$ = 7.7, 5.1, 4.7 Hz, ethyl 1-H), 3.82 (2H, hept, $J$ = 7.0 Hz, 2,6-isopropyl 1-$H_2$), 3.53 (1H, d, $J$ = 5.1 Hz, O-H), 3.16 (1H, dd, $J$ = 12.9, 4.7 Hz, ethyl 2-$H_a$), 3.03 (1H, dd, $J$ = 12.9, 7.7 Hz, ethyl 2-$H_b$), 2.87 (1H, hept, $J$ = 7.0 Hz, 4-isopropyl 1-H), 1.24 (6H, d, $J$ = 7.0 Hz, 4-isopropyl $(CH_3)_2$), 1.22 (12H, d, $J$ = 7.0 Hz, 2,6-isopropyl $(CH_3)_4$) ppm; $\delta_C$ (126 MHz, $CDCl_3$) 156.3 (pyrazine C-2), 153.0 (phenyl C-2 and C-6), 150.0 (phenyl C-4), 144.0, 143.5, 143.2 (pyrazine C-3, C-5 and C-6), 127.5 (phenyl C-1), 122.2 (phenyl C-3 and C-5), 71.2 (ethyl C-1), 45.9 (ethyl C-2), 34.4 (4-isopropyl C-1), 31.8 (2,6-isopropyl C-1), 24.7, 24.6, 24.0 (2,4,6-isopropyl $CH_3$) ppm; HRMS Found M+Na 381.1971. $C_{16}H_8F_4N_4Na$ requires M+Na 381.1971.

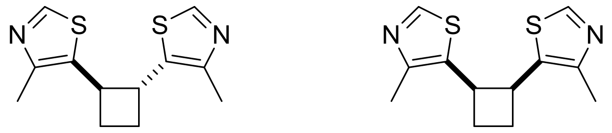

4-Methyl-5-[(1*R**,2*R**)-2-(4-methyl-1,3-thiazol-5-yl)cyclobutyl]-1,3-thiazole *anti*-4 and 4-methyl-5-[(1*R**,2*S**)-2-(4-methyl-1,3-thiazol-5-yl)cyclobutyl]-1,3-thiazole *syn*-4

To a solution of Ir[dF($CF_3$)ppy]$_2$(dtbpy)$PF_6$ (2.2 mg, 0.002 mmol) in MeCN (1 mL) in a 7 mL scintillation vial was added 4-methyl-5-vinylthiazole (34 µL, 0.3 mmol). The mixture was stirred for 16 h with a blue LED lamp ~10 cm away, then concentrated *in vacuo*. The crude residue was purified by column chromatography (98:2 $CH_2Cl_2$/MeOH) to give *cyclobutanes anti*-4 and *syn*-4 (64:36 d.r). *anti*-4 (18.2 mg, 49% yield) as a yellow oil; $\delta_H$ (500 MHz, $CDCl_3$) 8.57 (2H, s, thiazole 2-H and 2'-H), 3.70-3.61 (2H, m, cyclobutane 1-H and 2-H), 2.54-2.40 (2H, m, cyclobutane 3-$H_a$ and 4-$H_a$), 2.24 (6H, s, $(CH_3)_2$) 2.14-2.02 (2H, m, cyclobutane 3-$H_b$ and 4-$H_b$) ppm; $\delta_C$ (126 MHz, $CDCl_3$) 149.4 (thiazole

C-2), 148.3 (thiazole C-4), 135.2 (thiazole C-5), 43.8 (cyclobutane C-1), 29.0 (cyclobutane C-3), 15.2 ($CH_3$) ppm; HRMS Found M+H 251.0674. $C_{12}H_{15}N_2S_2$ requires M+H 251.0671. *syn*-4 (10.6 mg, 28% yield) as a yellow oil; $\delta_H$ (500 MHz, $CDCl_3$) 8.48 (2H, s, thiazole 2-H and 2'-H), 4.25-4.18 (2H, m, cyclobutane 1-H and 2-H), 2.71-2.61 (2H, m, cyclobutane 3-$H_a$ and 4-$H_b$), 2.33-2.25 (2H, m, cyclobutane 3-$H_b$ and 4-$H_b$), 2.18 (6H, s, $(CH_3)_2$) ppm; $\delta_C$ (126 MHz, $CDCl_3$) 149.54 (thiazole C-2), 149.48 (thiazole C-4), 132.2 (thiazole C-5), 37.9 (cyclobutane C-1), 28.9 (cyclobutane C-3), 15.2 ($CH_3$) ppm; HRMS Found M+H 251.0677. $C_{12}H_{15}N_2S_2$ requires M+H 251.0671.

A mixture of diastereomers was also obtained (1.6 mg, 4% yield) as a yellow oil. The configuration of the minor diastereomer, *syn*-4, was determined by the collection of X-ray crystallographic data. CCDC deposition number 2157885.

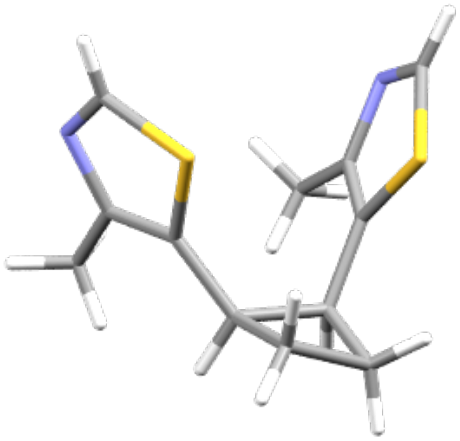

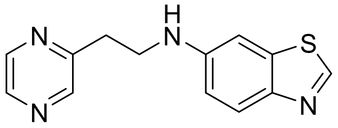

*N*-[2-(Pyrazin-2-yl)ethyl]-1,3-benzothiazol-6-amine **5**

To a solution of Ir[dF($CF_3$)ppy]$_2$(dtbpy)$PF_6$ (2.2 mg, 0.002 mmol) in MeCN (1 mL) in a 7 mL scintillation vial was added 2-vinylpyrazine (10.6 mg, 0.1 mmol), 6-aminobenzothiazole (45 mg, 0.3 mmol) and TFA (15.3 µL, 0.2 mmol). The mixture was stirred for 16 h in darkness, then concentrated *in vacuo*. The crude residue was purified by column chromatography (50:50 up to 70:30 EtOAc/hexane) to give *amine* 5 (10.5 mg, 41% yield) as a yellow amorphous solid. $\delta_H$ (500 MHz, $CDCl_3$) 8.66 (1H, s, benzothiazole 2-H), 8.54 (1H, dd, *J* = 2.6, 1.5 Hz, pyrazine 5-H), 8.50 (1H, d, *J* = 1.5 Hz, pyrazine 3-H), 8.46 (1H, d, *J* = 2.6 Hz, pyrazine 6-H), 7.87 (1H, d, *J* = 8.8 Hz, benzothiazole 4-H), 7.09 (1H, d, *J* = 2.3 Hz, benzothiazole 7-H), 6.81 (1H, dd, *J* = 8.8, 2.3 Hz, benzothiazole 5-H), 4.39 (1H, br. s, N-H), 3.64 (2H, t, *J* = 6.6 Hz, ethyl 1-$H_2$), 3.16 (2H, t, *J* = 6.6 Hz, ethyl 2-$H_2$) ppm; $\delta_C$ (126 MHz, $CDCl_3$)

155.4 (pyrazine C-2), 149.3 (benzothiazole C-2), 146.34, 146.27 (benzothiazole C-3a and C-6), 145.1 (pyrazine C-3), 144.3 (pyrazine C-5), 143.0 (pyrazine C-6), 136.0 (benzothiazole C-7a), 124.0 (benzothiazole C-4), 115.0 (benzothiazole C-5), 102.3 (benzothiazole C-7), 43.3 (ethyl C-1), 34.5 (ethyl C-2) ppm; HRMS Found M+H 257.0856; $C_{13}H_{13}N_4S$ requires M+H 257.0855.

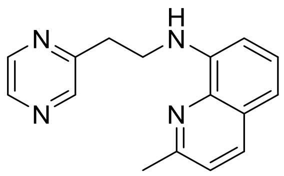

2-Methyl-*N*-[2-(pyrazin-2-yl)ethyl]quinolin-8-amine **7**

To a solution of Ir[dF($CF_3$)ppy]$_2$(dtbpy)$PF_6$ (2.2 mg, 0.002 mmol) in MeCN (1 mL) in a 7 mL scintillation vial was added 2-vinylpyrazine (10.6 mg, 0.1 mmol), 8-aminoquinaldine (43.2 mg, 0.3 mmol) and TFA (15.3 µL, 0.2 mmol). The mixture was stirred for 16 h in darkness, then concentrated *in vacuo*. The crude residue was purified by column chromatography (20:80 up to 40:60 EtOAc/hexane) to give *amine* **7** (7.1 mg, 27% yield) as a yellow oil. $\delta_H$ (500 MHz, $CDCl_3$) 8.56 (1H, dd, $J$ = 2.6, 1.5 Hz, pyrazine 5-H), 8.51 (1H, d, $J$ = 1.5 Hz, pyrazine 3-H), 8.44 (1H, d, $J$ = 2.6 Hz, pyrazine 6-H), 7.93 (1H, d, $J$ = 8.3 Hz, quinoline 4-H), 7.31 (1H, app. t, $J$ = 7.9 Hz, quinoline 6-H), 7.23 (1H, d, $J$ = 8.3 Hz, quinoline 3-H), 7.02 (1H, dd, $J$ = 8.1, 1.1 Hz, quinoline 5-H), 6.72 (1H, dd, $J$ = 7.6, 1.1 Hz, quinoline 7-H), 6.43 (1H, br. s, N-H), 3.78 (2H, t, $J$ = 7.1 Hz, ethyl 1-$H_2$), 3.26 (2H, t, $J$ = 7.1 Hz, ethyl 2-$H_2$), 2.68 (3H, s, $CH_3$) ppm; $\delta_C$ (126 MHz, $CDCl_3$) 155.8, 155.7 (quinoline C-2 and pyrazine C-2), 145.1 (pyrazine C-3), 144.4 (pyrazine C-5), 143.9 (quinoline C-8), 142.8 (pyrazine C-6), 137.7 (quinoline C-8a), 136.3 (quinoline C-4), 126.8 (quinoline C-6), 122.3 (quinoline C-3), 114.2 (quinoline C-5), 105.0 (quinoline C-7), 42.9 (ethyl C-1), 35.1 (ethyl C-2), 25.3 ($CH_3$) ppm. 15 of 16 expected signals observed; HRMS Found M+H 265.1451. $C_{16}H_{17}N_4$ requires M+H 265.1448.

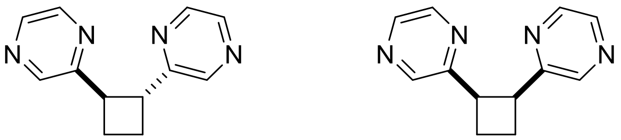

2-[(1*R**,2*R**)-2-(Pyrazin-2-yl)cyclobutyl]pyrazine *anti*-8 and 2-[(1*R**,2*S**)-2-(pyrazin-2-yl)cyclobutyl]pyrazine *syn*-8

To a solution of Ir[dF($CF_3$)ppy]$_2$(dtbpy)$PF_6$ (2.2 mg, 0.002 mmol) in MeCN (1 mL) in a 7 mL scintillation vial was added 2-vinylpyrazine (31.8 mg, 0.3 mmol). The mixture was stirred for 16 h with a blue LED lamp ~10 cm away, then concentrated *in vacuo*. The crude residue was purified by column chromatography (98:2 $CH_2Cl_2$/MeOH) to give *cyclobutanes anti*-8 and *syn*-8 as an inseparable mixture of diastereomers. *anti*-8 and *syn*-8 (14.9 mg, 47% yield, 73:27 d.r) as a yellow oil. $\delta_H$ (500 MHz, $CDCl_3$) 8.54 (1.5H, dd, *J* = 2.6, 1.5 Hz, pyrazine 5-$H^{anti}$ and 5'-$H^{anti}$ ), 8.42 (1.5H, d, *J* = 1.5 Hz, pyrazine 3-$H^{anti}$ and 3'-$H^{anti}$), 8.40 (1.5H, d, *J* = 2.6 Hz, pyrazine 6-$H^{anti}$ and 6'-$H^{anti}$), 8.27 (0.5H, dd, *J* = 2.5, 1.6 Hz, pyrazine 5-$H^{syn}$ and 5'-$H^{syn}$), 8.26 (0.5H, d, *J* = 1.6 Hz, pyrazine 3-$H^{syn}$ and 3'-$H^{syn}$), 8.17 (0.5H, d, *J* = 2.5 Hz, pyrazine 6-$H^{syn}$ and 6'-$H^{syn}$), 4.31-4.24 (0.5H, m, cyclobutane 1-$H^{syn}$ and 2-$H^{syn}$), 4.10-4.00 (1.5H, m, cyclobutane 1-$H^{anti}$ and 2-$H^{anti}$), 2.85-2.74 (0.5H, m, cyclobutane 3-$H_a^{syn}$ and 4-$H_a^{syn}$), 2.60-2.51 (0.5H, m, cyclobutane 3-$H_b^{syn}$ and 4-$H_b^{syn}$), 2.49-2.35 (3H, m, cyclobutane 3-$H_2^{anti}$ and 4-$H_2^{anti}$) ppm; $\delta_C$ (126 MHz, $CDCl_3$) 157.9 (pyrazine C-$2^{anti}$), 156.7 (pyrazine C-$2^{syn}$), 144.6 (pyrazine C-$5^{syn}$), 144.5 (pyrazine C-$5^{anti}$), 143.9 (pyrazine C-$3^{anti}$), 143.8 (pyrazine C-$3^{syn}$), 142.8 (pyrazine C-$6^{anti}$), 141.9 (pyrazine C-$6^{syn}$), 44.8 (cyclobutane C-$1^{anti}$), 43.9 (cyclobutane C-$1^{syn}$), 24.6 (cyclobutane C-$3^{anti}$), 23.0 (cyclobutane C-$3^{syn}$) ppm; HRMS Found M+H 213.1133. $C_{12}H_{15}N_2S_2$ requires M+H 213.1135.

By analogy with the [2+2] cycloaddition of 4-methyl-5-vinylthiazole, the configuration of the major diastereomer was identified as the *anti*-isomer.

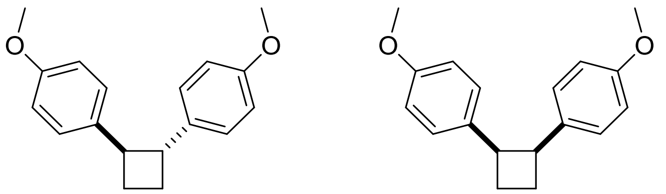

1-Methoxy-4-[(1*S**,2*S**)-2-(4-methoxyphenyl)cyclobutyl]benzene *anti*-9 and 1-methoxy-4-[(1*S**,2*R**)-2-(4-methoxyphenyl)cyclobutyl]benzene *syn*-9

To a solution of Ir[dF($CF_3$)ppy]$_2$(dtbpy)$PF_6$ (2.2 mg, 0.002 mmol) in MeCN (1 mL) in a 7 mL scintillation vial was added 4-methoxystyrene (40.2 mg, 0.3 mmol). The mixture was stirred for 16 h with a blue LED lamp ~10 cm away, then concentrated *in vacuo*. The crude residue was purified by column chromatography (98:2 hexane/acetone) to give *cyclobutanes anti*-9 and *syn*-9 as an inseparable mixture of diastereomers. *anti*-9 and *syn*-9 (7.9 mg, 20% yield, 86:14 d.r) as a yellow oil. $\delta_H$ (400 MHz, $CDCl_3$) 7.18-7.11 (3.44H, m, phenyl 3-$H_2^{anti}$ and 2'-$H_2^{anti}$), 6.88-6.78 (4H, m, phenyl 2-$H_2^{anti}$, 3'-$H_2^{anti}$, 3-$H_2^{syn}$ and 2'-$H_2^{syn}$), 6.69-6.62 (0.56H, m, phenyl 2-$H_2^{syn}$ and 3'-$H_2^{syn}$), 3.95-3.87 (0.28H, m, cyclobutane 1-$H^{syn}$ and 2-$H^{syn}$), 3.79 (5.16H, s, $(OMe)_2^{anti}$), 3.71 (0.84H, s, $(OMe)_2^{syn}$) 3.49-3.39 (1.72H, m, cyclobutane 1-$H^{anti}$ and 2-$H^{anti}$), 2.48-2.32 (0.56H, m, cyclobutane 3-$H_2^{syn}$ and 4-$H_2^{syn}$), 2.32-2.22 (1.72H, m, cyclobutane 3-$H_a^{anti}$ and 4-$H_a^{anti}$), 2.12-2.01 (1.72H, m, cyclobutane 3-$H_b^{anti}$ and 4-$H_b^{anti}$) ppm. $^1$H NMR for the major component (*anti*-9) was consistent with the reported literature (M. Riener and D. A. Nicewicz, *Chem. Sci.* 2013, 4, 2625-2629).

### NMR Spectra

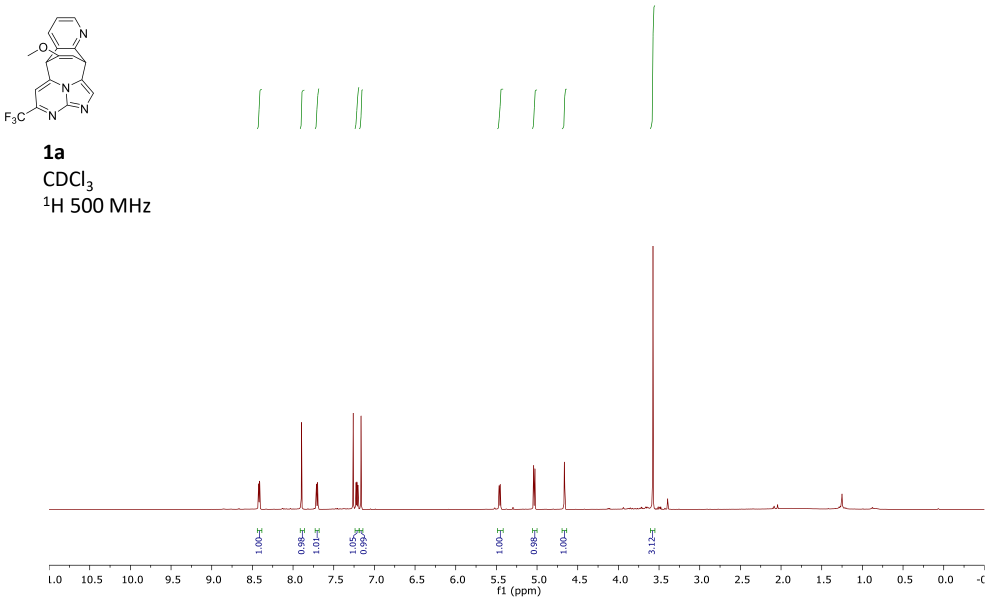

**1a**
$CDCl_3$
$^{1}H$ 500 MHz

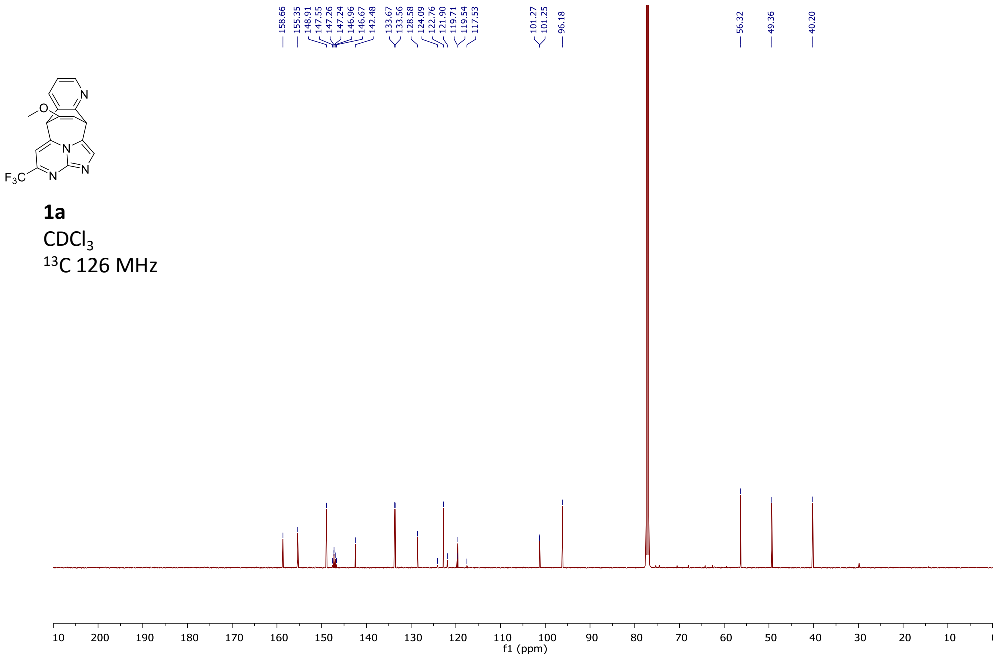

**1a**
$CDCl_3$
$^{13}C$ 126 MHz

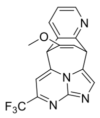

**1a**
$CDCl_3$
$^{19}F$ 376 MHz

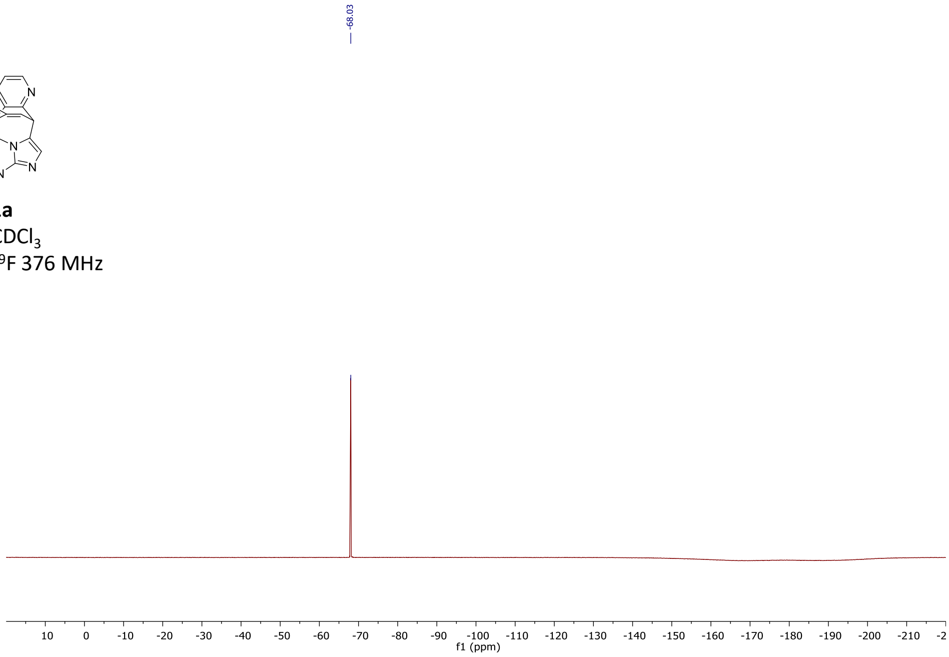

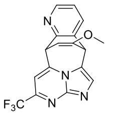

**1b**
$CD_3CN$
$^1H$ 500 MHz

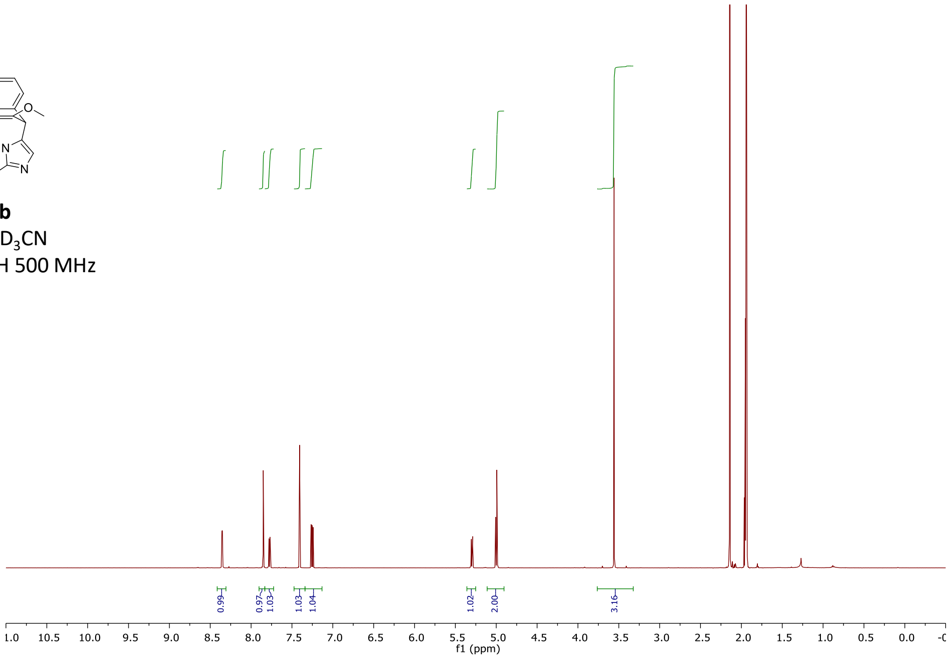

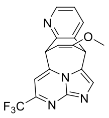

**1b**
$CD_3CN$
$^{13}C$ 126 MHz

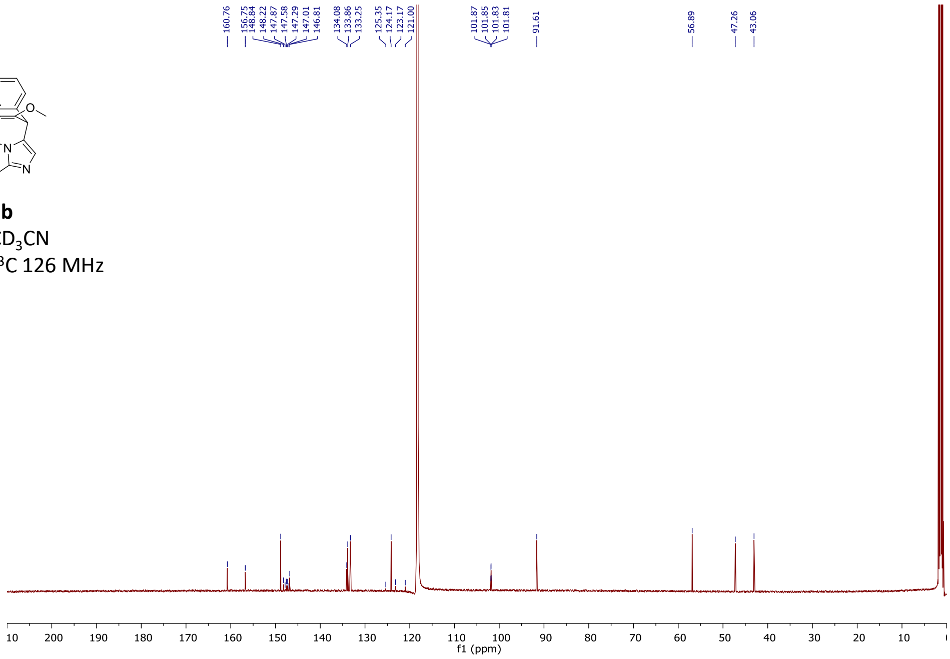

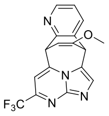

**1b**
$CD_3CN$
$^{19}F$ 376 MHz

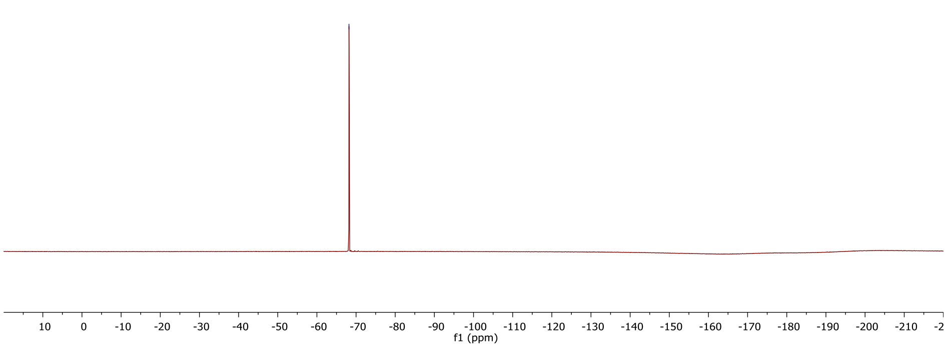

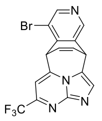

**2**
$CDCl_3$
$^1H$ 500 MHz

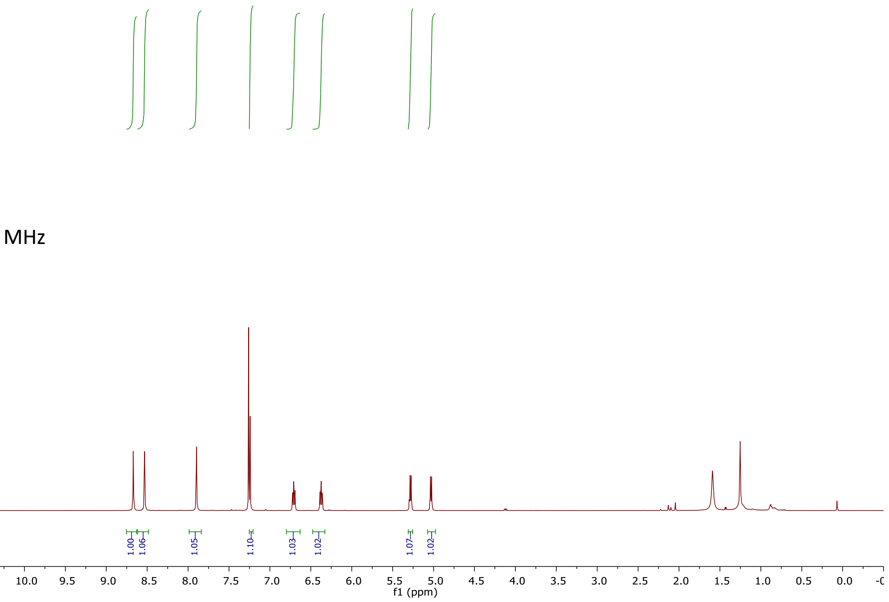

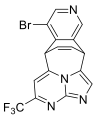

**2**
$CDCl_3$
$^{13}C$ 126 MHz

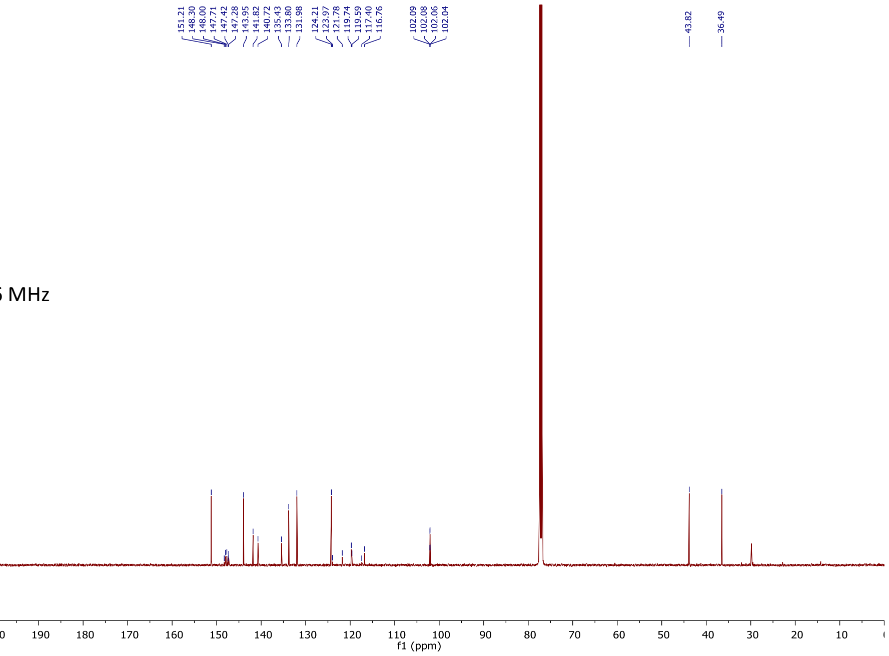

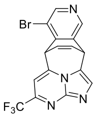

**2**
$CDCl_3$
$^{19}F$ 376 MHz

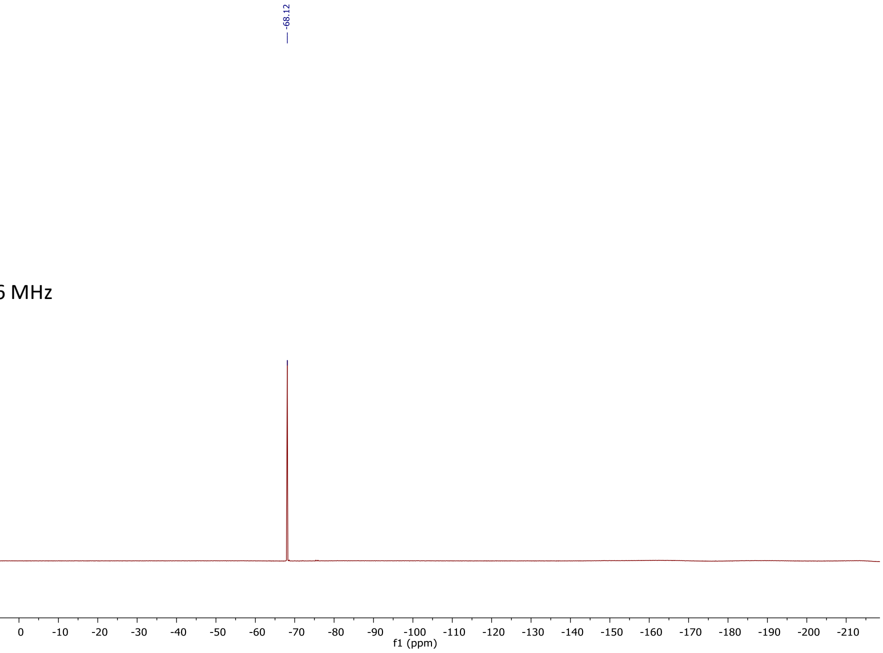

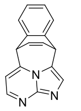

**3**
$CD_3CN$
$^{1}H$ 500 MHz

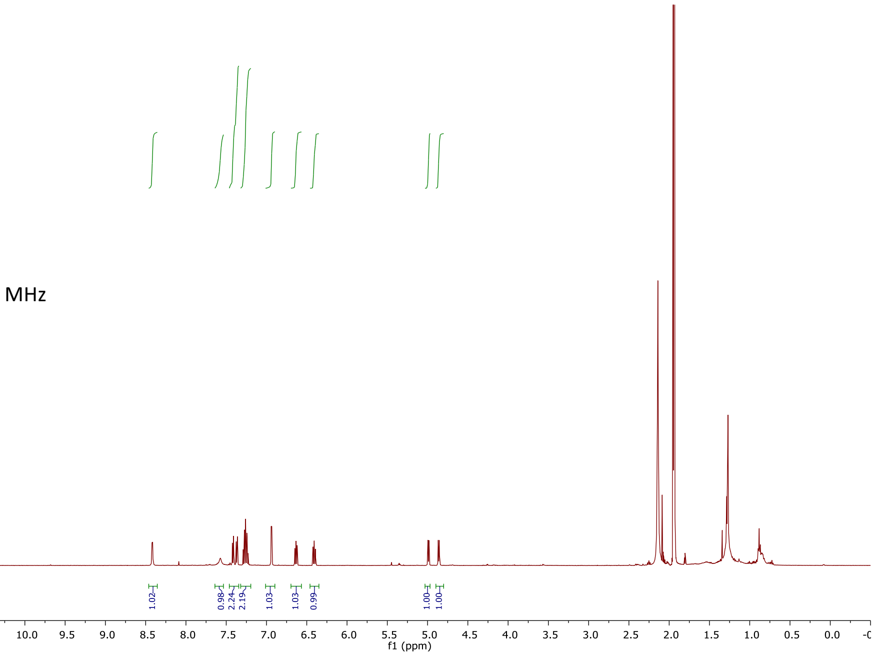

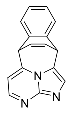

**3**
$CD_3CN$
$^{13}C$ 126 MHz

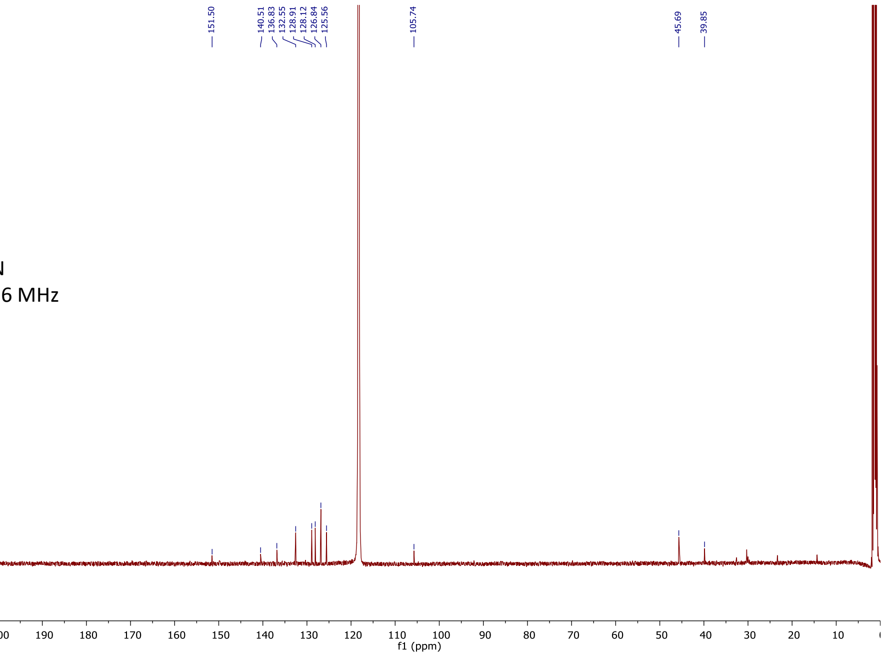

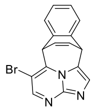

**4**
$CDCl_3$
$^{1}H$ 500 MHz

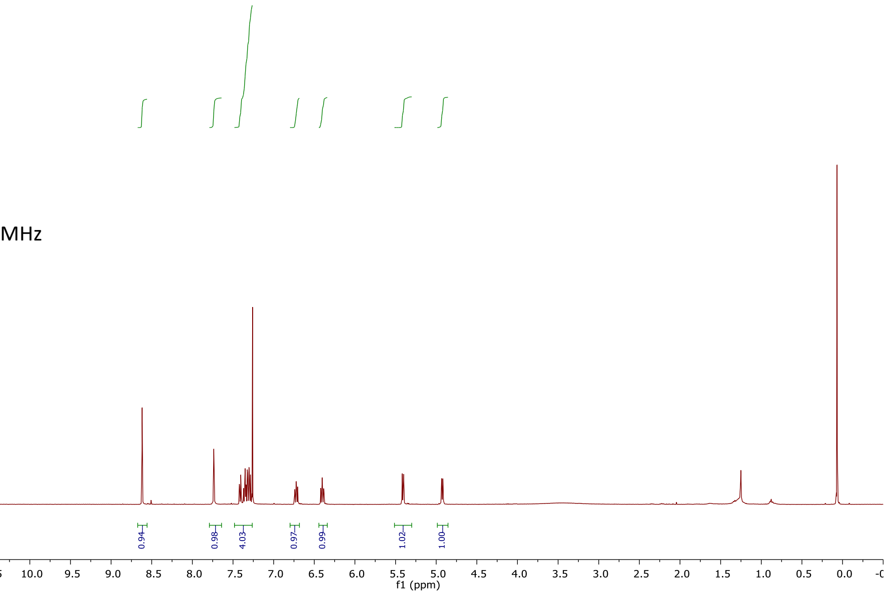

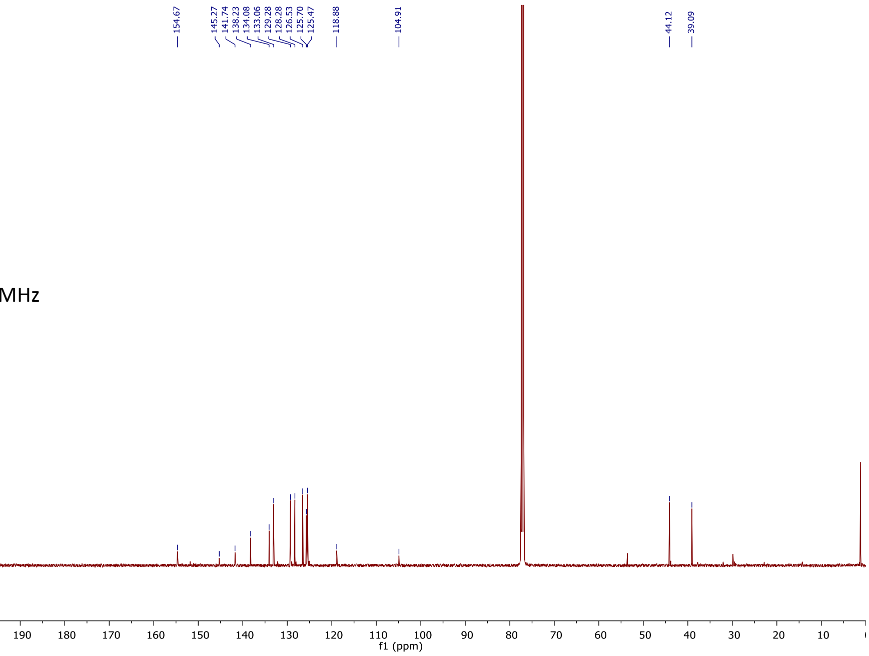

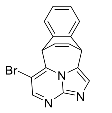

**4**
$CDCl_3$
$^{13}C$ 126 MHz

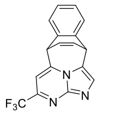

**5**
$CDCl_3$
$^{1}H$ 500 MHz

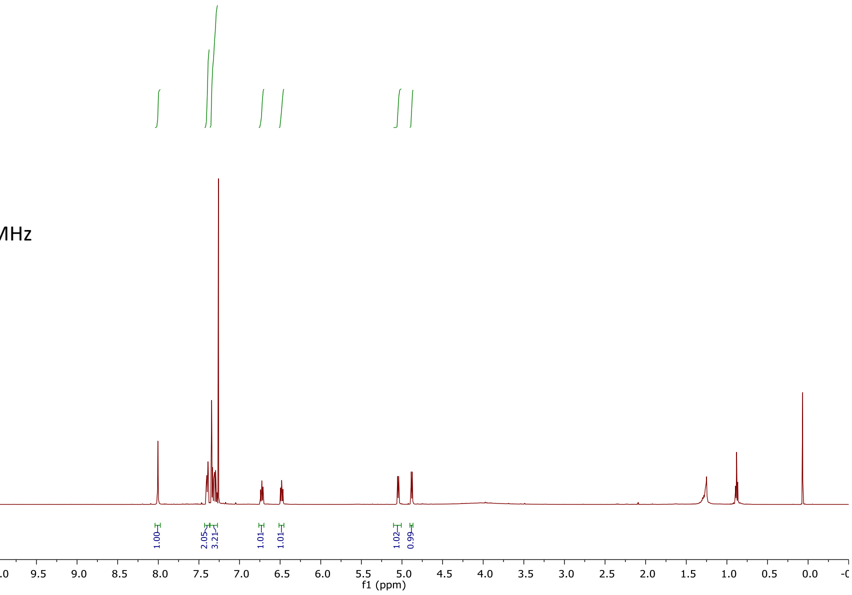

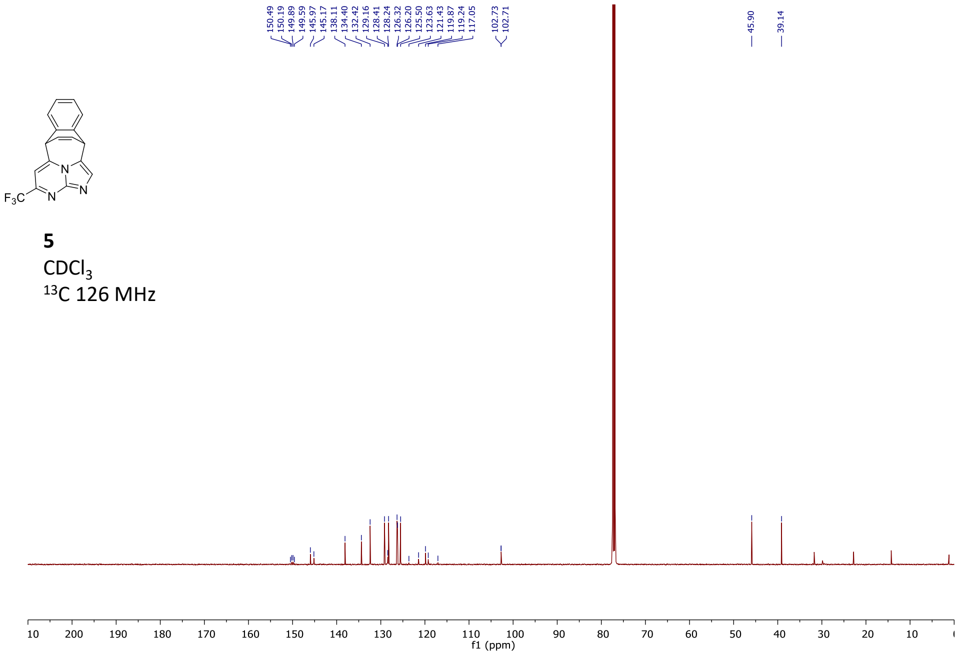

**5**
$CDCl_3$
$^{13}C$ 126 MHz

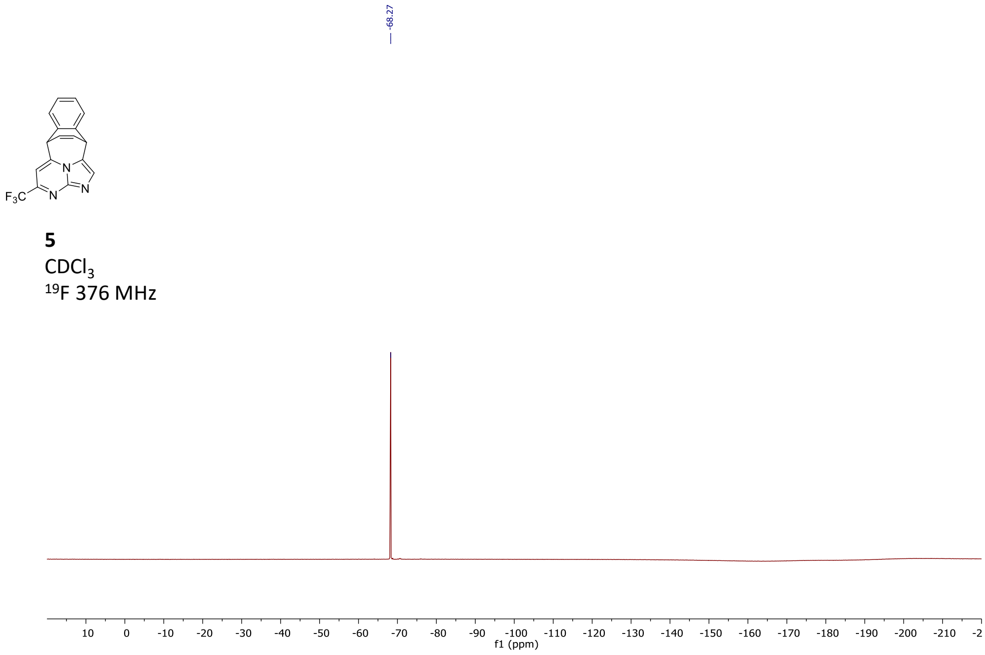

**5**
$CDCl_3$
$^{19}F$ 376 MHz

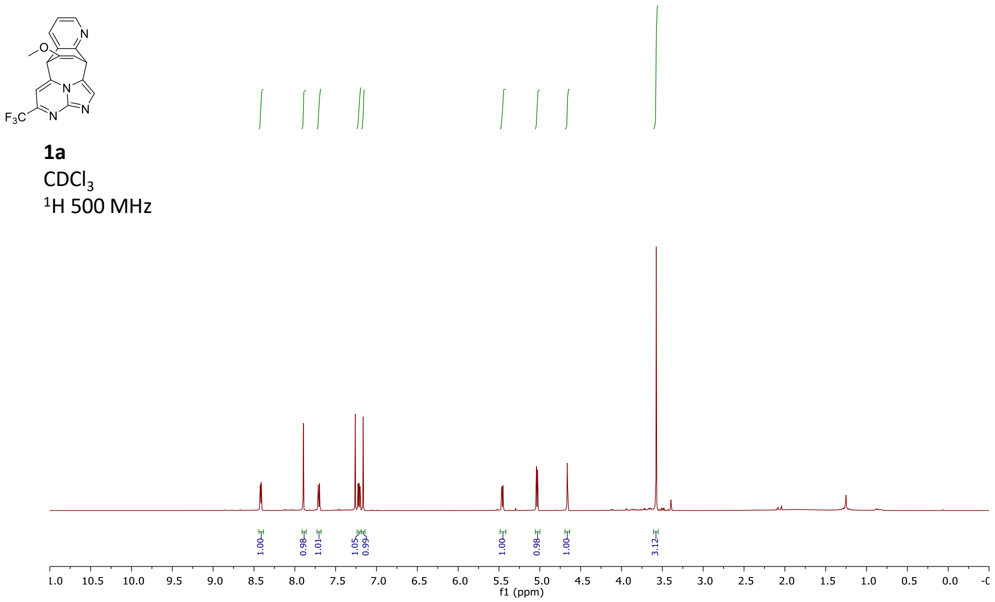

F$_3$C
1a
CDCl$_3$
$^1$H 500 MHz
1.00
0.98
1.01
1.05
0.99
1.00
0.98
1.00
3.12
f1 (ppm)

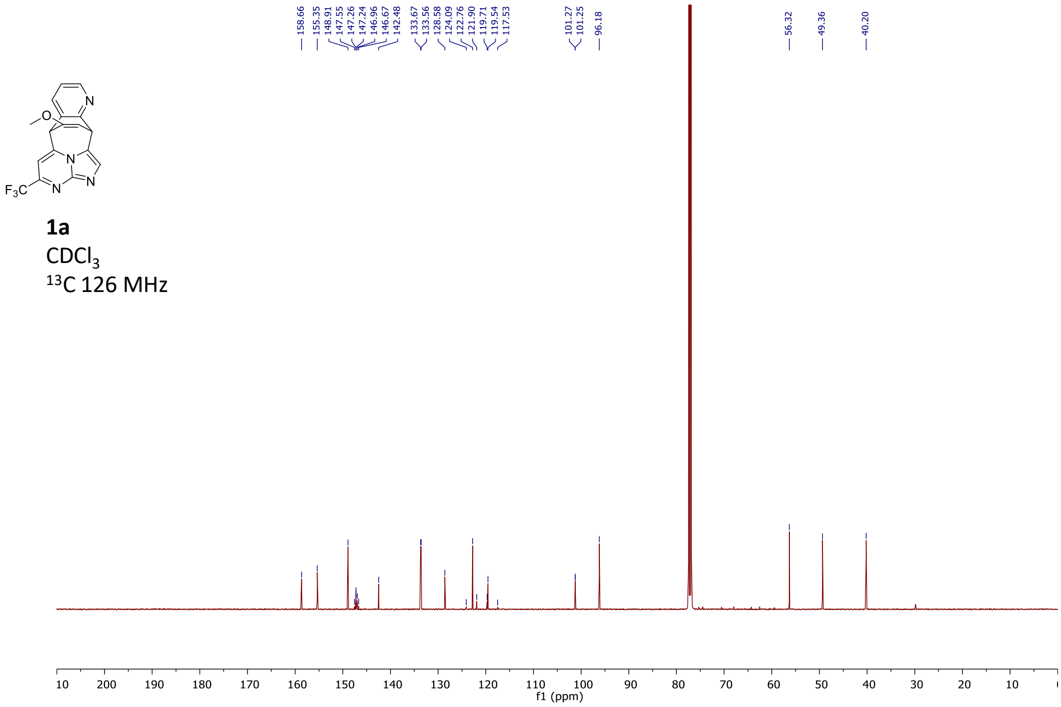

158.66
155.35
148.91
147.55
147.26
147.24
146.96
146.67
142.48
133.67
133.56
128.58
124.09
122.76
121.90
119.71
119.54
117.53
101.27
101.25
96.18
56.32
49.36
40.20
F$_3$C
1a
CDCl$_3$
$^{13}$C 126 MHz
f1 (ppm)

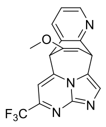

**1a**
$CDCl_3$
$^{19}F$ 376 MHz

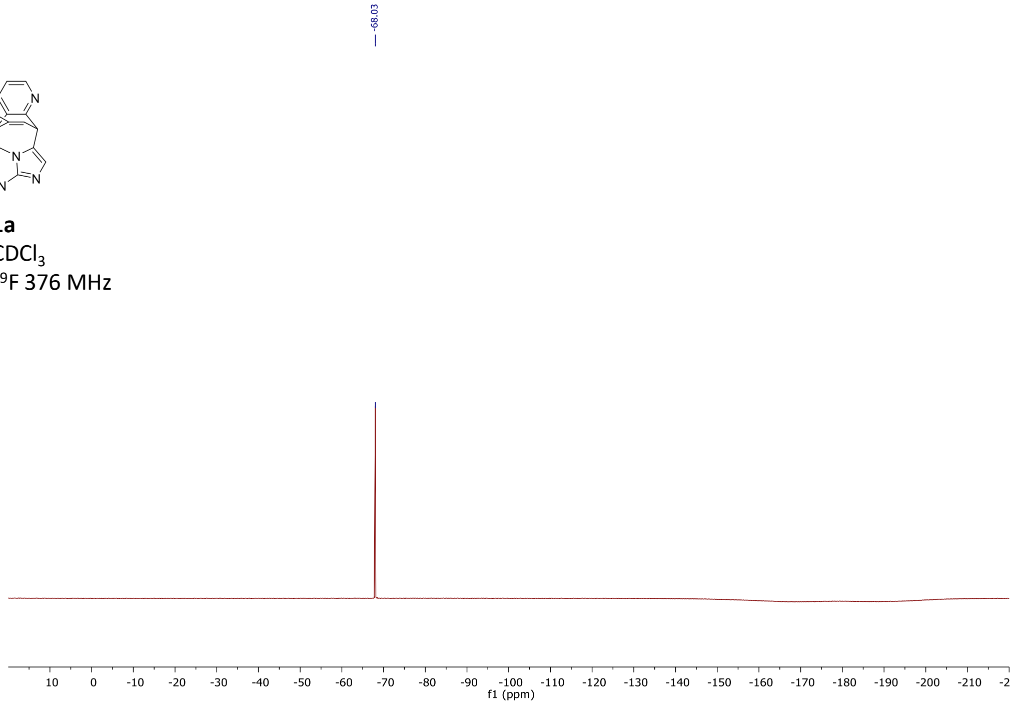

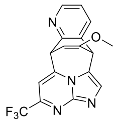

**1b**
$CD_3CN$
$^{1}H$ 500 MHz

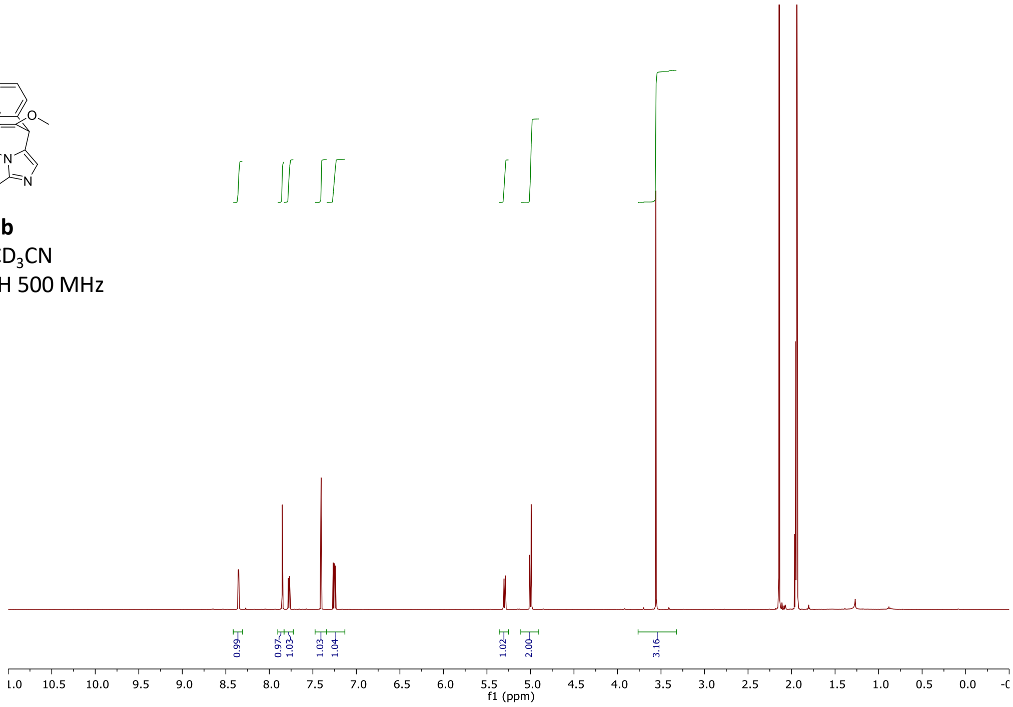

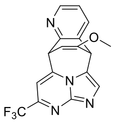

**1b**
$CD_3CN$
$^{13}C$ 126 MHz

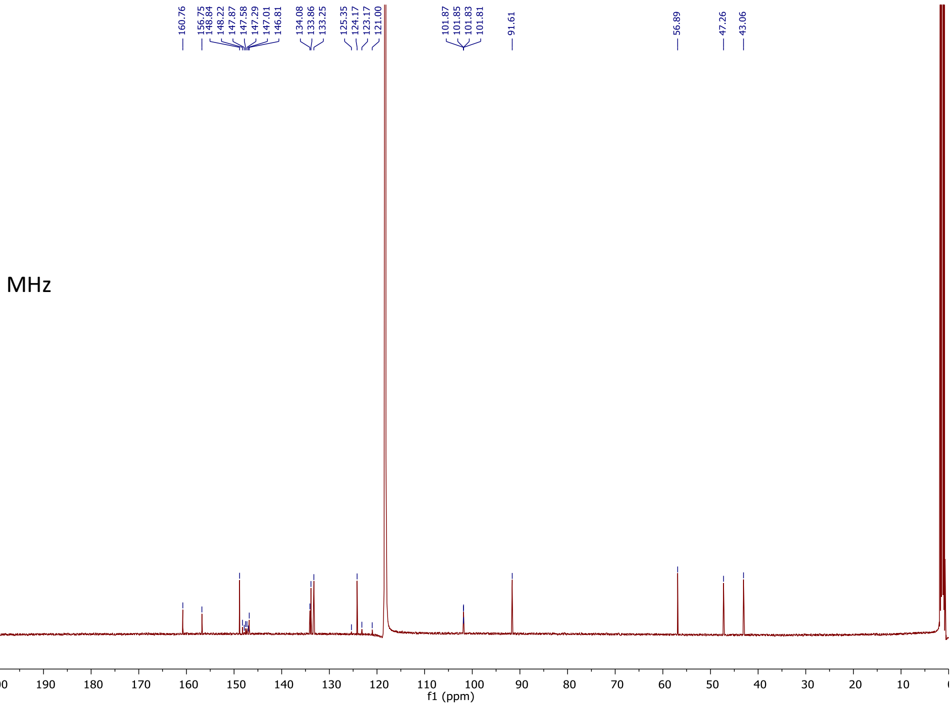

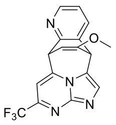

**1b**
$CD_3CN$
$^{19}F$ 376 MHz

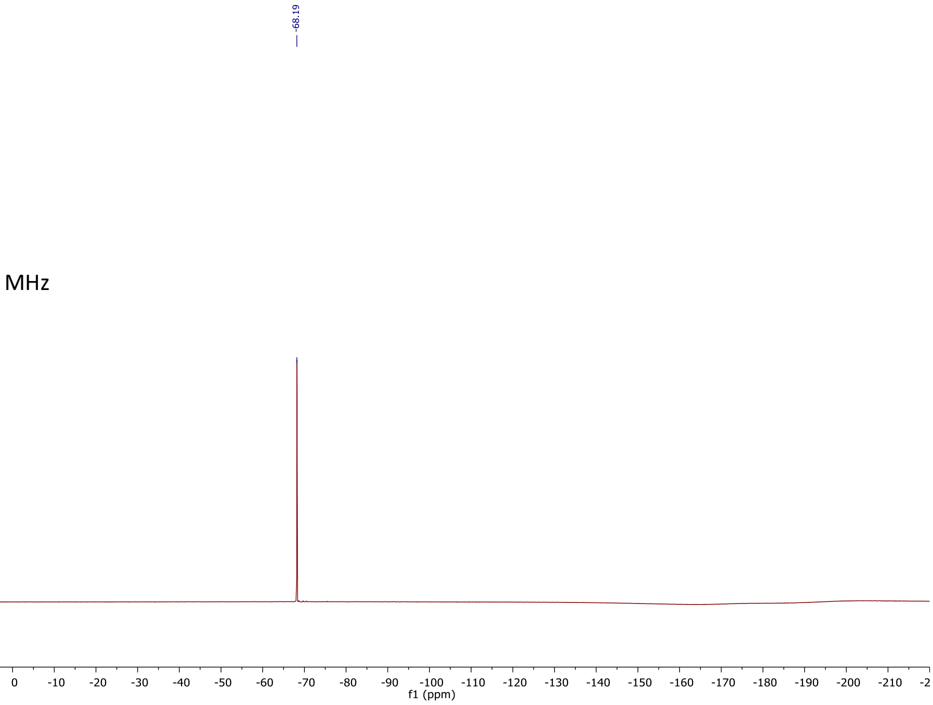

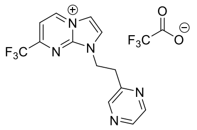

**2**
$D_2O$
$^{1}H$ 500 MHz

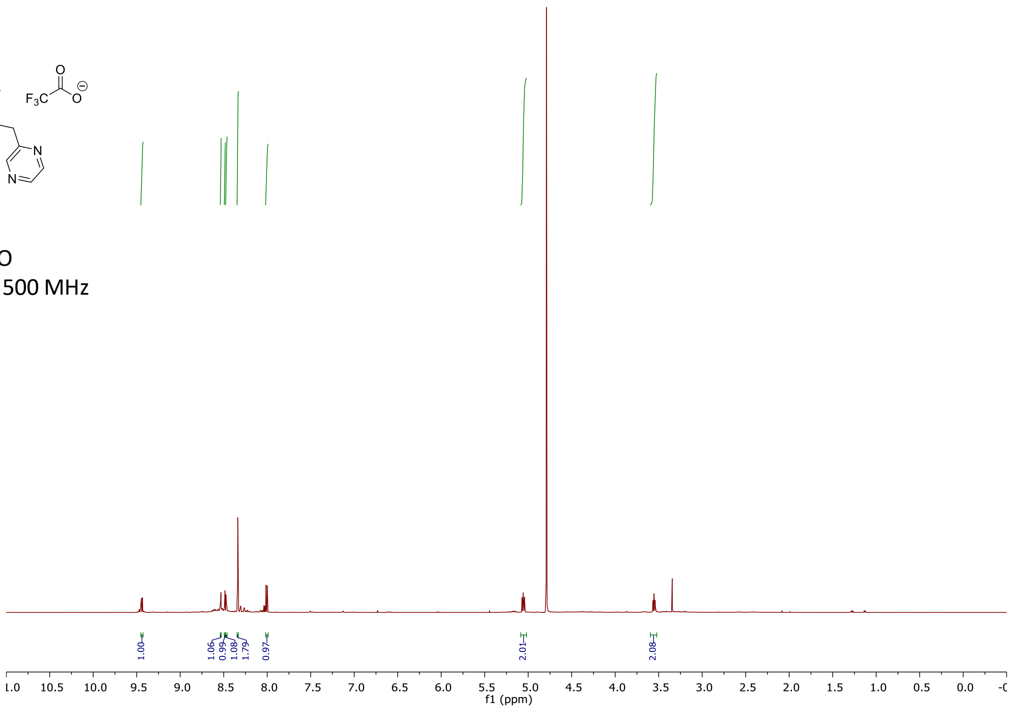

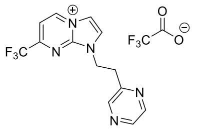

**2**
$D_2O$
$^{13}C$ 126 MHz

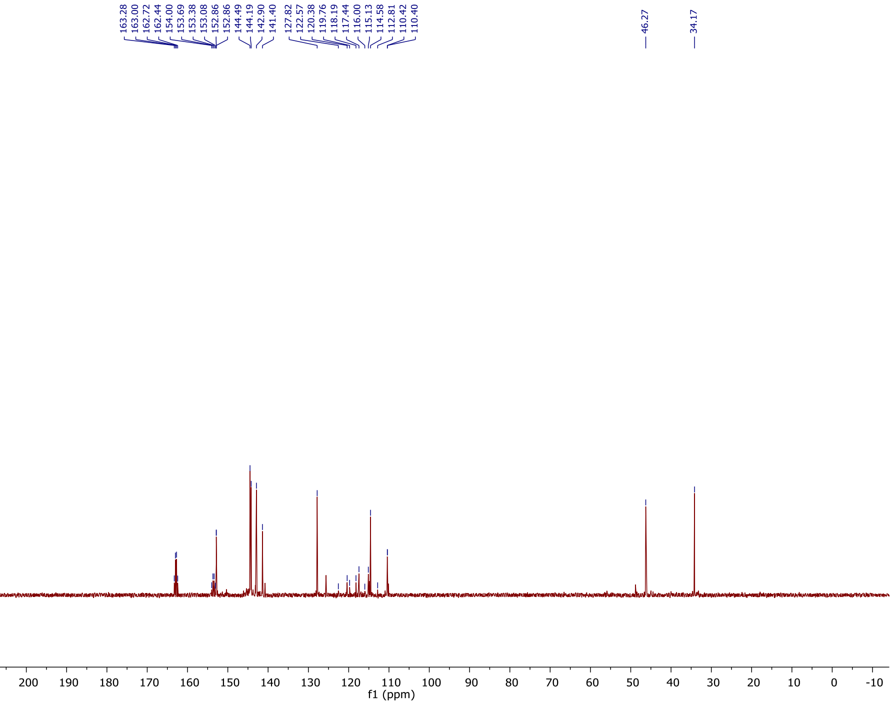

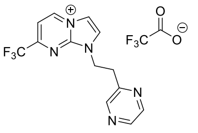

**2**
$D_2O$
$^{19}F$ 376 MHz

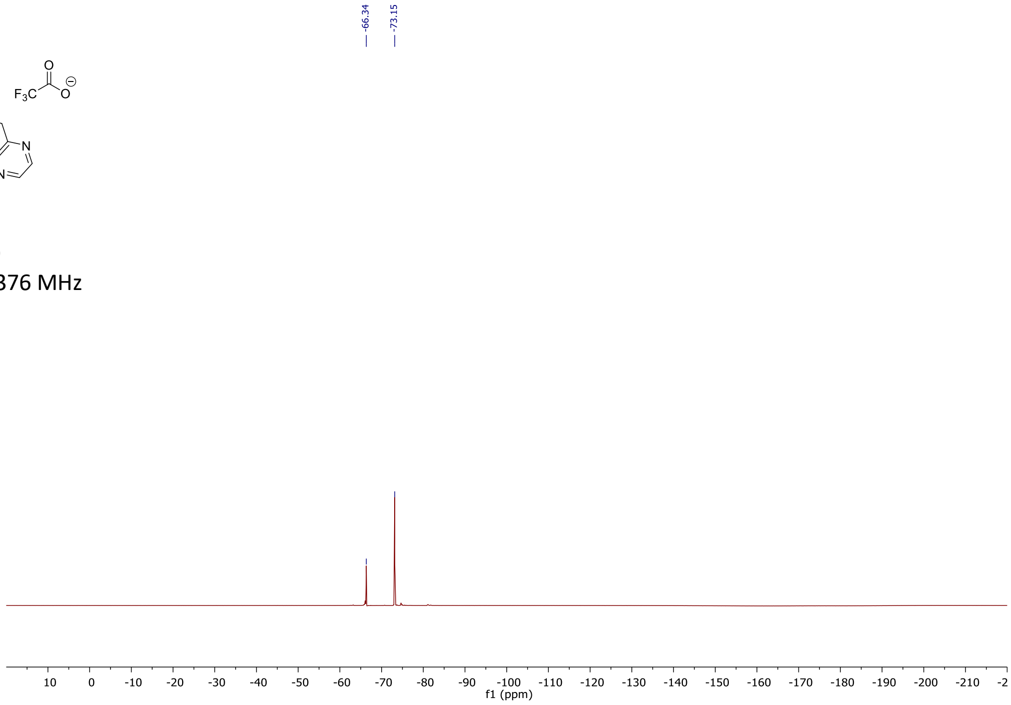

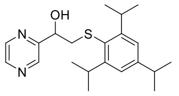

**3**
$CDCl_3$
$^{1}H$ 500 MHz

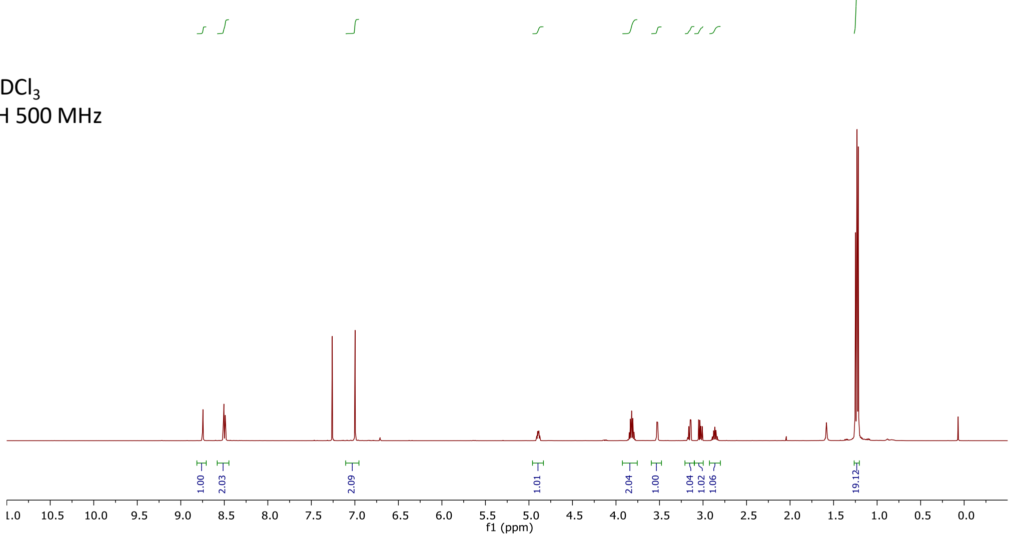

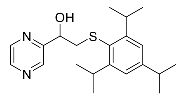

**3**
$CDCl_3$
$^{13}C$ 126 MHz

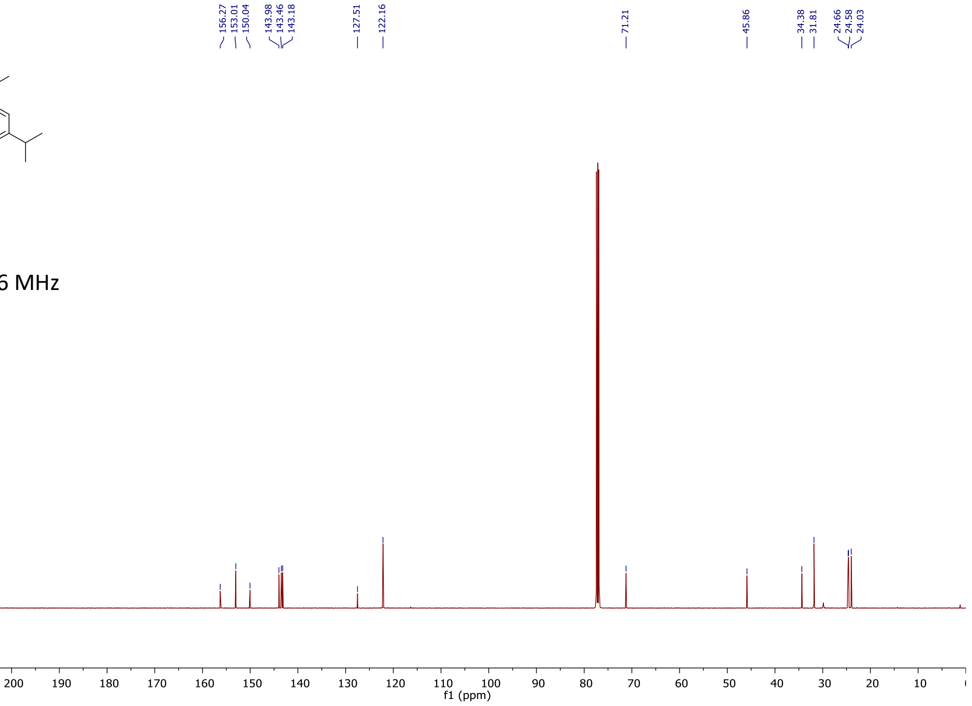

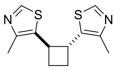

***anti*-4**
$CDCl_3$
$^{1}H$ 500 MHz

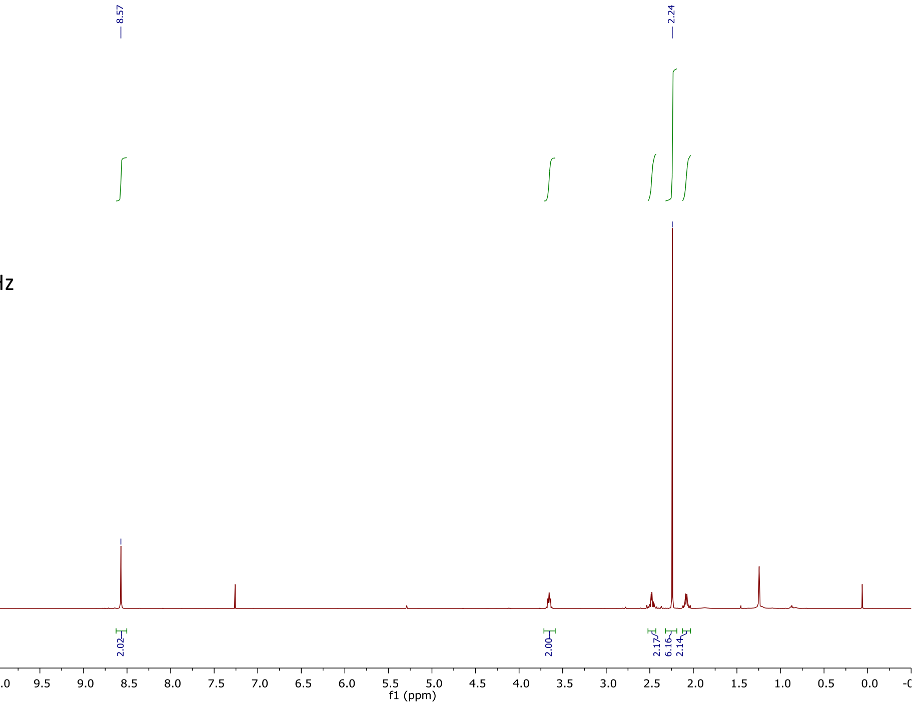

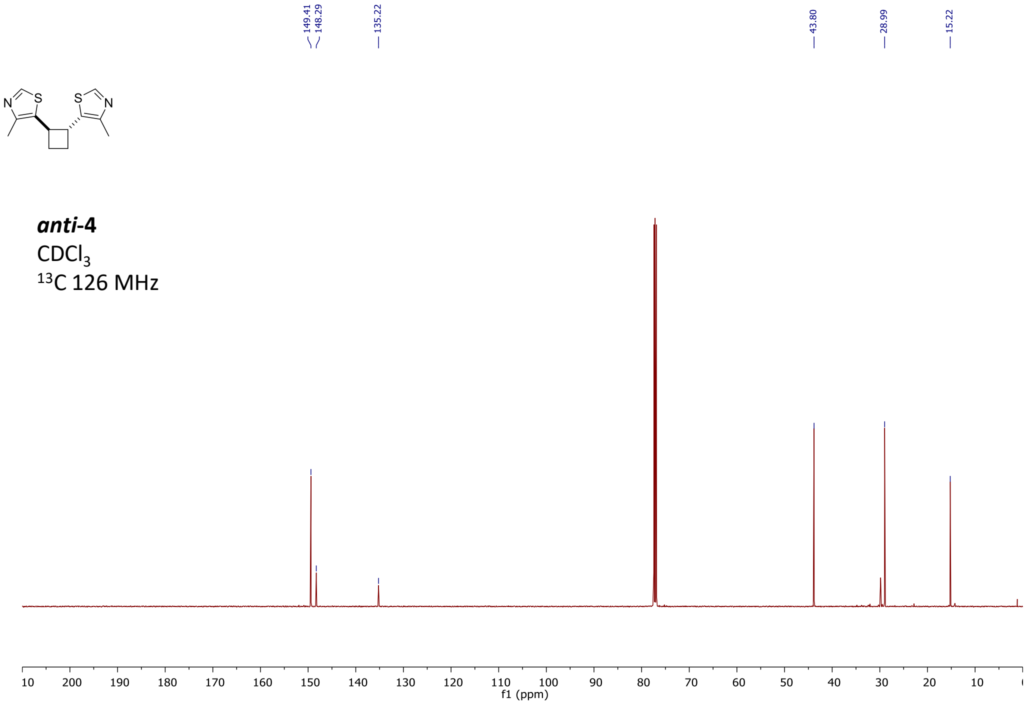

149.41
148.29
135.22
43.80
28.99
15.22
N
S
S
N
anti-4
CDCl3
13C 126 MHz
f1 (ppm)

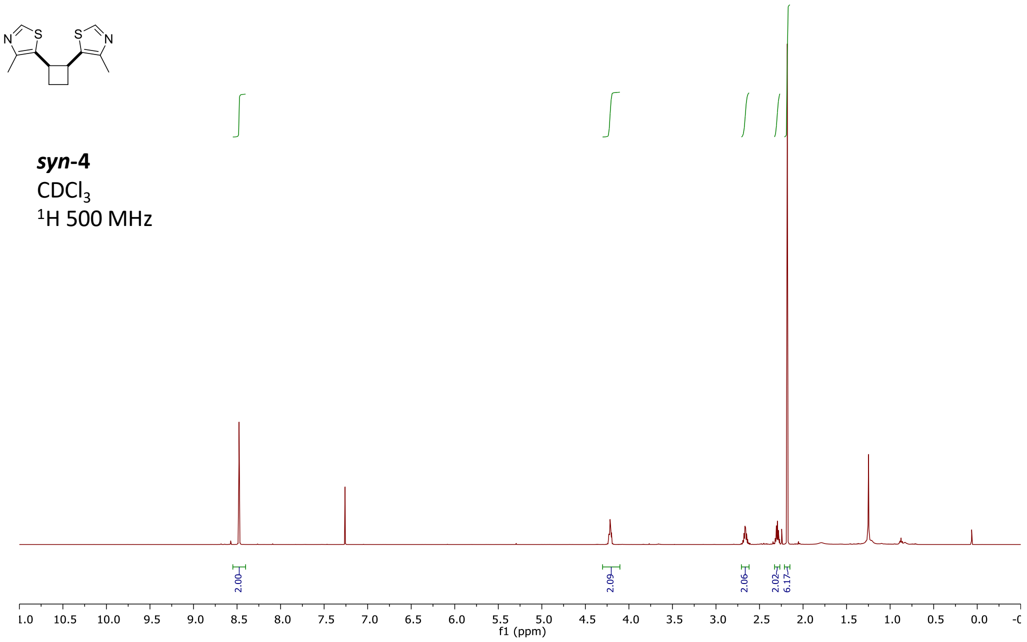

N
S
S
N
syn-4
CDCl3
1H 500 MHz
2.00
2.09
2.06
2.02
6.17
f1 (ppm)

***syn*-4**
$CDCl_3$
$^{13}C$ 126 MHz

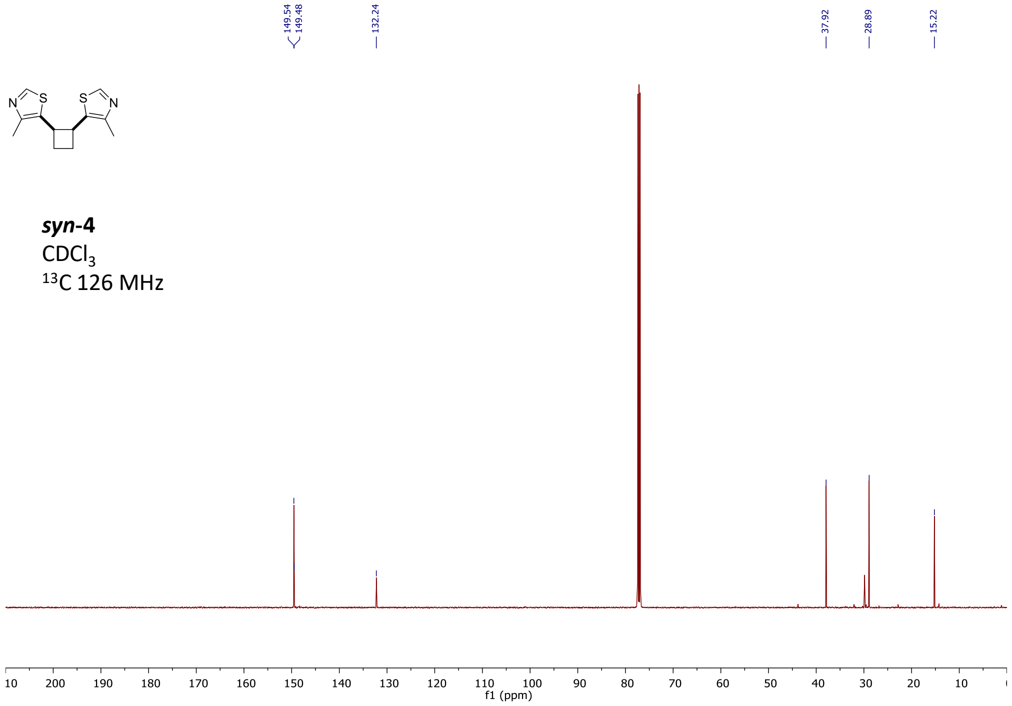

**5**
$CDCl_3$
$^{1}H$ 500 MHz

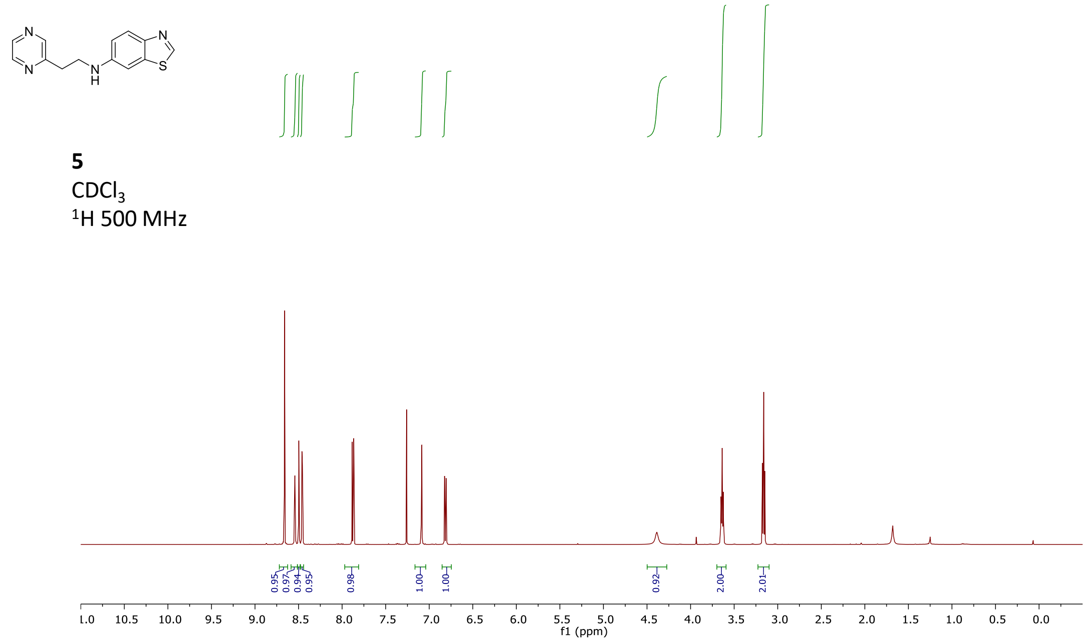

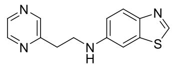

**5**
$CDCl_3$
$^{13}C$ 126 MHz

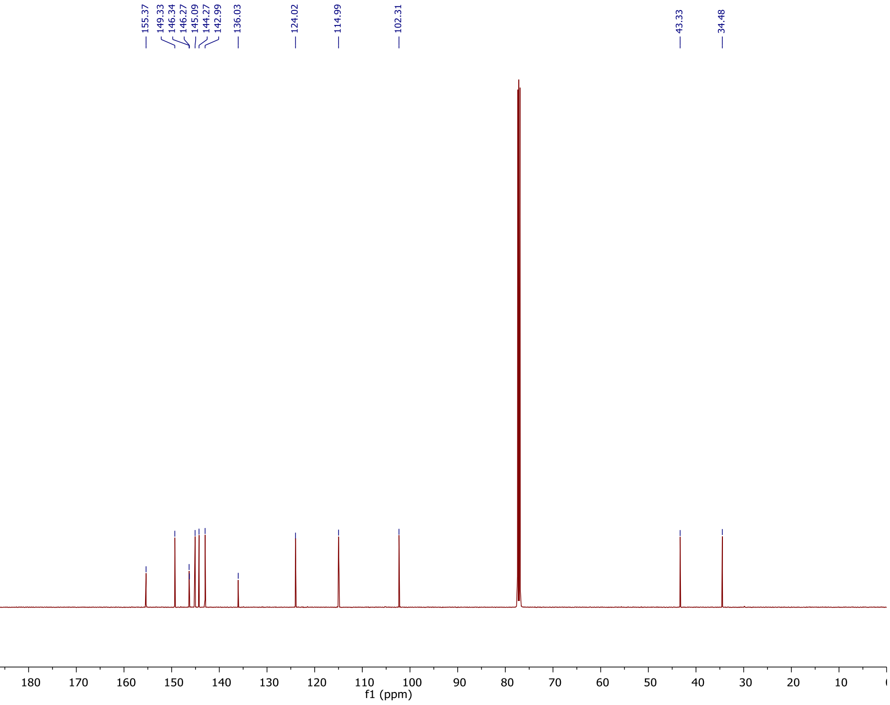

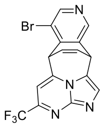

**6**
$CDCl_3$
$^{1}H$ 500 MHz

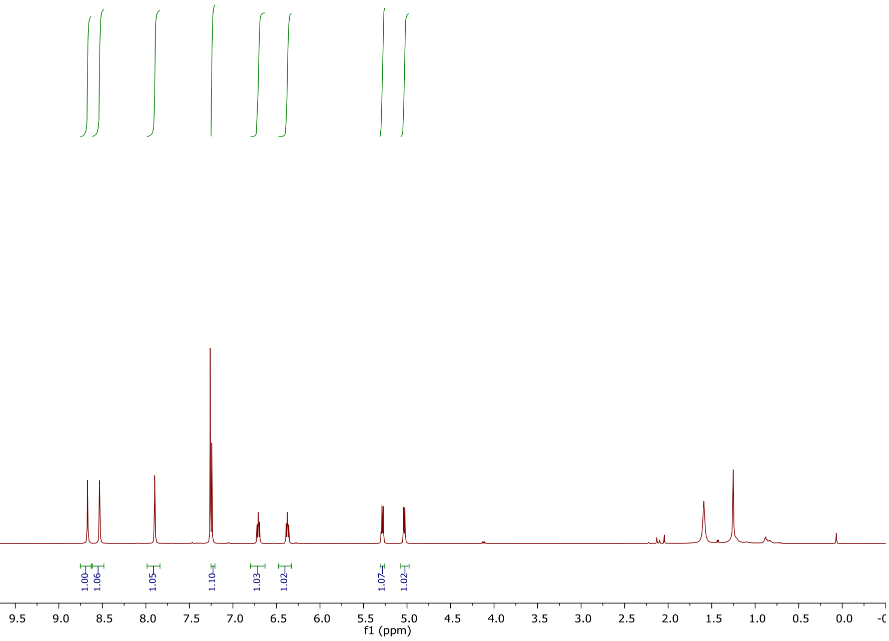

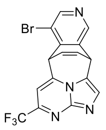

**6**
$CDCl_3$
$^{13}C$ 126 MHz

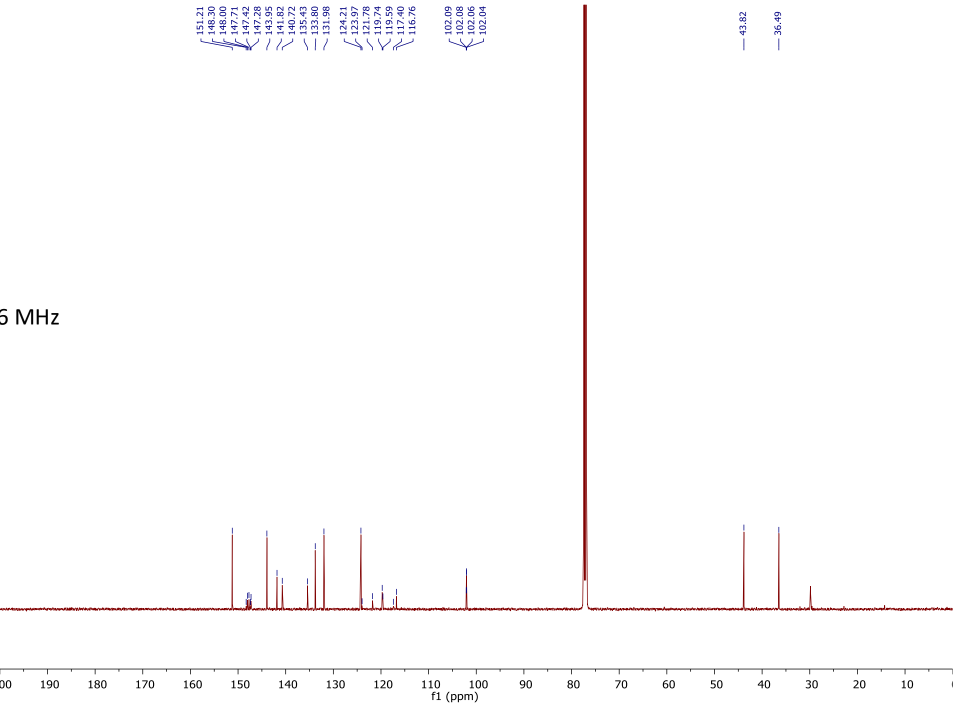

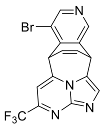

**6**
$CDCl_3$
$^{19}F$ 376 MHz

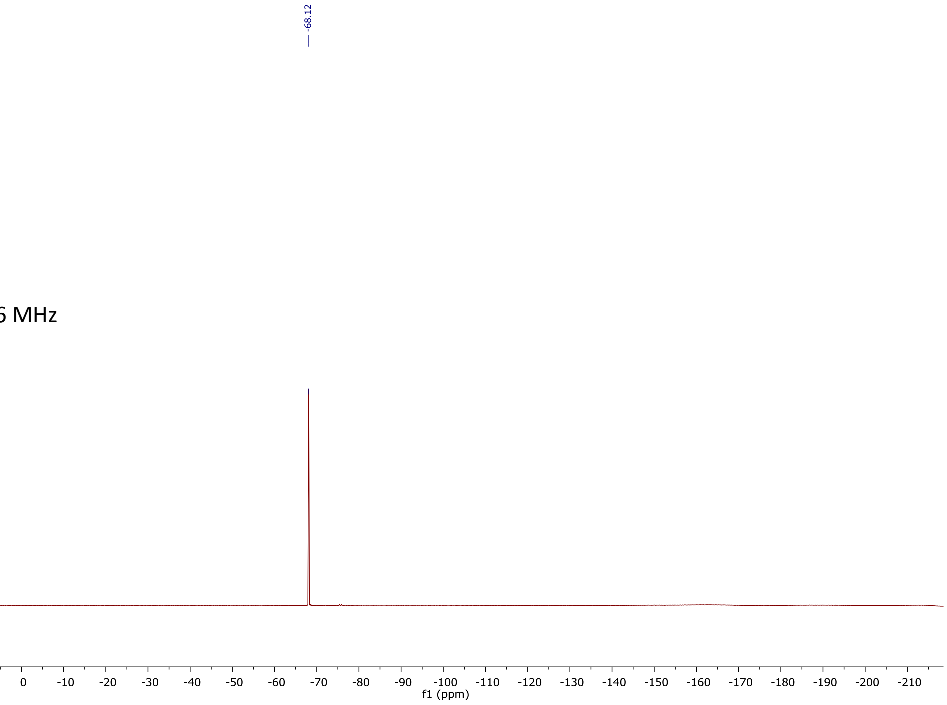

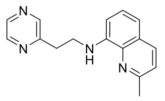

**7**
$CDCl_3$
$^1H$ 500 MHz

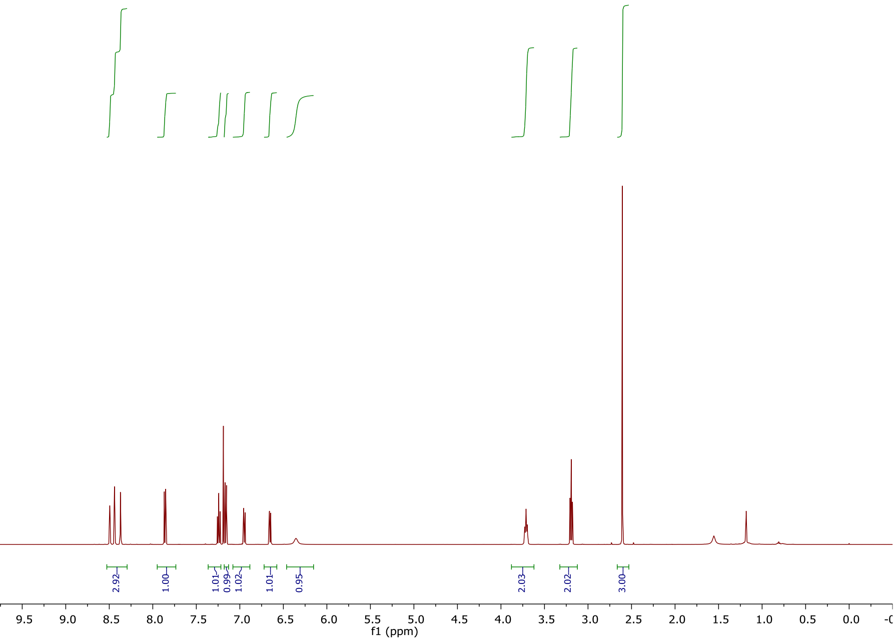

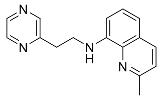

**7**
$CDCl_3$
$^{13}C$ 126 MHz

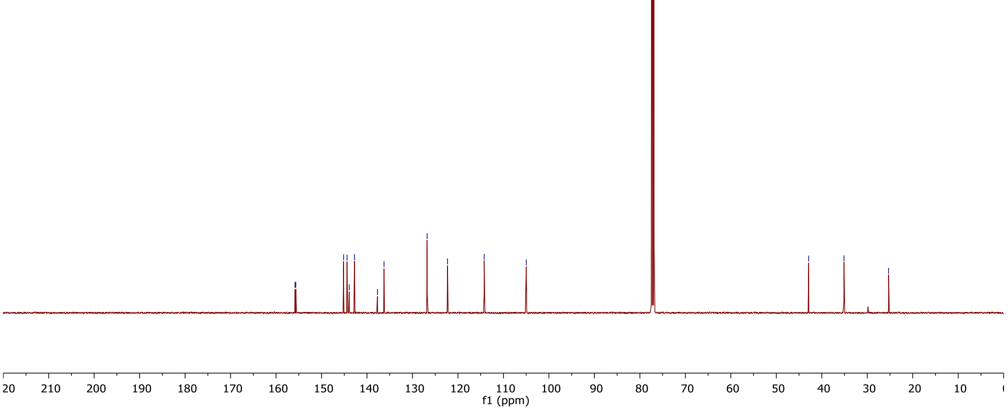

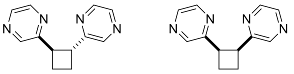

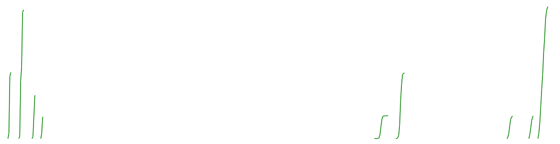

***anti*-8, *syn*-8**
$CDCl_3$
[1]H 500 MHz

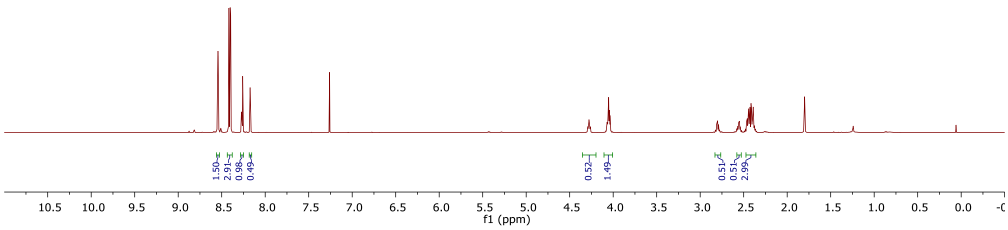

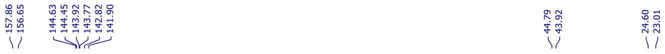

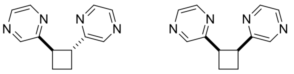

***anti*-8, *syn*-8**
$CDCl_3$
[13]C 126 MHz

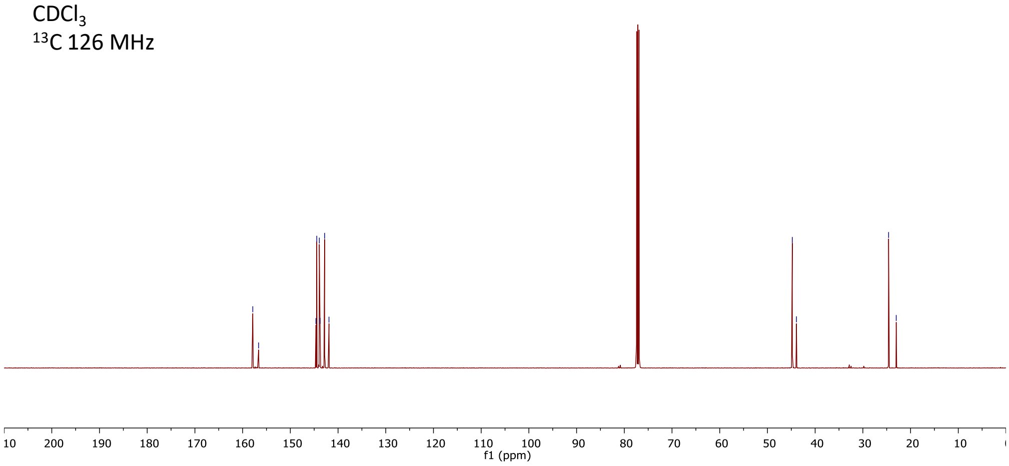

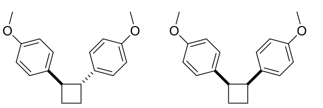

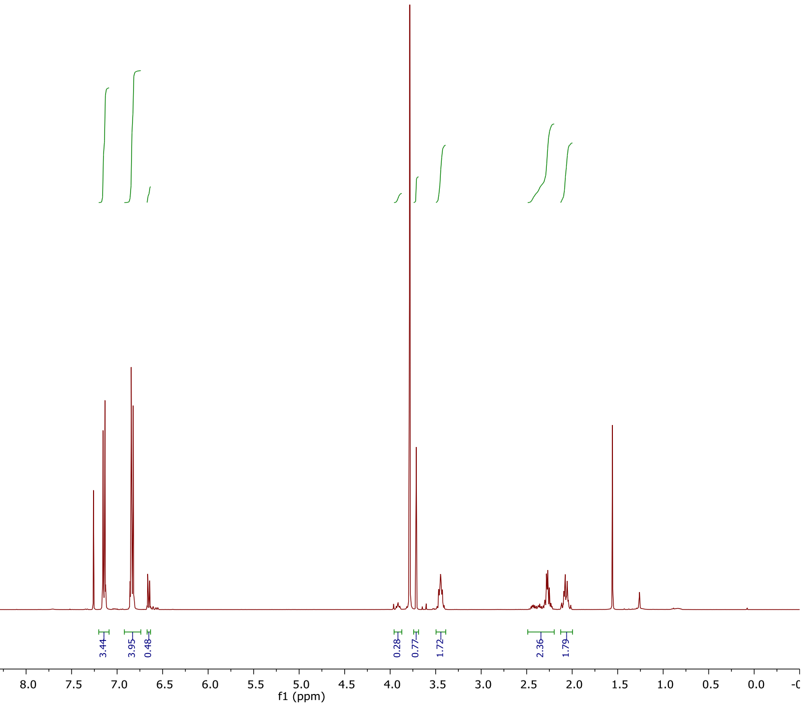

***anti*-9, *syn*-9**
$CDCl_3$
$^1H$ 400 MHz

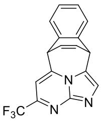

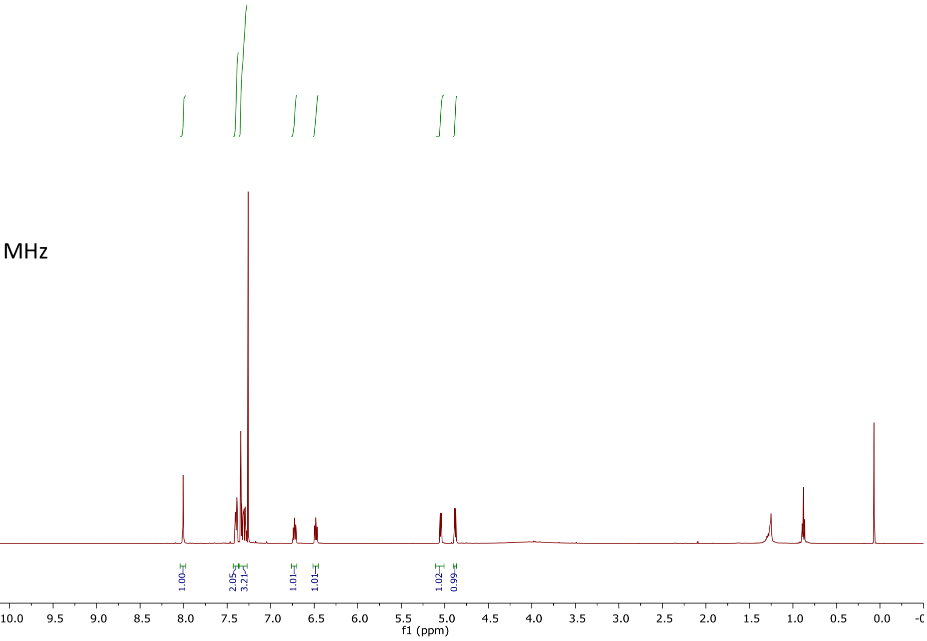

**10**
$CDCl_3$
$^1H$ 500 MHz

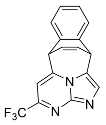

**10**
$CDCl_3$
$^{13}C$ 126 MHz

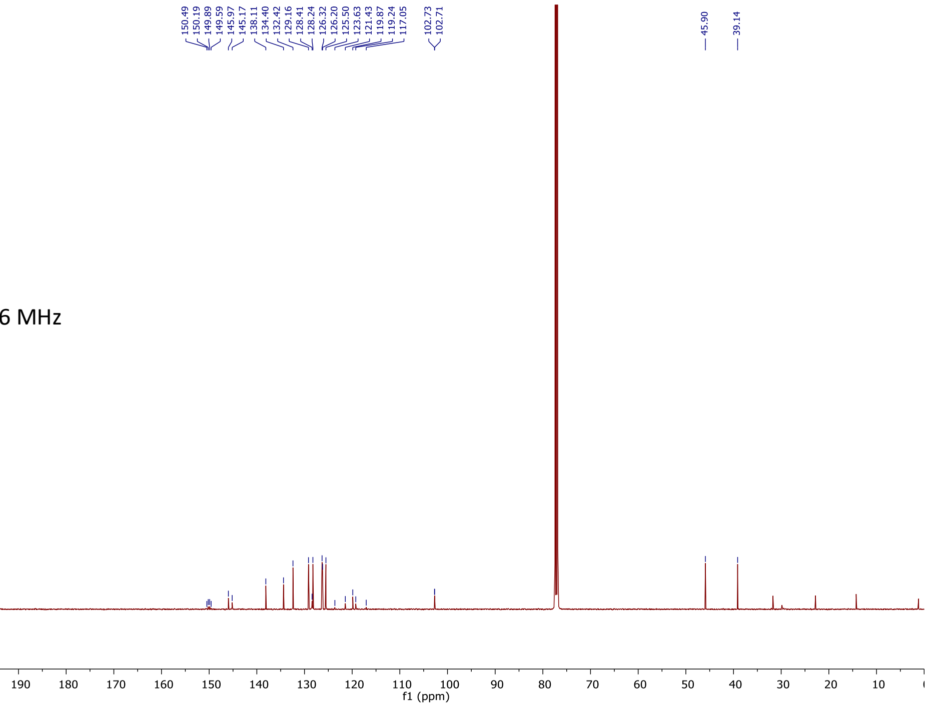

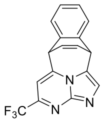

**10**
$CDCl_3$
$^{19}F$ 376 MHz

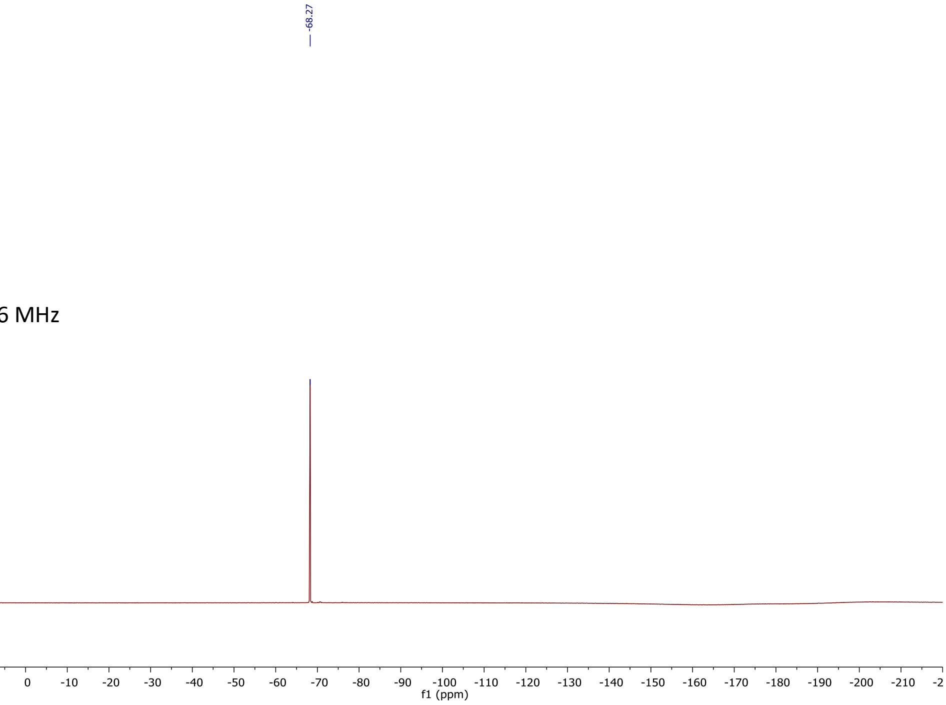

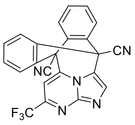

**11**
$CDCl_3$
$^{1}H$ 500 MHz

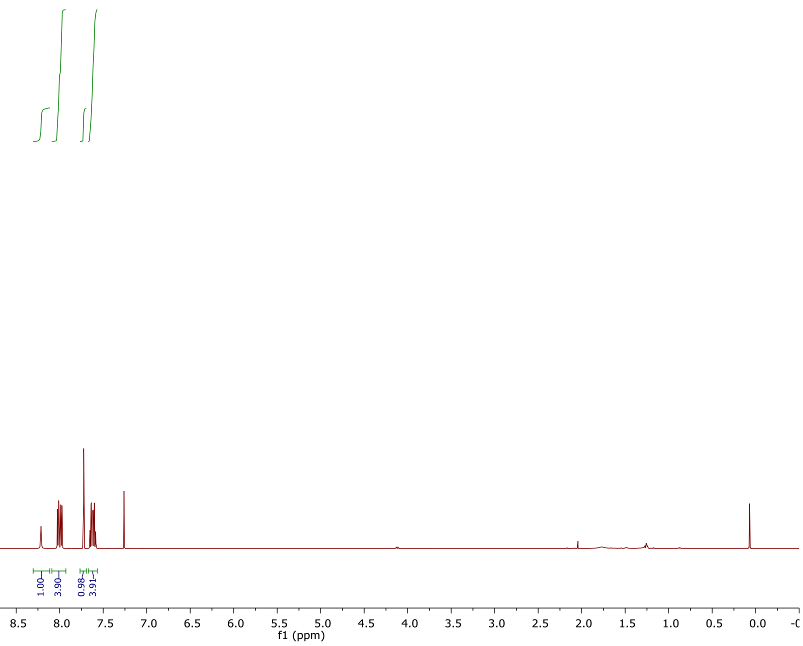

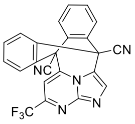

**11**
$CDCl_3$
$^{13}C$ 126 MHz

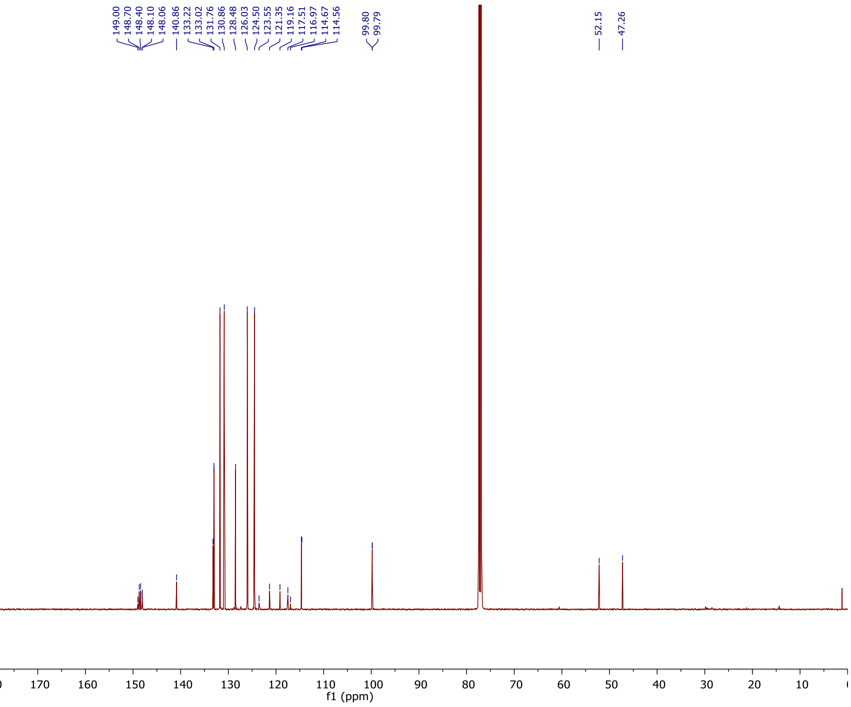

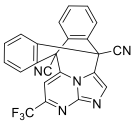

**11**
$CDCl_3$
$^{19}F$ 376 MHz

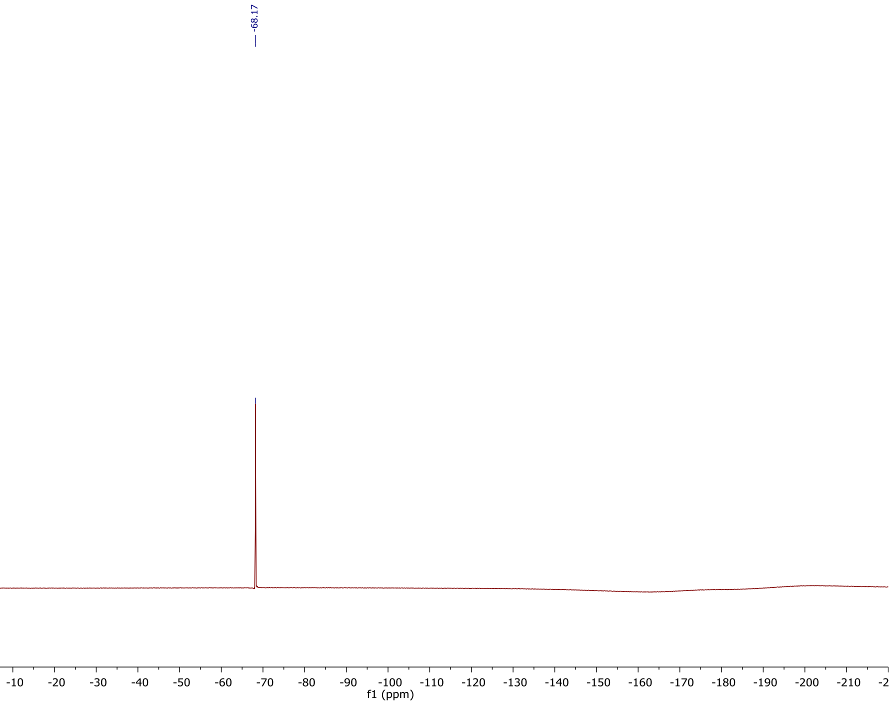

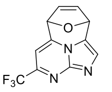

**12**
$CDCl_3$
$^{1}H$ 500 MHz

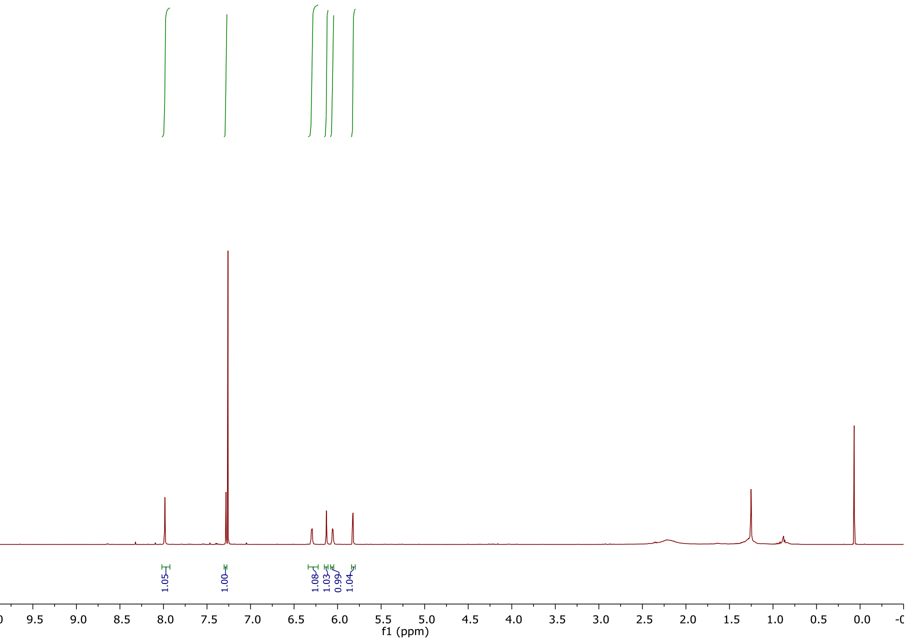

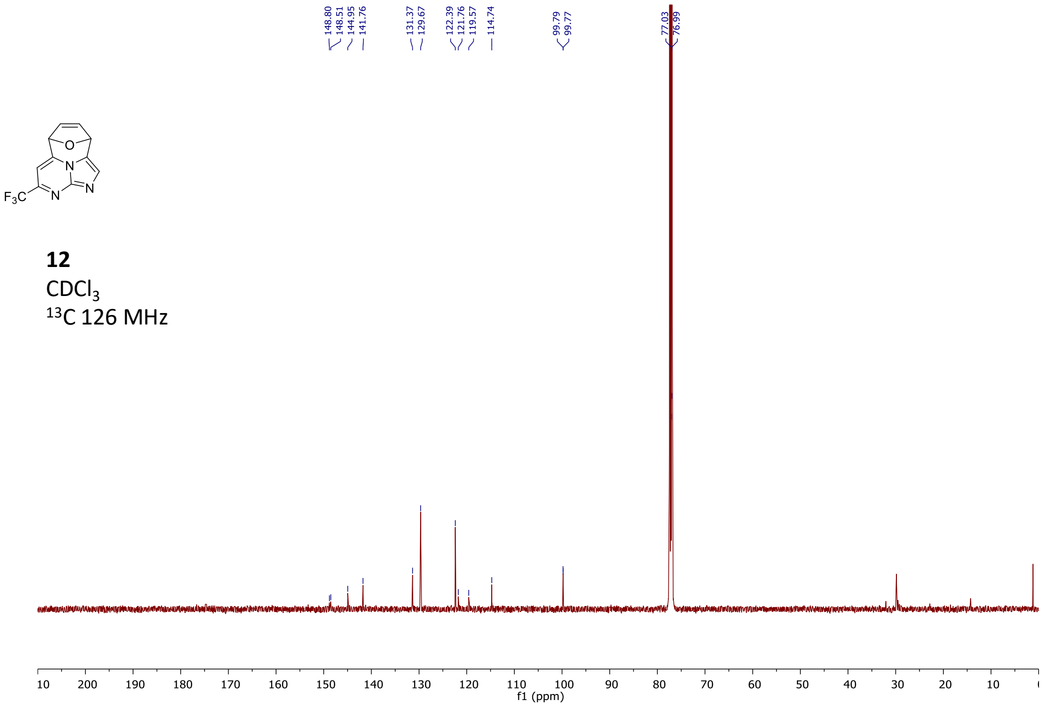

**12**
$CDCl_3$
$^{13}C$ 126 MHz

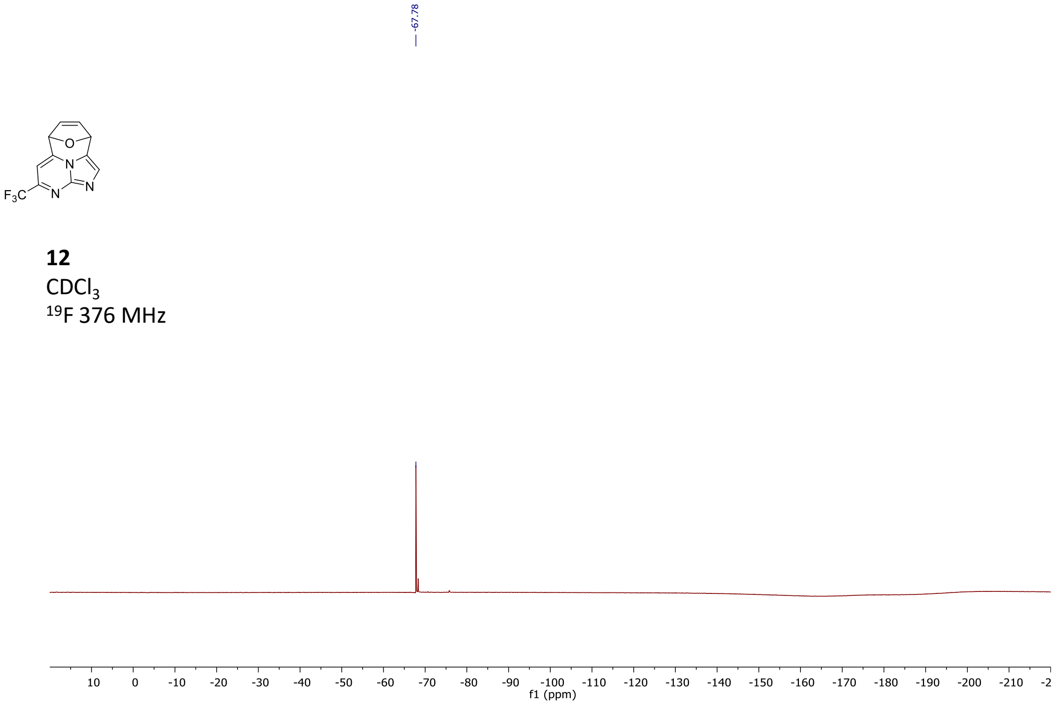

**12**
$CDCl_3$
$^{19}F$ 376 MHz

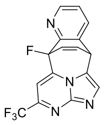

**13**
$CD_3CN$
$^{1}H$ 500 MHz

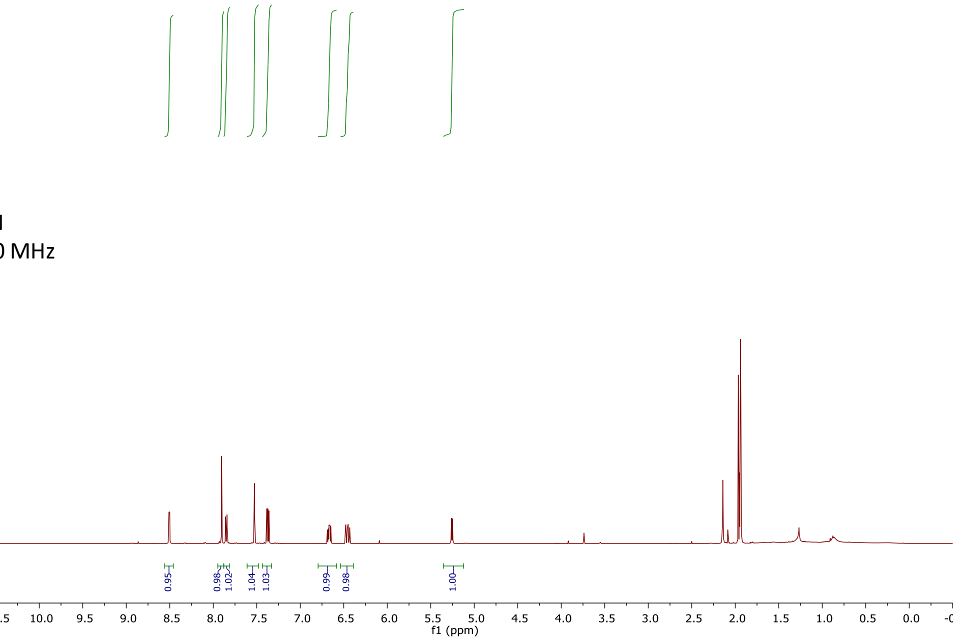

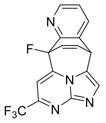

**13**
$CD_3CN$
$^{13}C$ 126 MHz

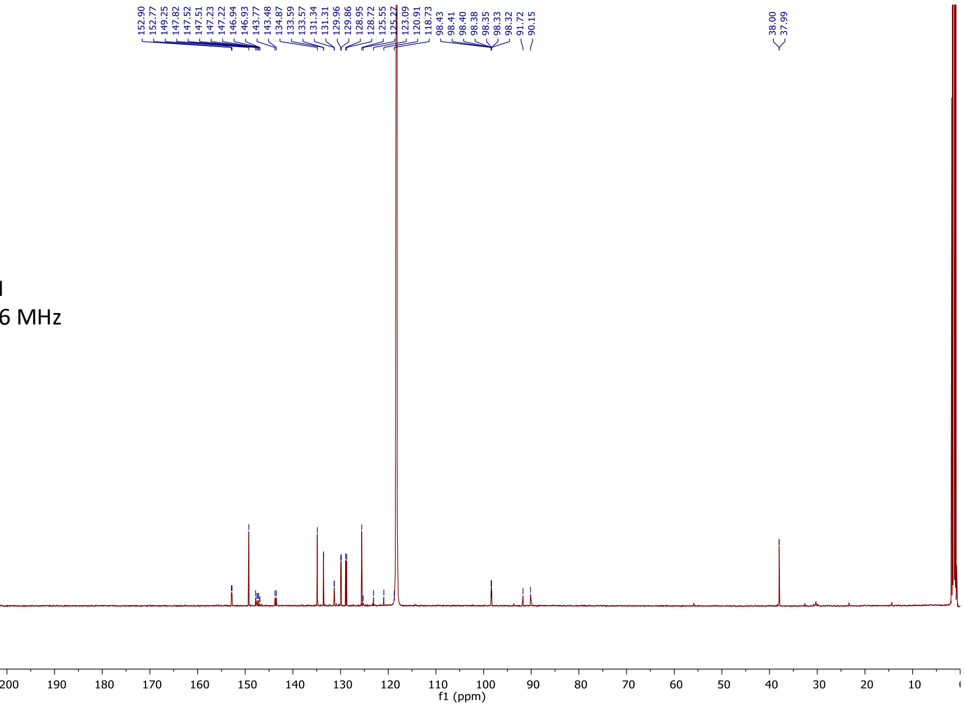

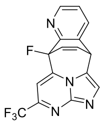

**13**
$CDCl_3$
$^{19}F$ 376 MHz

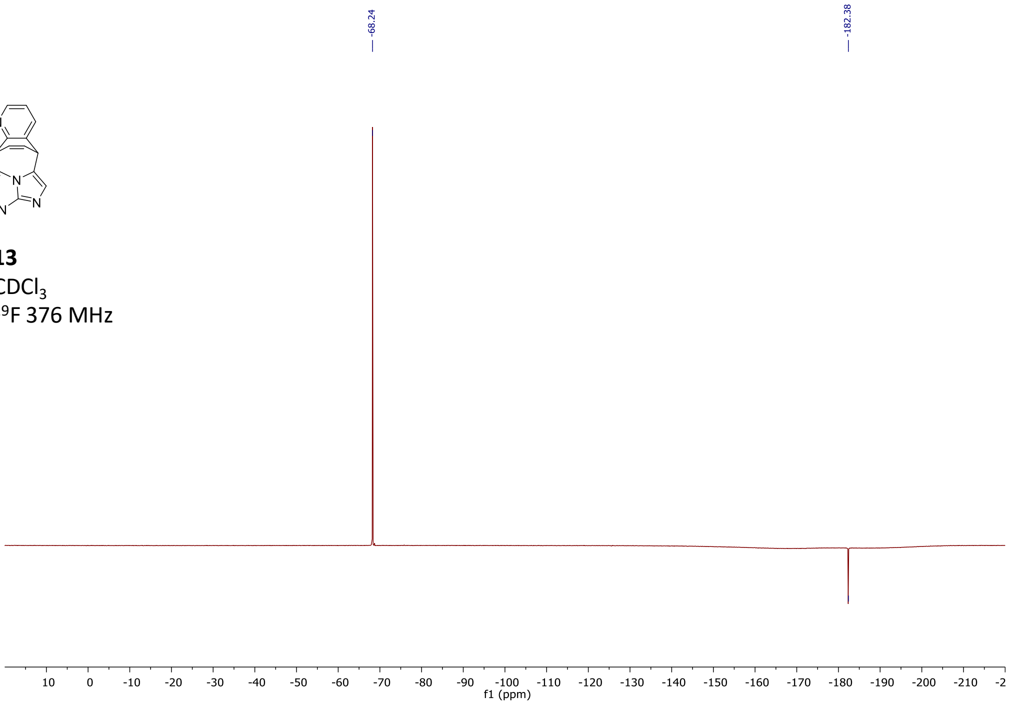

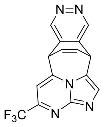

**14**
$CD_3CN$
$^{1}H$ 500 MHz

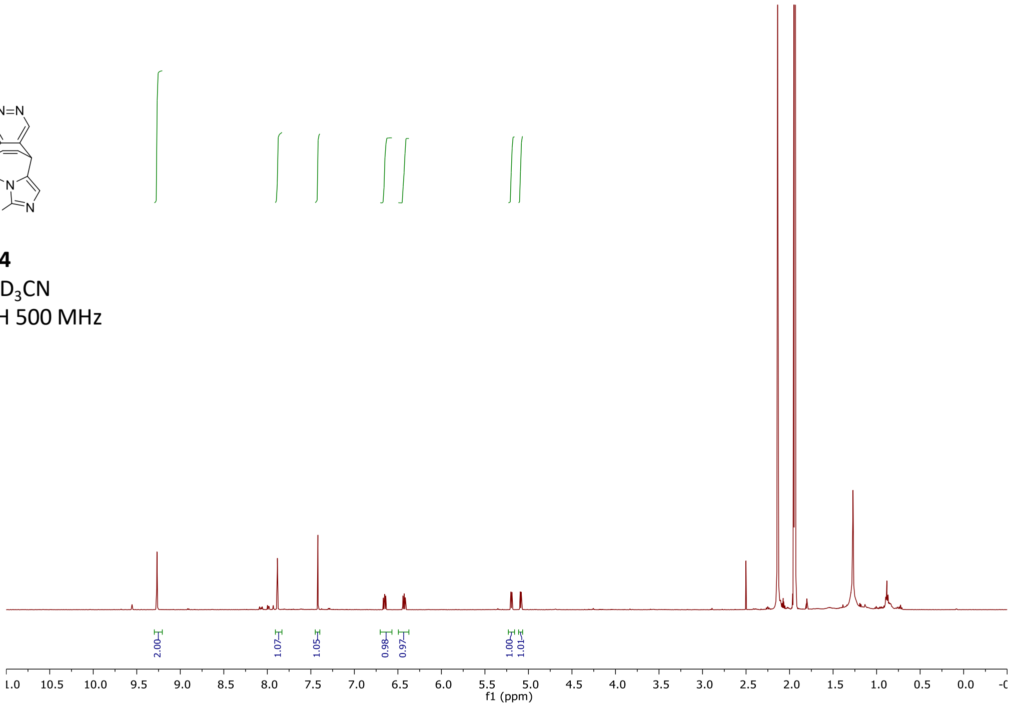

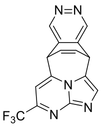

**14**
$CD_3CN$
$^{13}C$ 126 MHz

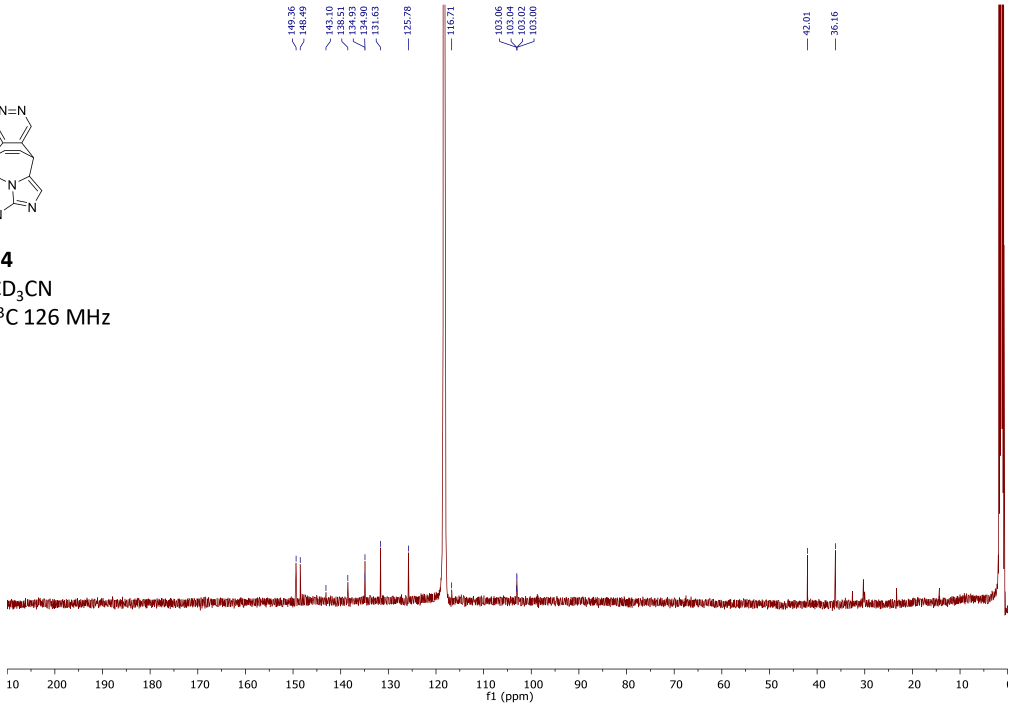

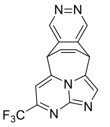

**14**
$CD_3CN$
$^{19}F$ 376 MHz

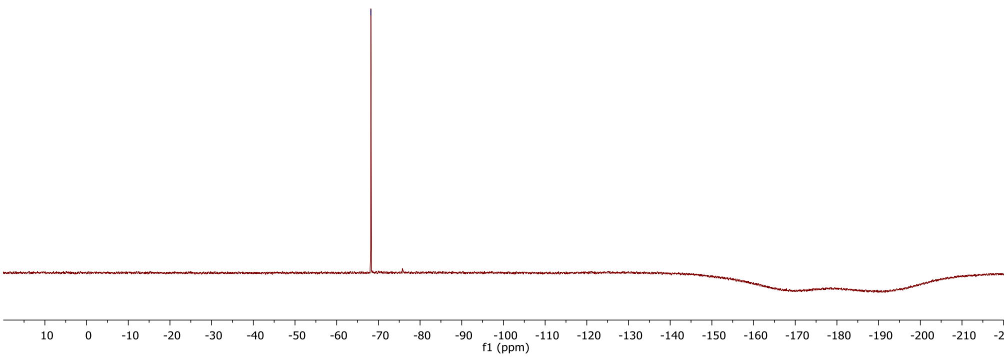

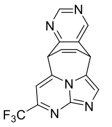

**15**
$CDCl_3$
$^{1}H$ 500 MHz

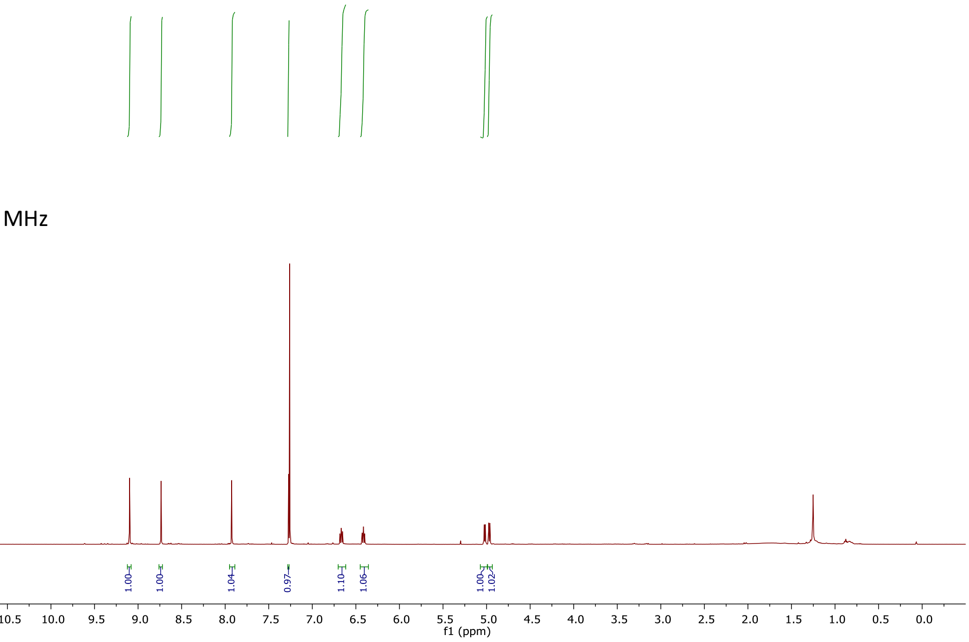

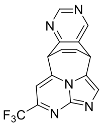

**15**
$CDCl_3$
$^{13}C$ 126 MHz

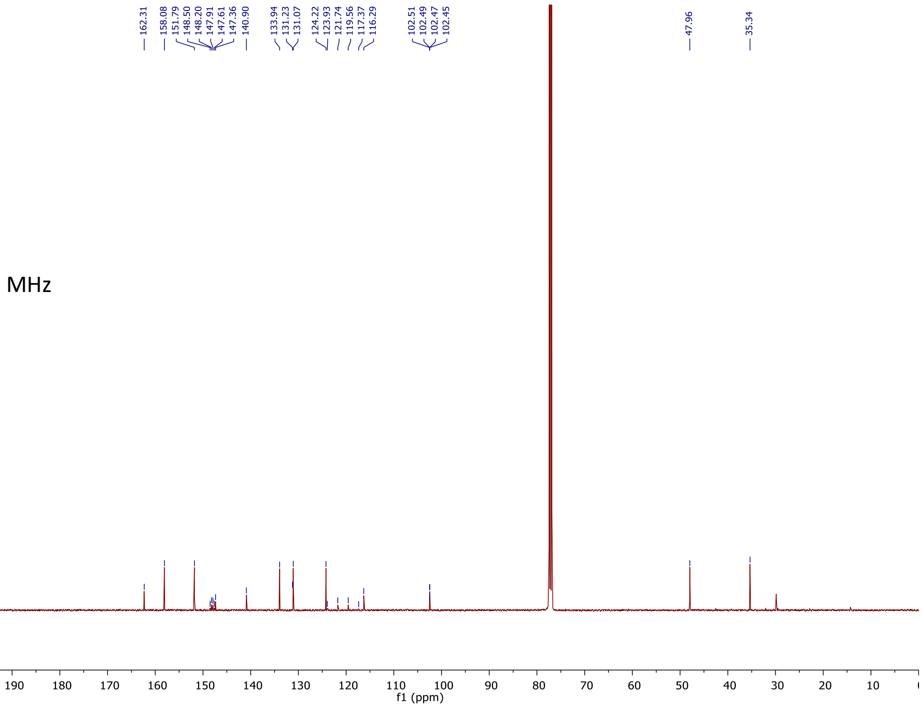

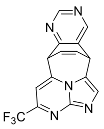

**15**
$CDCl_3$
$^{19}F$ 376 MHz

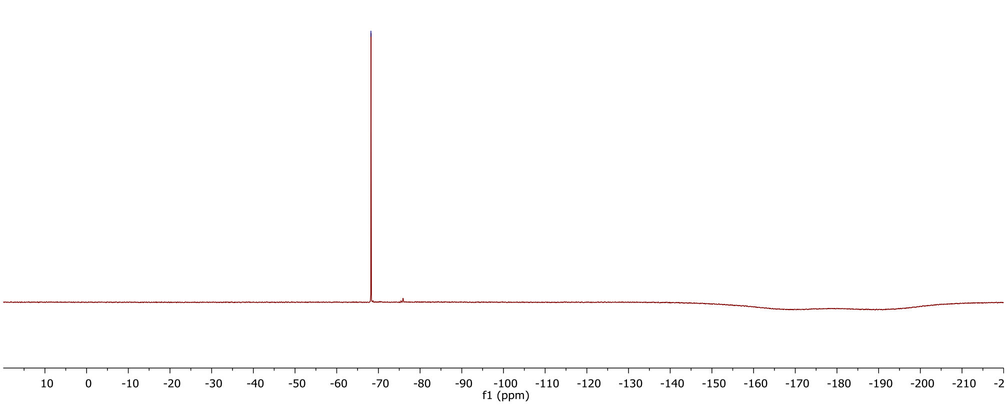

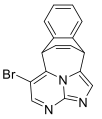

**16**
$CDCl_3$
$^{1}H$ 500 MHz

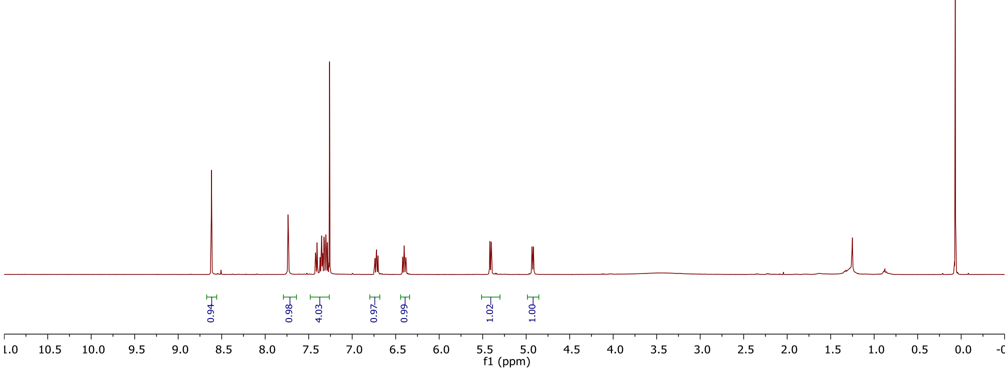

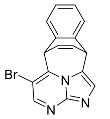

**16**
$CDCl_3$
$^{13}C$ 126 MHz

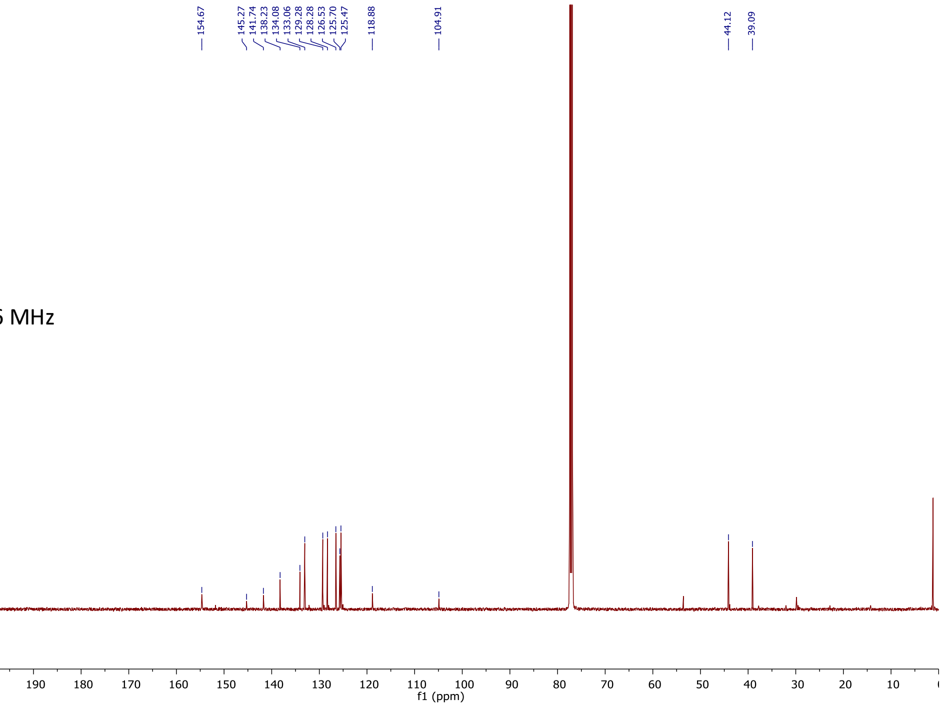

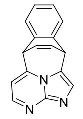

**17**
$CD_3CN$
$^{1}H$ 500 MHz

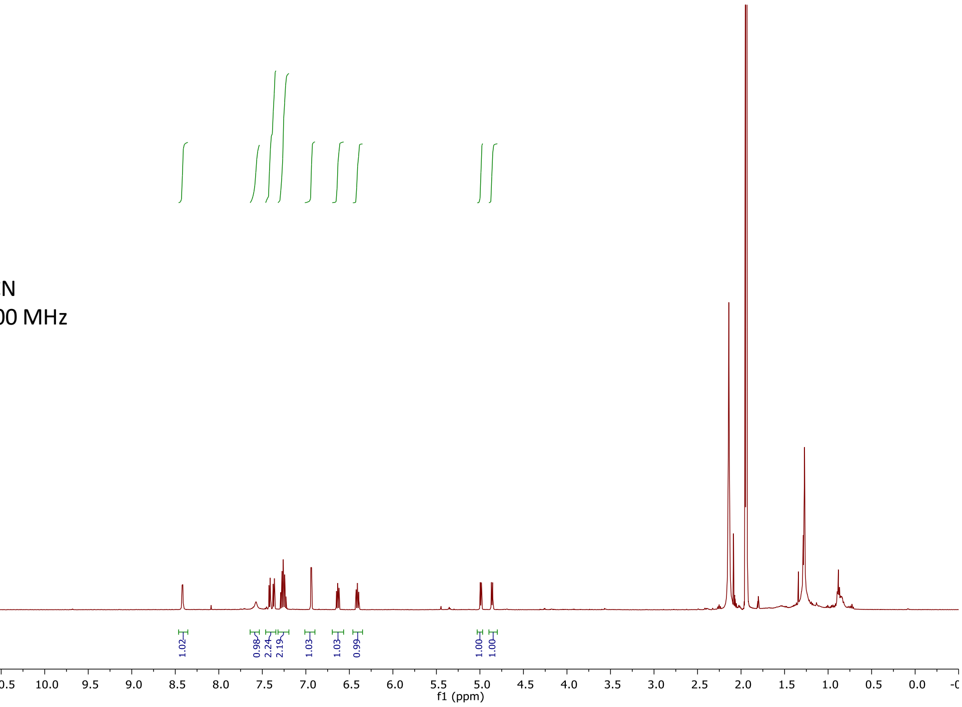

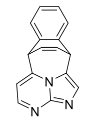

**17**
$CD_3CN$
$^{13}C$ 126 MHz

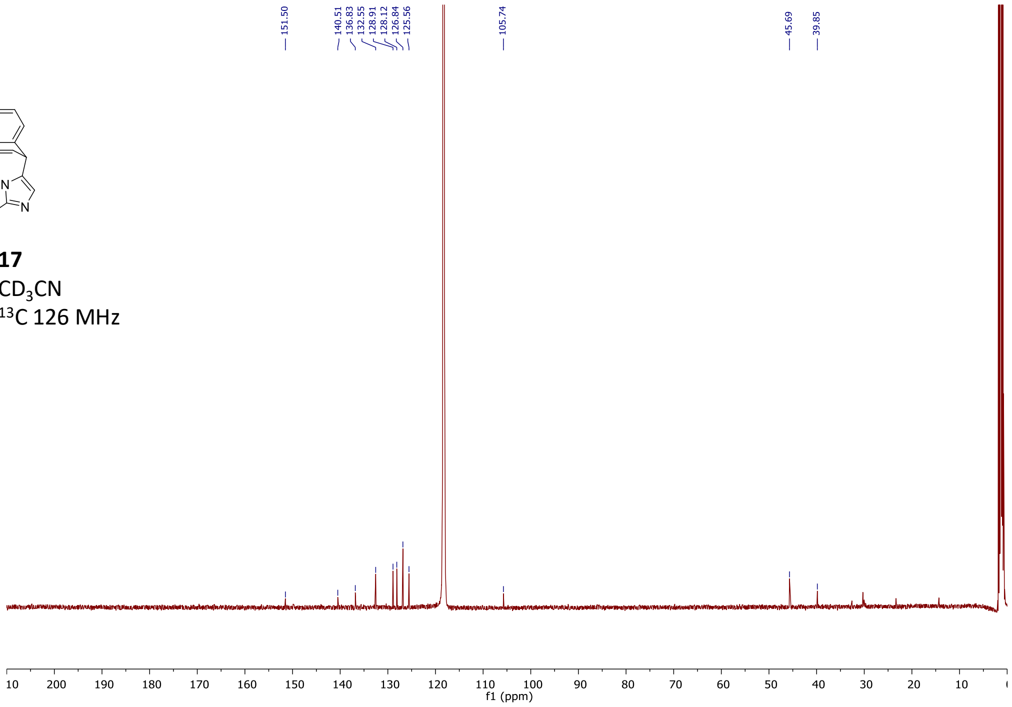

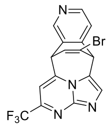

**18a**
$CD_3CN$
$^{1}H$ 500 MHz

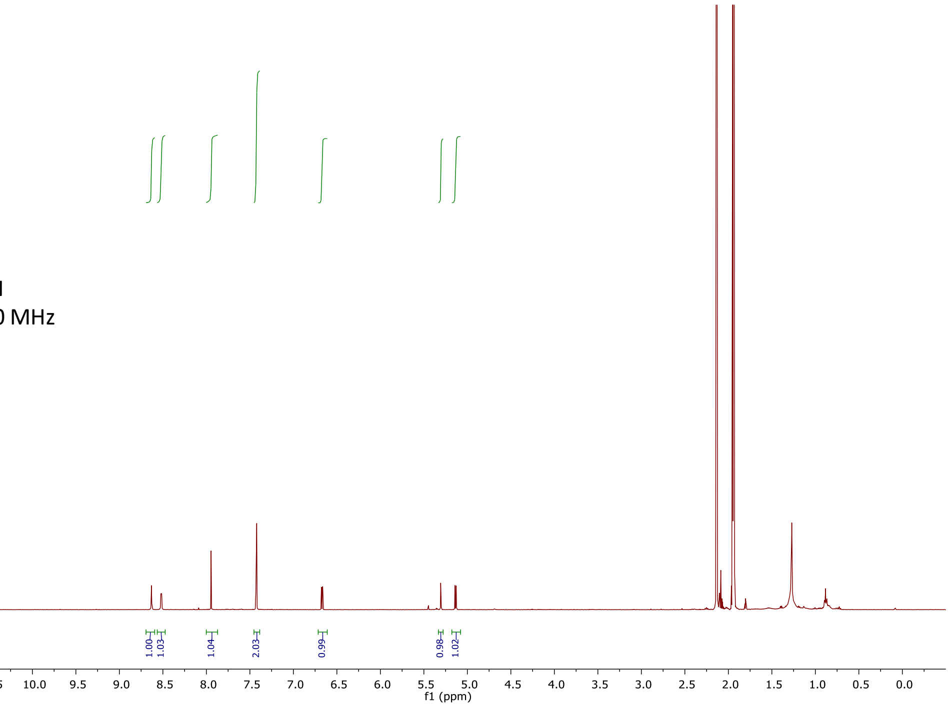

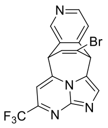

**18a**
$CD_3CN$
$^{13}C$ 126 MHz

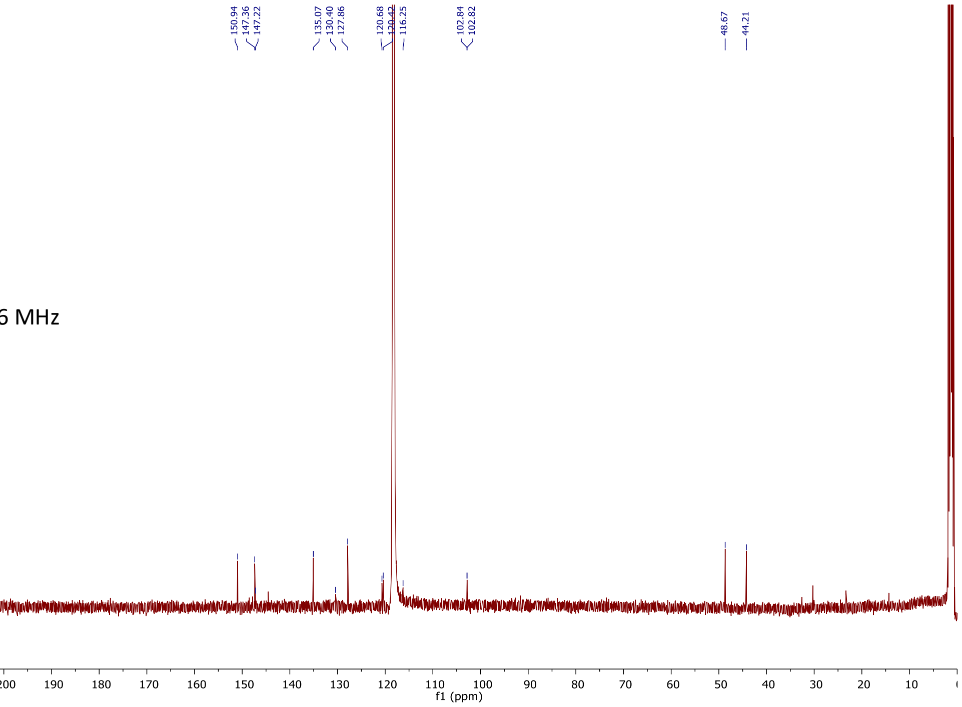

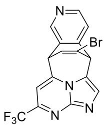

**18a**
$CD_3CN$
$^{19}F$ 376 MHz

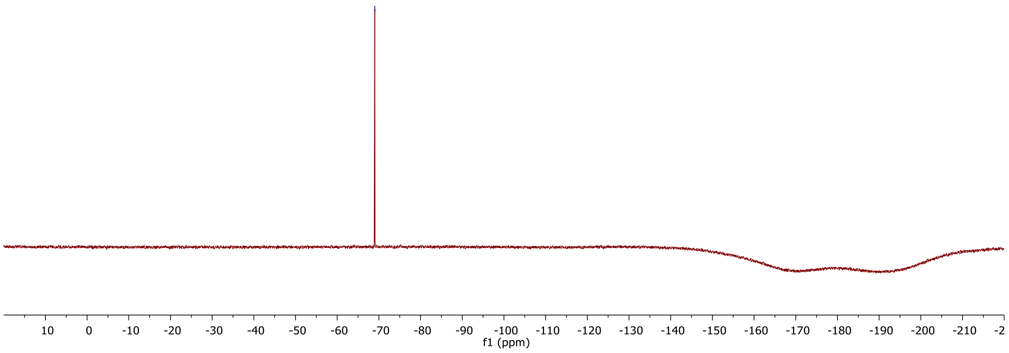

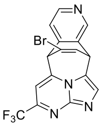

**18b**
$CD_3CN$
$^1H$ 500 MHz

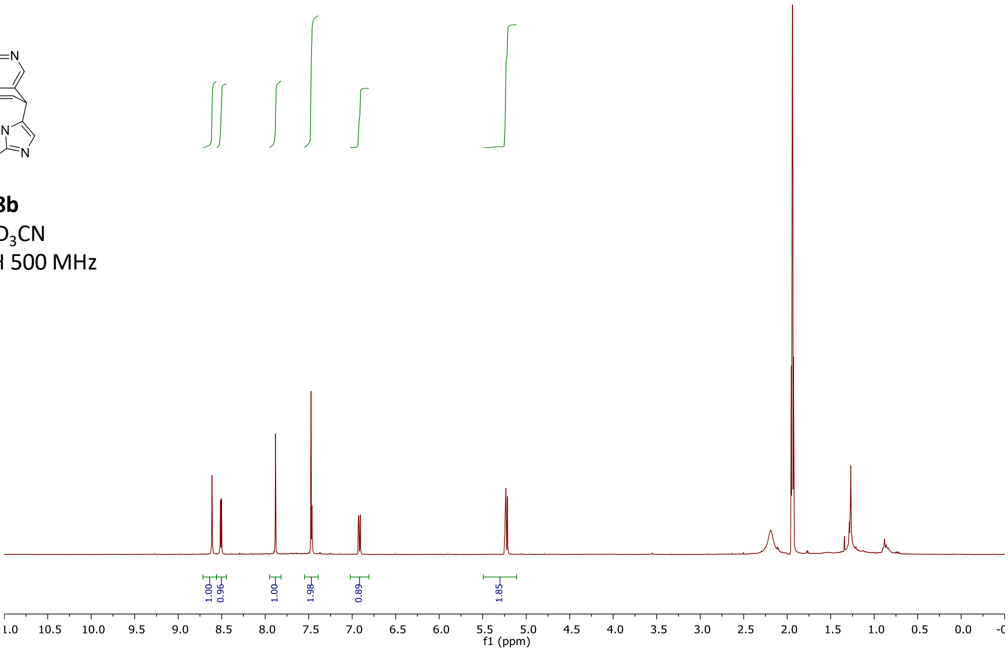

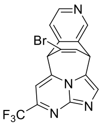

**18b**
$CDCl_3$
$^{13}C$ 126 MHz

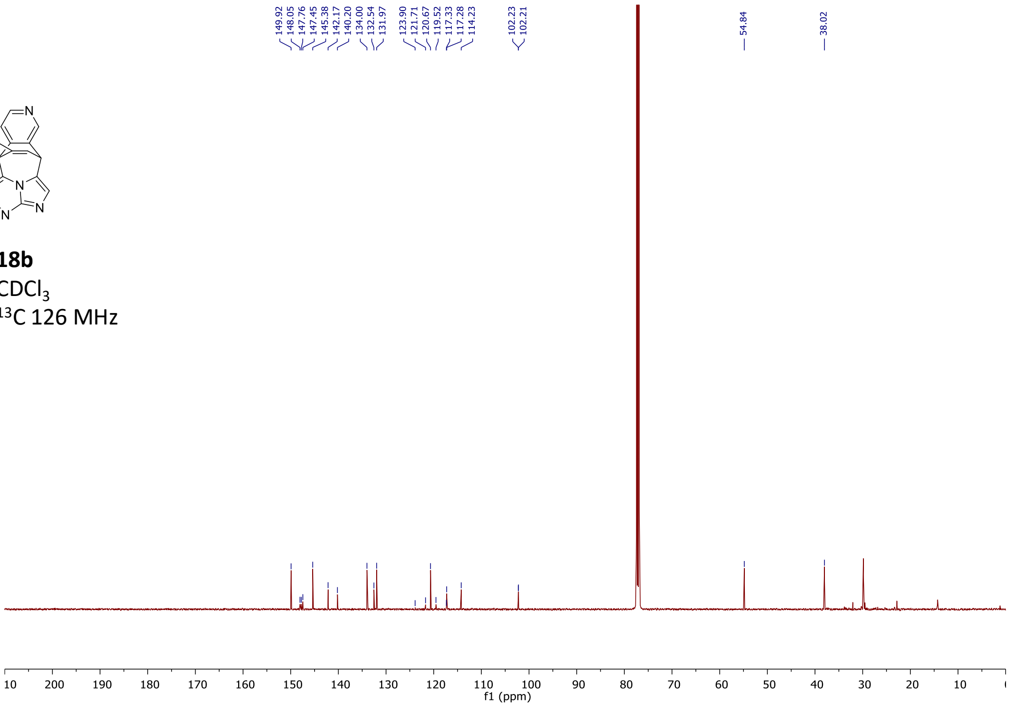

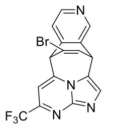

**18b**
$CD_3CN$
$^{19}F$ 376 MHz

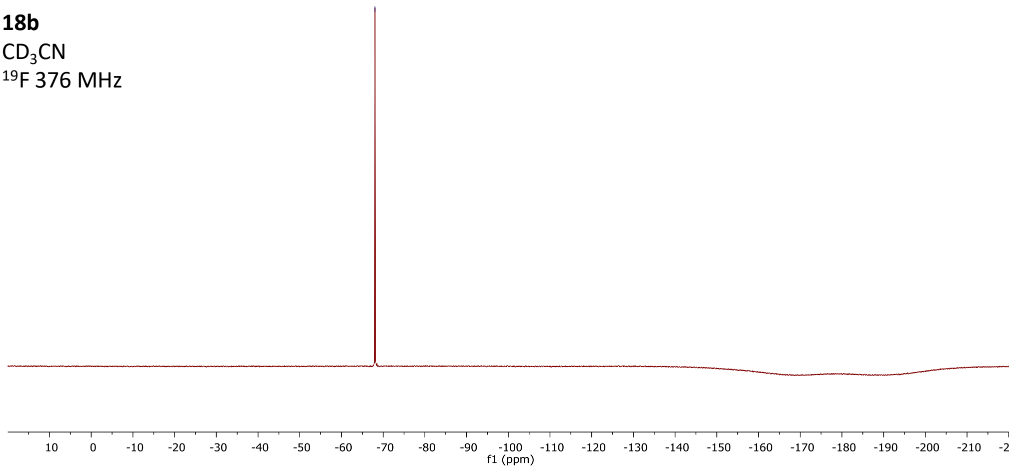

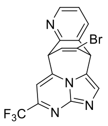

**19a**
$CDCl_3$
$^{1}H$ 400 MHz

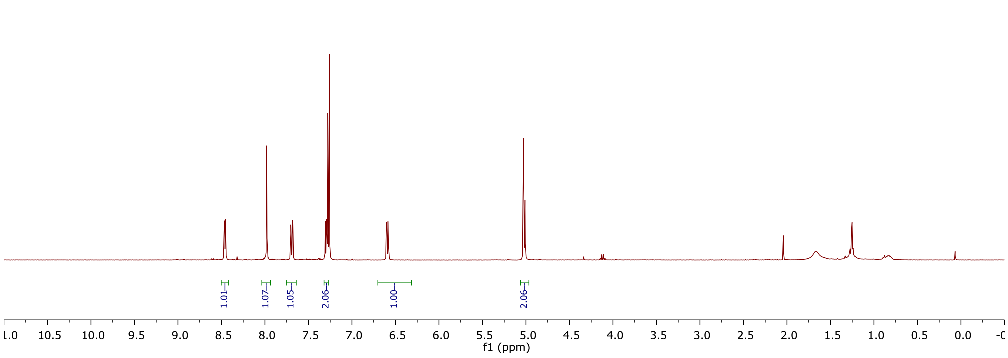

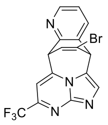

**19a**
$CDCl_3$
$^{13}C$ 126 MHz

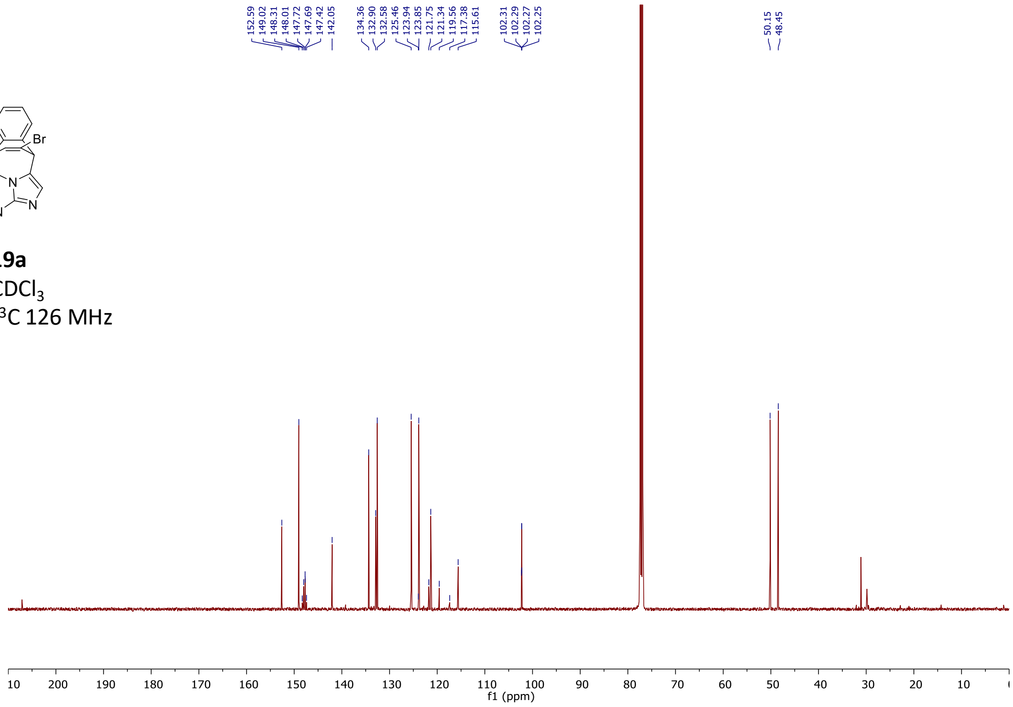

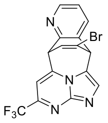

**19a**
$CDCl_3$
$^{19}F$ 376 MHz

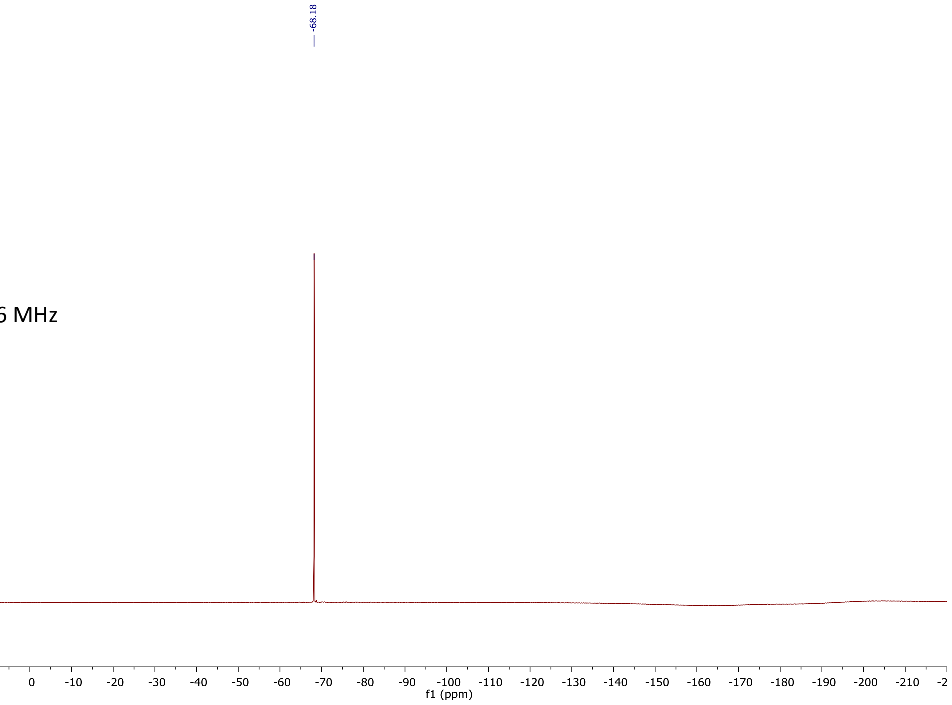

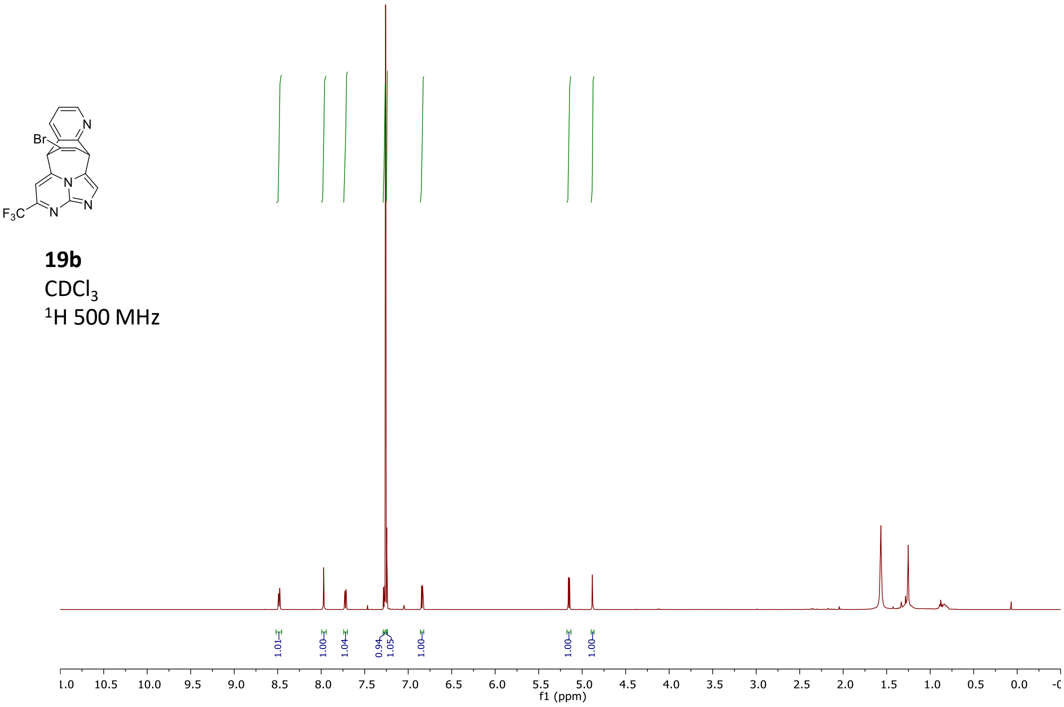
Br
N
F3C
19b
CDCl3
1H 500 MHz
1.01
1.00
1.04
0.94
1.05
1.00
1.00
1.00
f1 (ppm)

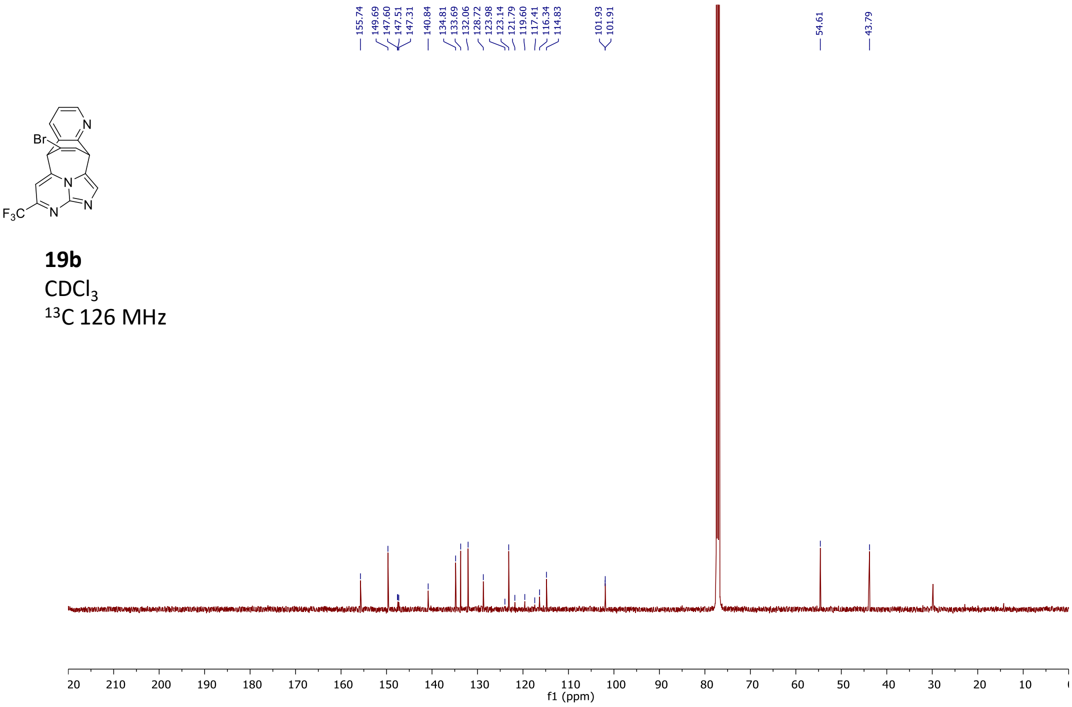
155.74
149.69
147.60
147.51
147.31
140.84
134.81
133.69
132.06
128.72
123.98
123.14
121.79
119.60
117.41
116.34
114.83
101.93
101.91
54.61
43.79
Br
N
F3C
19b
CDCl3
13C 126 MHz
f1 (ppm)

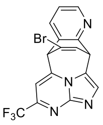

**19b**
$CDCl_3$
$^{19}F$ 376 MHz

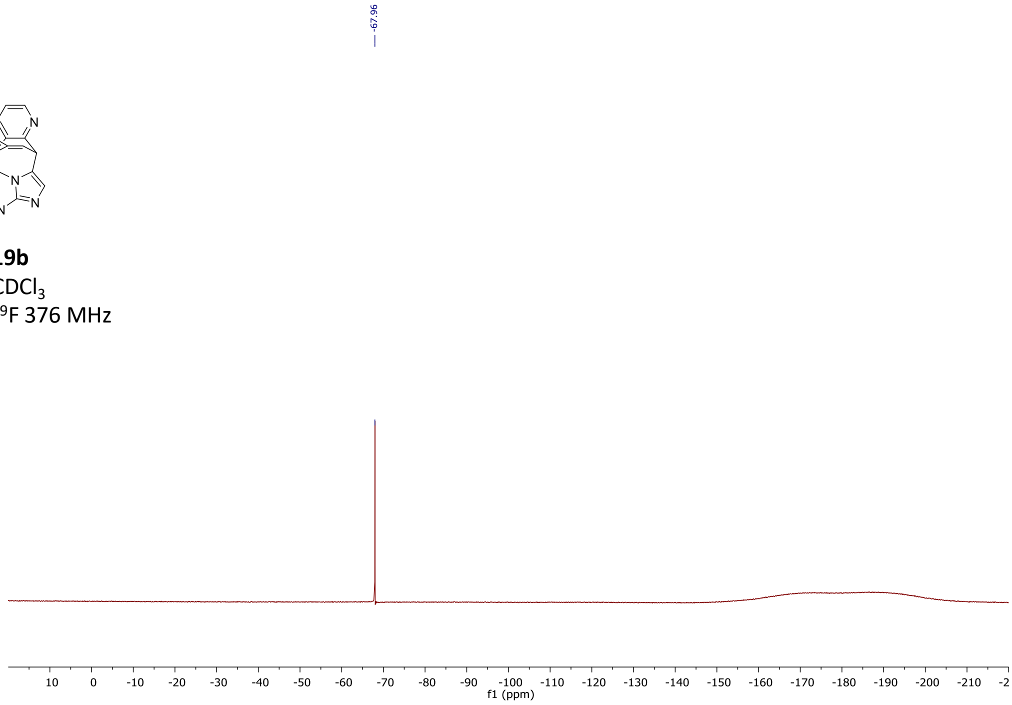

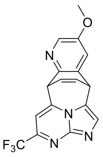

**20a**
$CD_3CN$
$^{1}H$ 500 MHz

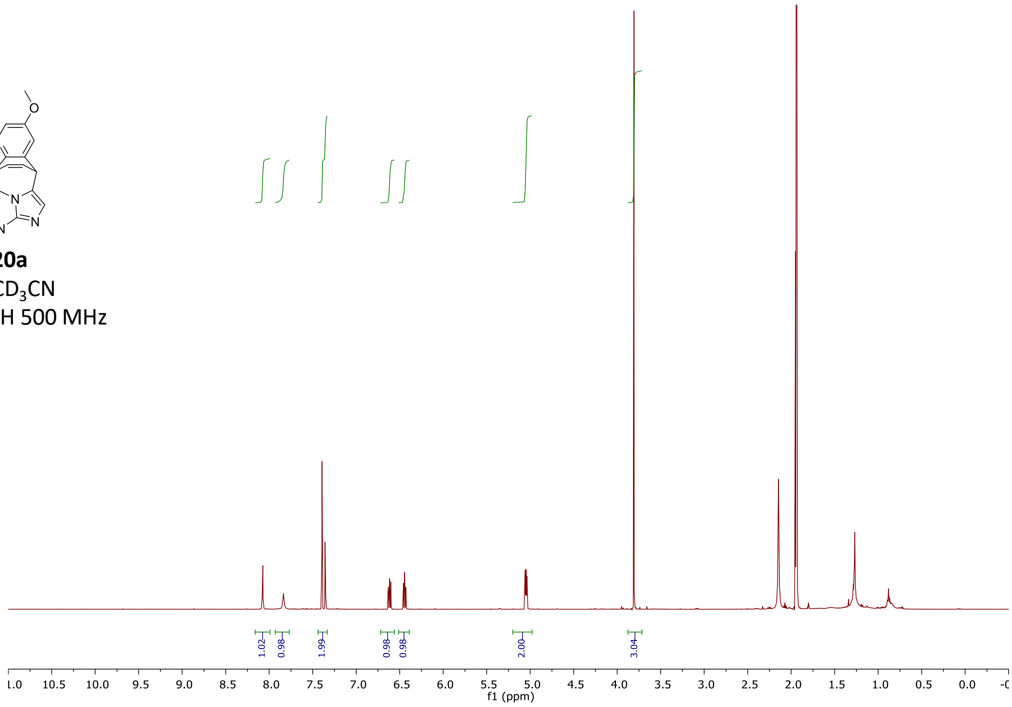

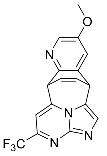

**20a**
$CD_3CN$
$^{13}C$ 126 MHz

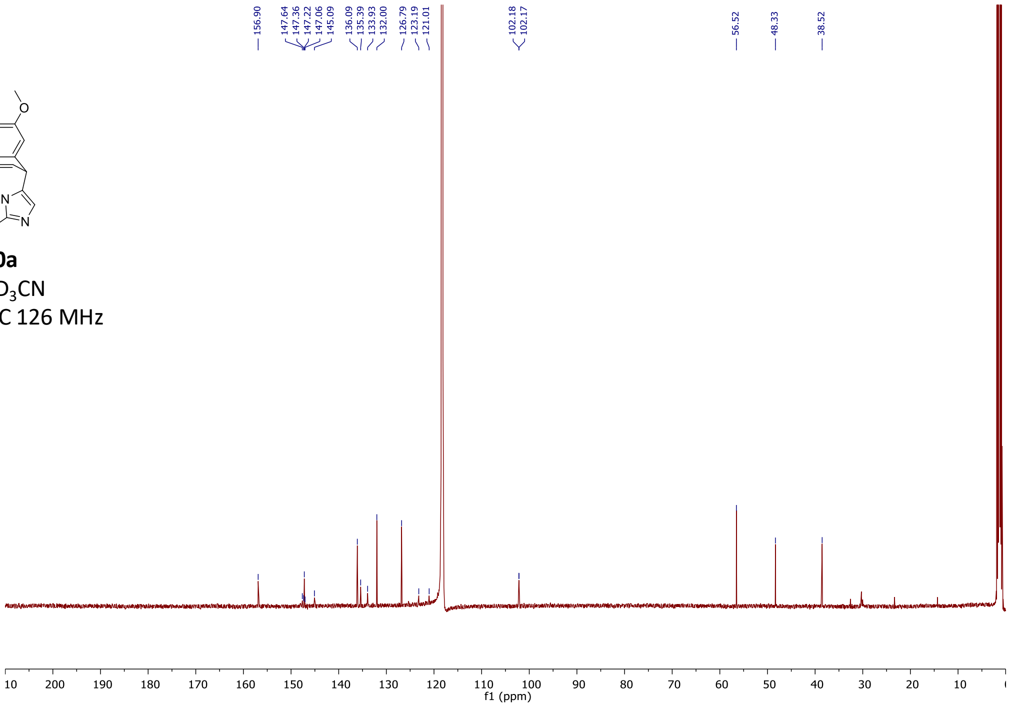

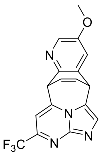

**20a**
$CD_3CN$
$^{19}F$ 376 MHz

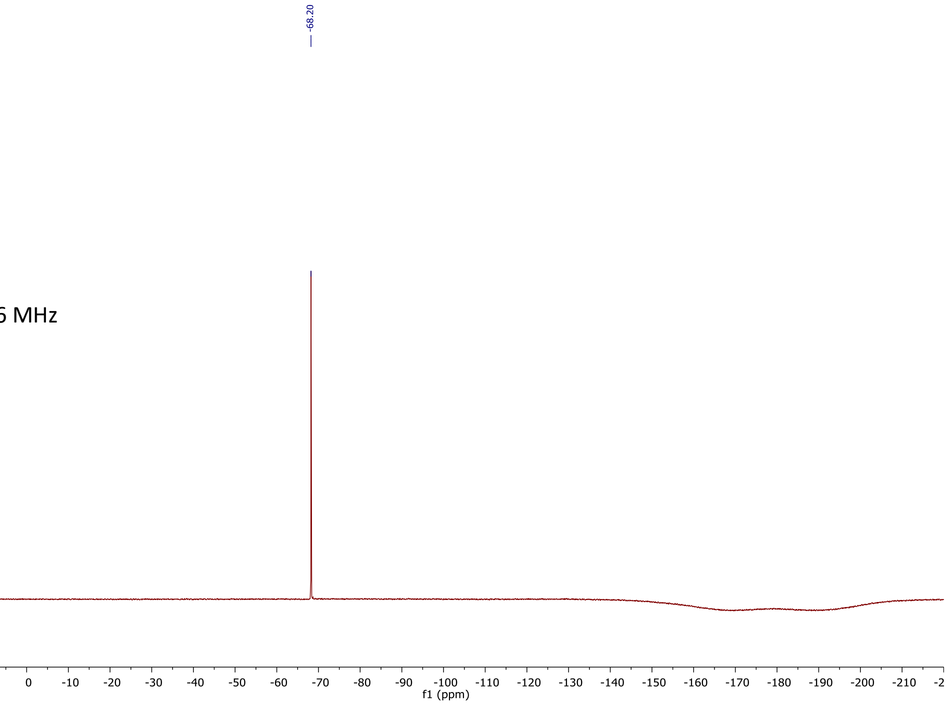

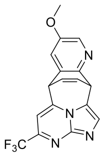

**20b**
$CDCl_3$
$^{1}H$ 500 MHz

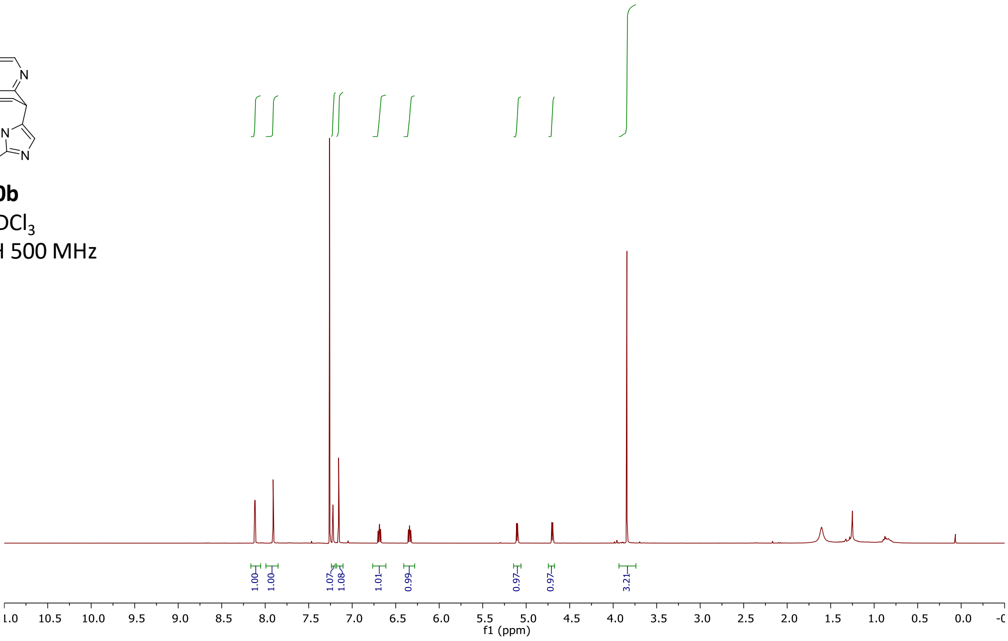

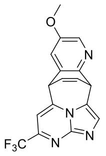

**20b**
$CDCl_3$
$^{13}C$ 126 MHz

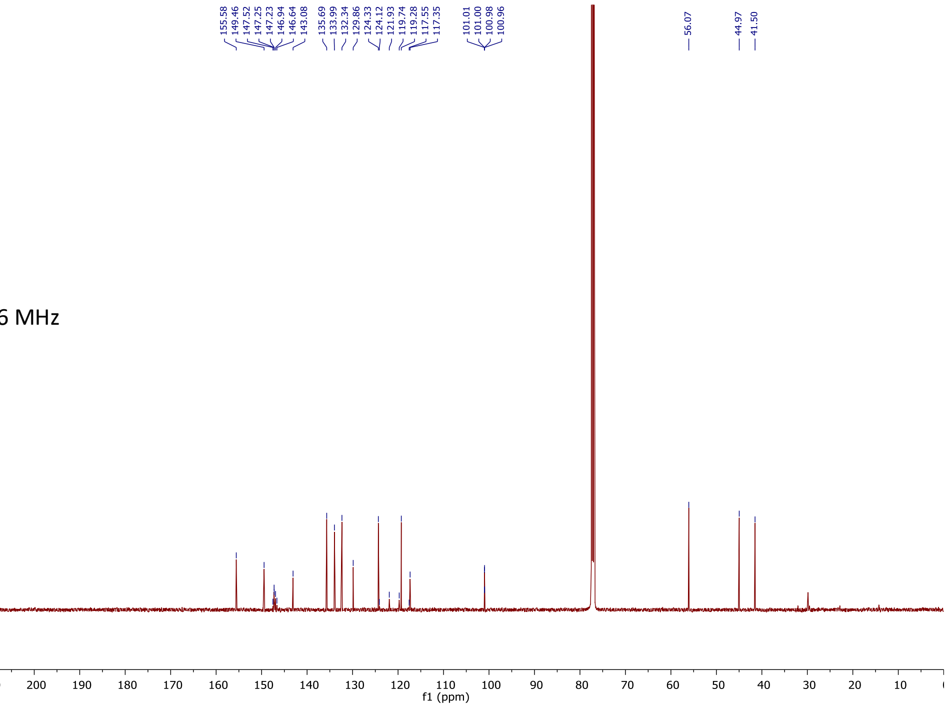

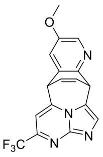

**20b**
$CDCl_3$
$^{19}F$ 376 MHz

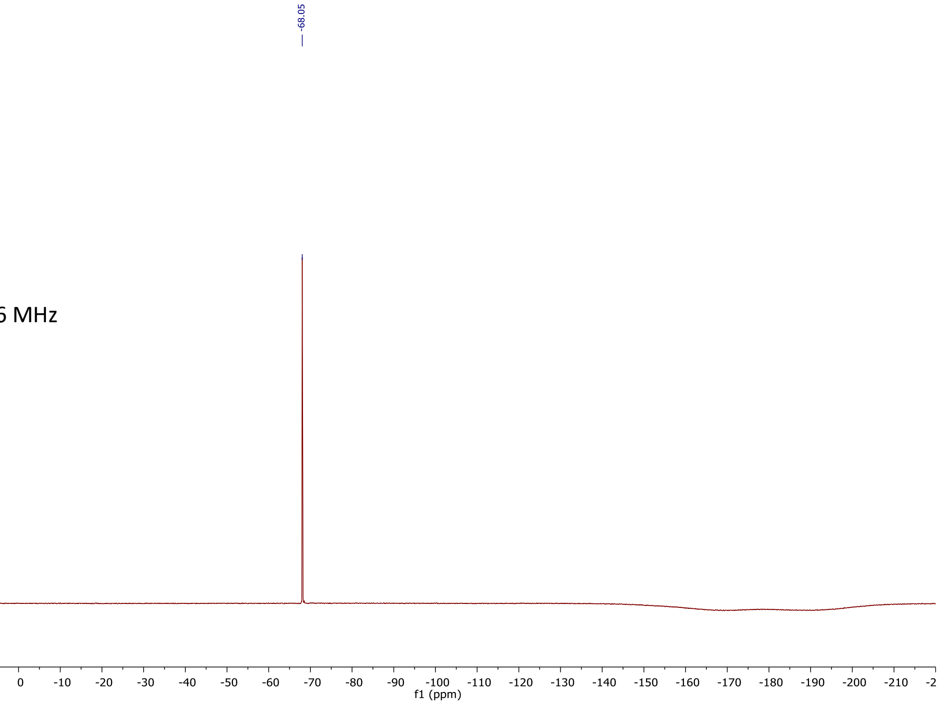

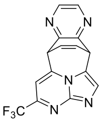

**21**
$CD_3CN$
$^{1}H$ 500 MHz

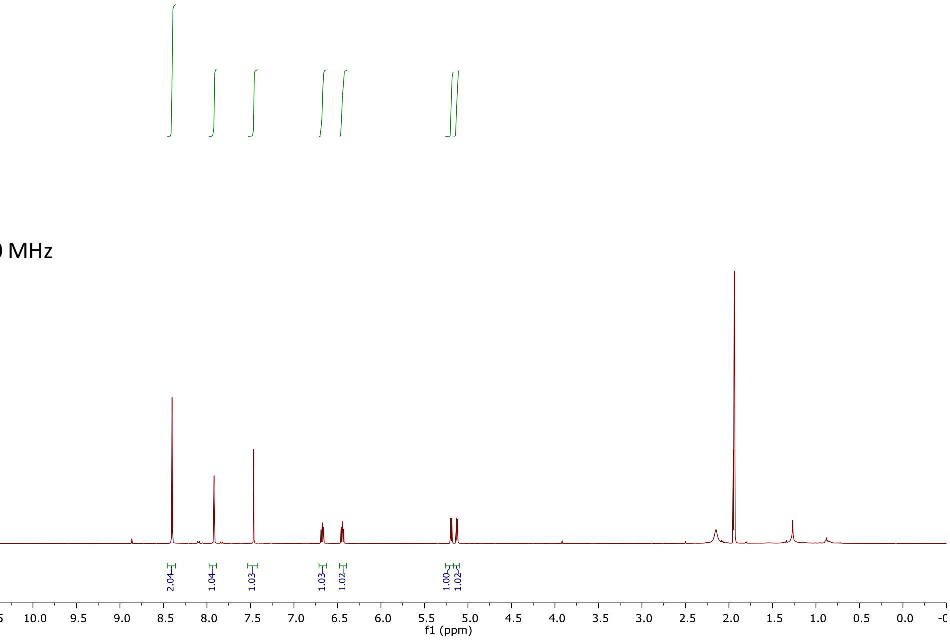

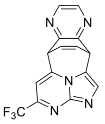

**21**
$CD_3CN$
$^{13}C$ 126 MHz

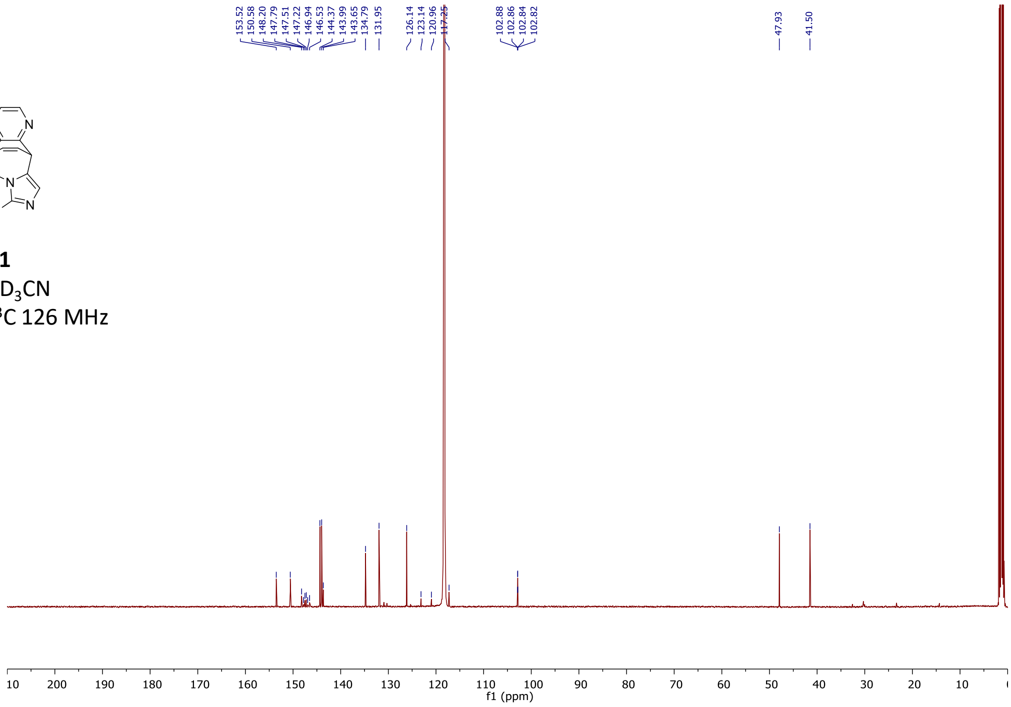

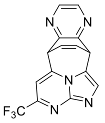

**21**
$CD_3CN$
$^{19}F$ 376 MHz

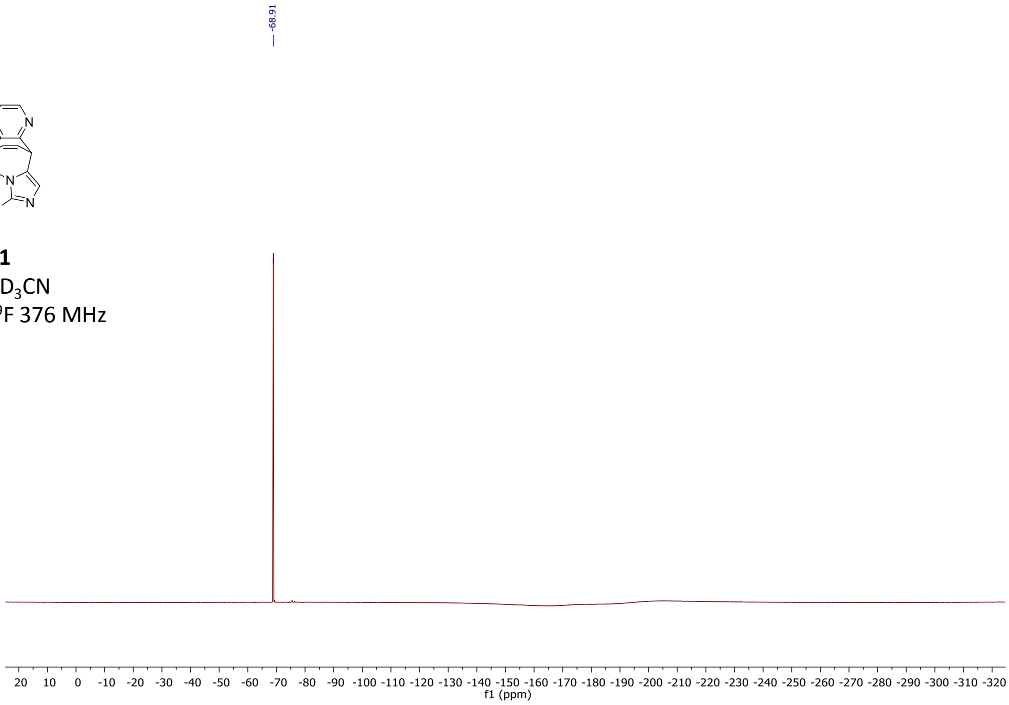

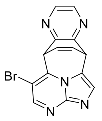

**22**
$CD_3CN$
$^{1}H$ 500 MHz

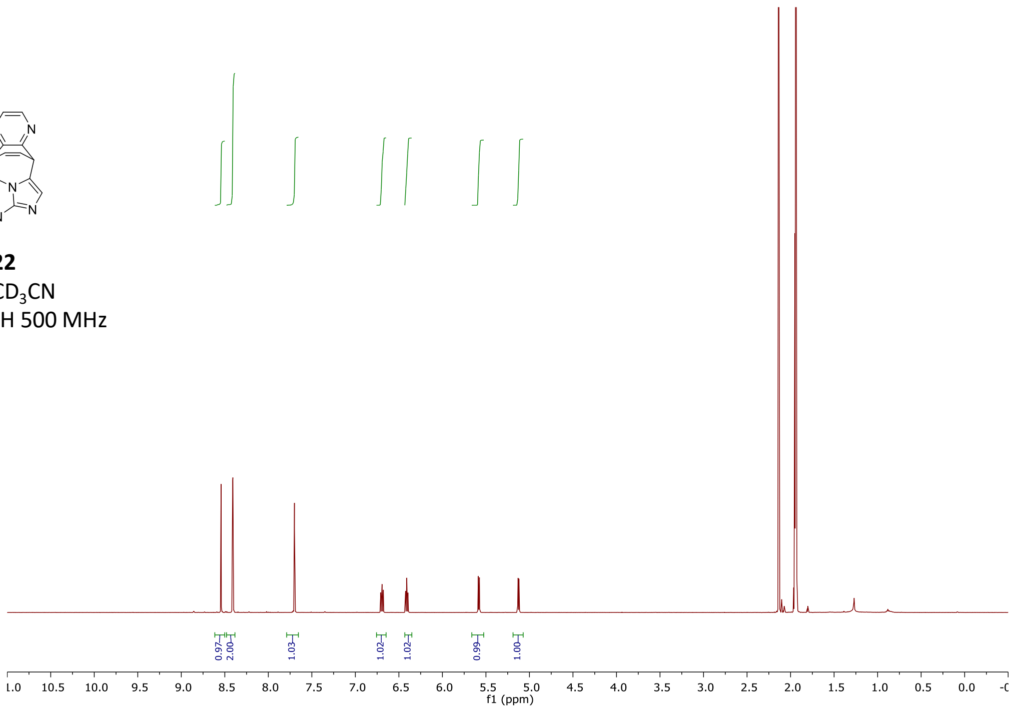

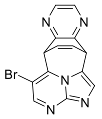

**22**
$CD_3CN$
$^{13}C$ 126 MHz

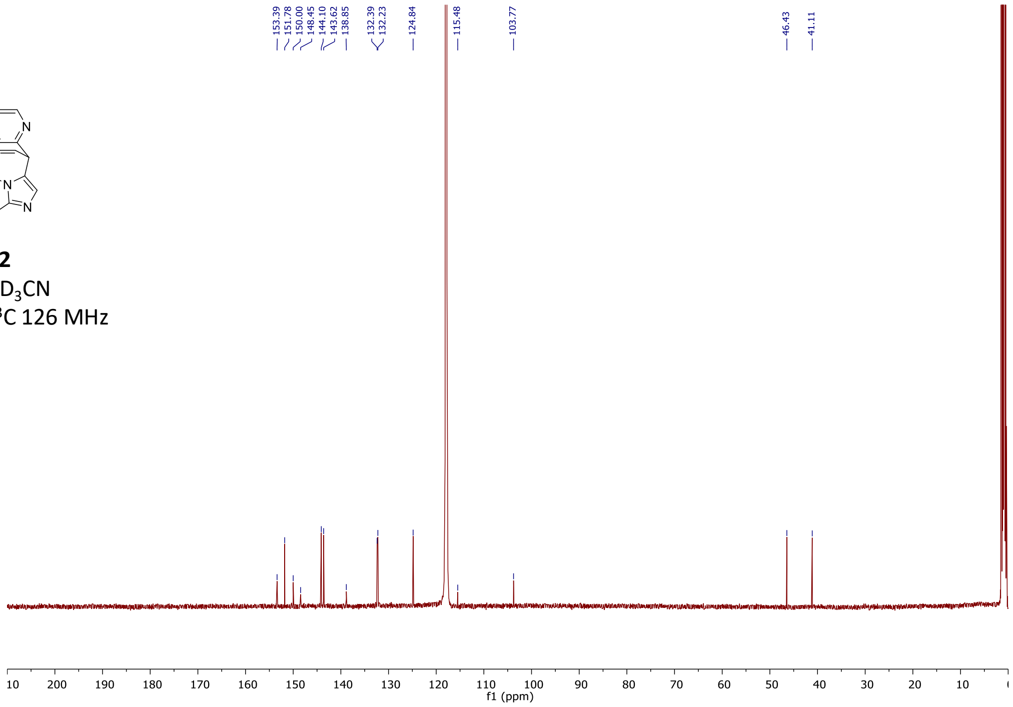

**General procedures for [4+3] cycloaddition on proteins**

***General procedure C for the [4+3] cycloaddition reaction on eH3-Npa9:*** The reaction mixture was prepared open to atmosphere in 1.2 mL glass vials. To 29.7 µL eH3-Npa9 (50 µM) in denaturing buffer (500 mM ammonium acetate, 3 M guanidine-HCl, pH 6.2) were added 0.3 µL of a 500 mM stock solution of small molecule in DMSO (5 mM, 100 equiv., 1% (*v/v*) residual DMSO) resulting in a final volume of 30 µL. The vial was transferred into a DeSyre photoreaction block, containing holes in the bottom plate to allow for light transmittance. The block was placed in the Zinsser Lumidox® photoreactor and the mixture was irradiated (365 nm, 65 mW/well) for 1 h while stirring (400 rpm) and cooling of the reaction block holder (4 °C).

Note: This protocol resulted in good conversion for 7-Trifluoromethyl imidazo[1,2-*a*]pyrimidine to the observed on-protein product, however, the MS spectra indicated significant side chain oxidations.

***General procedure D for the [4+3] cycloaddition reaction on eH3-Npa9 under exclusion of $O_2$:*** The optimized general procedure **D** involved preparing the reaction mixtures strictly under inert atmosphere in a glove box. For this, all vessels and stock solutions were left open to the nitrogen atmosphere of the glove box overnight before conducting the experiment. The protein stock solution was kept at 4 °C at all time.

To 99 µL eH3-Npa9 (50 µM) in denaturing buffer (500 mM ammonium acetate, 3 M guanidine-HCl, pH 6) in 1.2 mL glass vials was added 1 µL of a 500 mM small molecule stock solution in DMSO (5 mM, 100 equiv., 1% (v/v) residual DMSO) resulting in a final volume of 100 µL per reaction. Four technical replicates were prepared. The vials were capped and placed into a 96-well Desyre® photoreaction block with holes in the bottom plate to allow for light transmittance. Subsequently, the lid was applied with sufficient pressure to ensure air-tight sealing of the capped glass vials. The block was taken out of the glove box and put into the Zinsser Lumidox® photoreactor. The reaction mixtures were irradiated for minimum of 15 minutes with blue light (365 nm, 220 mW/well) while agitating at 400 rpm and cooling of the block holder to 4 °C. The block was transferred back into the glove box where an analytical sample was prepared to confirm the conversion to the desired product with protein-LC-MS. For this, 1 µL of crude reaction mixture was diluted with 49 µL of $H_2O$ containing 0.1% formic acid and analyzed utilizing the Waters XEVO-QTof LC-MS. Subsequently, the vials were resealed and irradiated for another 15 min in the photoreactor.

For MS/MS experiments and/or photophysical measurements the technical replicates with confirmed conversion to the product were combined and purified (desalted) using SEC (PD Minitrap G25, gravity protocol) with the respective reaction buffer as elution buffer. The final concentration of purified product was determined using a nanodrop-spectrophotometer.

***General procedure E for the [4+3] Cycloaddition on AnxV-Npa187 without Ca-salt:*** Reactions on Annexin V were strictly run under inert atmosphere in a glove box after all vessels and solutions had been gently flushed with nitrogen overnight. For each reaction 15 µM of protein (final reaction concentration) in the respective buffer solution in 1.2 mL glass vials was mixed with 1.5 mM of small molecule (100 equiv.) out of a DMSO stock solution (2% (*v/v*) residual DMSO). The vials were capped and placed into a 96-well Desyre® photoreaction block with holes in the bottom plate to allow for light transmittance. Subsequently, the lid was applied with sufficient pressure to ensure air-tight sealing of the capped glass vials. The block was taken out of the glove box and put into the Zinsser Lumidox® photoreactor. The reaction mixtures were irradiated for a minimum of 30 minutes with blue light (365 nm, 280-305 mW/well) while agitating at 400 rpm with cooling of the reaction block holder (4 °C). An analytical sample was prepared to confirm the conversion to the product with protein-LC-MS. For this, 2 µL of crude reaction mixture were diluted with 48 µL of $H_2O$ containing 0.1% formic acid and analyzed utilizing the Waters XEVO-QTof LC-MS.

Note: The procedure **E** was followed with all three imidazo[1,2-*a*]pyrimidine analogs (7-$CF_3$, 6-Br, and unsubstituted). None of these reactions resulted in desired product formation.

***General procedure F for the [4+3] Cycloaddition on AnxV-Npa187 with 40 µM Ca-salt:*** Reactions on Annexin V were strictly run under inert atmosphere in a glove box after all vessels and solutions had been gently flushed with nitrogen overnight. For each reaction 15 µM of protein (final reaction concentration) in the respective buffer solution in 1.2 mL glass vials was mixed with 40 µM of aqueous $CaCl_2$ solution (2.67 equiv.) and 1.5 mM of small molecule (100 equiv.) out of a DMSO stock solution (2% (*v/v*) residual DMSO). The vials were capped and placed into a 96-well Desyre® photoreaction block with holes in the bottom plate to allow for light transmittance. Subsequently, the lid was applied with sufficient pressure to ensure air-tight sealing of the capped glass vials. The block was taken out of the glove box and put into the Zinsser Lumidox® photoreactor. The reaction mixtures were irradiated for a minimum of 15 minutes with blue light (365 nm, 280-305 mW/well) while agitating at 400 rpm with cooling of the reaction block holder (4 °C). An analytical sample was prepared to confirm the conversion to the product with protein-LC-MS. For this, 2 µL of crude reaction mixture were diluted with 48 µL of $H_2O$ containing 0.1% formic acid and analyzed utilizing the Waters XEVO-QTof LC-MS.

Note: The procedure **F** was followed with all three imidazo[1,2-*a*]pyrimidine analogs (7-$CF_3$, 6-Br, and unsubstituted), as well as without small molecule as a control reaction. Only the reaction with the unsubstituted imidazo[1,2-*a*]pyrimidine resulted in traces of product, merely detected by protein LC-MS.

***General procedure G for the [4+3] Cycloaddition on AnxV-Npa187 with 30 mM Ca-salt:*** Reactions on Annexin V were strictly run under inert atmosphere in a glove box after all vessels and solutions had been gently flushed with nitrogen overnight. For each reaction 15 µM of protein (final reaction concentration) in Tris buffer solution in 1.2 mL glass vials was mixed with 30 mM of aqueous $CaCl_2$ solution (2000 equiv.) and 1.5 mM of small molecule (100 equiv.) out of a DMSO stock solution (2% (*v/v*) residual DMSO). Throughout various experiments, the concentrations of respective stock solutions of reagents were varied. The following gives one example of a detailed synthesis protocol: To 15 µL of AnxV-Npa187 (100 µM, 20 mM Tris buffer, pH 7.8) were added 24 µL aqueous $CaCl_2$ solution (125 mM), 2 µL small molecule (75 mM in DMSO), and 59 µL Tris buffer (20 mM, pH 7.8) to a final volume of 100 µL. Four technical replicates were prepared. The vials were capped and placed into a 96-well Desyre® photoreaction block with holes in the bottom plate to allow for light transmittance. Subsequently, the lid was applied with sufficient pressure to ensure air-tight sealing of the capped glass vials. The block was taken out of the glove box and put into the Zinsser Lumidox® photoreactor. The reaction mixtures were irradiated for a minimum of 15 minutes with blue light (365 nm, 280-305 mW/well) while agitating at 400 rpm with cooling of the reaction block holder (4 °C). An analytical sample was prepared to confirm the conversion to the product with protein-LC-MS. For this, 2 µL of crude reaction mixture were diluted with 48 µL of $H_2O$ containing 0.1% formic acid and analyzed utilizing the Waters XEVO-QTof LC-MS.

For further experiments (MS/MS, photophysical and/or biological measurements) the technical replicates with confirmed conversion to the desired product were combined and purified (desalted) using SEC (PD Minitrap G25, gravity protocol) with the respective reaction buffer as elution buffer.

**Synthesis on modified protein**

***eH3-Tha$_{CF3}$9***

Following general procedure **D** 7-Trifluoromethyl imidazo[1,2-*a*]pyrimidine was used in four technical replicates. After 80 min reaction time the replicates were combined and purified (SEC) resulting in a final concentration of 12 µM purified on-protein product in 1 mL of buffer as determined with the nanodrop photo-spectrometer. $M_{exp.}$= 19508 *m/z* (aromatized product), $M_{obs.}$= 19528 *m/z*.

***eH3-Tha$_{Br}$9***

Following general procedure **D** 6-Bromoimidazo[1,2-*a*]pyrimidine was used in four technical replicates. After 30 min reaction time the replicates were combined and purified (SEC) resulting in a final concentration of 45 µM purified unsubstituted on-protein product in 1 mL of buffer as determined with the nanodrop photospectrometer. $M_{exp.}$= 19518 *m/z* (aromatized product), $M_{obs.}$= 19440 *m/z* (HBr eliminated aromatized product)

***eH3-Tha$_{H}$9***

Following general procedure **D** imidazo[1,2-*a*]pyrimidine was used in four technical replicates. After 15 min reaction time the replicates were combined and purified (SEC) resulting in a final concentration of 82 µM purified on-protein product in 1 mL of buffer as determined with the nanodrop photospectrometer. $M_{exp.}$= 19440 *m/z* (aromatized product), $M_{obs.}$= 19460 *m/z*

***AnxV-Tha$_{H}$187***

Following general procedure **G** imidazo[1,2-*a*]pyrimidine was used in four technical replicates and irradiated at 365 nm with 305 mW/well. After 30 min reaction time and confirming the formation of product, the replicates were combined and purified (SEC) resulting in a final concentration of 2 µM purified on-protein product in 1 mL of buffer as determined with the nanodrop photospectrometer. $M_{exp.}$= 35934 *m/z* (aromatized product), $M_{obs.}$= 35953 *m/z*

#### Biological Methods

##### Construction of Npa-tag *via* amber codon suppression system

*Construction of pEVOL- SS12-TyrRS(Y32L/D158P/I159A/L162Q/A167V)/tRNATyr plasmid*

pEVOL expression vector was generously donated by Prof. P.G. Schultz (Scripps Institute). A dural *glnS'* (constitutive) / *araBAD* (arabinose-inducible) promoter system was constructed to flank the genes of TyrRS. A single $tRNA^{Tyr}$ gene was located between the *proK* promoter and terminator. The mutations on *ss12-tyrRS* (Y12L/D158P/I159A/L162Q/A167V) gene were generated step wisely with site-directed mutagenesis kit. The two identical *ss12-tyrRS* genes were inserted into pEVOL vector between BglII and SalI, or NdeI and PstI sites. Human histone *eH3.1 (C96A/C110A)* gene with a C-terminal FLAG, HA and His-tag was subcloned into pET3d vector using NcoI and BamHI sites. The eH3.1/K9TAG and annexin V/W187TAG mutant were generated by site-directed mutagenesis (QuickChange II site-directed mutagenesis kit, Agilent). A list of primers for generating all constructs is provided below.

*SS12-TyrRS(Y32L/D158P/I159A/L162Q/A167V) mutant*

**DNA sequence of *ss12-tyrRS gene***

ATGGACGAATTTGAAATGATAAAGAGAAACACATCTGAAATTATCAGCGAGGAAGAGTTAAGAGAGGTTTTAAAA
AAAGATGAAAAATCTGCTCTGATAGGTTTTGAACCAAGTGGTAAAATACATTTAGGGCATTATCTCCAAATAAAA
AAGATGATTGATTTACAAAATGCTGGATTTGATATAATTATATTGTTGGCTGATTTACACGCCTATTTAAACCAG
AAAGGAGAGTTGGATGAGATTAGAAAAATAGGAGATTATAACAAAAAAGTTTTTGAAGCAATGGGGTTAAAGGCA
AAATATCTTTATGGAAGTGAGTTCCAGCTTGATAAGGATTATACACTGAATGTCTATAGATTGGCTTTAAAAACT
ACCTTAAAAAGAGCAAGAAGGAGTATGGAACTTATAGCAAGAGAGGATGAAAATCCAAAGGTTGCTGAAGTTATC
TATCCAATAATGCAGGTTAATCCTGCTCATTATCAGGGCGTTGATGTTGTAGTTGGAGGGATGGAGCAGAGAAAA
ATACACATGTTAGCAAGGGAGCTTTTACCAAAAAAGGTTGTTTGTATTCACAACCCTGTCTTAACGGGTTTGGAT
GGAGAAGGAAAGATGAGTTCTTCAAAAGGGAATTTTATAGCTGTTGATGACTCTCCAGAAGAGATTAGGGCTAAG
ATAAAGAAAGCATACTGCCCAGCTGGAGTTGTTGAAGGAAATCCAATAATGGAGATAGCTAAATACTTCCTTGAA
TATCCTTTAACCATAAAAAGGCCAGAAAAATTTGGTGGAGATTTGACAGTTAATAGCTATGAGGAGTTAGAGAGT
TTATTTAAAAATAAGGAATTGCATCCAATGGATTTAAAAAATGCTGTAGCTGAAGAACTTATAAAGATTTTAGAG
CCAATTAGAAAGAGATTATAA

**Protein sequence of SS12-TyrRS**

MDEFEMIKRNTSEIISEEELREVLKKDEKSALIGFEPSGKIHLGHYLQIKKMIDLQNAGFDIIILLADLHAYLNQ
KGELDEIRKIGDYNKKVFEAMGLKAKYLYGSEFQLDKDYTLNVYRLALKTTLKRARRSMELIAREDENPKVAEVI
YPIMQVNPAHYQGVDVVVGGMEQRKIHMLARELLPKKVVCIHNPVLTGLDGEGKMSSSKGNFIAVDDSPEEIRAK
IKKAYCPAGVVEGNPIMEIAKYFLEYPLTIKRPEKFGGDLTVNSYEELESLFKNKELHPMDLKNAVAEELIKILE
PIRKRL

**Oligonucleotides for constructing *ss12-tyrRS*(Y32L/D158P/I159A/L162Q/A167V) mutant gene**

ss12-tyrRS-Y32L-forward: 5′-TAAAAAAAGATGAAAAATCTGCTCTGATAGGTTTTGAACCAAGTGGTA-3′

ss12-tyrRS-Y32L-reverse: 5′- TACCACTTGGTTCAAAACCTATCAGAGCAGATTTTTCATCTTTTTTTA-3′

ss12-tyrRS-D158PI159AL162Q-forward:

5′-TCTATCCAATAATGCAGGTTAATCCTGCTCATTATCAGGGCGTTGATGTTGCAGTTGGA-3′

ss12-tyrRS-D158PI159AL162Q-reverse:

5′-TCCAACTGCAACATCAACGCCCTGATAATGAGCAGGATTAACCTGCATTATTGGATAGA-5′

ss12-tyrRS-A167V-forward: 5′-TCATTATCAGGGCGTTGATGTTGTAGTTGGAGGGATGGAGCAGAGAA-3′

ss12-tyrRS-A167V-reverse: 5′-TTCTCTGCTCCATCCCTCCAACTACAACATCAACGCCCTGATAATGA-3′

*Construction of pEVOL-SS12-TyrRS/tRNA$^{Tyr}$ plasmid*

**tRNA$^{Tyr}$ sequence**

CCGGCGGTAGTTCAGCAGGGCAGAACGGCGGACTCTAAATCCGCATGGCAGGGGTTCAAATCCCCTCCGCCGGAC
CA

**Oligonucleotides for inserting ss12-tyrRS genes into a pEVOL vector between BglII/SalI and NdeI/PstI sites**

TyrRS-BglII-Forward: 5′-AGGAGGAATTAGATCTATGGACGAATTTGAAATGATAAAGA-3′

TyrRS-SalI-Reverse: 5′-GATGATGATGGTCGACTTATAATCTCTTTCTAATTGGCTCTAAAAT-3′

TyrRS-NdeI-Forward: 5′-TGAGGAATCCCATATGGACGAATTTGAAATGATAAAGA-3′

TyrRS-PstI-Reverse: 5′-AGCGTTTGAAACTGCAGTTATAATCTCTTTCTAATTGGCT-3′

**L-3-(2-Naphthyl)alanine (Npa) incorporation into proteins**

The plasmid harboring human eh3/k7tag gene or annexin V/W187tag gene was co-transformed with pEVOL-SS12-TyrRS/tRNA$^{Tyr}$ into *E. coli* BL21(DE3) cells. A 30 ml of overnight culture from a single colony was amplified into a 1 L fresh LB media supplemented with 100 µg/ml Ampicillin (Amp) and 25 µg/ml Chloramphenicol (Cm). Cells were grown at 37 °C till $OD_{600}$ reached 0.4~0.6. Then 1.0 g L-arabinose (0.2%) and 2 mM Npa were added to induce the expression of tRNA synthetase and aminoacylated tRNA. The incubator temperature was reduced to 30 °C. An hour later, cells are induced with 1 mM IPTG and incubated at 30 °C for 16 h. The cell pellets were harvested by centrifugation and stored in -80 °C for later purification.
In order to express wild type annexin V as a control, the plasmid harboring wild type *annexin V* gene was transformed into *E. coli* BL21(DE3) competent cells. A 10 ml of overnight culture was added into 1 L fresh LB media in the presence of 100 µg/ml Amp, when the $OD_{600}$ reached 0.4~0.6, 1 mM IPTG was added, and the incubation was carried out for extra 18 hr with the temperature at 30 °C.

*Histone eH3-Npa9*

**DNA sequence of *eh3-k9tag***

ATGGCACGTACCAAACAGACCGCACGTTAGAGCACCGGTGGTAAAGCACCGCGTAAACAGCTGGCAACCAAAGCA
GCCCGTAAAAGCGCACCGGCAACCGGTGGTGTTAAAAAACCGCATCGTTATCGTCCGGGTACAGTTGCACTGCGT
GAAATTCGTCGTTATCAGAAAAGTACCGAACTGCTGATTCGTAAACTGCCGTTTCAGCGTCTGGTTCGTGAAATT
GCACAGGATTTCAAAACCGATCTGCGTTTTCAGAGCAGCGCAGTTATGGCACTGCAAGAAGCAGCAGAAGCATAT
CTGGTTGGCCTGTTTGAAGATACCAATCTGGCAGCAATTCATGCAAAACGTGTTACCATTATGCCGAAAGATATT
CAGCTGGCACGTCGTATTCGTGGTGAACGTGCCGGTGGTGATTATAAAGATGATGATGATAAAAGTGCAGCCGGT
GGTTATCCGTATGATGTTCCGGATTATGCCCATCATCATCATCATCATCATCATCATCATCACTAATAA

**Protein sequence of eH3-K9TAG**

ARTKQTARXSTGGKAPRKQLATKAARKSAPATGGVKKPHRYRPGTVALREIRRYQKSTELLIRKLPFQRLVREIA
QDFKTDLRFQSSAVMALQEAAEAYLVGLFEDTNLAAIHAKRVTIMPKDIQLARRIRGERAGGDYKDDDDKSAAGG
YPYDVPDYAHHHHHHHHHHH

* The amber stop codon site is highlighted in red

**Oligonucleotides for replacing K9 into amber stop codon *tag***

eH3-K9TAG-Forward: 5′-GCCCGTACCAAGCAGACCGCCCGTTAGTCCACCGGAGGGAAGGCTCCCCGC-3′

eH3-K9TAG-Reverse: 5′-GCGGGGAGCCTTCCCTCCGGTGGACTAACGGGCGGTCTGCTTGGTACGGGC-3′

*Annexin V-Npa187*

**DNA sequence of *annexin V-w187tag***

ATGGCACAGGTTCTCAGAGGCACTGTGACTGACTTCCCTGGATTTGATGAGCGGGCTGATGCAGAAACTCTTCGG
AAGGCTATGAAAGGCTTGGGCACAGATGAGGAGAGCATCCTGACTCTGTTGACATCCCGAAGTAATGCTCAGCGC
CAGGAAATCTCTGCAGCTTTTAAGACTCTGTTTGGCAGGGATCTTCTGGATGACCTGAAATCAGAACTAACTGGA
AAATTTGAAAAATTAATTGTGGCTCTGATGAAACCCTCTCGGCTTTATGATGCTTATGAACTGAAACATGCCTTG
AAGGGAGCTGGAACAAATGAAAAAGTACTGACAGAAATTATTGCTTCAAGGACACCTGAAGAACTGAGAGCCATC
AAACAAGTTTATGAAGAAGAATATGGCTCAAGCCTGGAAGATGACGTGGTGGGGGACACTTCAGGGTACTACCAG
CGGATGTTGGTGGTTCTCCTTCAGGCTAACAGAGACCCTGATGCTGGAATTGATGAAGCTCAAGTTGAACAAGAT
GCTCAGGCTTTATTTCAGGCTGGAGAACTTAAATAGGGGACAGATGAAGAAAAGTTTATCACCATCTTTGGAACA
CGAAGTGTGTCTCATTTGAGAAAGGTGTTTGACAAGTACATGACTATATCAGGATTTCAAATTGAGGAAACCATT
GACCGCGAGACTTCTGGCAATTTAGAGCAACTACTCCTTGCTGTTGTGAAATCTATTCGAAGTATACCTGCCTAC
CTTGCAGAGACCCTCTATTATGCTATGAAGGGAGCTGGGACAGATGATCATACCCTCATCAGAGTCATGGTTTCC
AGGAGTGAGATTGATCTGTTTAACATCAGGAAGGAGTTTAGGAAGAATTTTGCCACCTCTCTTTATTCCATGATT
AAGGGAGATACATCTGGGGACTATAAGAAAGCTCTTCTGCTGCTCTGTGGAGAAGATGACTAA

**Protein sequence of Annexin V-W187TAG**

MAQVLRGTVTDFPGFDERADAETLRKAMKGLGTDEESILTLLTSRSNAQRQEISAAFKTLFGRDLLDDLKSELTG
KFEKLIVALMKPSRLYDAYELKHALKGAGTNEKVLTEIIASRTPEELRAIKQVYEEEYGSSLEDDVVGDTSGYYQ
RMLVVLLQANRDPDAGIDEAQVEQDAQALFQAGELKXGTDEEKFITIFGTRSVSHLRKVFDKYMTISGFQIEETI
DRETSGNLEQLLLAVVKSIRSIPAYLAETLYYAMKGAGTDDHTLIRVMVSRSEIDLFNIRKEFRKNFATSLYSMI
KGDTSGDYKKALLLLCGEDD

* the amber stop codon at position Trp 187 is highlighted in red.

**Oligonucleotides for introducing amber codon TAG at position 187**

ANV187TAG-Forward: 5′-ATTTCAGGCTGGAGAACTTAAATAGGGGACAGATGAAGAAAAGTTTA-3′
ANV187TAG-Reverse: 5′-TAAACTTTTCTTCATCTGTCCCCTATTTAAGTTCTCCAGCCTGAAAT-3′

#### Protein purification

For histone eH3/K9Npa protein, cell pellets were re-suspended and lysed in 20 ml of wash buffer (50 mM Tris, 100 mM NaCl, pH 7.5) with a cOmplete™ protease inhibitor and 1 mg DNase. The mixture was sonicated using a microtip and centrifuged at 20,000 rpm for 20 mins at 4 °C. The supernatant was discarded, and the pellets were resuspended, which were washed once with 20 ml of Tris/1% Triton X-100 and another 20 ml of Tris buffer. The pellets were resuspended with 1 ml 100% DMSO at room temperature for 10 min. A 10 ml unfolding buffer (7 M guanidinium chloride, 20 mM Tris, pH 7.5) was added and the mixture was shaken vigorously at room temperature for 1 hr. Following centrifugation at 20,000 rpm for 20 mins, the supernatant was incubated with Ni-NTA resin, which was pre-equilibrated with a Tris/urea buffer. The impurities and endogenous proteins were washed with Tris/urea buffer (50 mM Tris, 500 mM NaCl, 7 M urea, pH 7.5) with increased amounts of imidazole (20-60 mM). The final protein was eluted with 150 to 500 mM imidazole in Tris/urea buffer and identified on SDS-PAGE. Purified fractions were dialyzed against water (with 2 mM β-mercapto-ethanol) and lyophilized.

For annexin V proteins, cell pellets were suspended in a 30 ml $CaCl_2$ buffer (50 mM Tris, 10 mM $CaCl_2$, pH 7.2) with a tablet of cOmplete™ protease inhibitor cocktail and 1 mg DNaseI. The mixture was sonicated using a microtip and centrifuged at 20,000 rpm for 15 mins at 4 °C. The supernatant was discarded. The cell pellets were re-suspended in 40 ml EDTA buffer (50 mM Tris, 20 mM EDTA, pH 7.2) and the cell debris was removed by centrifugation (20, 000 rpm, 15 mins, 4 °C). The supernatant containing annexin V was dialyzed against 4L Tris buffer (20 mM Tris, pH 7.8) for 3 times at 4 °C. The dialyzed protein solution was loaded onto a pre-equilibrated 5 ml HiTrap Q HP anion exchange chromatography column. The column was applied to an ÄKTA FPLC system (GE Healthcare) using solvent A (20 mM Tris, pH 7.8) and solvent B (20 mM Tris, 500 mM NaCl, pH 7.8). The purified fraction was identified by SDS-PAGE and concentrated with Vivaspin centrifugal concentrator (10 kDa cutoff, Cytiva™) to 1 ml. Further purification was performed on a pre-equilibrated Superdex 75 16/600 size-exclusion column with a Tris buffer (20 mM Tris, pH 7.8). The successful incorporation of Npa in annexin V protein was confirmed by LC-MS spectra.

**Labelling of annexin Vs with Alexa Fluor 488 NHS ester dye**

Annexin V proteins were desalted into 1X PBS buffer via PD SpinTrap G25 column (Cytiva™ life sciences) for reaction with NHS ester. To synthesize annexin V-AF488 with only one fluorophore conjugate, 1 molar equivalent of AF488-NHS ester (1.6 µl of 2.5 mM stock in DMSO) was incubated with 40 µM Annexin V solutions at room temperature for 1 h to give a final volume of 100 µl. The reaction was monitored by LC-MS spectra. The percentage of conjugation was determined based on the intensity of deconvoluted peaks. The unreacted ester dye in each sample was removed by PD SpinTrap™ G-25 desalting column.

**Fluorescence plate assay for the determination of AnxV binding affinities to phosphatidylserine (PS)**

To compare the binding affinities of each annexin V to PS, the plate assay was tested. In general, PS (POPS16:0-18:1, 1-palmitoyl-2-oleoyl-sn-glycero-3-phopho-L-serine, Avanti Polar Lipids) was immobilized on a 96-well F-bottom microplate (µCLEAR®, black, Greiner BIO-ONE) after solubilization in methanol at a concentration of 1.0 mg/ml. Each well was filled with 200 µl of PS solution. After methanol evaporation at room temperature, a film of PS was formed at the bottom of each well. A 50 µl of AF488-conjugated annexin V with serial dilutions (ranging from 500 nM to 1 nM) was added to the PS-coated wells and incubated for 1 h at room temperature in the dark. Triplicate experiments were performed in the Tris buffer (20 mM Tris, pH 7.8) with or without 5 mM $CaCl_2$. The plates were washed three times with 50 µl Tris buffer. Bound annexin V proteins were detected by a fluorescence plate reader (Ex, 488-14 nm; Em, 535-30 nm; CLARIOstar$^{Plus}$, BMG Labtech) and CLARIOstar-Data Analysis software was used for data collection. The quantity of bound AF488-annexin V proteins on PS layer obtained from the relative fluorescent units (RFU) was plotted against the logarithmic concentration of total AF488-annexin V proteins. The curves were fitted according to a sigmoidal dose-response profile using Origin 2022. The protein concentration at half-saturation corresponds to the dissociation constant ($K_d$).

**Analysis of apoptosis cells stained with AF488-conjugated annexin V proteins**

*Confocal microscopy:* Confocal microscopy analysis was performed to determine the cellular localization of each of the AF488-conjugated annexin V proteins. Hela cells ($50x10^3$ cells) were seeded into 8-well chamber (300 µl/well, µ-sldie 8 well ibiTreat, Ibidi) and incubated at 37 ºC for 24 hrs in an incubator humidified with 5% $CO_2$. Cells were washed once with 1X PBS, and then treated with 0.3 µg (1 µg/ml/$10^6$ cell) actinomycin D at 37 ºC for 30 mins to induce cell death. Cells were washed twice with an annexin binding buffer (10 mM HEPES, 140 mM NaCl, 5 mM $CaCl_2$, pH 7.4), and then added with AF488-conjugated annexin V proteins (0.35 µM/well). After incubation at room temperature in the dark for 15 mins, unbounded annexin Vs were washed away with binding buffer. Cells were analyzed using a Leica SP8 confocal microscope (63X/1.40 oil, Argon laser 488).
*Flow cytometry:* Hela cells ($6X10^6$ in a T75 flask) were incubated with actinomycin D (1 µg/ml in 5 ml 1X PBS) at 37 ºC for 30 mins. After 2 washes with 10 ml 1X PBS, cells were detached from T75 flask with 5 ml 1XPBS (containing 10 mM EDTA) by centrifugation at 300 g for 5 mins and resuspended in a 2 ml annexin binding buffer (10 mM HEPES, 140 mM NaCl, 5 mM $CaCl_2$, pH 7.4). The cells were aliquoted into 1.5 ml Eppendorf tubes (200 µl/tube). A 4 µM AF-conjugated annexin Vs was added to each tube and incubated at room temperature for 15 mins in the dark. Unbound annexin V was removed by centrifugation. The treated cells were resuspended with FACS buffer (annexin binding buffer with 1% BSA) containing 5 µl of propidium iodide staining solution (eBioscience™ propidium idode, ThermoFisher Scientific) and analyzed by flow cytometry (CytoFLEX LX, BECKMAN COULTER). The data was collected with software CytExpert 2.4 and processed by FlowJo v10.9.0. Hela cells untreated with apoptosis reagent actinomycin D and without annexin V staining as control. For the cells incubated with annexin Vs-AF488, the FSC-SSC (forward scatter-side scatter) plots were gated according to untreated cells and percentages felt into four regions were calculated. Annexin V-positive cells are denoted as early apoptotic, PI-positive cells are specified as necrotic, while positive cells for both staining indicate late apoptosis in a population.

#### Spectroscopy and spectrometry

##### Fluorescence measurements

All the measurements were performed on Cary Eclipse Fluorescence Spectrophotometer (Agilent Technologies) using a fluorimeter cuvette with path length of 10 mm. For different measurements, excitation and emission slits are varied. During experiments for the comparison of emission intensities, excitation and emission slits are kept constant.

##### Tandem mass spectrometry

***Protein digestion for MSMS analysis:*** 5-10 ug of each protein were digested using a modified single-pot solid phase assisted sample preparation method (SP3) using 200 ug magnetic carboxyl coated magnetic beads (Cytiva). Proteins in Luche buffer were precipitated directly, samples in Tris buffer were supplemented with sodium dodecyl-sulfate (2% final concentration). Proteins were precipitated onto the beads using acetonitrile (80% final concentration) and incubated for 30 min with shaking. Once the proteins were immobilised, the beads were washed three times with 80% Ethanol, followed by three washes with 100% acetonitrile before incubation with digestion buffer (50 mM TEAB, containing Trypsin Gold (Promega) at 1:25 enzyme-to-protein for four hours. Supernatant was collected and beads were washed with 2% DMSO solution and combined with supernatant before acidification with formic acid (5% final concentration) and centrifugation at 16,000 x g for 10 minutes to remove any undigested enzyme. Peptides were desalted on Oasis HLB cartridges prior to analysis.

***LC/MS:*** Analysis of peptides was carried out using an Ultimate 3000 nano-LC 1000 system coupled to an Orbitrap Ascend (Thermo Fisher Scientific). Supernatant from SP3 beads was diluted 10x in ultra-pure water with 5% formic acid and 5% DMSO. Peptides were initially trapped on a C18 PepMap100 pre-column (300 µm inner diameter x 5 mm, 100A) and then separated on an in-house constructed C18 column (Reprosil-Gold, Dr. Maisch, 1.9 µm particle size) column (ID: 50 µm, length: 50 cm) at a flow rate of 100 nL/min. Peptides were separated over 15 min (12-38%B) using mobile phase A (water and 0.1% formic acid) and mobile phase B (acetonitrile and 0.1% formic acid). Separated peptides were directly electrosprayed into an Orbitrap Ascend mass spectrometer (Thermo Fisher Scientific). Mass spectra were acquired in the orbitrap (350-1400 m/z, resolution 60000, AGC target 3 x 106, maximum injection time 50 ms) in a data-dependent mode. The top 40 most abundant peaks in the survey scan were fragmented using CID (resolution 7500, AGC target 4 x104, maximum injection time 64 ms).

Peptide identification and quantification were performed using Andromeda search engine implemented in MaxQuant (2.3.0.0)[2,3]. Peptides were searched against an *E.coli* reference database (Uniprot, downloaded 2022.09.07) with additional sequence of recombinant H3 or Annexin V with mutated residues marked as C. Default settings were used apart from the following additional variable modifications: Carbamidomethylation (+57.0215), Npa incorporation, as well as target modifications at Cys were used as variable modifications; no fixed modifications were applied. Exact mass differences used for the search for each modification are listed in Table SS1.

***Table SS1.*** *Exact mass differences used for the search to confirm Npa incorporation or successful on-protein chemistry*

| **Protein** | **Modification** | **Mass** |
|---|---|---|
| eH3 | Cys -> Npa | 94.0749 |
| | Cys -> $Tha_{CF3}9$ | 299.1212 |
| | Cys -> $Tha_{H}9$ | 231.1338 |
| | Cys -> $Tha_{Br}9$ | 211.1076 |
| AnxV | Trp -> Npa | 11.0048 |
| | Trp -> $Tha_{H}187$ | 148.0637 |

##### *eH3 histone sequence:*

ARTKQTARCSTGGKAPRKQLATKAARKSAPATGGVKKPHRYRPGTVALREIRRYQKSTEL
LIRKLPFQRLVREIAQDFKTDLRFQSSAVMALQEAAEAYLVGLFEDTNLCAIFAKRVTIM
PKDIQLARRIRGERAGGDYKDDDDKSAAGGYPYDVPDYAHHHHHHHHHH

##### *Annexin V sequence:*

MAQVLRGTVTDFPGFDERADAETLRKAMKGLGTDEESILTLLTSRSNAQRQEISAAFKTLF
GRDLLDDLKSELTGKFEKLIVALMKPSRLYDAYELKHALKGAGTNEKVLTEIIASRTPEE
LRAIKQVYEEEYGSSLEDDVVGDTSGYYQRMLVVLLQANRDPDAGIDEAQVEQDAQALFQ
AGELKCGTDEEKFITIFGTRSVSHLRKVFDKYMTISGFQIEETIDRETSGNLEQLLLAVV
KSIRSIPAYLAETLYYAMKGAGTDDHTLIRVMVSRSEIDLFNIRKEFRKNFATSLYSMIK
GDTSGDYKKALLLLCGEDD
